# Monolingual and bilingual infants rely on the same brain networks: Evidence from resting-state functional connectivity

**DOI:** 10.1101/2020.04.10.035469

**Authors:** Borja Blanco, Monika Molnar, Manuel Carreiras, Liam H. Collins-Jones, Ernesto Vidal, Robert J. Cooper, César Caballero-Gaudes

**Author notes:** Correspondence should be addressed to Borja Blanco.

## Abstract

This study examines whether bilingual exposure has a profound effect on the functional organization of the developing human brain during infancy. Recent behavioural research attests that monolingual vs. bilingual experience affects cognitive and linguistic processes already during the first months of life. However, to what extent the intrinsic organization of the infant human brain adapts to monolingual vs. bilingual environments is unclear. We measured spontaneous hemodynamic brain activity using functional near-infrared spectroscopy (fNIRS) in a large cohort (N=99) of 4-month-old monolingual and bilingual infants. We implemented well-established analysis approaches of functional brain imaging that enabled us to reveal the functional organization of the infant brain in large-scale cortical networks, and to perform group-level comparisons (i.e., monolingual vs. bilingual groups) in a reliable manner. Our results revealed no differences between the intrinsic functional organization of the developing monolingual and bilingual infant brain at 4 months of age.

## 1. Introduction

One way to understand the intrinsic functional organization of the human brain is through the measurement of resting-state functional connectivity (RSFC). RSFC reflects spontaneous but synchronized fluctuations in cerebral hemodynamic activity between brain regions that share a common role in supporting various, functionally relevant sensory and cognitive processes (Fox and Raichle, 2007; Damoiseaux et al., 2006). RSFC can be measured in infants, children, and adults, thus providing a window into neural specialization across the life span (Gao et al., 2017). The intrinsic functional organization of the infant brain described by RSFC might show changes depending on various pre- and postnatal factors. As measured by functional magnetic resonance imaging (fMRI) studies, premature and full-term infants show different RSFC patterns (Damaraju et al., 2010; Smyser et al., 2010); moreover, the configuration and maturational course of functional connectivity differs across neurotypical infants and toddlers in comparison to those of high risk of developmental disorders, such as autism spectrum disorder (Dinstein et al., 2011; Keehn et al., 2013). Relevant to the current study, RSFC also reflects differences in neural adaptation associated with environmental factors, such as the caregivers’ education level or socioeconomic status (Gao et al., 2015).

A specific external factor is growing up in a bilingual environment. Language acquisition begins as soon as infants are able to hear spoken language, about 3 months prior to birth (e.g., Werker, 2018). In a bilingual learning environment, infants are exposed to the linguistic regularities (e.g., speech sounds, words, grammar) of not one, but two inputs simultaneously. Therefore, the linguistic statistical regularities present in a bilingual environment are different than in a monolingual one. In addition, differently from monolinguals, bilinguals need to discriminate between two languages. While bilingual infants’ overall language exposure should be comparable to that of monolinguals, bilingual infants likely receive less exposure to each of their languages –compared to their monolingual peers– because their exposure time is split between two inputs. A bilingual environment has been shown to have consequences on infants’ brain responses and behaviour when cognitive functions and spoken language processing are considered. Dissimilar patterns of brain activation have been observed in monolingual and bilingual 4-month-olds in a language discrimination/recognition task (Nacar-García et al., 2018) and when infants are presented with native and non-native speech sounds (García-Sierra et al., 2011; Petitto et al., 2012; Ferjan Ramírez et al., 2017) or words (Conboy and Mills, 2006). Importantly, these differences have been observed across different imaging modalities such as electro- and magnetoencephalography, and functional near-infrared spectroscopy (fNIRS). Bilingualism has an impact at the behavioural level as well, for instance when attention allocation to languages are considered: bilingual 4-month-olds orient slower to their languages (Bosch and Sebastián-Gallés 1997; Sebastián-Gallés et al., 2012) and they exhibit longer sustained attention periods than monolinguals when processing spoken language (Molnar et al., 2014). It has been also proposed that bilingual infants possess increased executive control abilities (Kovacs and Mehler 2009; Byalistok, 2015). These brain and behavioural differences across monolingual and bilingual infants have been conceptualized as different types of adaptation patterns to monolingual vs. bilingual environments. From the perspective of bilingual infants, this adaptation might facilitate the acquisition of two languages as opposed to one.

Does growing up in a bilingual environment have consequences when RSFC is considered? This question is relevant, given that differences between monolingual and bilingual infants neural responses have been only observed in the presence of explicit cognitive and/or linguistic tasks. Whether bilingualism has a more profound effect on the developing human brain, such as its functional organization, is currently unknown. Measuring RSFC in monolingual and bilingual infants can reflect whether bilingual experience during the first months of life leads to specific adaptations in the intrinsic properties of brain’s function that are observable in the absence of any task or stimuli (i.e., at rest). Testing whether functional adaptation to a bilingual context is evident at the earliest stages of human development is crucial for our understanding of how bilingualism interacts with general brain maturation patterns beyond task-specific language and cognitive processing. There is evidence that long-term exposure to two languages might alter the brain’s functional (Parker Jones et al., 2011; Berken et al., 2016) and structural organization (Mechelli et al., 2004; García-Pentón et al., 2014; Mohades et al., 2015), as demonstrated by MRI studies in adults. For instance, stronger functional connectivity in bilingual adults as compared to monolinguals has been observed in long-range bilateral and anterior-posterior connections on both hemispheres (Luk et al., 2011), and in brain networks associated with language and executive control processes (Grady et al., 2015; Berken et al., 2016). The aim of the current study is to assess whether the brain’s functional architecture begins to adapt to a bilingual environment as early as 4 months of age, by the time neural and behavioural responses to external stimuli often differ across monolinguals and bilinguals. Specifically, the current goal is to contrast the RSFC across monolingual and bilingual 4-month-old infants using functional near-infrared spectroscopy (fNIRS).

Due to its practical advantages, fNIRS has emerged as an alternative to fMRI to characterize RSFC in infant populations (Obrig et al., 2000). Despite the visual (White et al., 2009; Mesquita et al., 2010), sensorimotor (Lu et a., 2010; Mesquita et al., 2010; Zhang H. et al., 2010) and auditory/language networks (Lu et a., 2010) have already been identified in adults using multichannel fNIRS, the number of RSFC studies in infants is scarce. Using a whole-cortex fNIRS setup Homae et al., (2010) described changes in cortical network organization from birth to 6 months of age. They observed that functional connectivity was constrained to spatially adjacent cortical regions in neonates, with this pattern progressively evolving towards increased interhemispheric connectivity between homologous brain regions during the first months of life. The same authors also demonstrated a modulation in RSFC patterns induced by task execution (Homae et al., 2011). They measured RSFC in 3-month old infants before and after presenting them with speech sounds, and showed an increase in fronto-temporal connectivity after stimuli presentation. Watanabe et al. (2017) and Taga et al. (2000; 2017) investigated the associative properties of resting-state oxyhemoglobin (HbO) and deoxyhemoglobin (HbR) time series as potential biomarkers for vascular and metabolic function in the infant brain. These studies shown that the phase relationship between HbO and HbR time series shifts from an in-phase state to an antiphase state during the first months of life (Taga et al., 2017), and that the developmental trajectory of this shift might be delayed in infants born preterm (Watanabe et al., 2017). Other works have further demonstrated the potential of fNIRS-RSFC to investigate typical and atypical functional brain development. Besides assessing the effect of premature birth in functional connectivity development (White et al., 2012; Fuchino et al., 2013; Watanabe et al., 2017), functional connectivity studies using fNIRS have also revealed alterations in the strength and synchronization of spontaneous HbO and HbR fluctuations in infants with Down’s syndrome (Imai et al., 2014). Moreover, there has been evidence of a gradual increase of fronto-temporoparietal connectivity in resting state between 11 and 36 months of age as a prospective precursor of the developing default-mode network (Bulgarelli et al., 2020), which might be also related to the ability of self-recognition at 18 months of age (Bulgarelli et al., 2019). High-density diffuse optical tomography systems, which employ the same principles as fNIRS, have also been used to obtain RSFC maps of the occipital and temporal cortices in newborns at the bedside (White et al., 2012; Ferradal et al., 2015), showing large spatial agreement with the patterns obtained using fMRI (Ferradal et al., 2015).

Some of these previous functional connectivity studies with fNIRS identified resting-state networks (RSNs) at the single-subject level (White et al., 2009; Lu et a., 2010; Mesquita et al., 2010; White et al., 2012; Novi et al., 2016). However, when attempting to describe RSFC at the group-level, and to quantitatively compare the observed RSFC patterns across different experimental conditions or groups, established methods of fNIRS data analysis present some limitations. For example, fNIRS spatial resolution is low, and a channel localization method that ensures consistency for seed-based correlation analysis is often not available. Thus, the brain region under investigation may vary across individuals, reducing statistical power. On the other hand, group-level functional connectivity studies based on independent component analysis (ICA) have often been computed by averaging those subject-specific components that match an a priori defined spatial configuration (e.g., bilateral and covering sensorimotor regions). However, individual data are usually affected by noise components of different characteristics, which might result in an ICA separation that differs across subjects. The ultimate consequence of these limitations is that most previous fNIRS studies have evaluated group differences in RSFC using qualitative comparisons (Homae et al., 2010; White et al., 2012), or performing statistical analysis on specific connectivity indexes only (Homae et al., 2010; Imai et al., 2014; Watanabe et al., 2017).

Consequently, an additional goal of this work is to overcome the aforementioned limitations of current group-level RSFC analysis methods with fNIRS, i.e. accurately describing RSFC at the group level and quantitatively comparing RSFC between experimental conditions/groups. For this, we implemented two data-driven methodologies based on ICA, which have been widely employed in resting-state fMRI research, to extract group-level large-scale functional connectivity patterns in our population of infants. These methods also provide a means to quantitatively compare the strength/prominence of these networks across our experimental groups. First, we used temporal group ICA (tGICA) with dual regression to compute temporally independent patterns of spontaneous hemodynamic activity (Beckmann et al., 2009; Smith et al., 2012). By concatenating the fNIRS channel time courses of multiple subjects, tGICA generates a set of group-level maximally independent temporal time courses and its common aggregated spatial maps (i.e., functional networks, FN), which quantify the presence of each particular independent component on each specific channel. Group-level spatial maps, which spatially represent the FN of interest, can be regressed out to the subject level to obtain subject-specific spatial maps using spatio-temporal or dual regression. Between group differences can then be assessed by performing statistical analyses across subject-specific maps on a channel by channel basis. As a complementary analysis, we also implemented connICA (Amico et al., 2017), a connectome-based ICA approach in which the individual functional connectivity matrices or connectomes are jointly decomposed to obtain latent group-level independent functional connectome components (FCC) and its associated weights that quantify the relative prominence of each FCC on each subject. Besides describing cortical network organization at the group level, the two methods employed in this study allow quantitatively investigating the link between particular clinical, cognitive or external variables (i.e., early bilingualism in the current work), and the relative presence of the extracted functional connectivity components on each experimental group under assessment.

In this study we measured spontaneous hemodynamic brain activity using fNIRS in a large cohort of 4-months old infants to study the effect of bilingual language acquisition on different functional brain systems, while avoiding potential confounds due to task interference. Assessing RSFC with fNIRS in developmental populations face challenges such as high attrition rate (i.e., low sample size), the difficulty to perform long recordings and data quality. In order to reduce the impact of these issues, we measured spontaneous hemodynamic brain activity in a large sample of infants during natural sleep and quality assurance methods were implemented during data preprocessing. With this procedure we were able to collect a sample of 99 infants with good data quality and continuous recordings with at least 9 minutes duration. We examined the configuration of group-level RSFC patterns using two data-driven methodologies based on ICA: tGICA and connICA. The presence of the identified functional networks and functional connectome components was quantified in each participant, and results were compared across two monolingual groups of infants—Spanish and Basque—and one bilingual group of infants—Spanish-Basque—in order to determine if simultaneously acquiring two languages from birth modulates functional network development at this early age.

## 2. Methods

### 2.1. Participants

123 healthy full-term infants participated in this study. In sixteen of these participants testing was not conducted because infants were not able to fall sleep. One participant was discarded for receiving a regular exposure to English. Two infants were excluded before data preprocessing because their recorded datasets were shorter than 600 seconds. Five infants (n = 2 Basque-Spanish bilingual infants, n = 1 Spanish monolingual infant and n = 2 Basque monolingual infants) were excluded during data preprocessing due to insufficient data quality. In the final sample — for which data was analyzed and results are presented — 99 participants were included: 36 Basque-Spanish bilingual (BIL) infants (21 girls; mean age = 125 ± 4 days), 30 Spanish (SP) monolingual infants (13 girls; mean age = 123 ± 3 days) and 33 Basque (BQ) monolingual infants (17 girls; mean age = 122 ± 4 days). Participants’ language background was assessed with a questionnaire filled by the parents, in which infants’ percentage of exposure to each language (SP and BQ) during the first months of life was measured. Participants exposed to a single language (SP or BQ), or less than 10% of the time to a second language (SP or BQ), were included in each of the monolingual groups. Infants raised in a Spanish-Basque bilingual environment, those that were exposed to their two native languages from birth, formed the bilingual group. Participants were recruited from the same region of the Basque Country (Gipuzkoa); a socioeconomic status questionnaire was completed to ensure that families showed similar levels of education, parental occupation and household income across groups. Parents were informed about the procedure of the study and signed a written informed consent before the experiment. The local ethical committee approved this study.

### 2.2. Data acquisition

Functional Near Infrared Spectroscopy (fNIRS) measurements were performed with a NIRScout system (NIRx Medical Technologies, CA, USA) at wavelengths 760 and 850 nm with a sampling rate of 8.93 Hz. fNIRS provides an indirect measure of local neural activity by measuring concentration changes of oxyhemoglobin (HbO) and deoxyhemoglobin (HbR) in the cerebral and extracerebral vascular tissue underlying each measurement channel (Obrig and Villringer, 2003). Sixteen light emitters and 24 detectors were positioned on a stretchy fabric cap (Easycap GmbH, Germany) over frontal, temporal, parietal and occipital regions of both hemispheres according to the international 10-20 system (Fig. 1a). Nasion, inion and preauricular points were used as external head landmarks, and caps of two different sizes (i.e., 40 and 42) were employed to adapt to individual head perimeter/size. Each pair of an adjacent light emitter and a detector formed a single measurement channel, which generated 52 channels for each hemoglobin oxygenation state (i.e., HbO and HbR). Occipital channels were discarded in all participants for being particularly prone to contain signal artifacts, since during data acquisition the back part of the infants’ head was leaning against the parent’s body, and any minor movement resulted in the misplacement of these particular optodes. Thus, only data from the remaining 14 sources and 19 detectors (i.e., 46 channels) was analyzed.

**Fig. 1.**
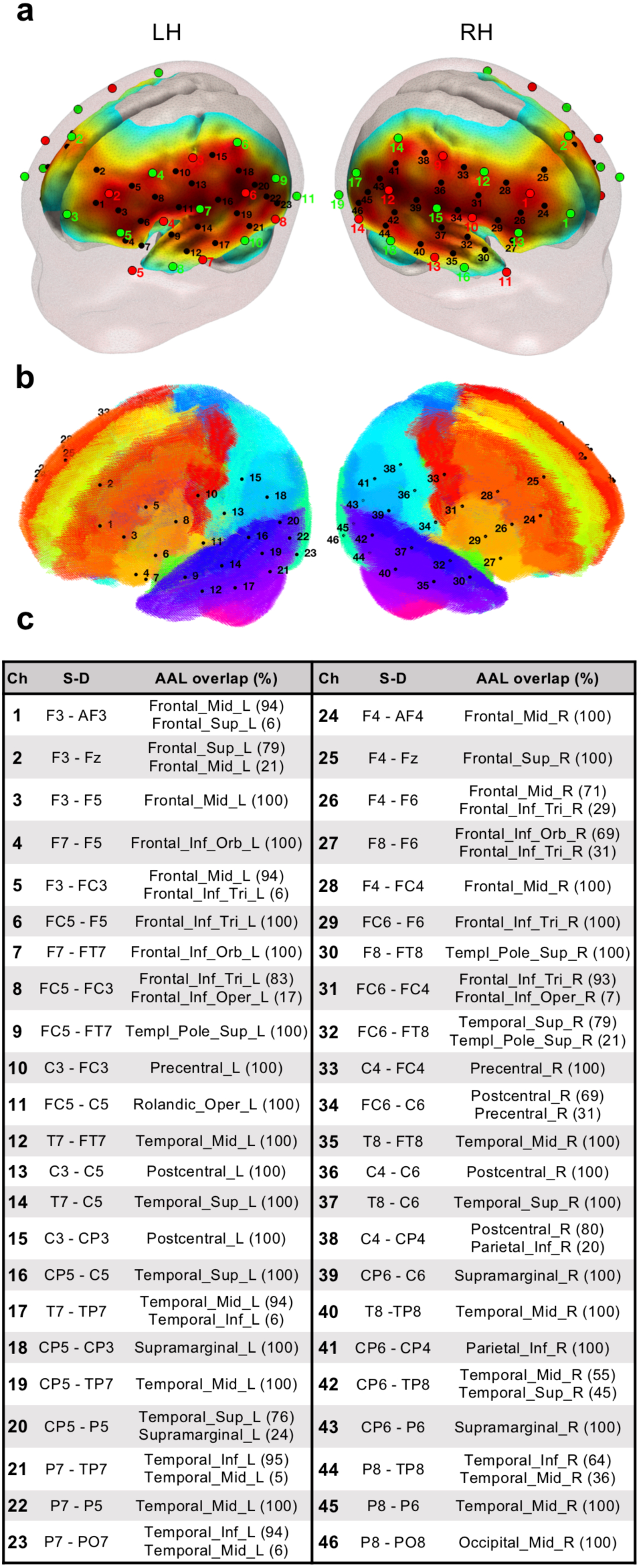
**a)** fNIRS optode (sources in red, detectors in green) and channel (black) localization in the current experimental setup. The normalized sensitivity profile of this configuration is displayed in a 6-month-old infant head model. **b)** Localization of the fNIRS channels in our setup registered to a 6-month-old infant AAL template. **c)** Table depicting the brain labels of our setup based on the probabilistic spatial registration of the fNIRS channels to a 6-month-old infant AAL template. Ch = Channel; S-D = Source-Detector pair.

The sensitivity profile of the fNIRS probe setup was computed to provide information of the brain areas under investigation—and for results visualization purposes. The probe setup was registered to an average 6-month-old infant template (Richards et al., 2016) to compute the sensitivity matrix of our source-detector configuration using Toast++ (Schweiger and Arridge, 2014). We obtained the aggregated sensitivity profile of our probe by summing the normalized cortical sensitivity profiles of each individual channel (Fig. 1a). Channel positions were defined as the grey matter node which coordinates were closest to the central point of the maximum sensitivity path along each source-detector pair. Next, a 6-month-old average atlas (Akiyama et al., 2013) was used to compute a probabilistic spatial registration of the cortical structures underlying each channel. Channel coordinates were first transformed to the Akiyama et al., (2013) average T1 template space using Advanced Normalization Tools (ANTs) (Avants et al., 2009), and then registered into the Akiyama et al., (2013) anatomic atlas, defined by 116 cortical regions based on Automated Anatomical Labeling (AAL) (Tzourio-Mazoyer et al., 2002). Finally, for each channel the AAL anatomical labels within a distance of 20 mm were defined, and the percentage of overlap with each AAL region was calculated (see Fig. 1b-c).

### 2.3. Experimental procedure

We measured infants’ spontaneous hemodynamic activity during natural sleep while leaning on their parents’ lap in a sound attenuated room. The only source of illumination in the room was the screen of the recording computer, which was attenuated to low brightness levels. Recordings started when infants were relaxed, accustomed to the fNIRS cap and clear signs of sleep were noticeable. Over the duration of the recordings, parents were asked to remain silent and to minimize movements in order to avoid involuntary cap or optode displacement. Recordings lasted between 10 and 25 minutes unless the infant woke up during the experiment. Recordings were interrupted if infants showed continued and excessive movement or signs of discomfort at any point.

### 2.3. Data preprocessing and quality assessment

All data preprocessing and analyses were performed in Matlab (R2012b and R2014b, Mathworks, MA, USA) using in-house scripts, as well as third-party toolboxes and algorithms. Optical density changes were calculated from raw light intensity data using the *hmrIntensity2OD* algorithm implemented in Homer2 (Huppert et al., 2009). Noisy periods at the beginning or/and at the end of the recordings, presumably matching awake activity of the infants (i.e., before the infant was completely asleep and after the infant woke up) were visually identified and removed. As participants were asleep during the acquisition, recordings displayed good data quality. However, some datasets showed brief, sparse motion-induced artefacts characterized by abrupt amplitude signal changes and/or artefactual signal drifts. These artefacts were corrected using the wavelet-based method described in Patel et al., (2014) which was adapted for fNIRS data. Optical density data was then converted into HbO and HbR concentration changes using the *hmrOD2Conc* function in Homer2, with differential pathlength factors of 5.3 and 4.2 (Scholkmann and Wolf, 2013). After this step all datasets were limited to 5000 samples (i.e., ~560 seconds) to ensure homogeneous effect estimate precision in the first level of the analysis (i.e., robust Pearson correlation coefficient). This step was performed by visually identifying the segment of the dataset displaying the best data quality. Temporal filtering and global signal regression were performed simultaneously in a unique nuisance regression model (Caballero-Gaudes and Reynolds, 2017). In detail, contribution of high-frequency physiological noise sources (e.g., respiration and cardiac pulsation) was accounted for by including Fourier terms for frequencies above 0.09 Hz in the model. Very slow frequency fluctuations and signal drifts were modelled by adding the first 4 order Legendre polynomials to the model. The average fNIRS signal across all channels was also included in the regression model to remove globally occurring hemodynamic processes in cerebral and extracerebral tissues assumed to largely reflect systemic hemodynamic changes (White et al., 2009; Mesquita et al., 2010). As HbO and HbR are differently affected by global systemic processes, data of each hemoglobin chromophore were filtered independently by including in the model either the global HbO or HbR signal (Tachtsidis and Scholkmann, 2016).

Data quality was evaluated in each participant at each preprocessing step by examining different indicators (see Supplementary materials). Specifically, we inspected channel time series (e.g., intensity, optical density, concentration) to detect motion-induced artifacts and signal drifts in the raw data and after wavelet despiking. We assessed the presence of physiological components, such as respiration and cardiac pulsation, in the power spectral density of HbO and HbR prior to temporal filtering. We also evaluated the statistical association between time series fluctuations of Hb chromophores—HbO and HbR—which is expected to be characterized by a strong negative correlation (Villringer and Chance, 1997; Obrig and Villringer, 2003) and an antiphase state (Watanabe et al., 2017). These properties describing the intrinsic relationship between HbO and HbR hemodynamic fluctuations have been confirmed in previous task-based (Wolf et al., 2002) and resting-state fNIRS studies in infants and adults (Watanabe et al., 2017), and even algorithms that maximize the negative correlation between Hb chromophores have been proposed as signal improvement—noise reduction— methods (Cui et al., 2010). Thus, a negative correlation between HbO and HbR signals (i.e., an antiphase state) (Watanabe et al., 2017) was considered as a valid indicator of good data quality. Finally, as part of our data quality assessment routine during preprocessing we also replicated—in each of our three experimental groups— the results of two previous fNIRS RSFC studies with infants. First, as we did at the subject level, we replicated the work by Watanabe and colleagues (2017) showing the expected antiphase state between HbO and HbR signals in our three experimental groups. In addition, following the work by Homae and colleagues (2010) we performed a hierarchical clustering analysis to spatially group channels based on the degree of similarity between their time series, which was measured pairwise as temporal correlation. This analysis was computed for HbO and HbR, and we obtained similar spatial clusters as in the original study, with frontal, temporal and parietal channels of each hemisphere clustering together. Quality assurance figures for each participant at different steps of the preprocessing, and figures of group-level replication analyses are given as **supplementary material**.

### 2.5. Functional connectivity analyses

#### 2.5.1. Temporal group ICA with dual regression

Group-level functional networks were computed by means of a tGICA (Fig. 2a, Beckmann et al., 2009), by temporally concatenating all participants’ datasets after time-series normalization to zero mean and unit variance—producing a single group dataset with dimensions *[channels (46) x Hb chromophores (2)] x [time points (5000) x participants (99)]*. The FastICA algorithm (Hyvärinen, 1999) was applied to the group dataset to extract 15 independent components (IC), which corresponds to the number of principal components explaining 60% of group data variance. The choice of this parameter and other specific details of the ICA analysis are further explained below (ICA order selection). Afterwards, the subject-specific spatial maps associated with each independent functional network were obtained using a dual-regression approach. This two-step method involves an initial spatial regression of the tGICA spatial maps of interest to the subject-specific fNIRS dataset, to obtain the subject-specific time courses associated with each group-level IC. Then, a linear model fit is computed between the estimated subject-specific time courses and the subject-specific fNIRS datasets, to estimate the subject-specific spatial maps (Beckmann et al., 2009). Statistically significant differences between groups were tested channelwise for each functional network by performing a one-way random effects ANOVA with language background as a factor (i.e., BIL, SP and BQ), resulting in 15 spatial maps describing between group differences (i.e., channelwise F-test). Statistical tests were corrected for multiple comparisons at the channel level using the false discovery rate (FDR; q<0.05) method (Benjamini and Hochberg, 1995).

**Fig. 2.**
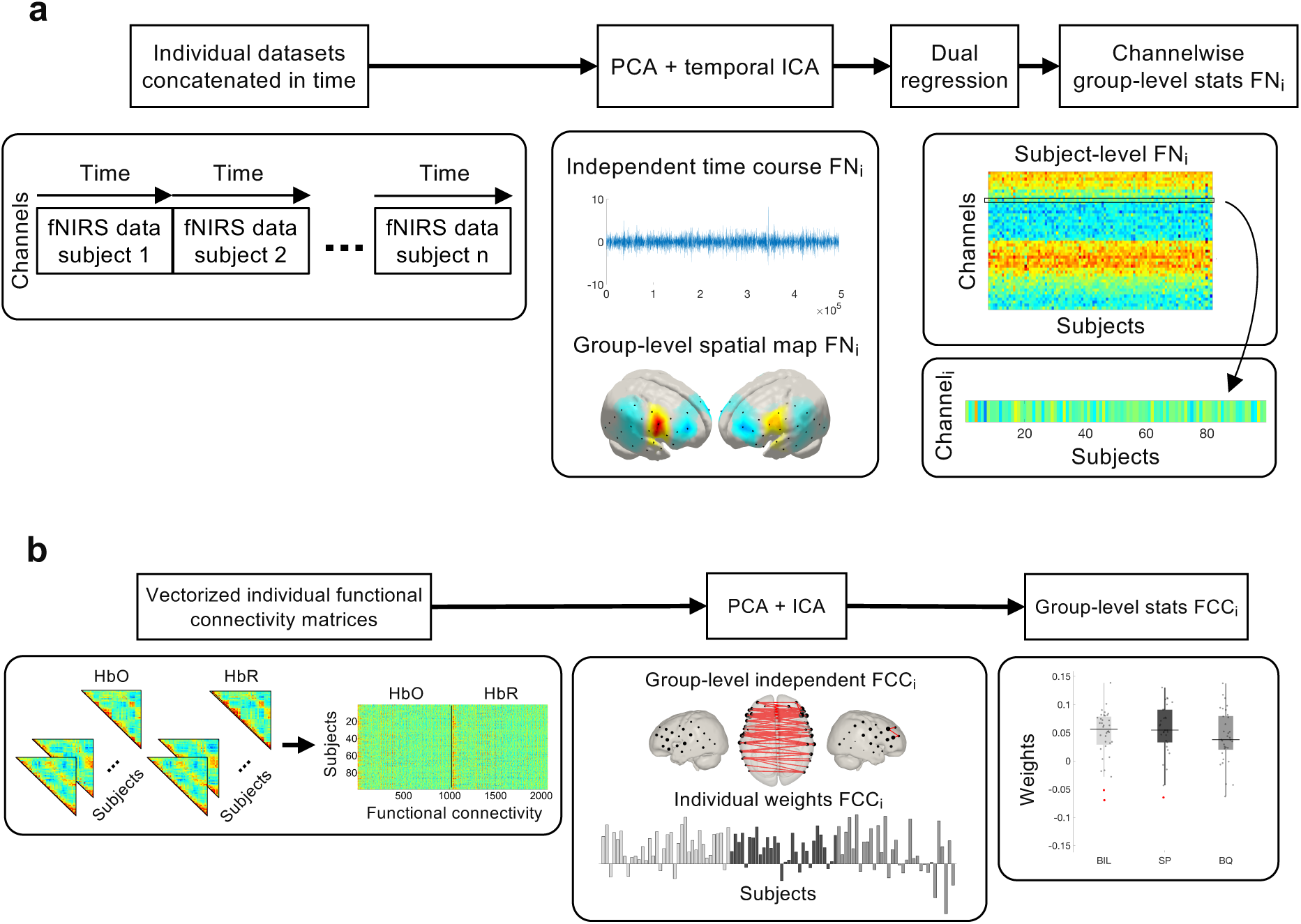
**a)** Processsing pipeline for temporal Group ICA (tGICA) method. **b)** Processing pipeline for connICA method.

#### 2.5.2. Connectome based analyses

A connectome based independent component analysis (connICA) (Amico et al., 2017) was performed based on the functional connectomes computed in all infants. For each individual, the temporal synchronization between channels was evaluated by computing a pairwise robust Pearson’s correlation between the time courses of the HbO and HbR signals separately at every channel for each infant. This robust correlation approach reduces the contribution of possible outlier time points (e.g. due to residual motion artefacts after preprocessing and wavelet denoising) in the correlation estimation (Santosa et al., 2017). Briefly, for each *i*, *j* element representing the preprocessed time series of channels *i* and *j*, a joint weighting matrix is calculated as

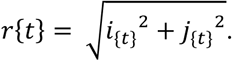

A weighting function (*S*) defined as

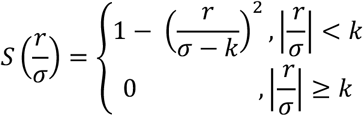

is applied to each *i,j*, element such that *i*_*W*_ = *Si* and *j*_*W*_ = *Sj*. The correlation for each entry of the functional connectivity matrix is computed between the preprocessed weighted signals *i*_*W*_, *j*_*W*_. In this study, we defined *k* = 4.685 as this is the value usually employed in the literature, and we calculated *σ* from the median absolute deviation (MAD) of the signal *r* as *σ* = 1.4826 *MAD*(*r*) (Santosa et al., 2017). Individual robust functional connectivity matrices representing the temporal association between all of the channels were defined for HbO and HbR, where the *i*_*W*_, *j*_*W*_ element reflects the robust Pearson’s correlation between channels *i*_*W*_ and *j*_*W*_. For the sake of simplicity, hereinafter the robust functional connectivity matrices will be referred to as functional connectivity matrices. Individual functional connectivity matrices were then converted from r values to z scores by Fisher’s r-to-z transform, and averaged across subjects within each experimental group. For illustration, the functional connectivity matrix of a representative infant and the average functional connectivity matrices for each group of infants (Spanish monolingual, Basque monolingual and Spanish-Basque bilingual) are presented in Fig. 3.

Individual functional connectivity matrices of HbO and HbR were input to a hybrid connectome-based ICA (connICA) (Fig. 2b., Amico et al., 2017; Amico and Goñi, 2018). First, the upper triangular part of the symmetric functional connectivity matrices of HbO and HbR were vectorized and concatenated for each individual. Then, these vectors were concatenated in rows to form a group-level functional connectivity matrix of dimensions *[99 participants] x [1035 connectivity pairs x 2 Hb chromophores]*. The integration of the information on functional connectivity provided by HbO and HbR was done under the premise that similar RSFC patterns should be observed across Hb chromophores (Mesquita et al., 2010; Homae et al., 2011; Ferradal et al., 2015). Next, the FastICA algorithm (Hyvärinen, 1999) was applied to the group-level functional connectivity matrix to obtain a set of latent group-level independent functional connectivity components, and their corresponding weights in each participant. From this analysis 11 IC were extracted, a number that is equal to the number of principal components necessary to explain 60% of the group data variance. The choice of this parameter is explained in the next section. Finally, the individual IC weights were evaluated as random effects, and an ANOVA was performed with language background as a factor (i.e., BIL, SP and BQ) to examine differences across experimental groups in the prominence of the extracted independent functional components. Statistical tests were corrected for multiple comparisons at the component level using the false discovery rate (FDR; q<0.05) method (Benjamini and Hochberg, 1995).

**Fig. 3.**
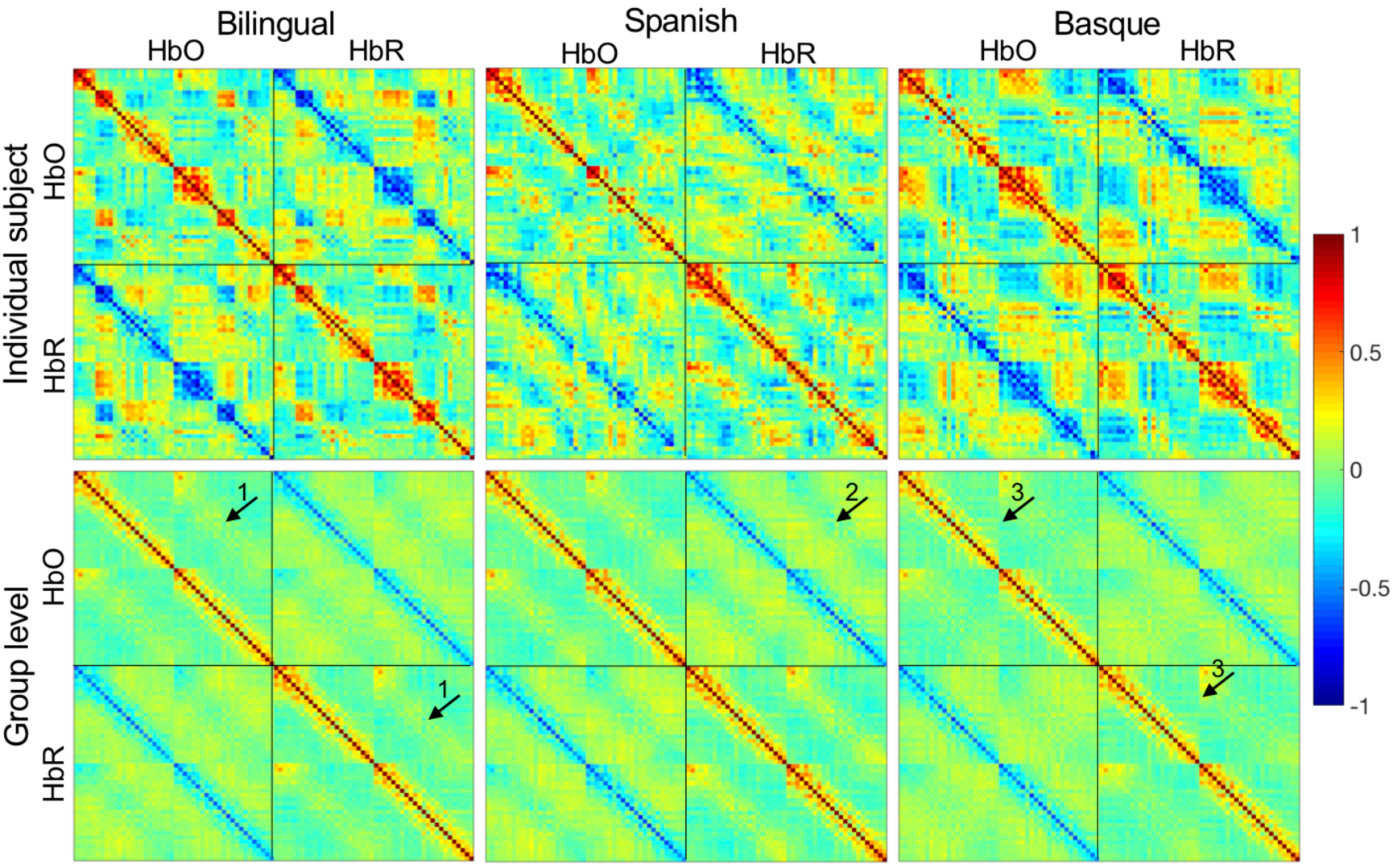
Robust functional connectivity matrices of representative individual subjects (first row) and at the group level (second row) for each of the three experimental groups. In each plot, the robust functional connectivity matrix for HbO and HbR is shown in the top-left part and bottom-right part, respectively. The robust functional connectivity matrix representing the correlation between HbO and HbR is shown in the top-right part. Note that plots are symmetric with respect to the main diagonal. In these plots, channels are ordered from anterior to posterior and from left to right. This allows visualizing an increased correlation between homotopic channels in HbO and HbR (see arrow 1), an increased negative correlation between homotopic channels between HbO and HbR (see arrow 2) and a clear delimitation of left and right hemispheres (see arrow 3).

### 2.6. ICA model order selection

A recurrent issue in studies using ICA to examine resting-state functional connectivity is how to determine the number of IC to be estimated. Here we propose a data-driven approach to determine the number of estimated ICA components by computing two metrics that exploit specific defining properties of ICA estimation and fNIRS data, namely the robustness of the components across multiple realization of the ICA (Himberg et al., 2004) and the assumed similarity between the HbO and HbR derived IC. First, a valid interpretation of the results requires selecting only those independent components that are considered robust; being robustness defined by the identifiability of the component across multiple runs of the ICA algorithm with different random initializations. Briefly, a principal component analysis (PCA) is commonly performed prior to ICA to reduce the dimensionality of the data and/or remove noise components. Then, the number of ICA components to be estimated usually corresponds with the number of PCA components that are retained. Here, we evaluated the robustness of the estimated IC across multiple PCA thresholds by using ICASSO (Himberg et al., 2004; Damaraju et al., 2014). Concretely, 100 realizations of ICA were computed for different PCA thresholds (i.e., 60, 65, 70, 75, 80, 85, 90, 95 and 99) representing the percentage of data variance explained. For the two ICA methods employed in the current work (tGICA and connICA), ICASSO quality index (*Iq*) values were obtained for each component and for each PCA threshold (Fig. 4). This index quantifies the robustness of the estimated components across ICA realizations, with values ranging between 0 (low robustness) and 1 (high robustness). In the tGICA approach, higher *Iq* values (i.e. more robustness) were obtained at the lower PCA thresholds (60% and 65% explained variance). For higher thresholds, the first 15-20 IC also showed high *Iq* values. However, these values start decaying as the number of IC increased, and were particularly low for PCA thresholds above 80%. A similar trend can be observed for connICA. The highest *Iq* values were obtained at the lower PCA thresholds (60% - 70% explained variance), whereas the *Iq* values for higher PCA thresholds were only high up to the first 15 IC and then progressively decreased.

**Fig. 4.**
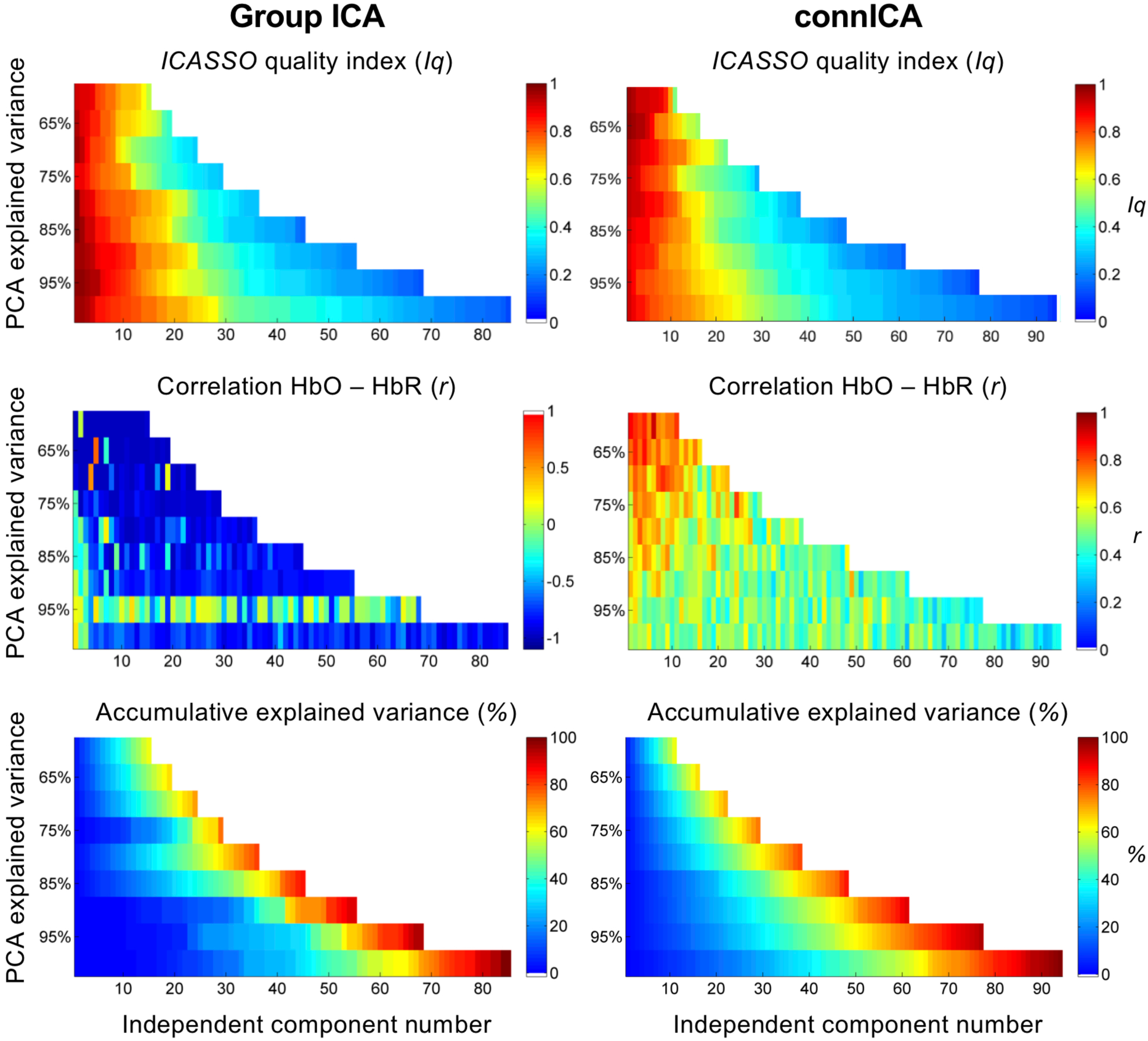
Model order selection criteria for tGICA (first column) and connICA (second column) methods at different thresholds of PCA explained variance (y axis). First row depicts the cluster quality index *(Iq)* for each component as estimated using ICASSO, which represents component stability across multiple ICA realizations. Second row shows the correlation between the estimated HbO and HbR components. Third row shows the components’ accumulative explained variance, being the maximum amount of explained variance determined by the specific PCA threshold.

Second, the expected statistical association between HbO and HbR chromophores can be used as a metric for evaluating the validity of the IC extracted from the ICA algorithm. More specifically, if the extracted IC or maps show a high correlation between HbO and HbR, this implies that they are consistent across Hb chromophores, and based on the reasoning presented above, more physiologically reliable. In the tGICA approach, IC are estimated from HbO and HbR time series, which are expected to be negatively correlated. Thus, a highly similar spatial configuration in the RSNs maps (i.e., channel weights) of both chromophores, but with opposite sign (i.e., negative correlation) is expected. In our tGICA analysis HbO and HbR spatial maps displayed a strong negative correlation at the lower PCA thresholds—up to 75% of explained variance. For higher thresholds, correlation between HbO and HbR components remained negative, but it decreased considerably. For the connICA approach, in contrast, a strong positive correlation between the HbO and HbR independent functional components is expected, as they are estimated from the HbO and HbR functional connectivity matrices, which show alike topology across chromophores (see Fig. 3). The analysis for the connICA method yielded similar results as the previous method. At the lower PCA thresholds (60% and 65% explained variance) the correlation between HbO and HbR components was notably high (*r* ≥ 0.7). At higher PCA thresholds, correlation between components remained positive, but with lower correlation values between HbO and HbR components.

Based on these two metrics (i.e., robustness of IC, and correlation between the extracted IC from HbO and HbR data) an appropriate choice for PCA model order, and in turn for the subsequent ICA method, corresponds to approximately 60% of the explained variance in both approaches (i.e., tGICA and connICA). To reinforce this choice, the variance explained by each IC was computed by calculating the accumulative data variance explained by the components. This analysis showed that for the two approaches with the selected 60% threshold we are able to explain the largest amount of data variance, while also obtaining the highest values in our robustness and consistency metrics. At higher PCA thresholds, explaining the same amount of variance would require including components that are not robust, or which do not show the expected association between HbO and HbR. Based on this information, in the tGICA approach we selected 15 principal components corresponding to 60% explained variance. As a reference, the Laplace approximation for this approach (Minka et al., 2001) yielded a result of 27 components (i.e., 73.5% explained variance). For the connICA method we considered 11 principal components, which corresponds to 60% explained variance. For this method the Laplacian approximation produced a value of 17 components (i.e., 66.34% explained variance).

## 3. Results

After preprocessing and detailed quality assessment of the data acquired in the 123 infants who were recruited to participate in this study, we were able to obtain 99 fNIRS recordings in infants during natural sleep with good data quality for both oxyhemoglobin (HbO) and deoxyhemoglobin (HbR) signals. The final sample included 36 Basque-Spanish bilingual (BIL) infants (21 girls; mean age = 125 ± 4 days), 30 Spanish (SP) monolingual infants (13 girls; mean age = 123 ± 3 days) and 33 Basque (BQ) monolingual infants (17 girls; mean age = 122 ± 4 days). For each infant, the data comprised 46 channels covering frontal, temporal and parietal regions of both hemispheres as shown in Fig. 1, which depicts the optode and channel location, and the sensitivity profile of our setup. All of these datasets showed clear peaks in the power spectrum related to the main frequency and harmonics of the respiratory and cardiac fluctuations (Tachtsidis and Scholkmann, 2016). An antiphase relationship between HbO and HbR signals within each channel was also considered as indication of good data quality (Watanabe et al., 2017). All the participants had recordings with a duration of 9 minutes, which were input for data analysis with temporal group ICA (tGICA) and connICA.

ICA spatial maps representing functional networks of temporally independent spontaneous hemodynamic activity are displayed in Fig. 5. Functional networks (FN) are depicted as t-stat maps from one-sample t-tests on the subject-specific reconstructed spatial maps at the channel level in order to take into account between subject variability. The observed components were robust across multiple realizations of the ICA algorithm based on ICASSO (Himberg et al., 2004) showing consistency values (*Iq*) ranging from 0.49 to 0.91. These components also exhibited high consistency across HbO and HbR, displaying correlation *r* values between −0.97 and −0.99 (supplementary Results - Table 1), as expected due to hemodynamic physiology. The first three functional networks, labelled as sensorimotor networks (FN 1-3), revealed a symmetric pattern over bilateral areas in the precentral and postcentral gyrus. FN 4 and FN 5 cover mainly areas located in the inferior frontal gyrus and the superior temporal gyrus that can be associated with the auditory and the language networks, respectively. Two functional networks were observed over frontal regions: FN 6 is confined to regions in the middle and superior frontal gyrus, and FN 7 comprised middle frontal areas and areas in the inferior parietal gyrus which can be related to the outer brain regions of the default-mode network that is typically observed in RSFC studies with fMRI. Similar to previous evidence with fMRI data, the observed functional networks also exhibit significant patterns of anticorrelated spontaneous activity. In FN 1-5 the spatial distribution of the anticorrelated patterns involved superior and middle frontal areas, in conjunction with posterior areas in the inferior and middle temporal gyrus and with inferior parietal regions. FN 6 showed anticorrelated activity with posterior temporal and inferior parietal regions. In FN 7 the negative spatial pattern was less prominent than the positive part, and included inferior frontal and superior temporal regions. The obtained functional networks for HbO and HbR were reconstructed to the subject space using dual regression (Beckmann et al., 2009) to yield subject-specific functional networks. Between group statistical analyses were conducted to assess the effect of early bilingual exposure on each channel and network (one-way ANOVA at the channel level, FDR corrected among 46 channels, q<0.05). Significant differences between experimental groups were not observed in any of the FN under assessment in HbO and HbR.

**Fig. 5.**
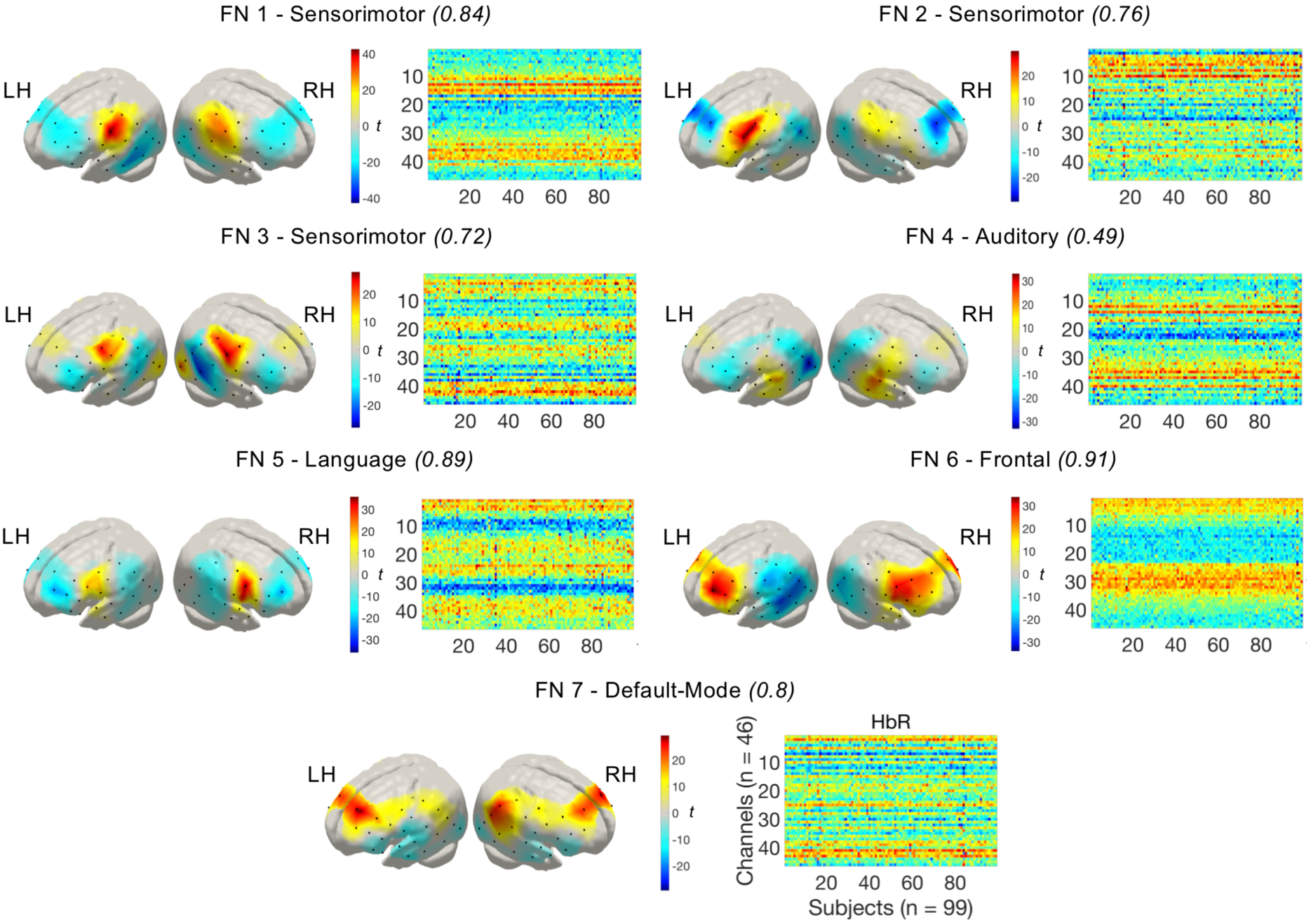
Functional networks (FN) representing the spatial maps derived from the tGICA method. Colorbar shows the t-value of the channel level one-sample t-test computed for each spatial map.

The input of the connICA method are the functional connectivity matrices obtained for each participant, which were computed based on a robust Pearson’s correlation approach as recommended in Santosa et al., 2017. A high degree of similarity was observed at the individual and at the group level in the configuration of the functional connectivity matrices (Fig. 3). A marked negative correlation between HbO and HbR and a stronger correlation between homotopic regions is also evident on the matrices. We consider these features indicative of the quality and reliability of our datasets. The group-level functional connectome components (FCC) extracted from the connICA analysis are depicted in Fig. 6. For the sake of representation, each plot only displays the 10% largest positive connections between nodes (i.e., fNIRS channels) of each FCC, and the size of the node represents the number of connections linked to it. Similar to the tGICA results, the FCC showed a high level of robustness based on the ICASSO algorithm, with consistency values (*Iq*) between 0.5 and 0.96, and a large degree of similarity between the HbO and HbR derived components, with correlation *r* values between 0.7 and 0.95 (supplementary Results - Table 2). FCC 1 is characterized by local, short-range, connections between adjacent nodes. It involves within hemisphere connections between nodes over the whole fNIRS setup, with interhemispheric connections constrained to the most anterior nodes. FCC 2 reflects functional connectivity between homotopic channels across hemispheres. FCC 3 and FCC 4 show a high degree of symmetry, displaying mainly short and long-range within hemisphere connections. FCC 5 and FCC 6 also show a highly symmetric pattern, revealing that the nodes located over superior temporal gyrus are functional hubs with a large number of intrahemispheric connections between temporal and frontal regions, and interhemispheric connections with frontal and posterior temporo-parietal regions. Finally, FCC 7 and FCC 8 are also highly symmetric with their main functional hubs located in precentral and inferior frontal regions, and showing intrahemispheric connections across frontal and precentral regions and interhemispheric connections between frontal, superior temporal and precentral regions. Figures with the complete positive and negative parts of the components, for all the FCC calculated with connICA, and for HbO and HbR, are available as supplementary material. Statistical analyses assessing significant differences across experimental groups were computed on the individual weights that quantify the prominence of each independent FCC in each individual. A one-way ANOVA at the FCC level (FDR corrected among 11 FCC, q<0.05) indicated no significant differences between monolingual and bilingual infants in any FCC.

**Fig. 6.**
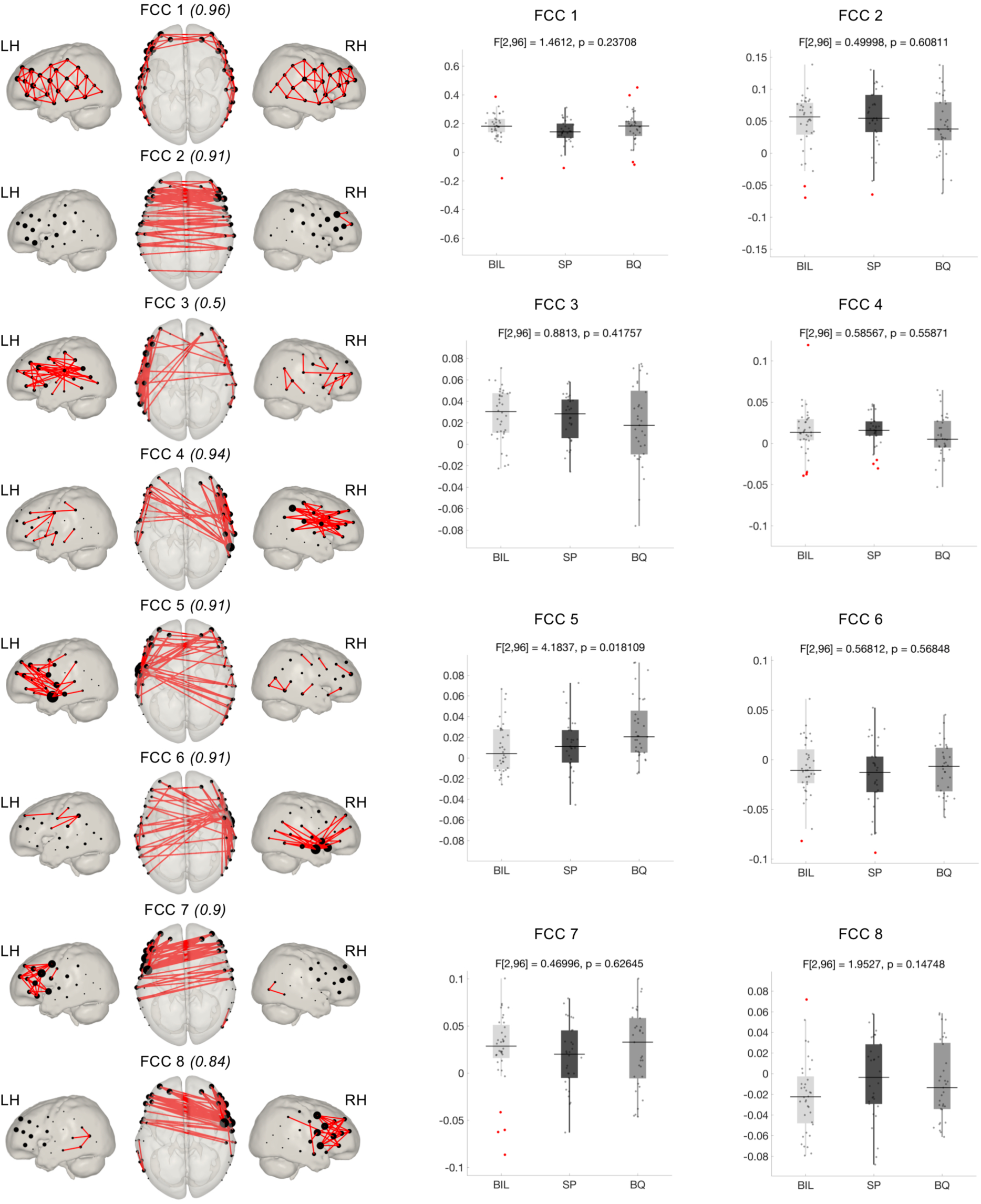
Functional connectome components (FCC) estimated using connICA. Components have been threshold to show only the top 10% of connections (absolute value). Node size was adjusted based on the number of connections reaching each node.

## 4. Discussion

Based on high quality fNIRS data acquired with a large cohort of 4-month-old monolingual and bilingual infants, this work evaluated the effect of bilingualism on functional brain connectivity at this early stage of development. To the best of our knowledge, our sample of 99 valid participants with long fNIRS recordings (9 minutes per infant) is the largest cohort of participants ever acquired to study RSFC in this age group. By applying methods commonly used in fMRI data analysis, we were able to identify large-scale group-level RSFC patterns in the infant brain. To extract these RSFC patterns we implemented two data analyses approaches based on independent component analysis (ICA) at the group level that search for independence between, either the time-courses of spontaneous hemodynamic activity measured with fNIRS (i.e., temporal group ICA - tGICA) (Beckmann et al., 2009; Smith et al., 2012), or in connectivity patterns across multiple individual functional connectivity matrices (connICA) (Amico et al., 2017). Our analyses showed no significant differences between the RSFC of monolingual and bilingual infants at 4 months of age.

The main goal of this study was to assess whether an early and continued exposure to a bilingual environment during the first months of life might impact the configuration of the emerging functional connectivity. The human brain’s capacity to adapt to long-term environmental factors manifests prominently during the first stages of development and it is particularly relevant during early language experience. For this reason, how long-term exposure to two languages affects cognitive and functional brain development received great attention in recent years (Petitto et al., 2012; Costa and Sebastián-Gallés, 2014; Byalistok, 2015; Kovacs et al., 2015). In the current work an effect of bilingualism in RSFC was not observed in infants at 4 months of age, suggesting that a bilingual environment might not affect the configuration of intrinsic functional connectivity at this age. Bilingualism has been only shown to modulate RSFC in adult participants thus far (Luk et al., 2011; Grady et al., 2015; Berken et al., 2016), and evidence of differences between monolingual and bilingual infants in previous neuroimaging studies manifested during explicit language tasks (Conboy and Mills, 2006; García-Sierra et al., 2011; Petitto et al., 2012; Ferjan Ramírez et al., 2017; Nacar-García et al., 2018). This particular age group (i.e., 4 months) was selected in the current study because this is the first point in development in which the behavioral and neural consequences of a bilingual effect have been described (Bosch et al., 1997; Sebastian-Gallés et al., 2012; Molnar et al., 2013; Nácar-García et al., 2018). Furthermore, a previous fNIRS study in infants at this age showed stronger brain responses to native as compared to foreign speech stimuli (Minagawa-Kawai et al., 2011), suggesting that the cerebral basis underlying language processing are already language-specific and might be mediated by the amount of language exposure during the first months of life. It is therefore a possibility that if differences between monolinguals and bilinguals exist at this age, they might only be observable during the performance of language-related tasks. Further research with monolingual and bilingual infants at different ages should also help clarify if the differences observed in adults’ functional connectivity might only emerge at a later stage in neural development. Alternatively, the spatial resolution of the current optode setup might have been insufficient to detect subtle variations in RSFC configuration induced by early bilingualism. Nonetheless, previous studies using similar fNIRS setups have reported differences on various RSFC indicators in infants between 0-6 months of age (Homae et al., 2010; Homae et al., 2011; Fuchino et al., 2013; Imai et al., 2014; Watanabe et al., 2017), demonstrating the feasibility of this technique to assess RSFC during the earliest stages of development.

It is also possible that testing sleeping infants prevented us from detecting subtle differences in RSFC properties across our experimental groups (Watanabe et al., 2014). However, to the best of our knowledge, all previous studies assessing RSFC in infants have been conducted with sleeping infants, irrespective of the imaging modality (i.e., fMRI or fNIRS), and were able to identify RSFC differences induced by the effect of different factors such as premature birth (Smyser et al., 2010) or socioeconomic status (Gao et al., 2015). Brain imaging techniques are extremely sensitive to motion induced artifacts that are commonly observed in acquisitions on awake participants. Collecting RSFC data from awake infants considerably degrades the reliability of the inferred temporal correlations between voxel or channel time courses (Santosa et al., 2017). Because our goal was to collect high quality and reliable RSFC data, we decided to test participants during natural sleep only, which consequently also allowed us to perform longer recordings. Because all infants were tested under similar conditions (i.e., immediately after they fall asleep), a homogenous sleep state was ensured (Gao et al., 2017), minimizing any possible confound due to different cognitive states across participants.

First, the functional networks (FN) extracted with tGICA in our group of infants yield evidence for the presence of a marked bilateral functional correlation in HbO and HbR fluctuations between homotopic brain regions. The spatial configuration of these FN indicates that, at this age, RSFC predominantly consists on correlated activity between anatomically and functionally similar regions across hemispheres, as already described in previous works (Fransson et al., 2007; Homae et al., 2010; Perani et al., 2011; Damaraju et al., 2014). We showed FN located in primary sensorimotor (FN 1-3) and auditory regions (FN 4) which have been repeatedly reported in infant studies with fMRI (Fransson et al., 2007; Damaraju et al., 2014; Gao et al., 2015), but from which evidence from infant studies using optical methods was still limited to studies with few individuals and a limited recording duration (Homae et al., 2010; White et al., 2012; Ferradal et al., 2015). Similarly, we observed a FN overlapping language related regions spreading over the inferior frontal gyrus and superior temporal gyrus (FN 5). Interestingly, a stronger involvement of the right hemisphere in the auditory (FN 5) and language (FN 4) networks can be observed in Fig. 5. This shared feature between language-relevant functional networks could be linked with previous observations showing increased activity in the right hemisphere for speech input, which previous studies have explained by the fact that infants in the first months of life mainly rely on prosodic information during language processing (Homae et al., 2006; Telkemeyer et al., 2009; Perani et al., 2011). Homotopic areas over frontal regions also demonstrated a high degree of functional synchronization. FN 6 was formed by multiple channels within frontal regions and across the midline. The spatial organization of this network supports existing evidence showing that frontal regions become functionally connected during the first year of life (Damaraju et al., 2014; Gao et al., 2015). FN 7 showed a symmetric functional connectivity pattern involving channels in middle frontal and inferior parietal regions, which is partly consistent with the spatial topology of the default-mode network, which exhibits its most prominent activity when the subject is not engaged in any particular task (Fox and Raichle, 2007). Evidence of a developing default-mode network has been observed in infants with fMRI (Gao et al., 2009; Damaraju et al., 2014; Gao et al., 2015), even though these results should be interpreted with caution due to the limited spatial resolution of the current experimental setup and the inability of fNIRS to measure deep medial and subcortical regions usually reported in fMRI studies, such as the posterior cingulate cortex and the precuneus.

The connICA method allowed us to identify macroscale properties of functional network organization at the group level based on the individual functional connectivity matrices (Amico et al., 2017). The functional relations between fNIRS channels form coherent interregional ensembles with distinct topological properties of large-scale functional connectivity. The first functional connectome component (FCC 1) showed short-range functional connectivity between adjacent channels, spanning our whole fNIRS optode setup. This functional connectivity pattern reflecting the intrinsic functional configuration of the infant brain has been shown to progressively decrease over the course of development, whereas long distance connections tend to increase towards a more distributed functional brain organization (Homae et al., 2010; Ouyang et al., 2017). The second component (FCC 2) displayed interhemispheric correlations between homotopic regions. This type of functional connectivity is prevalent in most of the studies that have assessed RSFC in infant subjects and has been linked with the interaction between functional and structural brain maturation (Fransson et al., 2007; Homae et al., 2010; Perani et al., 2011; Gao et al., 2015). Due to the marked spatial symmetry observed in components FCC 3 to FCC 8, we presented them in pairs in Fig. 6. FCC 3 and FCC 4 displayed mostly within hemispheric connectivity between anterior and posterior brain regions in the left and right hemispheres respectively. Functional networks extracted with our tGICA approach also showed patterns of long-range within hemisphere connectivity, but evidence from previous studies suggests that, at this age, this type of connectivity is still immature (Homae et al., 2010; Gao et al., 2015). FCC 5 and FCC 6 showed a functional hub in the left and right auditory cortices, which are densely interconnected with frontal and posterior temporal regions within and across hemispheres. In FCC 7 and FCC 8 connections converged over channels located in precentral and inferior frontal gyrus, which showed intra and interhemispheric connections with channels localized in frontal regions. Due to their spatial characteristics, these components might well represent the activity of sensorimotor, auditory or language regions, in which functional brain networks have been consistently identified in infant populations (Fransson et al., 2007; Perani et al., 2011; Damaraju et al., 2014; Ferradal et al., 2015).

Our results showed reliable patterns of correlated and anticorrelated activity within the observed functional networks and connectome components. One question that might arise from our findings is whether the observed patterns of negative functional connectivity are the result of our preprocessing pipeline including global signal regression, or if they reflect intrinsic, functionally meaningful properties of network organization (Murphy and Fox, 2017). To test this hypothesis, we run exactly the same analyses on our data without applying global signal regression. Using this pipeline, anticorrelated patterns of functional connectivity were still present, but functional networks and connectome components were less spatially interpretable. Considering these results, we decided to include this preprocessing step that we consider essential to account for the effects of systemic physiological confounds that are commonly described in fNIRS recordings (Tachtsidis and Scholkmann, 2016). Most previous optical imaging studies assessing RSFC reported only positive correlations, or presented both positive and negative correlations in the results but only discussed the former (Zhang H. et al., 2010; White et al., 2012). This has been in part due to the limited field of view of the fNIRS setup employed in these studies, but also due to the lack of a straightforward interpretation of the observed anticorrelated activity in the literature (Murphy and Fox, 2017). An interesting finding here is that the regions involved in the anticorrelated networks observed in most of our primary functional networks considerably overlap with the spatial configuration of the functional network we labelled as default-mode network (FN 7). It is therefore a possibility that this activity might reflect the interaction between task-positive and task-negative brain regions.

In summary, RSFC can be reliably measured in young infants using fNIRS. When appropriate analyses techniques are applied for group-level statistical comparison, no differences emerged between our monolingual and bilingual infants’ RSFC at 4 months of age. In light of previous research that demonstrated neural adaptation in bilingual infants in linguistic tasks at this age (e.g., Nacar-Garcia et al., 2018), our results suggest that intrinsic functional networks of the brain are not affected by bilingual experience during the earliest stages of life. Further, considering previously reported differences in adult monolingual vs. bilingual RSFC patterns (e.g., Luk et al., 2011; Grady et al., 2015; Berken et al., 2016), at what stage of development RSFC begin to show changes depending on language environment is open for future research.

## Supporting information

Supplementary Results

Supplementary Data Quality

## Acknowledgments

The authors would like to thank all the parents and infants who generously participate in our studies. The authors also would like to thank Elena Aguirrebengoa for her assistance on recruiting and testing participants.

## Ethics Statement

The study was approved by the local ethics committee. Parents were informed about the procedure of the study and signed a written informed consent before the experiment.

## Author Contributions

**Borja Blanco:** Conceptualization, Methodology, Investigation, Formal analysis, Writing – Original Draft, Writing – Review & Editing. **Monika Molnar:** Conceptualization, Methodology, Supervision, Writing – Review & Editing. **Manuel Carreiras:** Writing - Review & Editing. **Liam H. Collins-Jones:** Formal analysis - Visualization. **Ernesto Vidal:** Formal analysis - Visualization. **Robert J. Cooper:** Formal analysis – Visualization, Writing - Review & Editing. **César Caballero-Gaudes** Conceptualization, Methodology, Formal analysis, Supervision, Writing – Review & Editing.

