## Supplementary Results for "Monolingual and bilingual infants rely on the same brain networks: Evidence from resting-state functional connectivity"

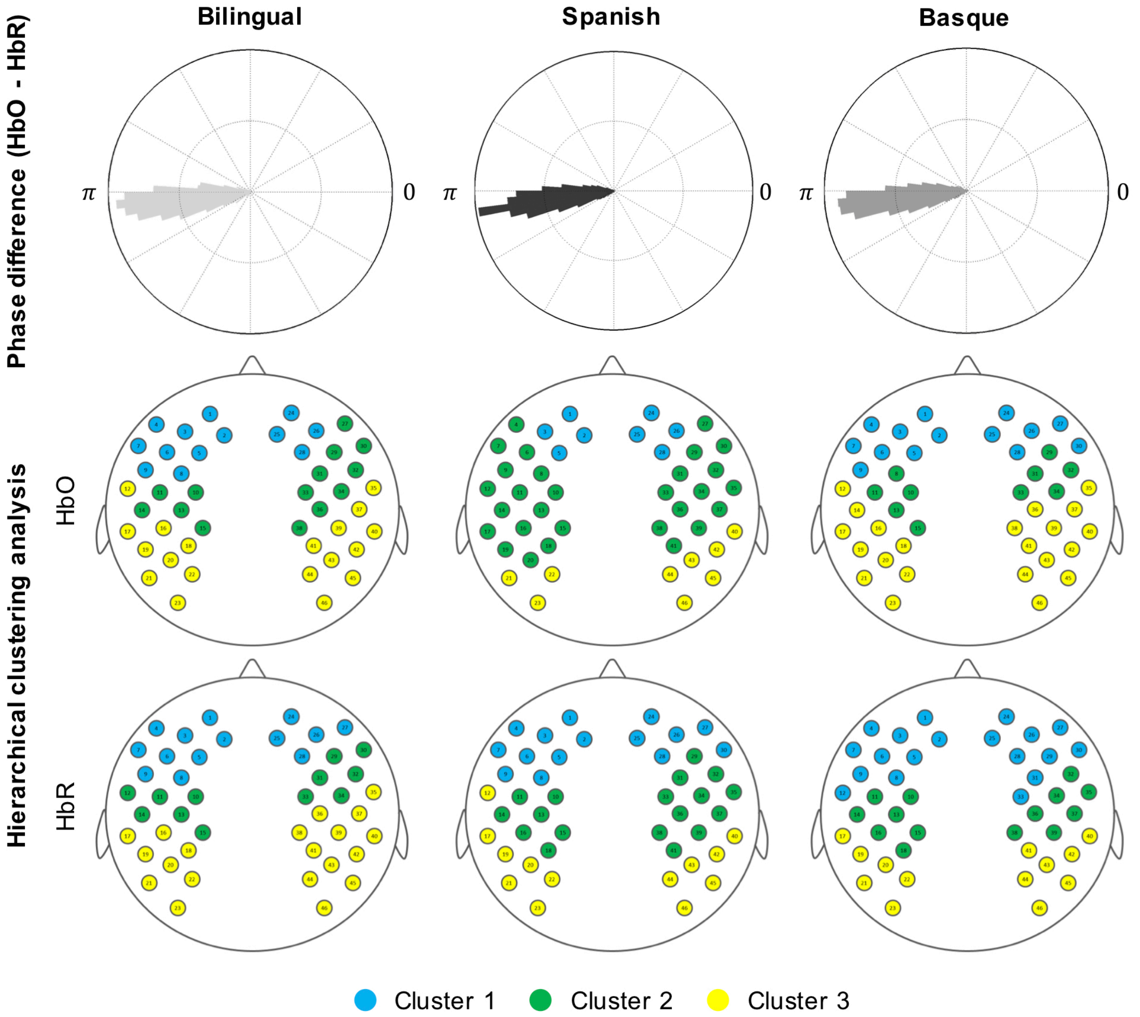


**Supplementary Figure 1.** Group-level data quality assurance plots replicating two previous infant studies assessing resting-state functional connectivity using fNIRS. First row shows the channelwise average phase difference (hPod value, Watanabe et al., 2017) between HbO and HbR in each experimental group. The three groups show a similar pattern characterized by an antiphase state between HbO and HbR, and which replicates previous outcomes. Second and third rows in this figure show the results of a hierarchical clustering approach (Homae et al., 2010) in which channels' time series are clustered based on similarity. A similar cluster configuration can be observed across groups in HbO and HbR. Cluster 1 is formed by channels located in the most anterior part of both hemispheres. Cluster 2 comprises channels located in middle brain regions. Channels located in the most posterior part of the setup are grouped together in cluster 3.

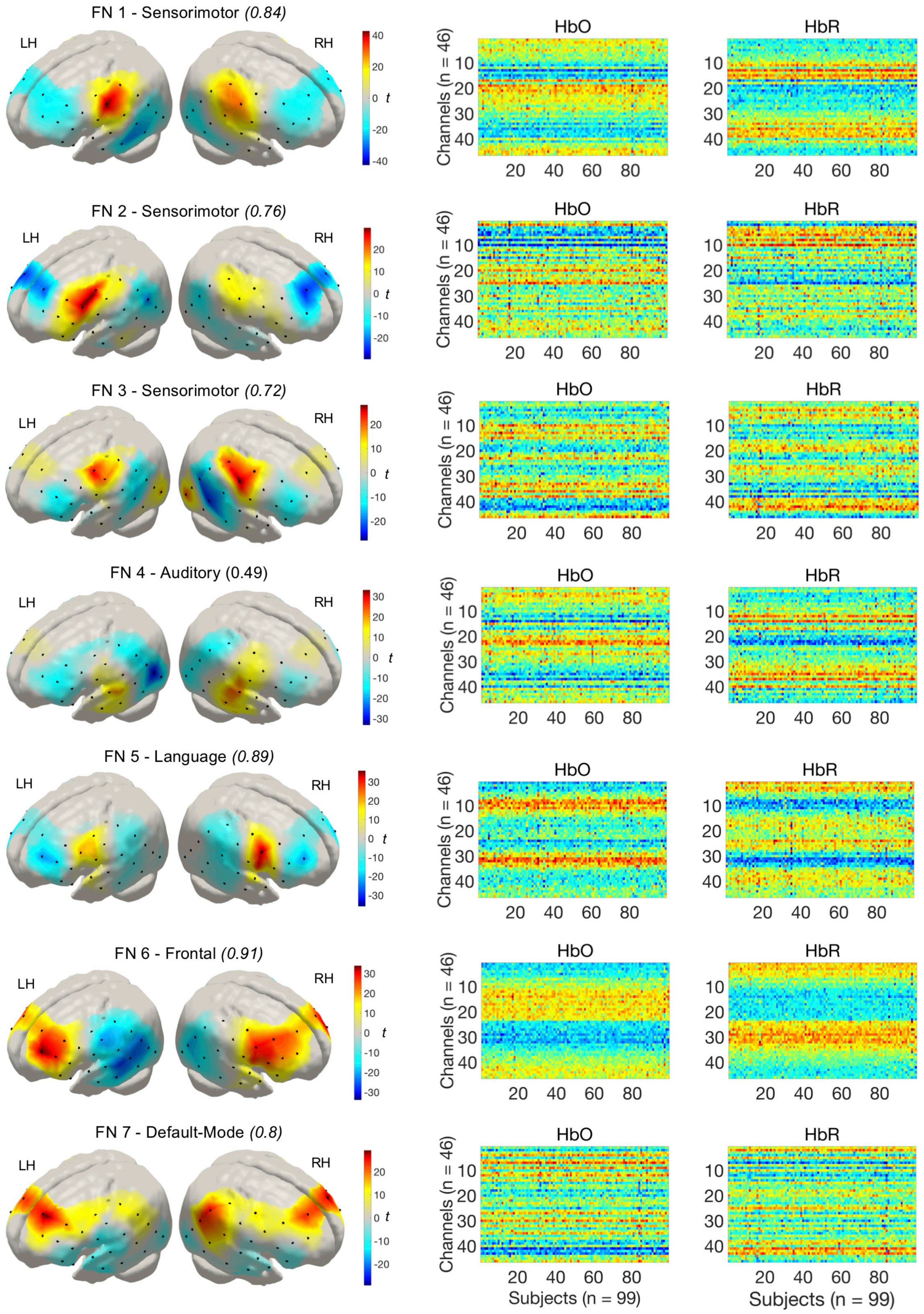


**Supplementary Figure 2.** Functional networks (n = 15) maps extracted with the tGICA approach. Each functional network map was reconstructed to the subject space using dual-regression. These subject-level maps for HbO and HbR are presented in the middle and rightmost columns. Due to the high similarity between HbO and HbR components, functional network maps are displayed for HbR only.

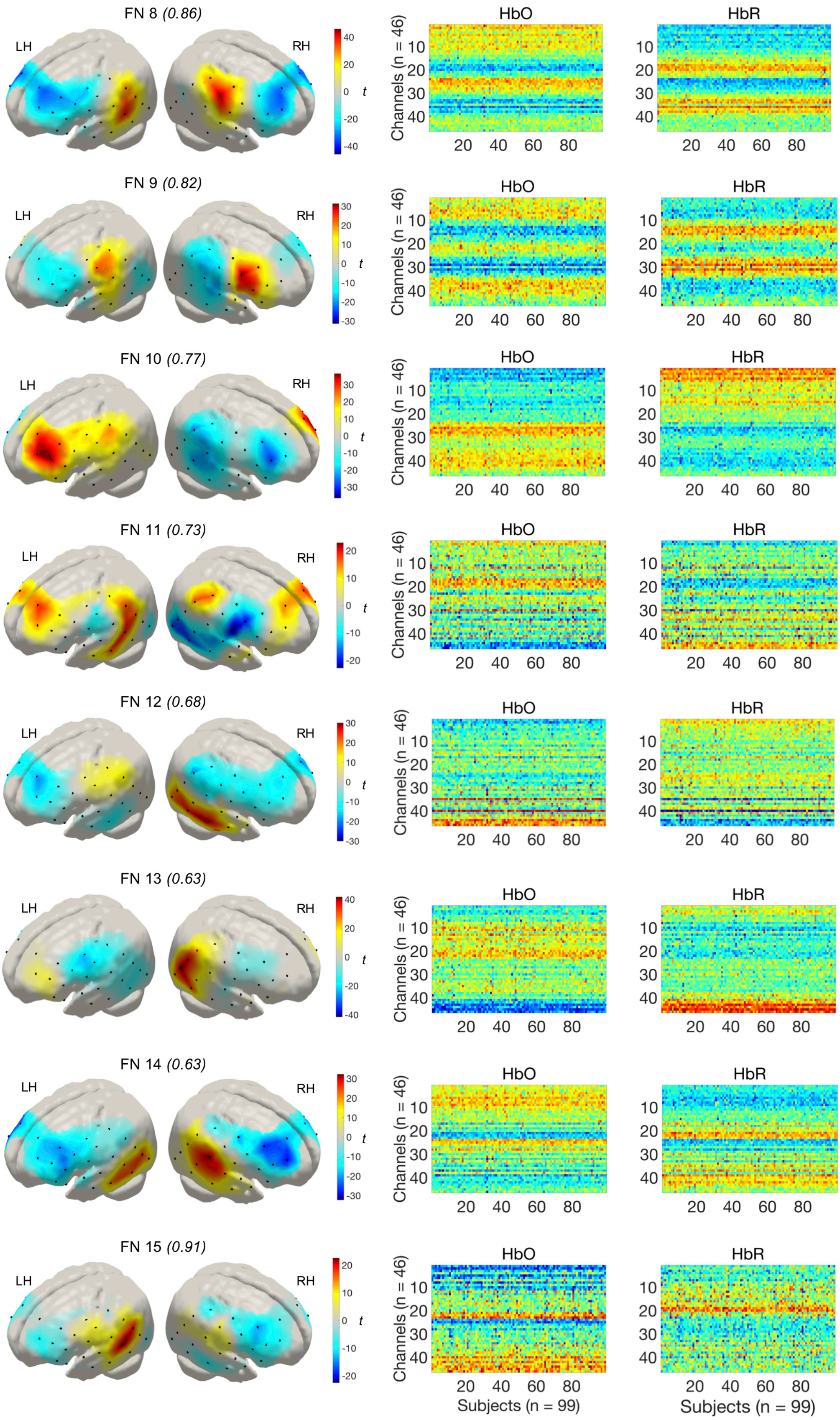
FN 8 (0.86)
FN 9 (0.82)
FN 10 (0.77)
FN 11 (0.73)
FN 12 (0.68)
FN 13 (0.63)
FN 14 (0.63)
FN 15 (0.91)
LH
RH
t
HbO
HbR
Channels (n = 46)
Subjects (n = 99)

| Funtional Network (FN) | *lq* | *r* | *ssd (%)* total = 100 | *ssd (%)* total = 60 |
|---|---|---|---|---|
| FN 1 Sensorimotor | 0.84 | -0.99 | 8.8 | 4.1 |
| FN 2 Sensorimotor | 0.76 | -0.98 | 4.3 | 3.9 |
| FN 3 Sensorimotor | 0.72 | -0.99 | 5.5 | 3.9 |
| FN 4 Auditory | 0.49 | -0.98 | 5.6 | 3.9 |
| FN 5 Language | 0.89 | -0.99 | 7.4 | 4.0 |
| FN 6 Frontal | 0.91 | -0.99 | 10.5 | 4.2 |
| FN 7 Default-Mode | 0.80 | -0.97 | 6.2 | 4.0 |
| FN 8 | 0.86 | -0.99 | 8.9 | 4.1 |
| FN 9 | 0.82 | -0.99 | 7.2 | 4.0 |
| FN 10 | 0.77 | -0.98 | 7.3 | 4.0 |
| FN 11 | 0.73 | -0.98 | 4.5 | 3.9 |
| FN 12 | 0.68 | -0.98 | 5.7 | 3.9 |
| FN 13 | 0.63 | -0.98 | 5.1 | 3.9 |
| FN 14 | 0.63 | -0.99 | 9.0 | 4.1 |
| FN 15* | 0.91 | 0.02 | 3.8 | 3.8 |

**Supplementary Table 1.** Temporal group ICA model order evaluation metrics for the PCA threshold selected (i.e., 60% - 15 ICs) in this study. Sum of squared differences (ssd) are computed with respect to the data after PCA (total = 100%) and with respect to the original data without PCA (total = 60%). Results of the channelwise statistical comparisons between groups on the prominence of the extracted networks showed no differences between groups (FDR corrected among 46 channels, $q<0.05$).

*Stats were not computed on this network because the spatial maps of HbO and HbR did not show a negative correlation.

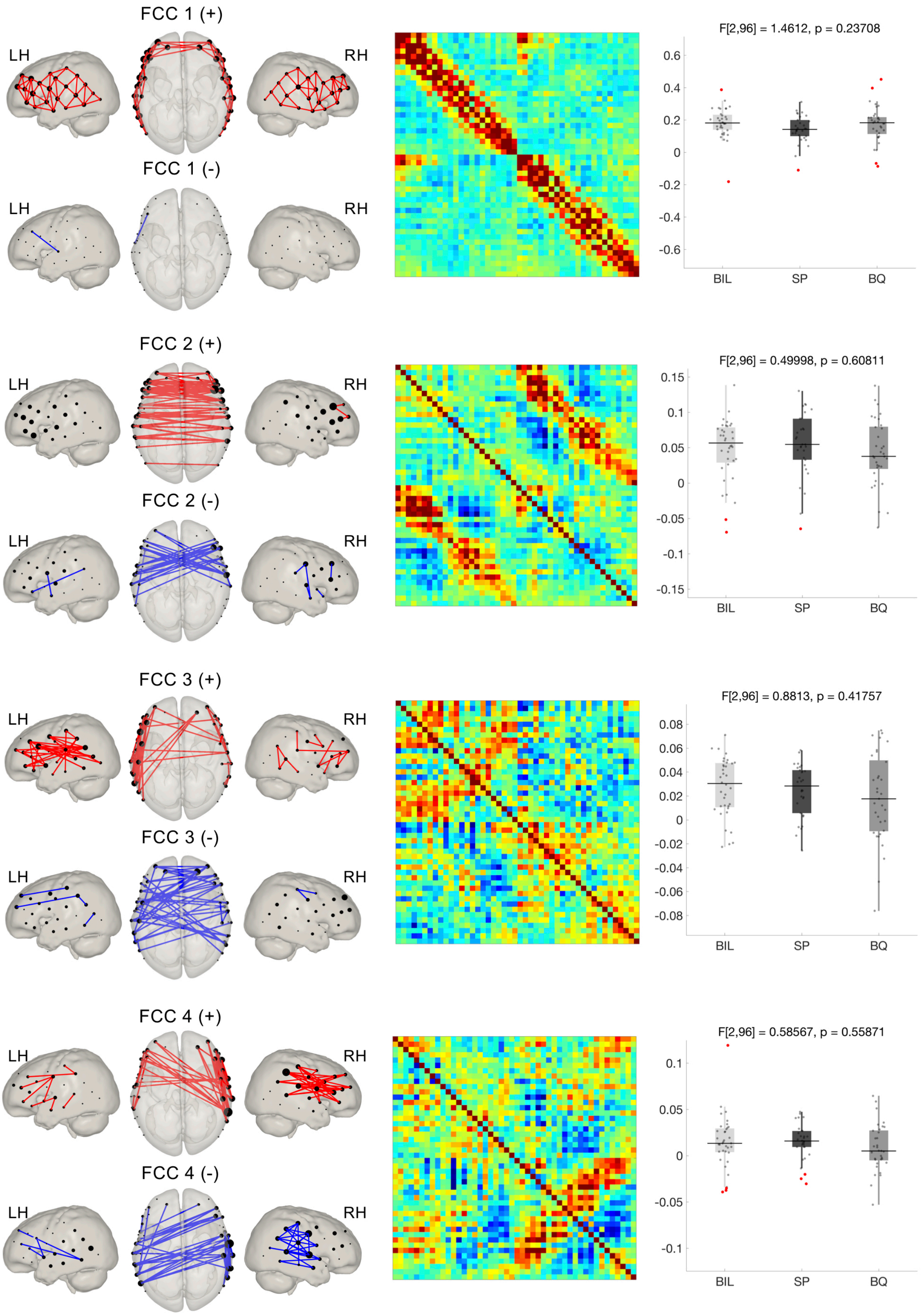


**Supplementary Figure 3.** Functional connectome components (n = 11) obtained with the connICA method. Leftmost column shows the positive and negative parts of the components are represented as nodes and edges (top 10% connections) in a brain template. The middle column shows the components as reconstructed in their original form (i.e., adjacency matrices), with HbO and HbR displayed in the upper and lower triangular sections of the matrix respectively. Statistical comparisons on the weights for each component (right column) revealed no differences between groups. Due to the high similarity between HbO and HbR components, brain maps are shown for HbR only.

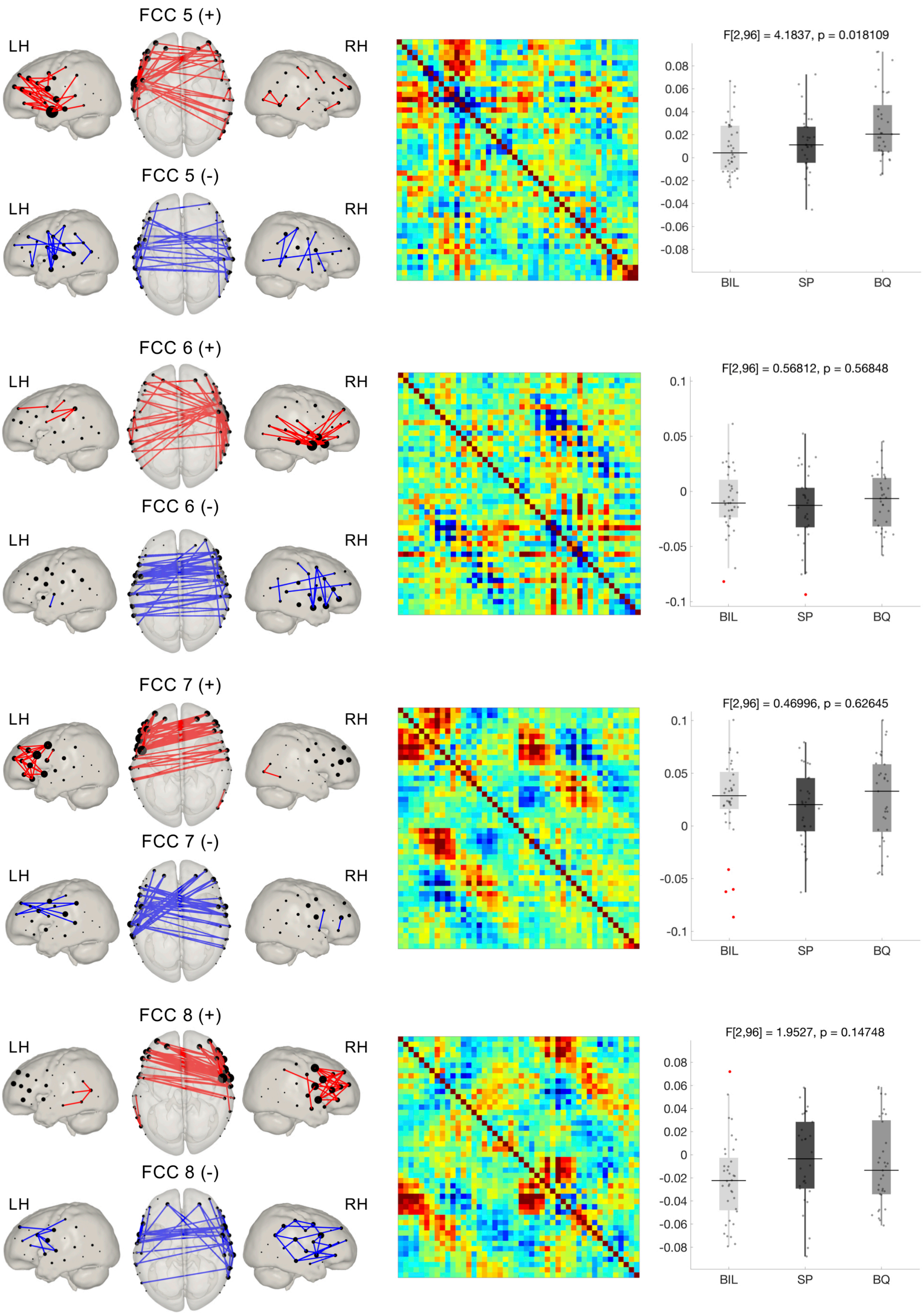
FCC 5 (+)
LH
RH
FCC 5 (-)
LH
RH
F[2,96] = 4.1837, p = 0.018109
0.08
0.06
0.04
0.02
0
-0.02
-0.04
-0.06
-0.08
BIL
SP
BQ
FCC 6 (+)
LH
RH
FCC 6 (-)
LH
RH
F[2,96] = 0.56812, p = 0.56848
0.1
0.05
0
-0.05
-0.1
BIL
SP
BQ
FCC 7 (+)
LH
RH
FCC 7 (-)
LH
RH
F[2,96] = 0.46996, p = 0.62645
0.1
0.05
0
-0.05
-0.1
BIL
SP
BQ
FCC 8 (+)
LH
RH
FCC 8 (-)
LH
RH
F[2,96] = 1.9527, p = 0.14748
0.08
0.06
0.04
0.02
0
-0.02
-0.04
-0.06
-0.08
BIL
SP
BQ

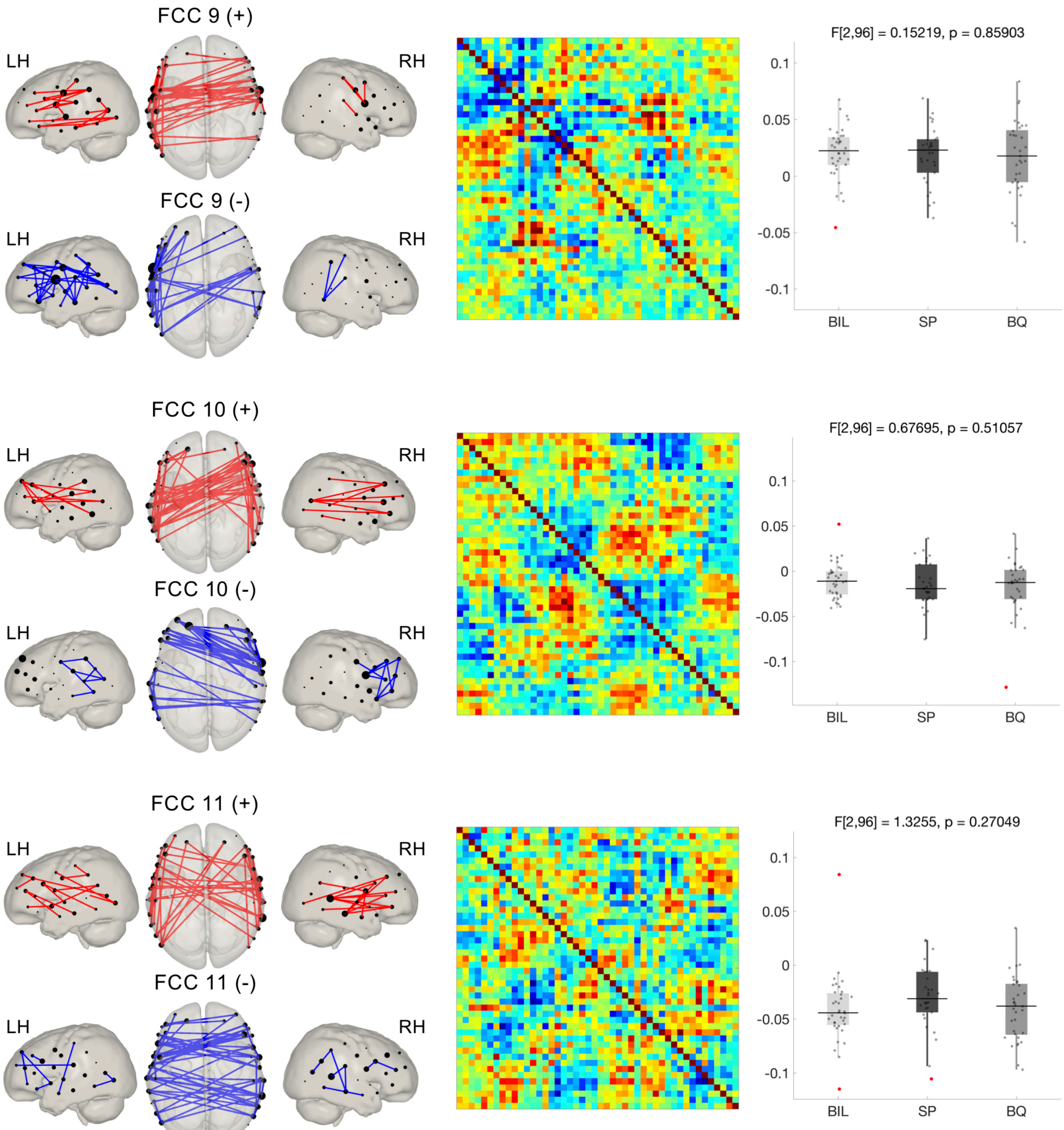
FCC 9 (+)
LH
RH
FCC 9 (-)
LH
RH
F[2,96] = 0.15219, p = 0.85903
0.1
0.05
0
-0.05
-0.1
BIL
SP
BQ
FCC 10 (+)
LH
RH
FCC 10 (-)
LH
RH
F[2,96] = 0.67695, p = 0.51057
0.1
0.05
0
-0.05
-0.1
BIL
SP
BQ
FCC 11 (+)
LH
RH
FCC 11 (-)
LH
RH
F[2,96] = 1.3255, p = 0.27049
0.1
0.05
0
-0.05
-0.1
BIL
SP
BQ

| Funtional Connectome Component (FCC) | *lq* | *r* | *ssd* (100 %) | *ssd* (60 %) |
|---|---|---|---|---|
| FCC 1 | 0.96 | 0.95 | 24 | 5.5 |
| FCC 2 | 0.91 | 0.88 | 7.2 | 5.5 |
| FCC 3 | 0.50 | 0.70 | 7.4 | 5.4 |
| FCC 4 | 0.94 | 0.72 | 7.8 | 5.4 |
| FCC 5 | 0.91 | 0.80 | 7.3 | 5.5 |
| FCC 6 | 0.91 | 0.83 | 7.3 | 5.4 |
| FCC 7 | 0.90 | 0.90 | 8.8 | 5.6 |
| FCC 8 | 0.84 | 0.84 | 7.3 | 5.4 |
| FCC 9 | 0.88 | 0.72 | 7.6 | 5.5 |
| FCC 10 | 0.90 | 0.74 | 8.0 | 5.4 |
| FCC 11 | 0.71 | 0.67 | 7.6 | 5.4 |

**Supplementary Table 2.** Model order evaluation metrics for the PCA threshold selected (i.e., 60% - 11 independent components) in the connICA approach. Sum of squared differences (ssd) are computed with respect to the data after PCA (total = 100%) and with respect to the original data without PCA (total = 60%). Results of the statistical comparisons between groups on the prominence of the extracted components showed no differences between groups (FDR corrected among 11 components, $q<0.05$).
