## Supplementary Data Quality for "Monolingual and bilingual infants rely on the same brain networks: Evidence from resting-state functional connectivity"

For each participant, two figures comprising various plots of the different steps of our data quality assessment routine are displayed in this supplementary material. A brief explanation of these figures is provided below.

**Figure 1**

This figure presents the time series of the complete recording (560 seconds) for the two wavelengths (760 and 850 nm) used by our fNIRS system.

**Top row:** Raw intensity data.

**Middle row:** Optical density data after conversion from raw intensity data.

**Bottom row:** Optical density data after artifact correction using the wavelet-based despike method (Patel et al., 2014).

**Figure 2**

This figure shows various data quality assessment plots at different preprocessing steps. Rows 1 and 2 display the HbO and HbR time series. The third row shows the power spectral density of HbO and HbR concentration data at each preprocessing step. In the bottom row, and for each column (i.e., preprocessing step), leftmost plot shows the functional connectivity matrix. In this plot, the functional connectivity matrix for HbO and HbR is shown in the top-left part and bottom-right part, respectively. The functional connectivity matrix representing the correlation between HbO and HbR is shown in the top-right part (expected negative). Note that these matrices are symmetric with respect to the main diagonal. For preprocessed data, after filtering and global signal regression, the standard functional connectivity matrix and the functional connectivity matrix computed using robust regression are presented. These matrices computed using robust regression are the input for connectome-based ICA method (connICA). The histogram plots in polar coordinates presented on the right part show the phase difference between HbO and HbR time-series (hPod value, Watanabe et al., 2017).

**Left column:** raw HbO and HbR data.

**Middle column:** HbO and HbR time series after filtering.

**Right column:** HbO and HbR preprocessed data, after filtering and global signal regression.

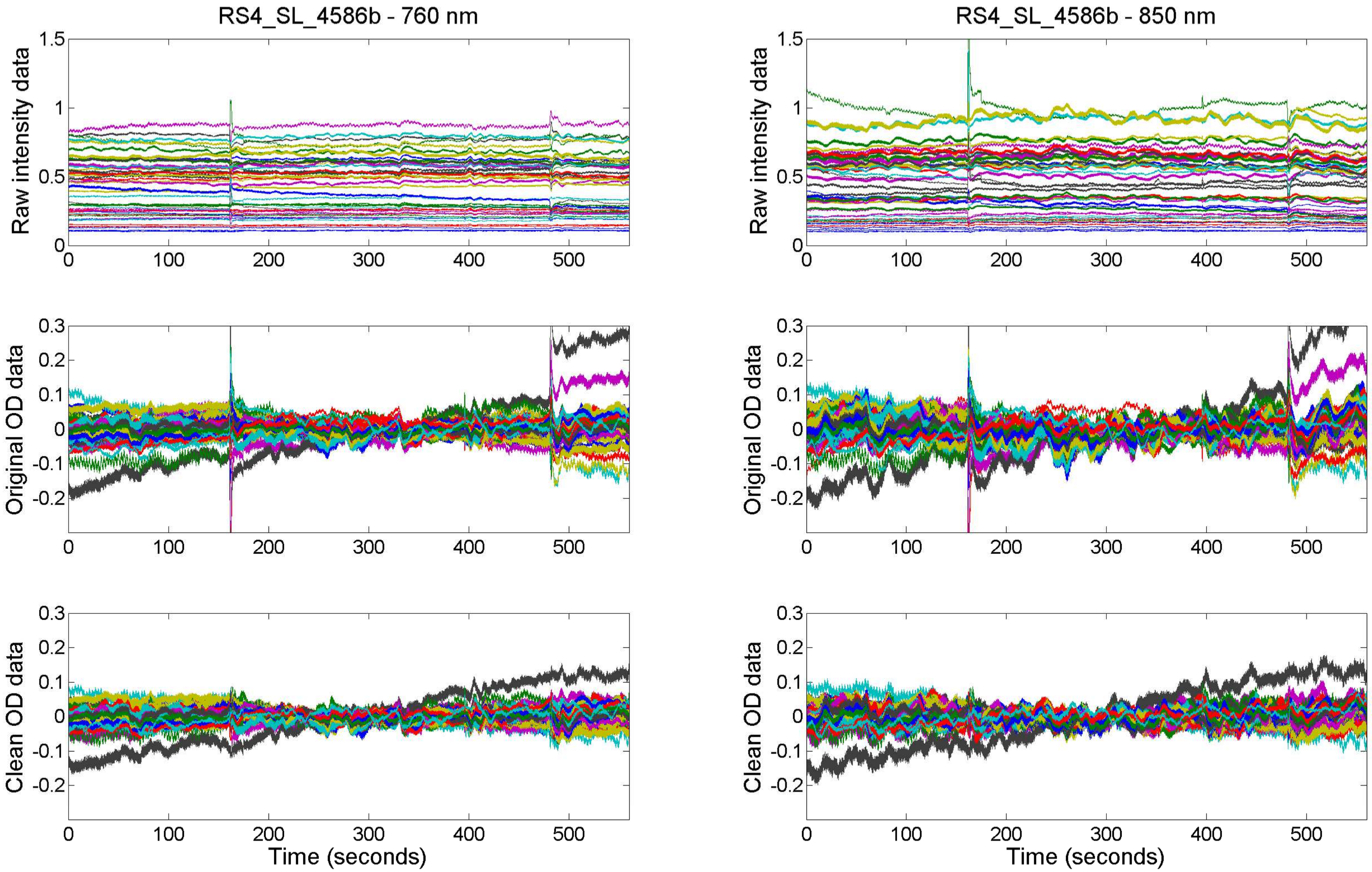
RS4_SL_4586b - 760 nm
RS4_SL_4586b - 850 nm
Raw intensity data
Original OD data
Clean OD data
Time (seconds)
0
100
200
300
400
500
1.5
1
0.5
0.3
0.2
0.1
-0.1
-0.2

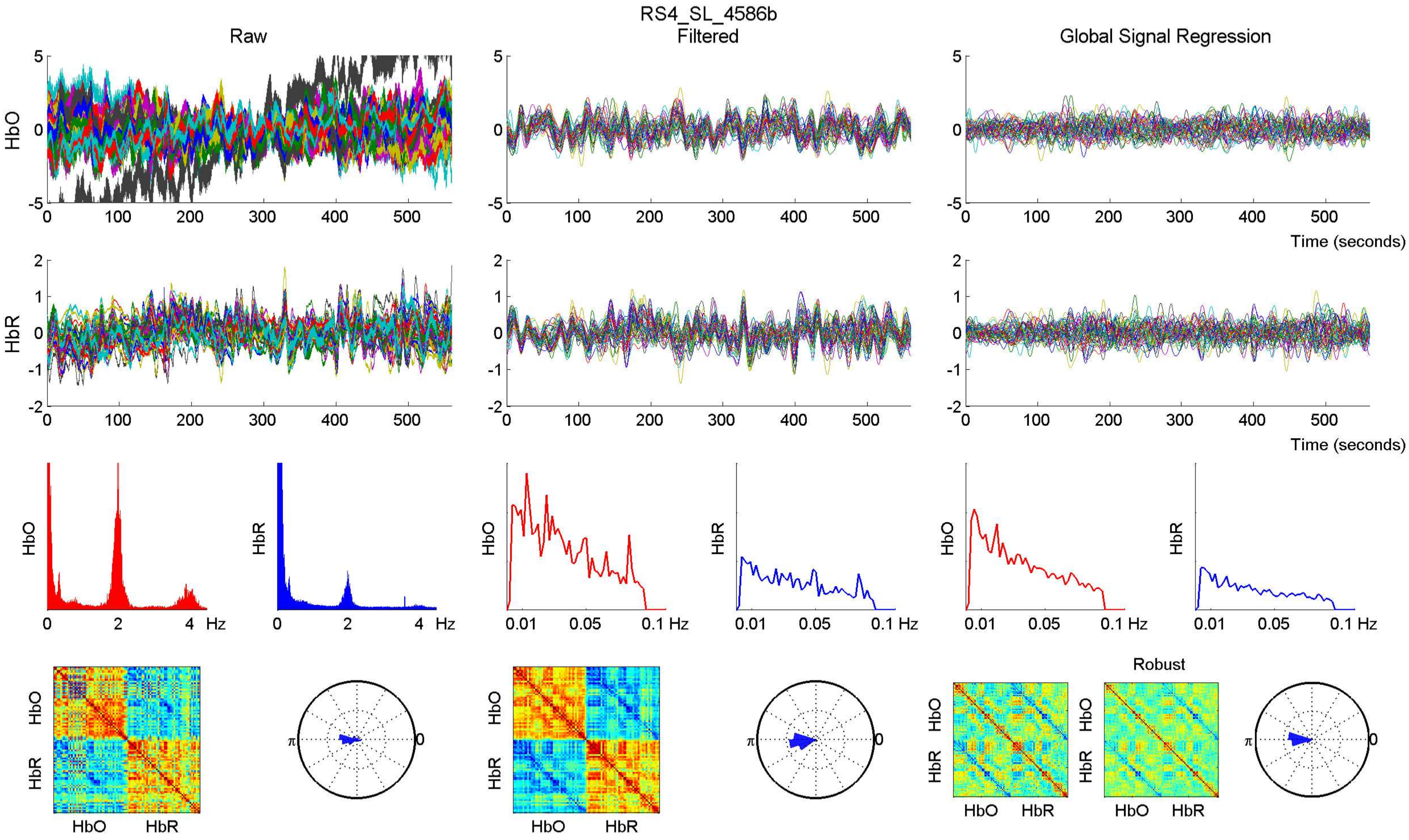
RS4_SL_4586b
Raw
Filtered
Global Signal Regression
HbO
HbR
Time (seconds)
Hz
Robust
HbO
HbR

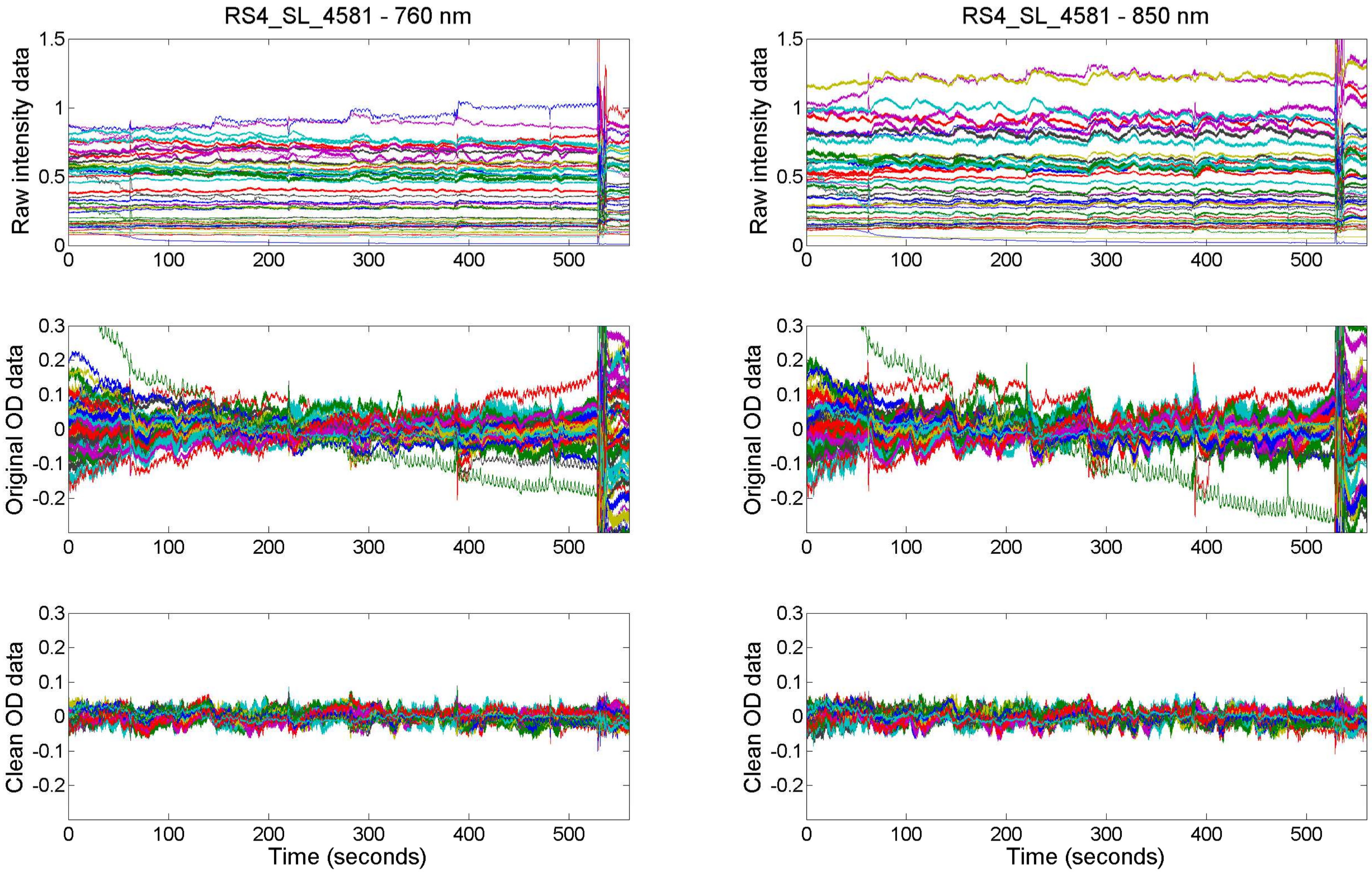
RS4_SL_4581 - 760 nm
RS4_SL_4581 - 850 nm
Raw intensity data
Original OD data
Clean OD data
Time (seconds)

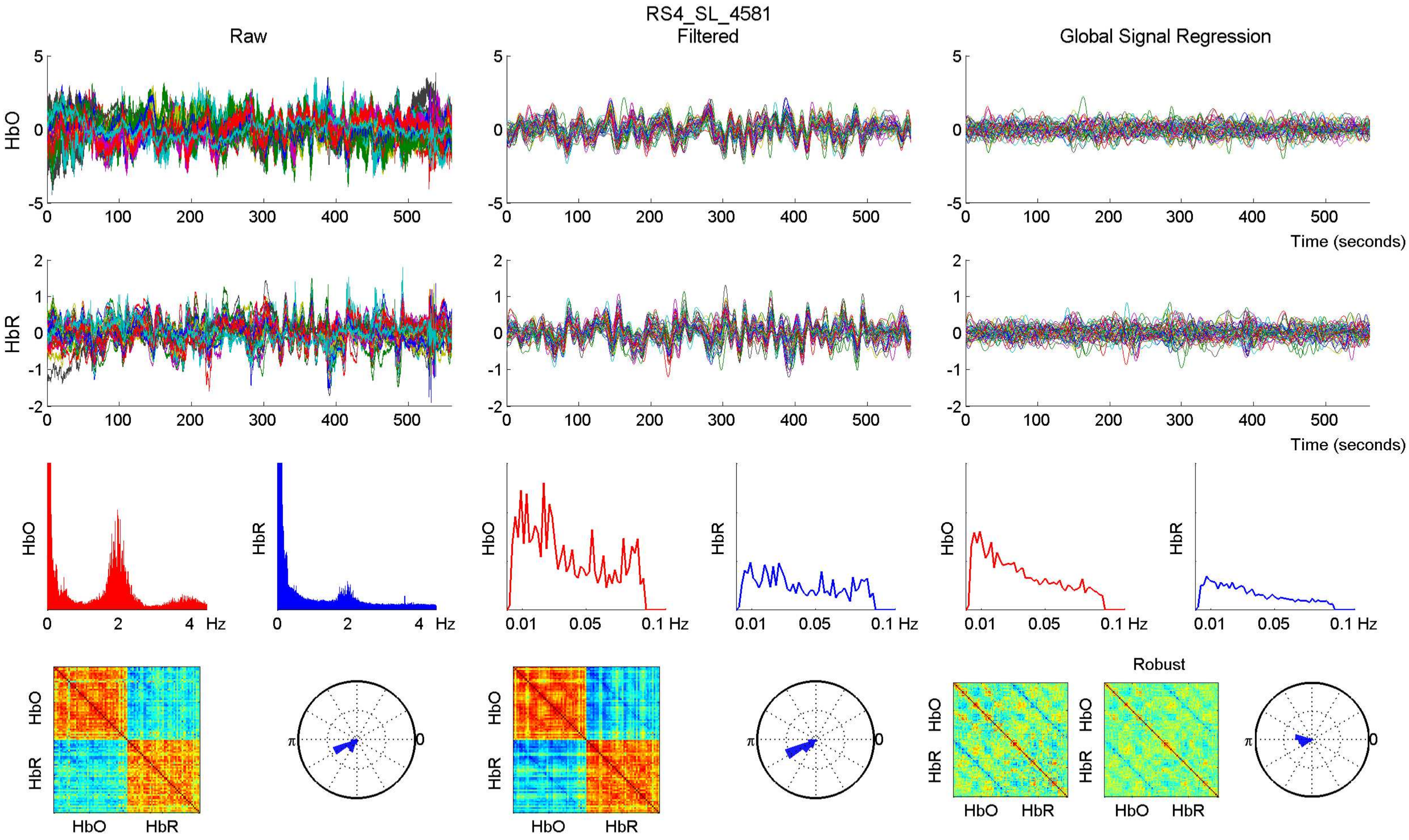
RS4_SL_4581
Raw
Filtered
Global Signal Regression
HbO
HbR
Time (seconds)
Hz
Robust
HbO
HbR

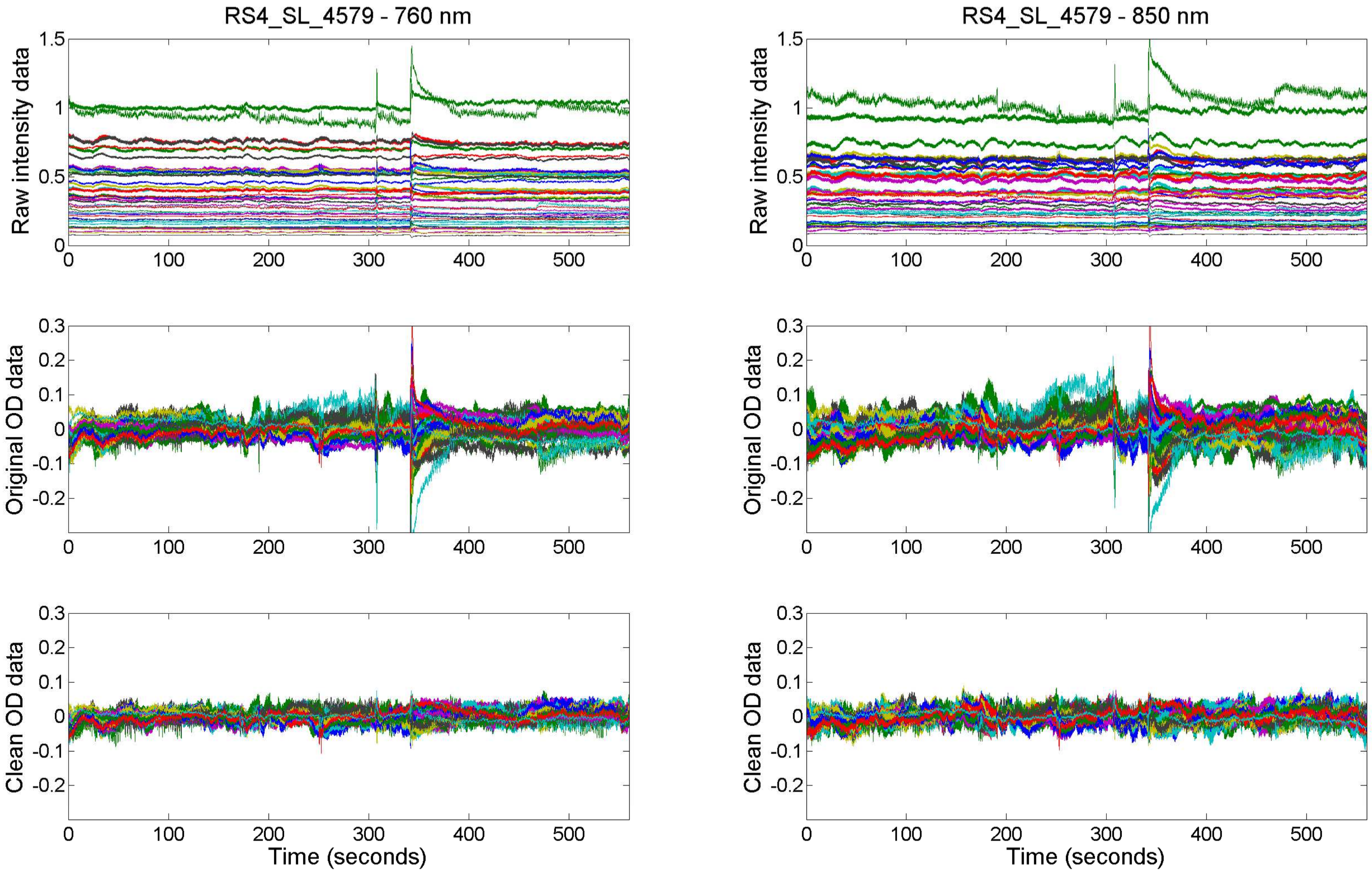
RS4_SL_4579 - 760 nm
RS4_SL_4579 - 850 nm
Raw intensity data
Original OD data
Clean OD data
Time (seconds)
0
100
200
300
400
500
1.5
1
0.5
0.3
0.2
0.1
-0.1
-0.2

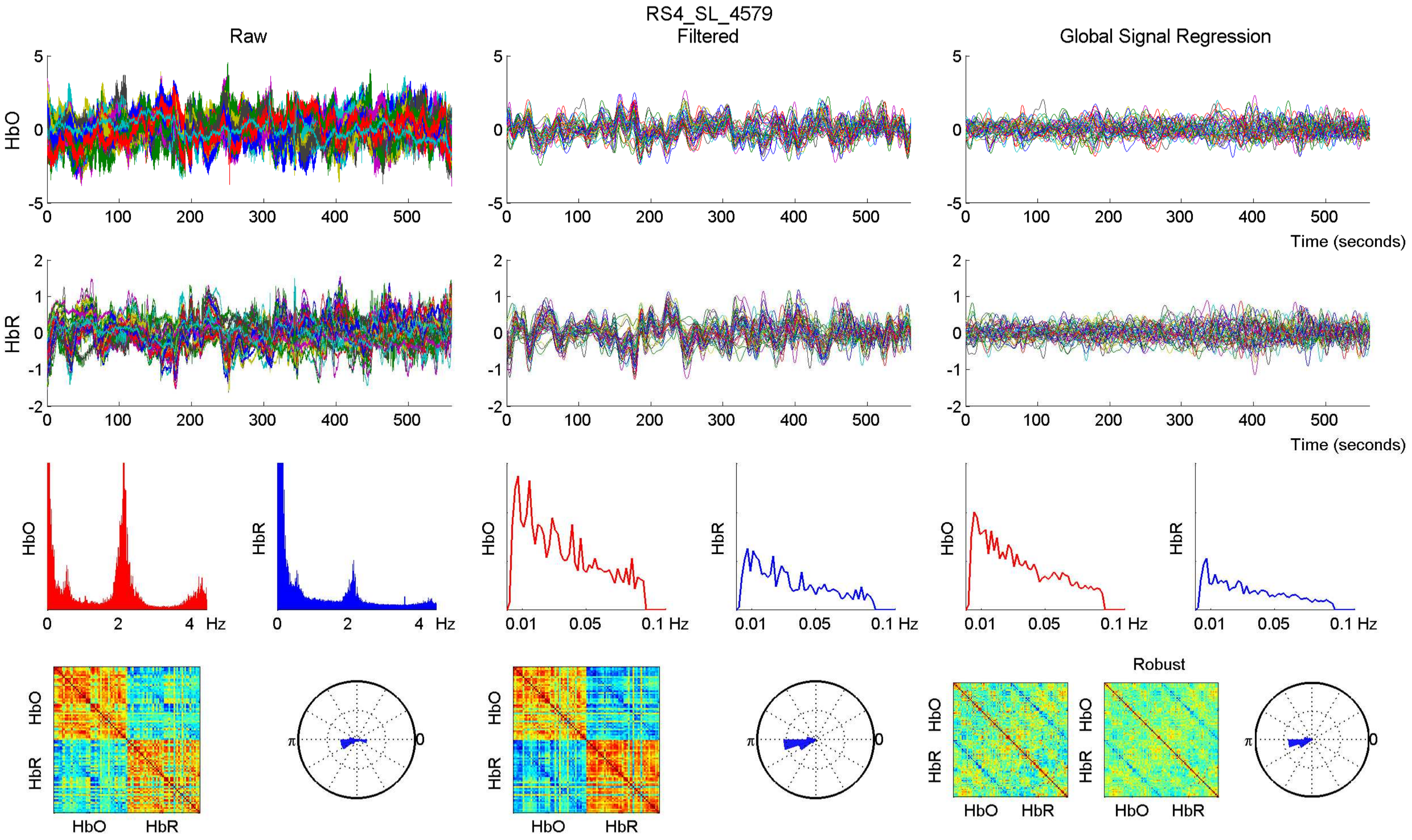

RS4_SL_4579
Raw
Filtered
Global Signal Regression
HbO
HbR
Time (seconds)
Hz
0.01
0.05
0.1 Hz
Robust
π
0

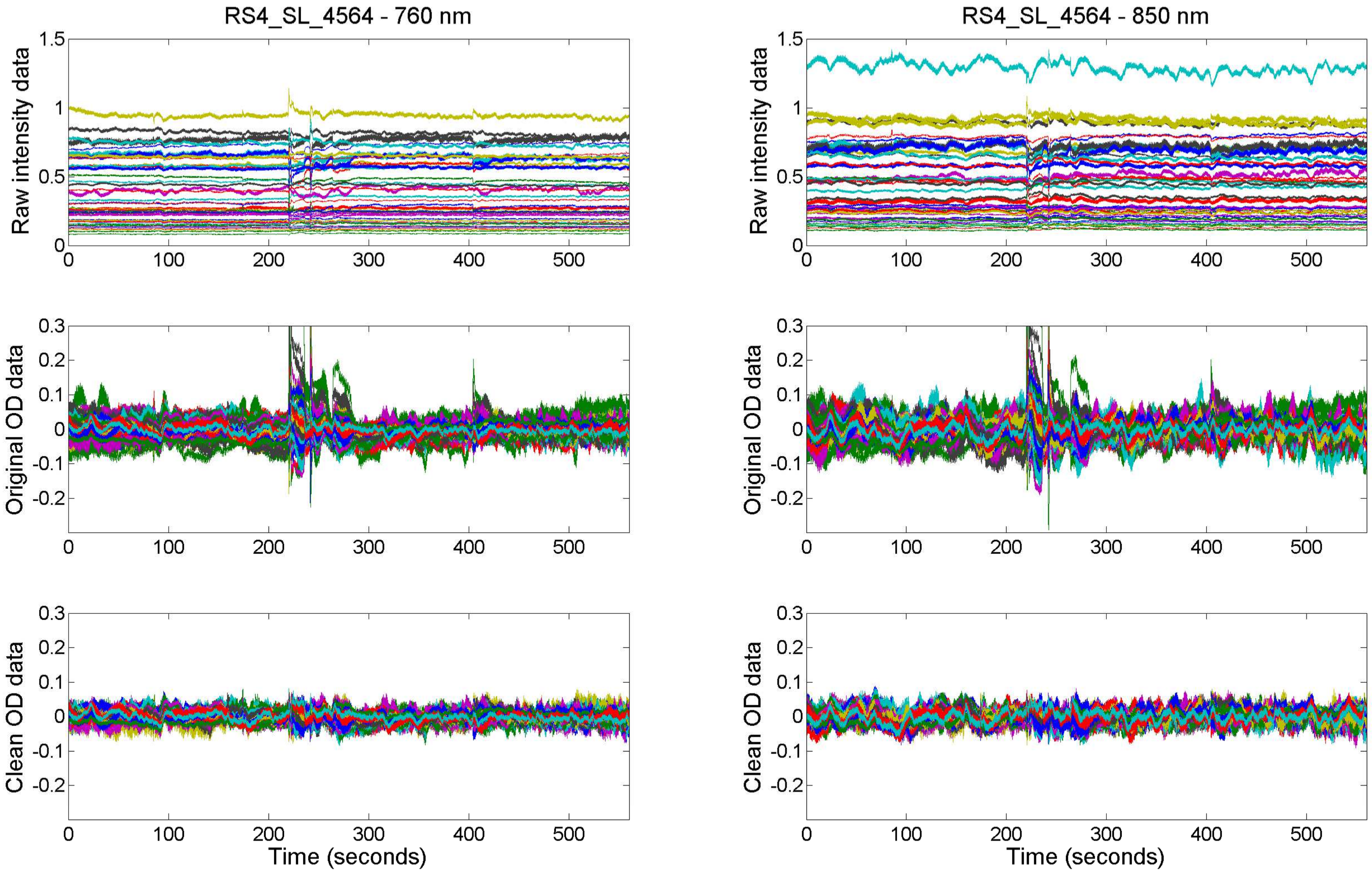

RS4_SL_4564 - 760 nm
RS4_SL_4564 - 850 nm
Raw intensity data
Original OD data
Clean OD data
Time (seconds)

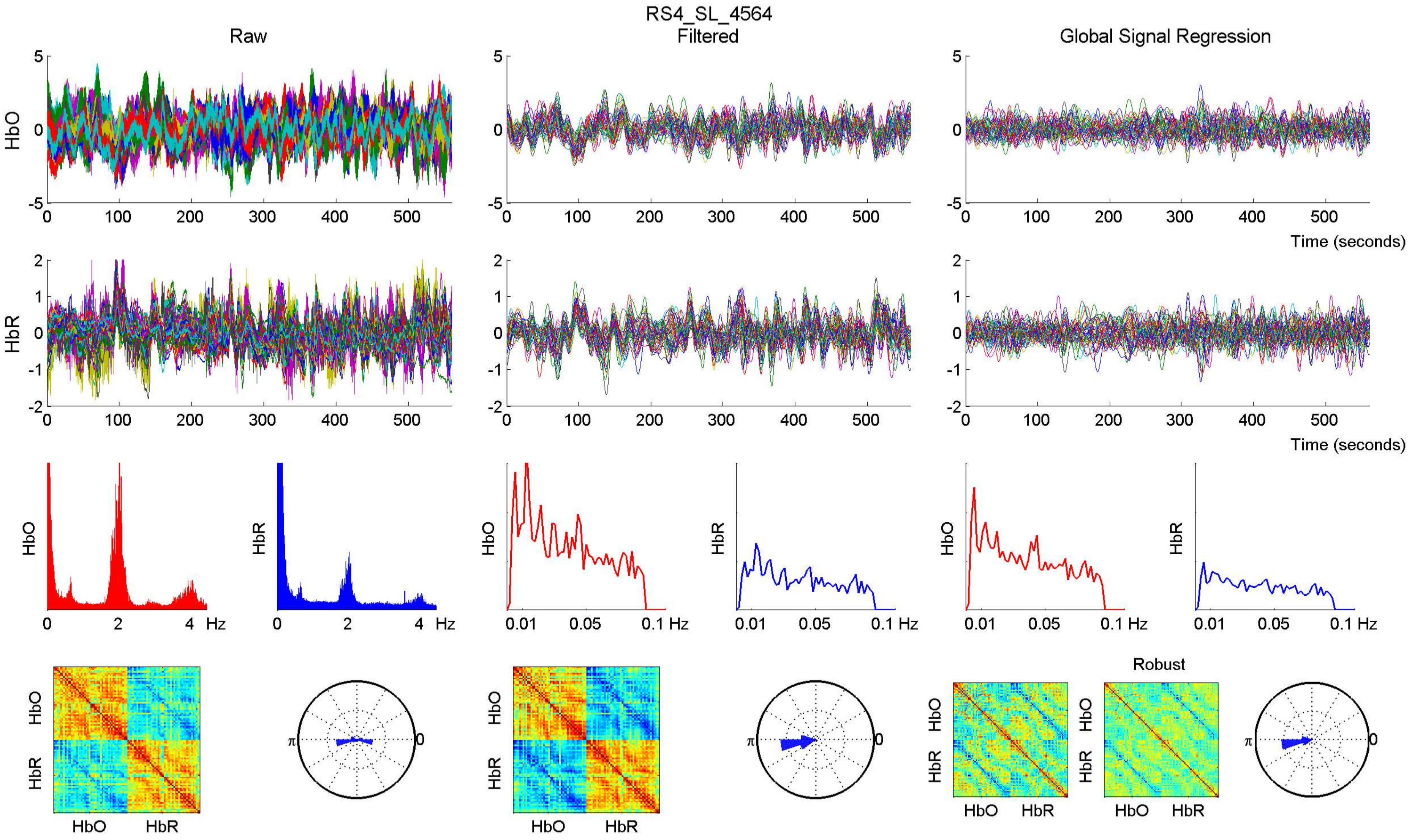
RS4_SL_4564
Raw
Filtered
Global Signal Regression
HbO
HbR
Time (seconds)
Hz
Robust

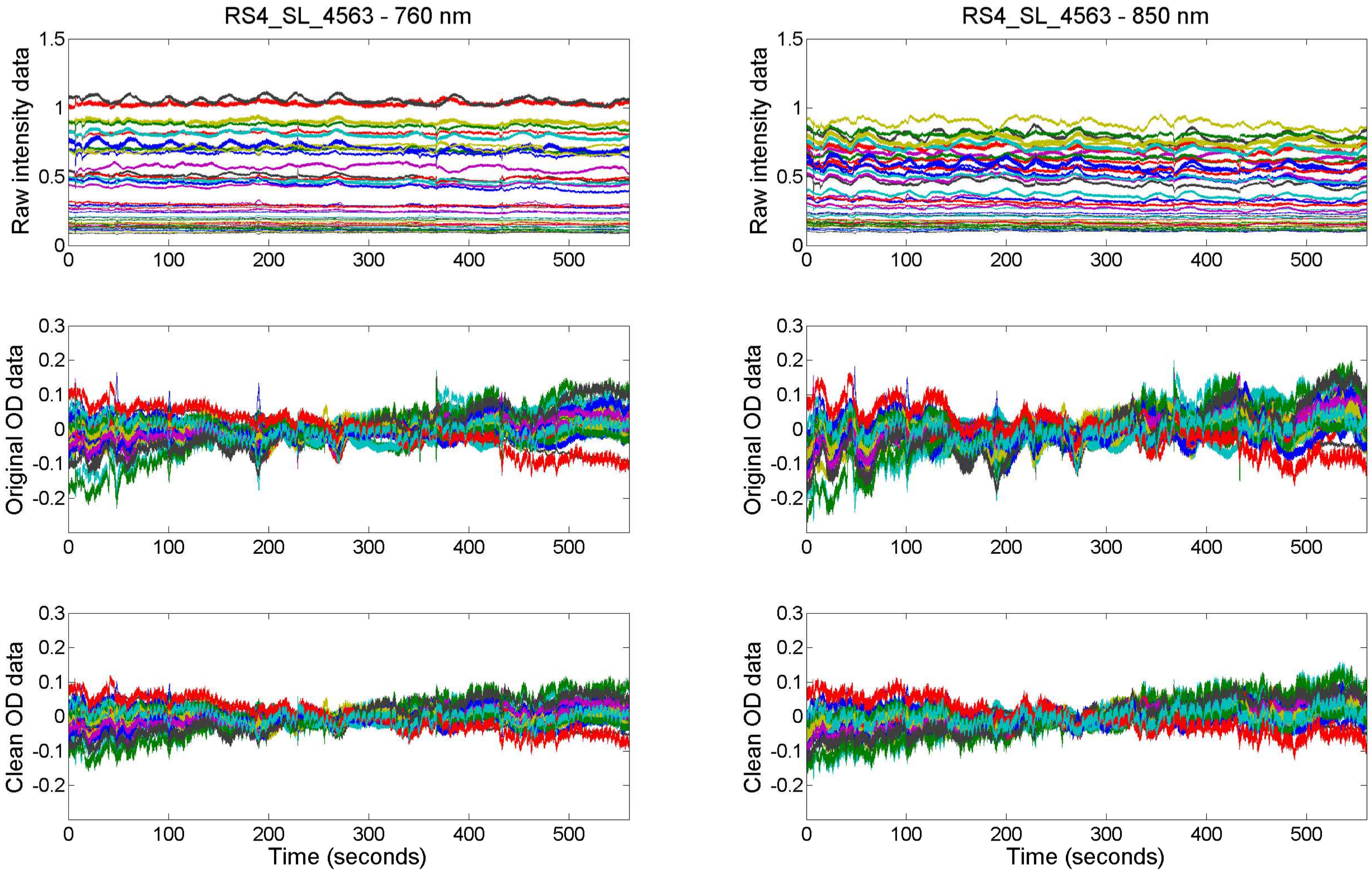
RS4_SL_4563 - 760 nm
RS4_SL_4563 - 850 nm
Raw intensity data
Original OD data
Clean OD data
Time (seconds)

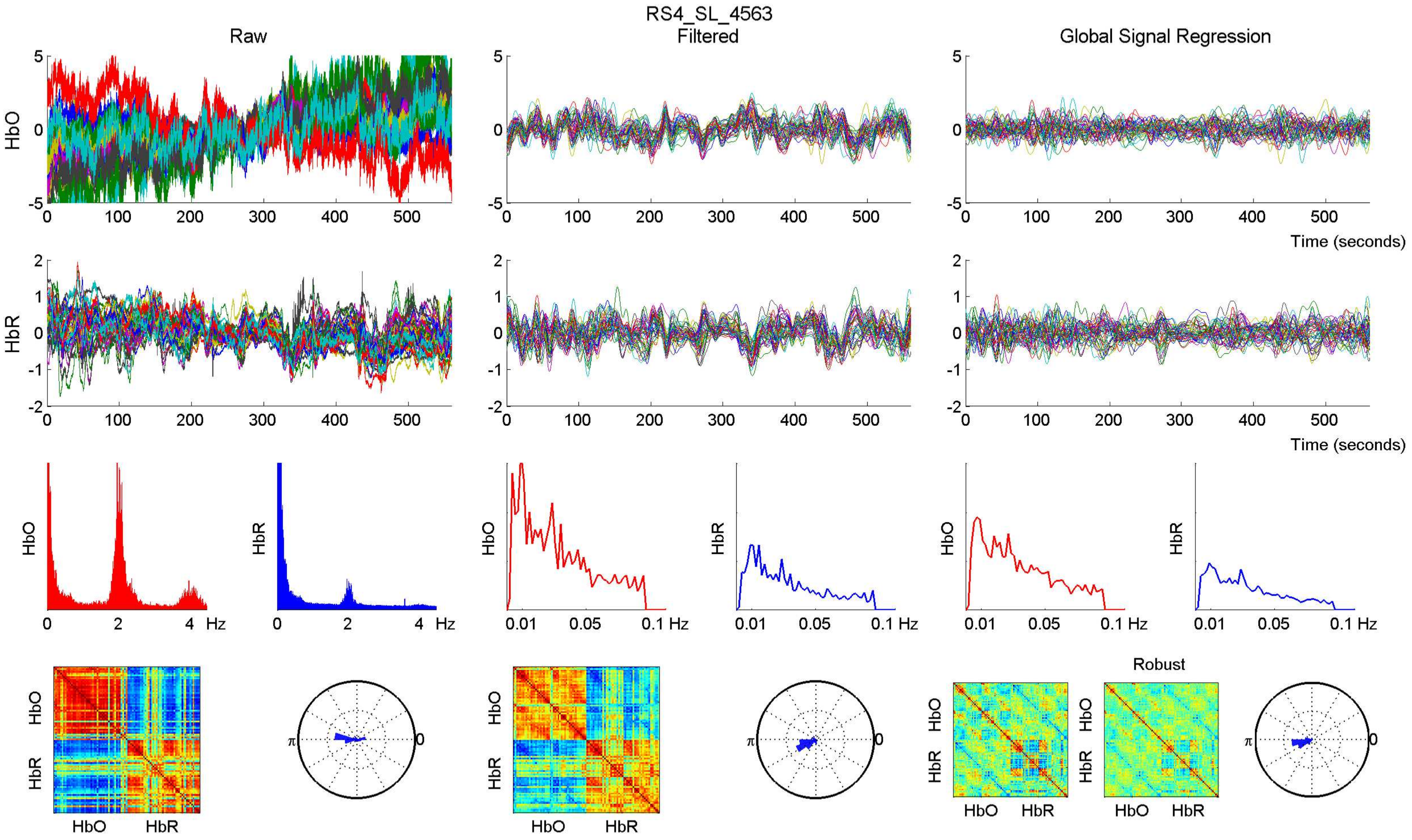
RS4_SL_4563
Raw
Filtered
Global Signal Regression
HbO
HbR
Time (seconds)
Hz
Robust
HbO
HbR

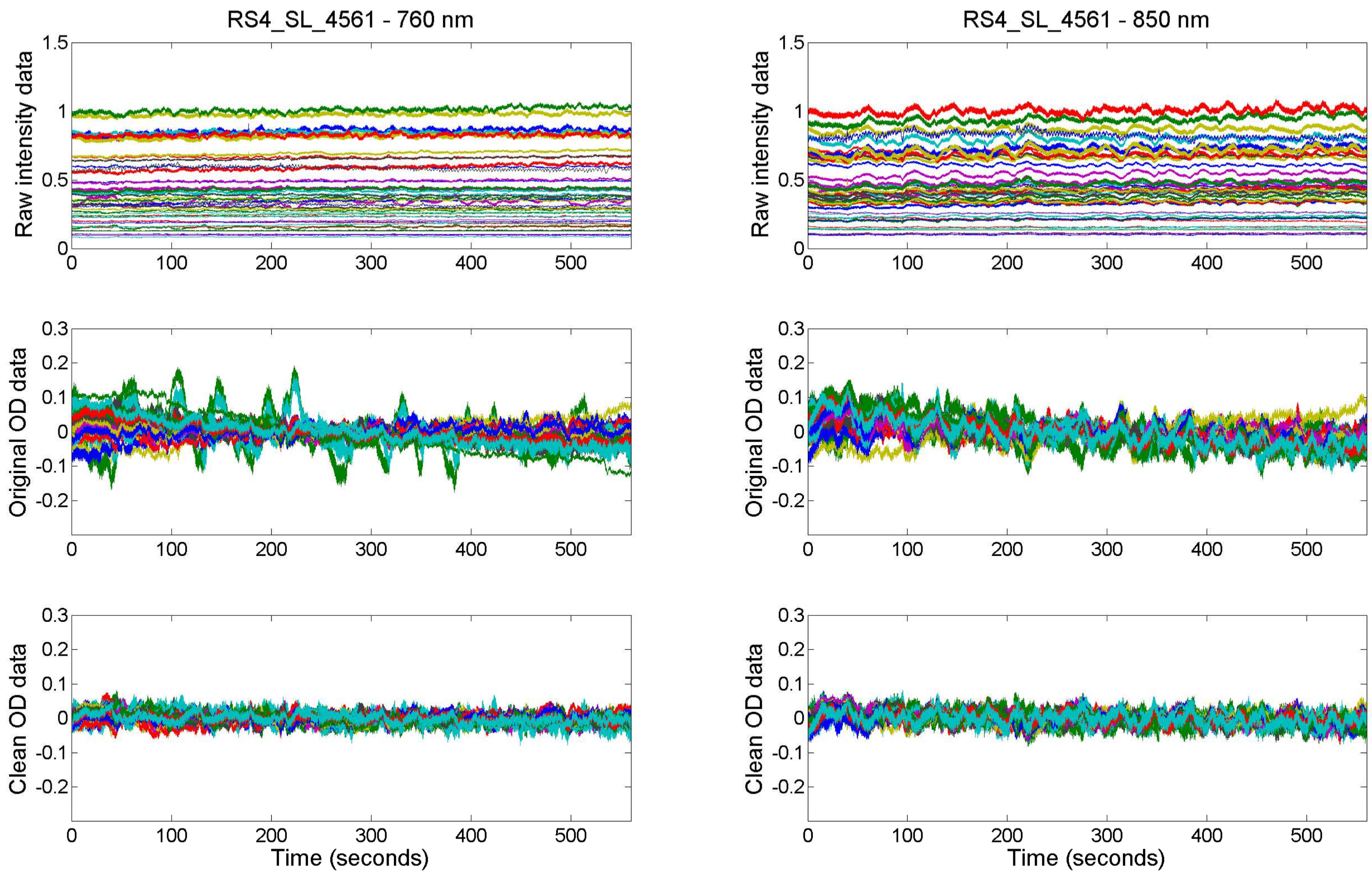

RS4_SL_4561 - 760 nm
RS4_SL_4561 - 850 nm
Raw intensity data
Original OD data
Clean OD data
Time (seconds)

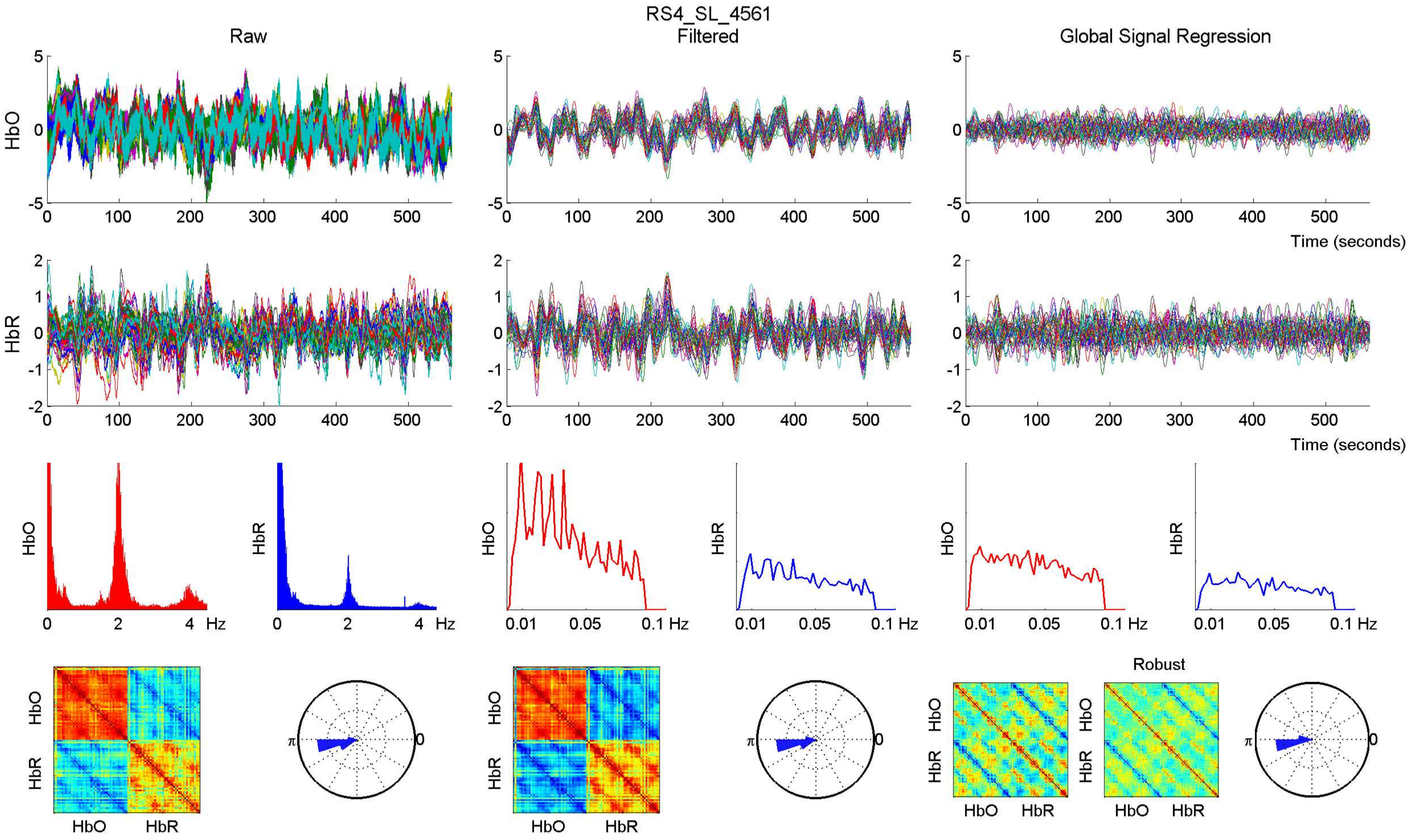

RS4_SL_4561
Raw
Filtered
Global Signal Regression
HbO
HbR
Time (seconds)
Hz
Robust
HbO
HbR

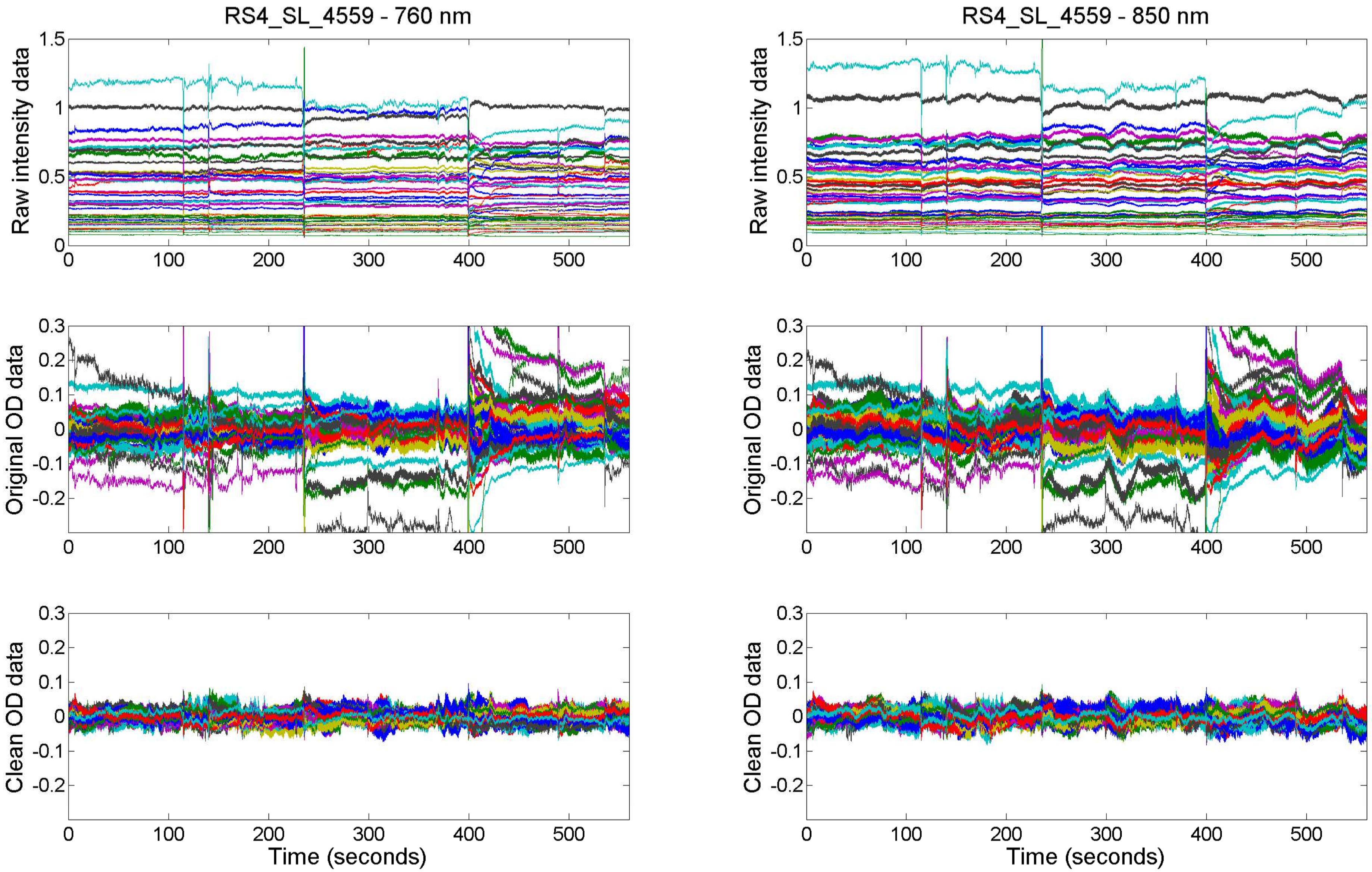

RS4_SL_4559 - 760 nm
RS4_SL_4559 - 850 nm
Raw intensity data
Original OD data
Clean OD data
Time (seconds)

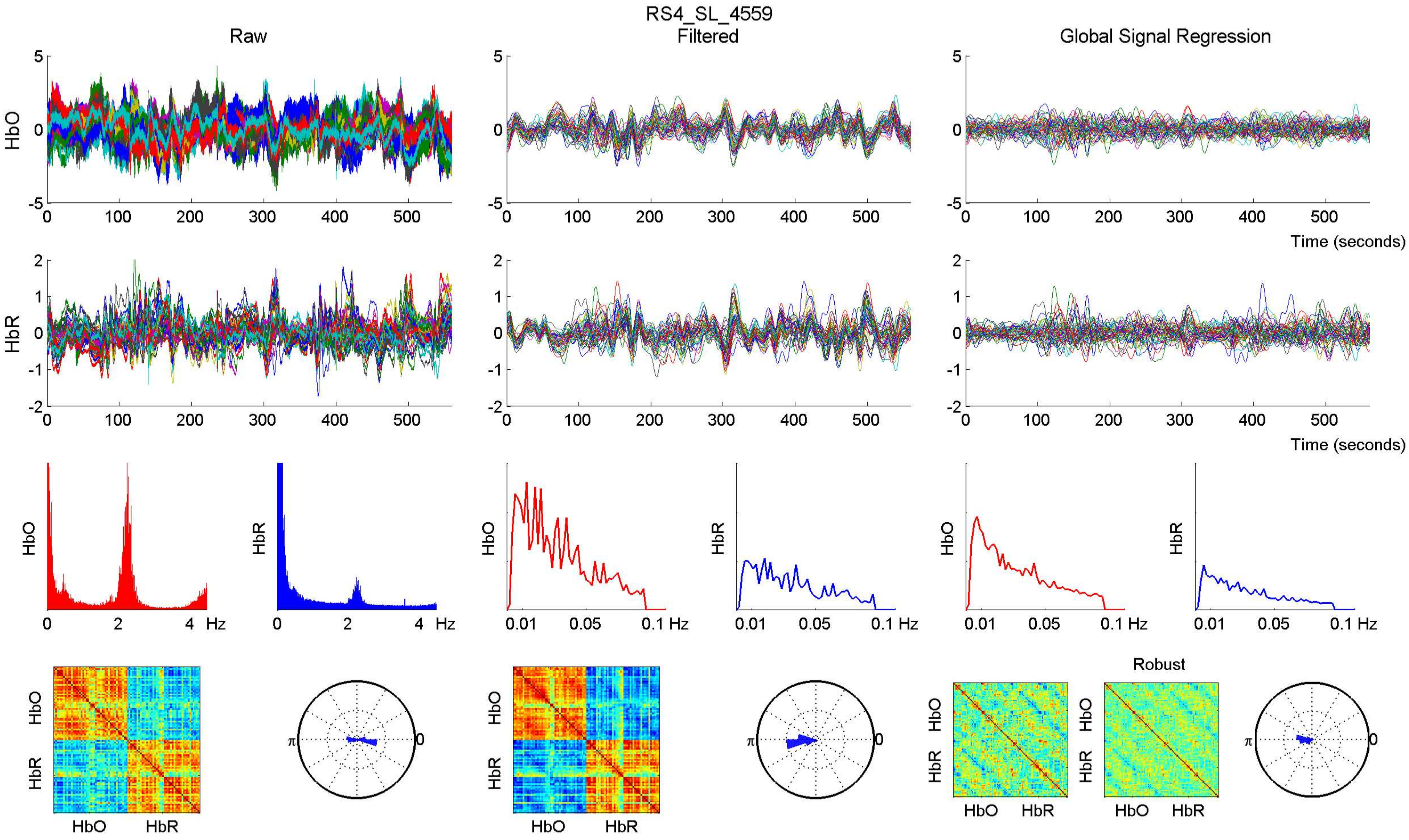
RS4_SL_4559
Raw
Filtered
Global Signal Regression
HbO
HbR
Time (seconds)
Hz
Robust

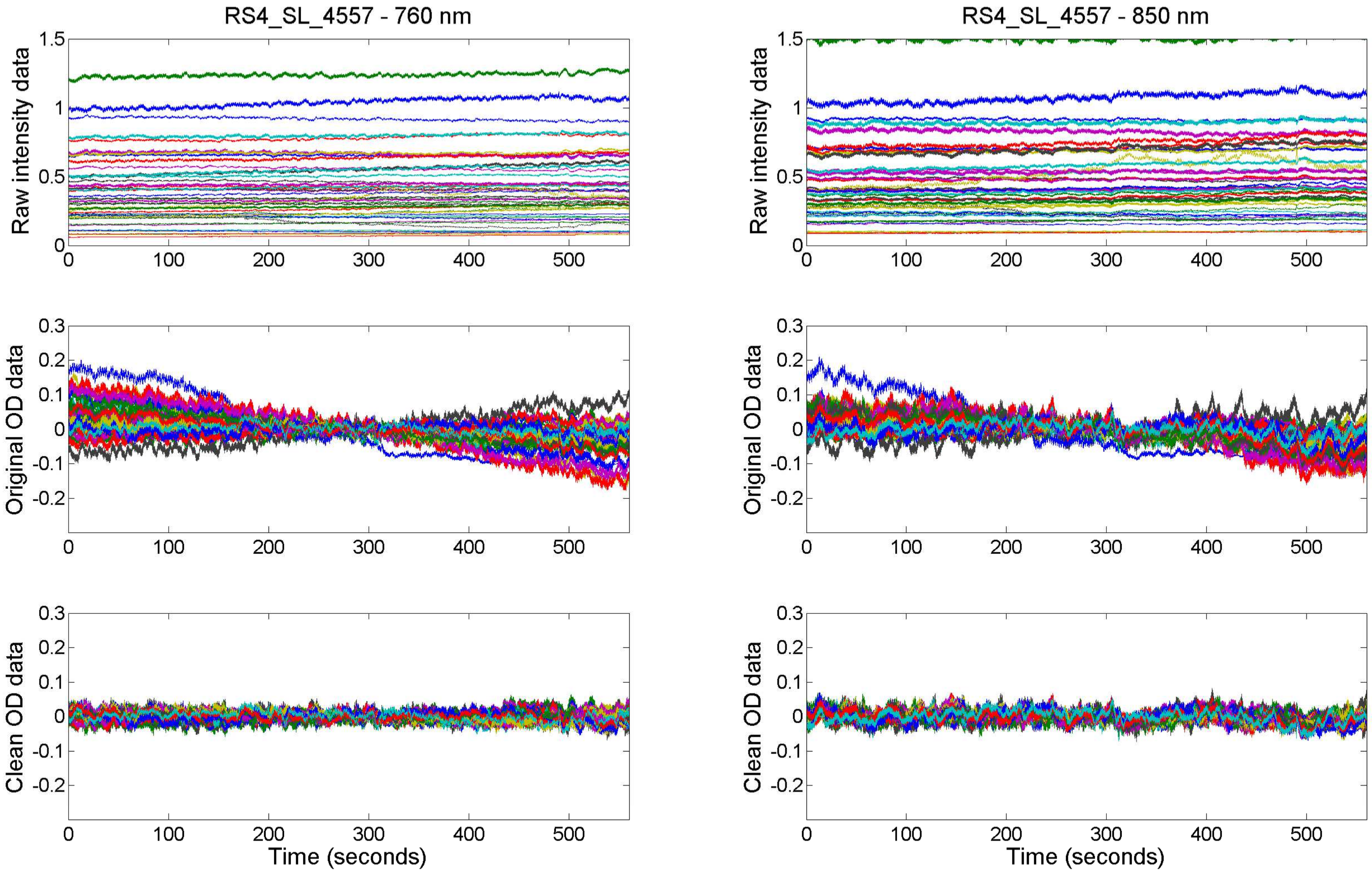
RS4_SL_4557 - 760 nm
RS4_SL_4557 - 850 nm
Raw intensity data
Original OD data
Clean OD data
Time (seconds)
Time (seconds)

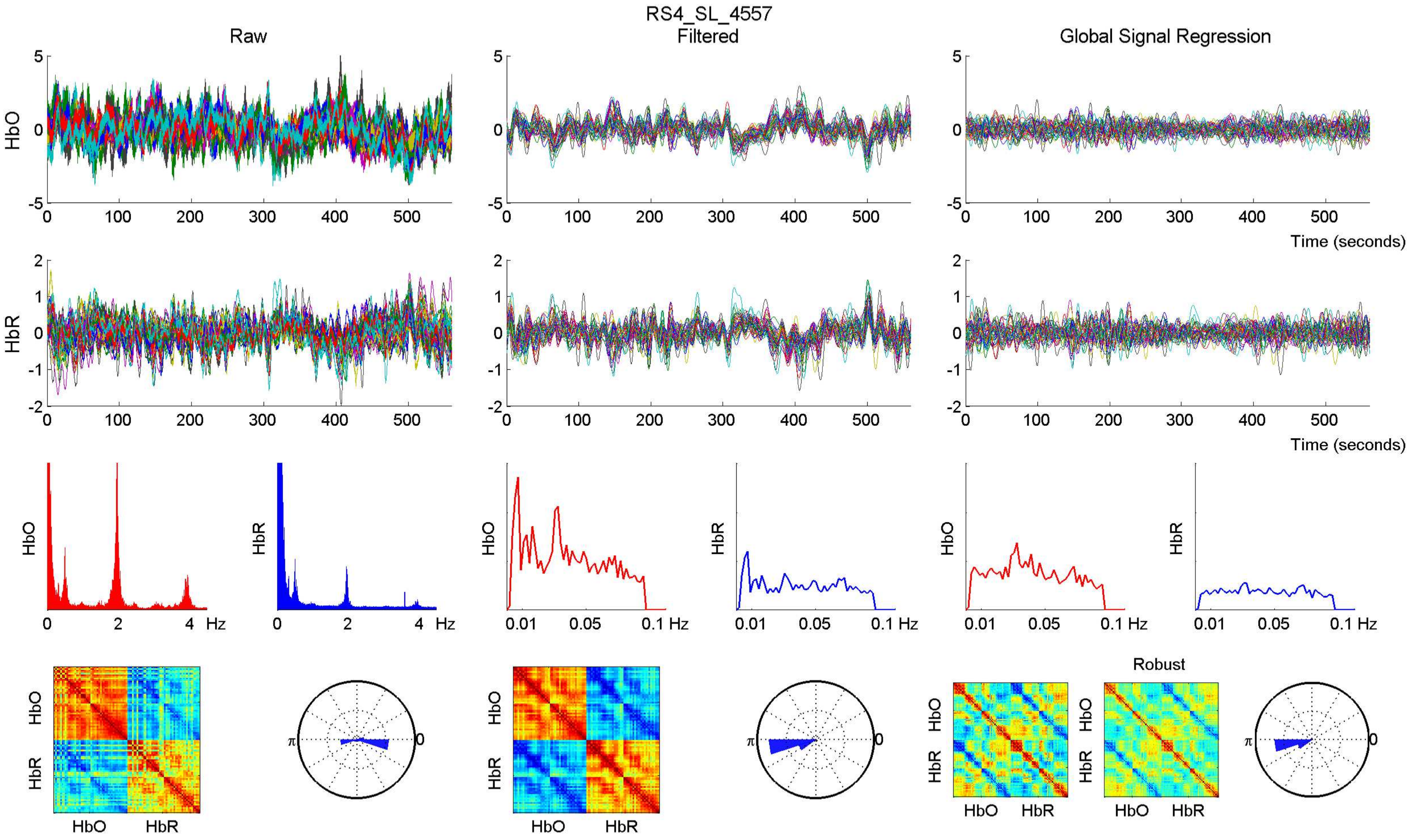

RS4_SL_4557
Raw
Filtered
Global Signal Regression
HbO
HbR
Time (seconds)
Hz
0.01
0.05
0.1 Hz
Robust
HbO
HbR
π
0

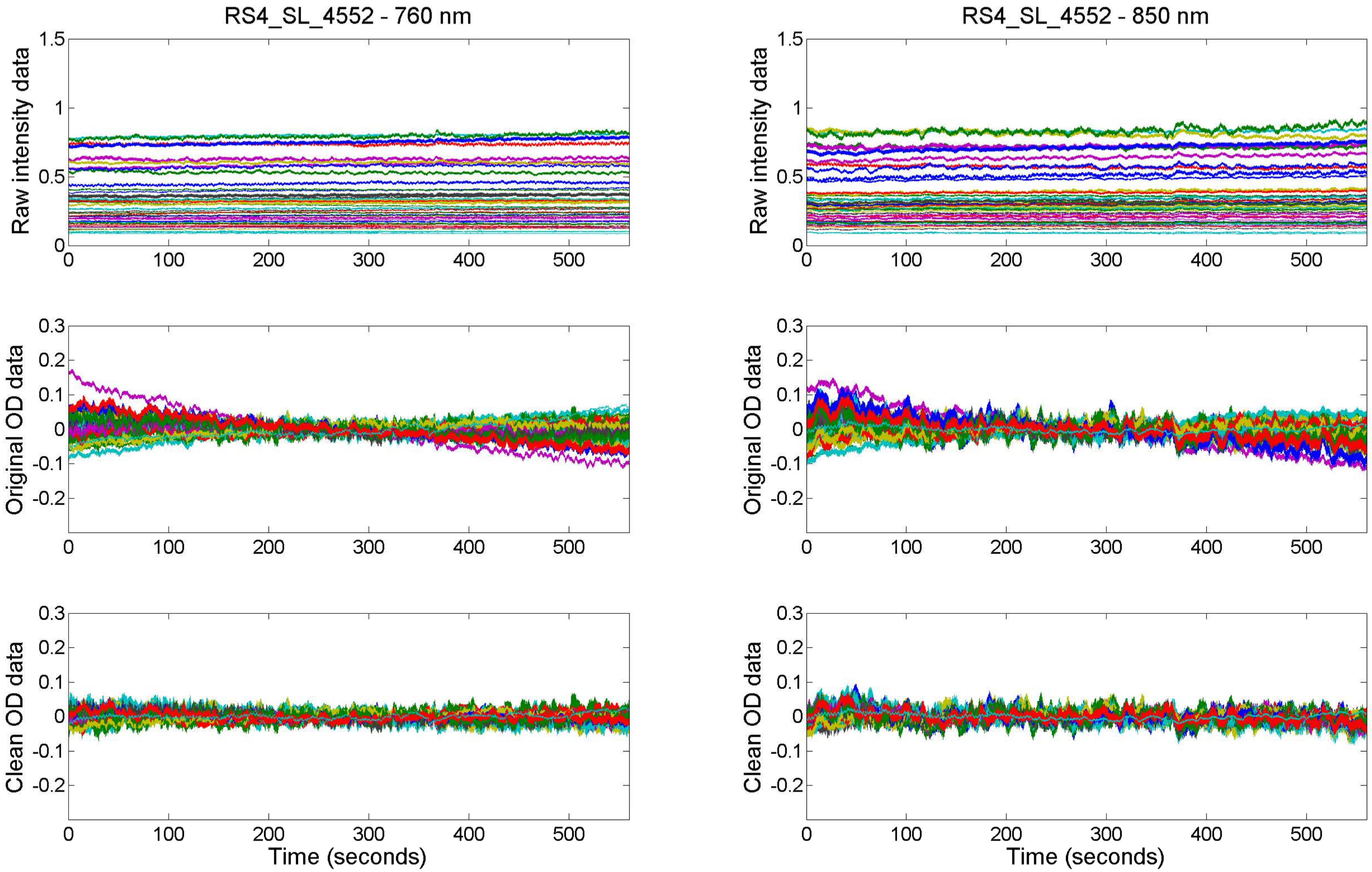

RS4_SL_4552 - 760 nm
RS4_SL_4552 - 850 nm
Raw intensity data
Original OD data
Clean OD data
Time (seconds)

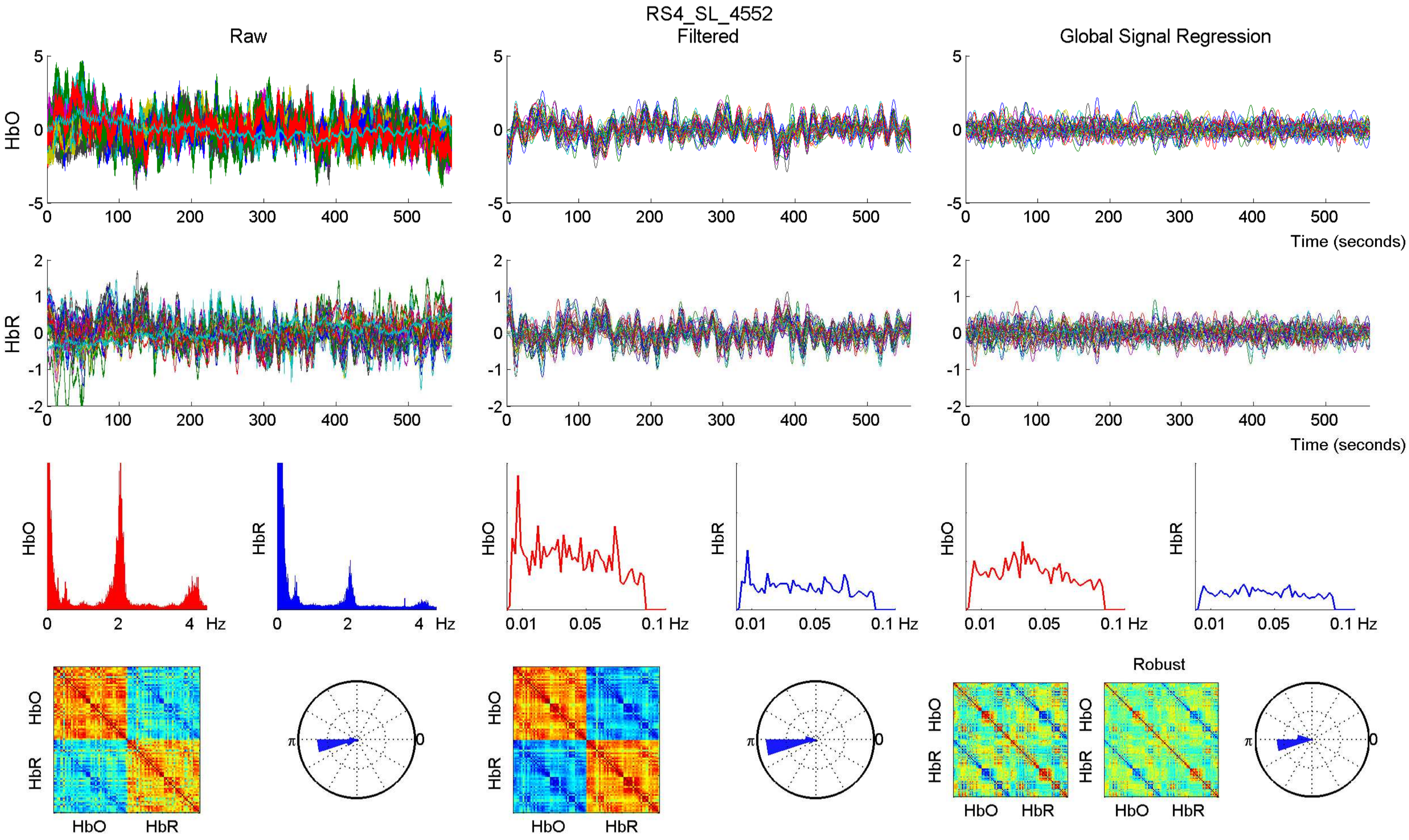

RS4_SL_4552
Raw
Filtered
Global Signal Regression
HbO
HbR
Time (seconds)
Hz
Robust

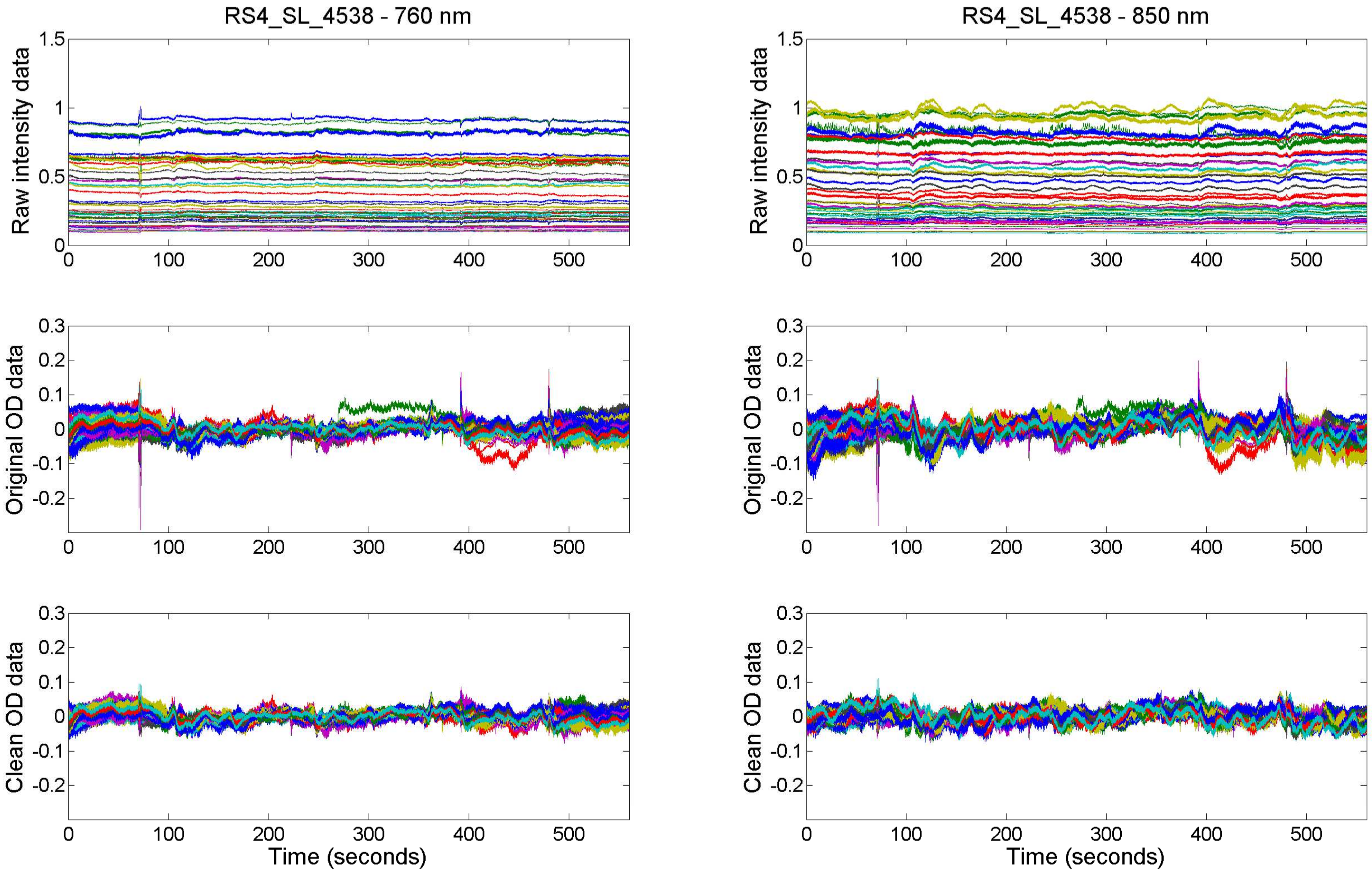
RS4_SL_4538 - 760 nm
RS4_SL_4538 - 850 nm
Raw intensity data
Original OD data
Clean OD data
Time (seconds)
0
0.5
1
1.5
-0.2
-0.1
0.1
0.2
0.3
100
200
300
400
500

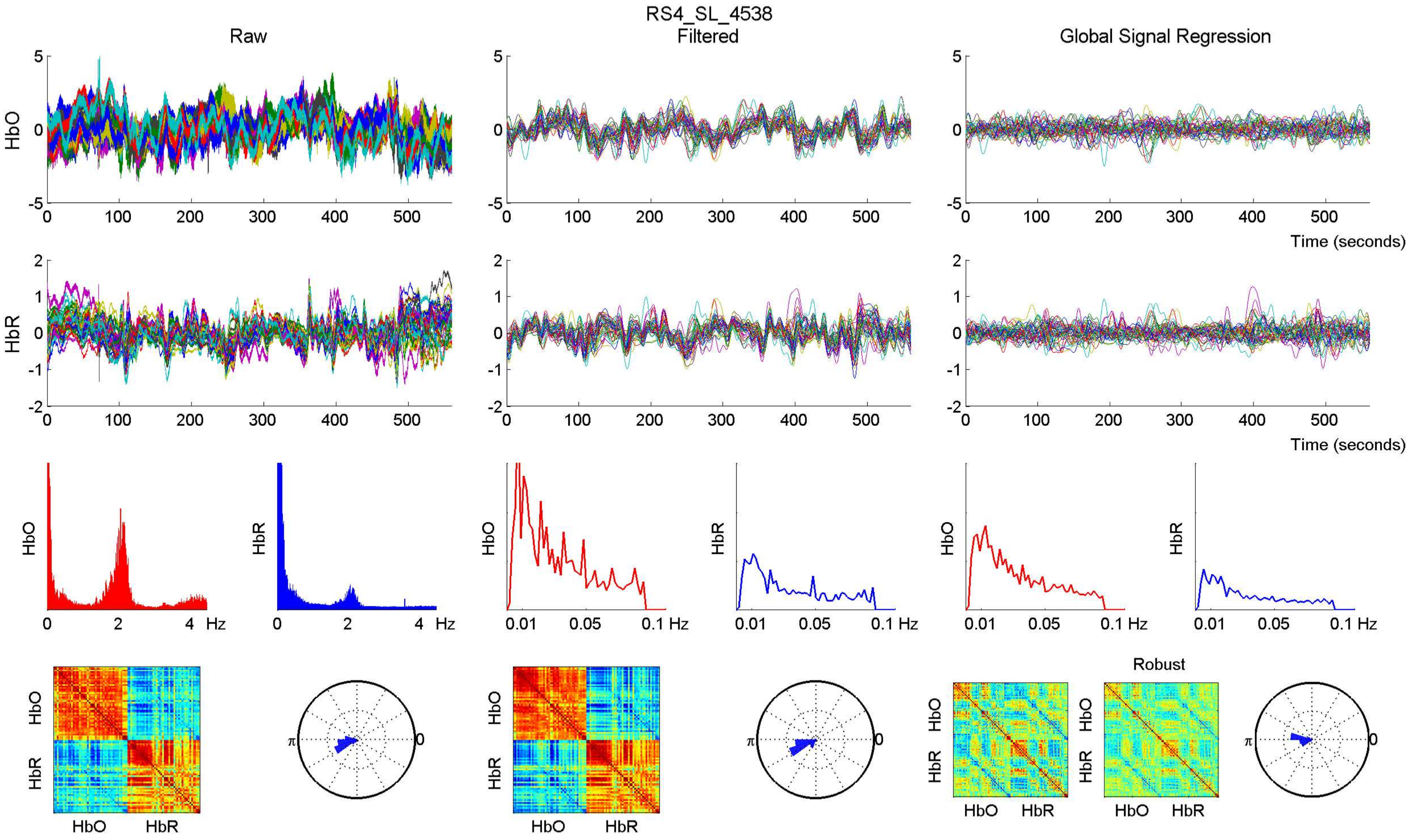

RS4_SL_4538
Raw
Filtered
Global Signal Regression
HbO
HbR
Time (seconds)
Hz
Robust

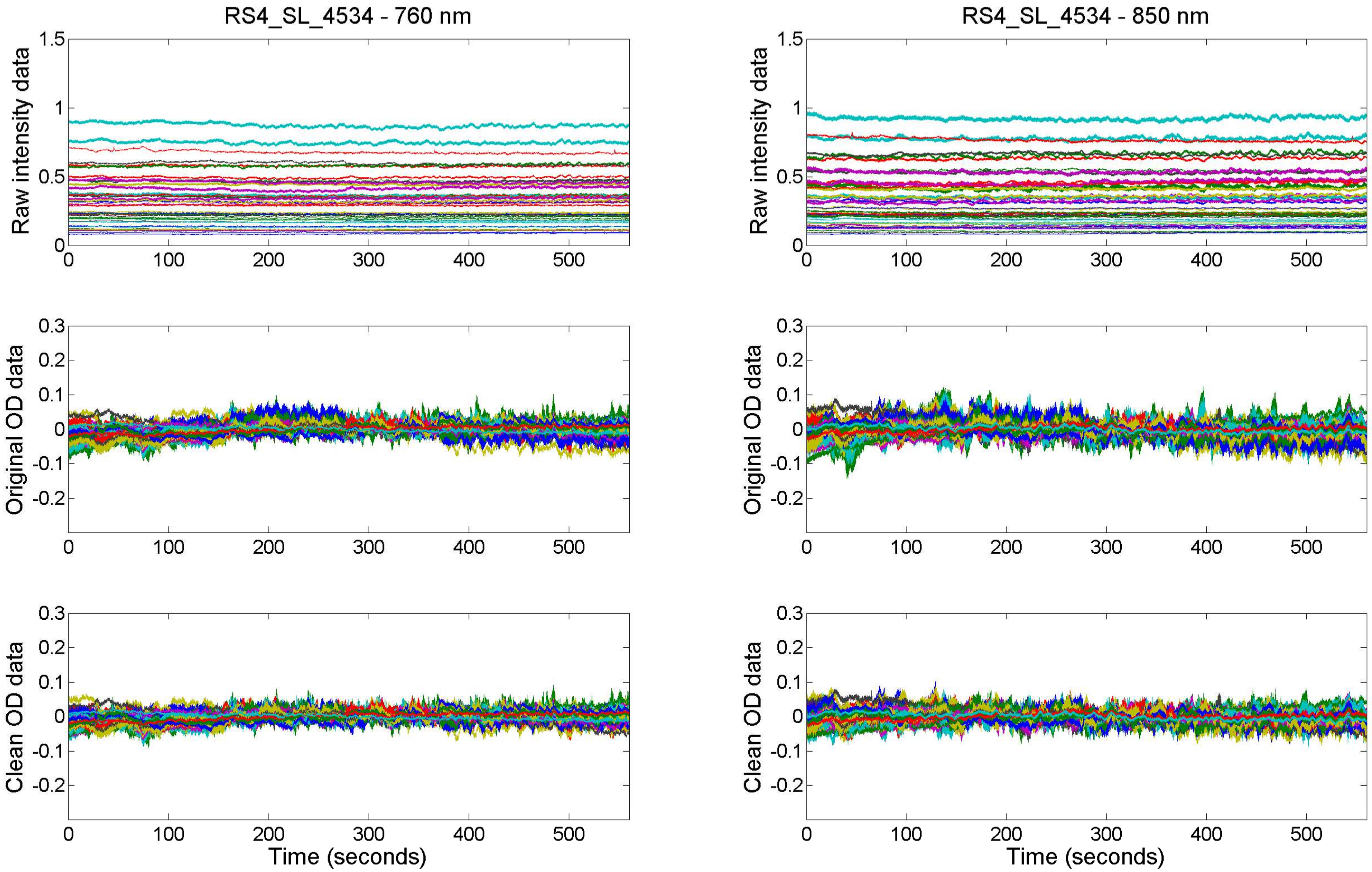

RS4_SL_4534 - 760 nm
RS4_SL_4534 - 850 nm
Raw intensity data
Original OD data
Clean OD data
Time (seconds)

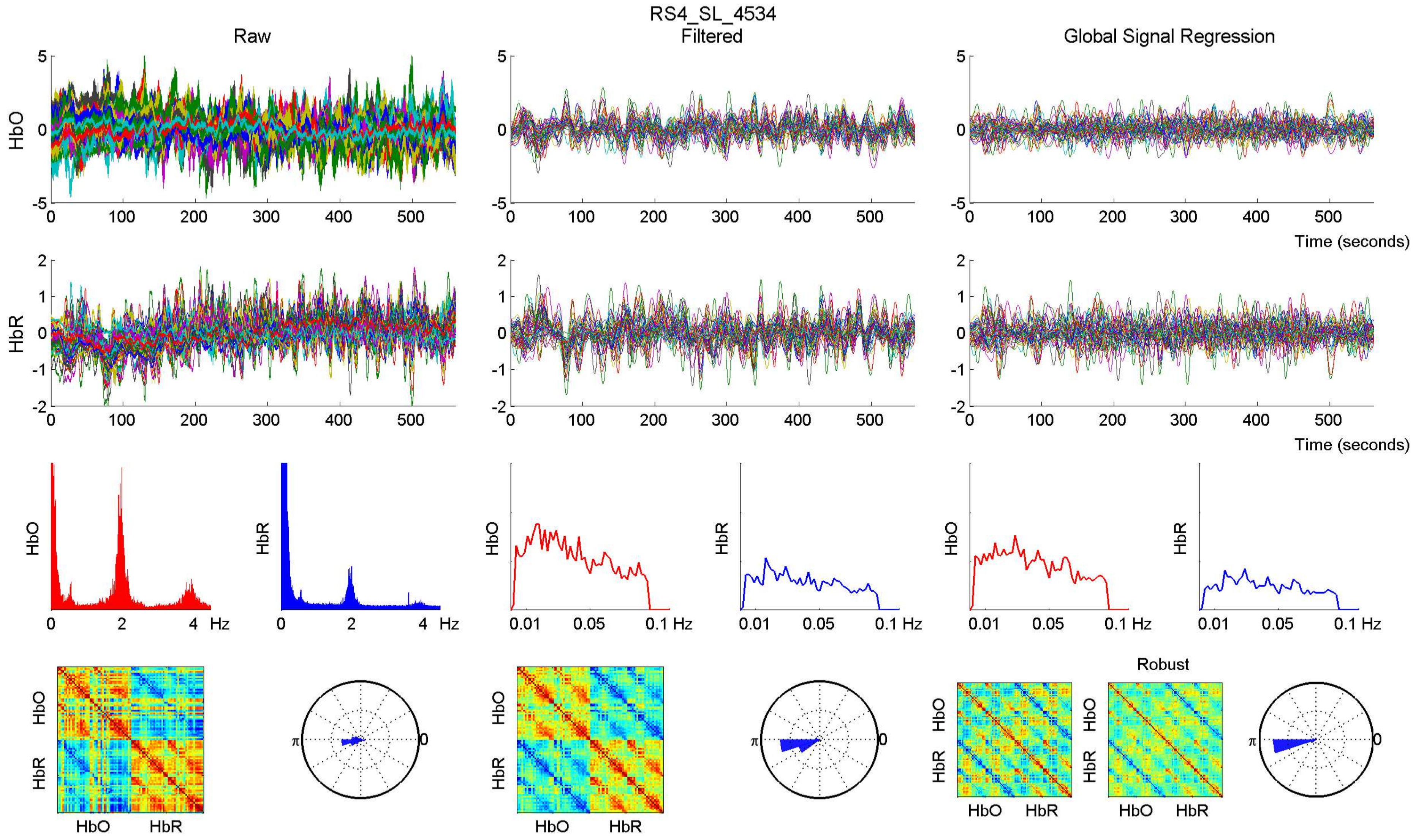
RS4_SL_4534
Raw
Filtered
Global Signal Regression
HbO
HbR
Time (seconds)
Hz
Robust

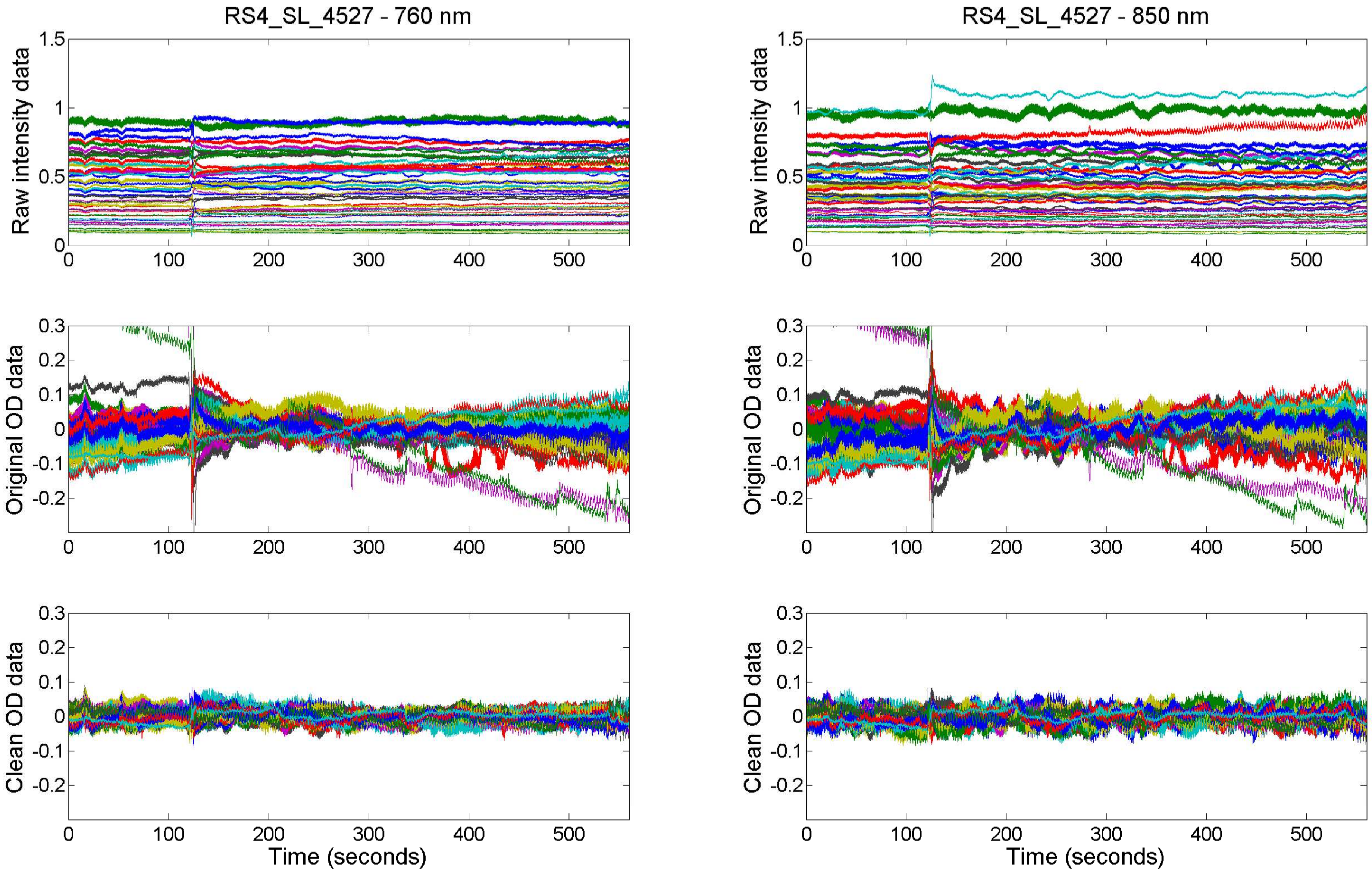

RS4_SL_4527 - 760 nm
RS4_SL_4527 - 850 nm
Raw intensity data
Original OD data
Clean OD data
Time (seconds)

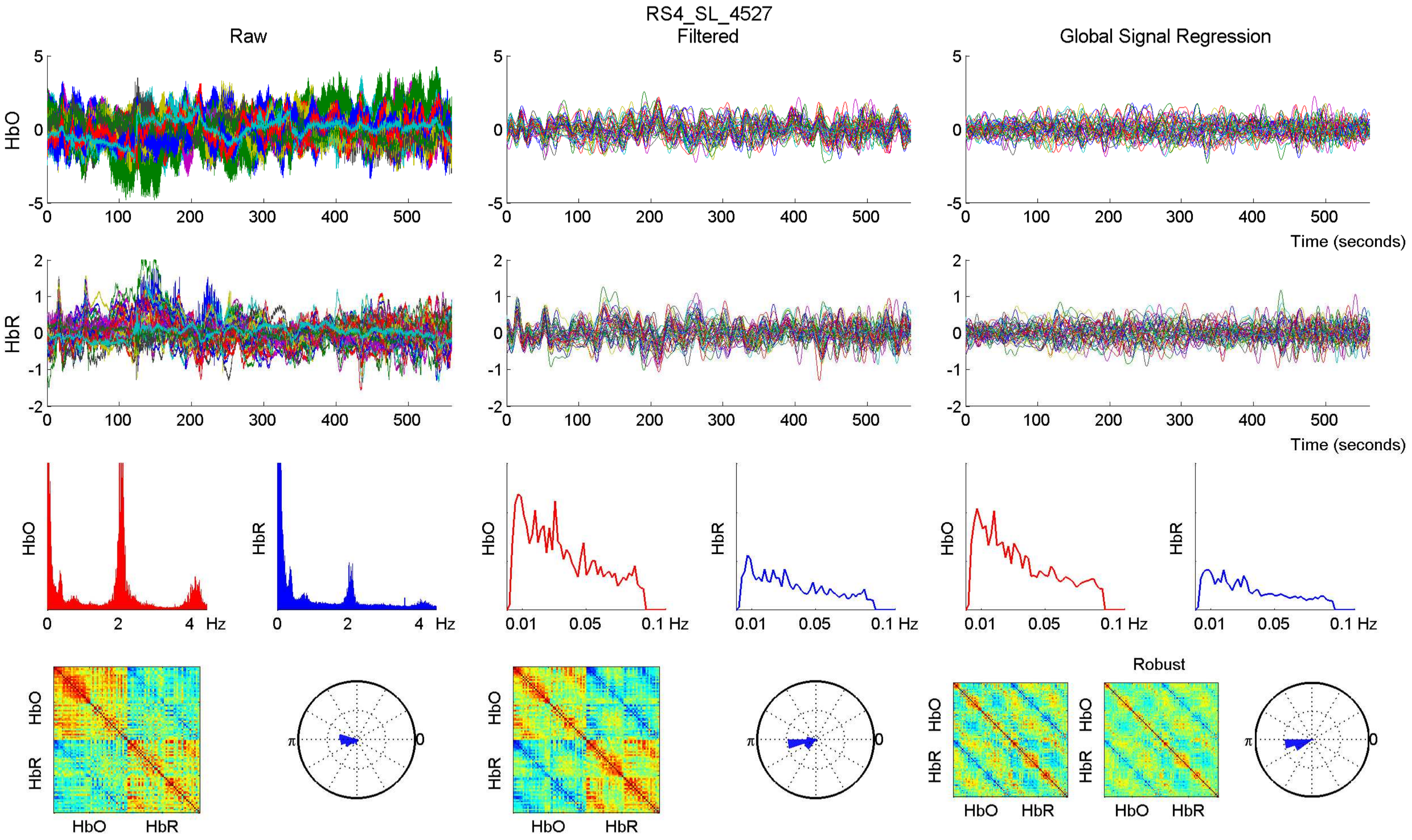

RS4_SL_4527
Raw
Filtered
Global Signal Regression
HbO
HbR
Time (seconds)
Hz
Robust

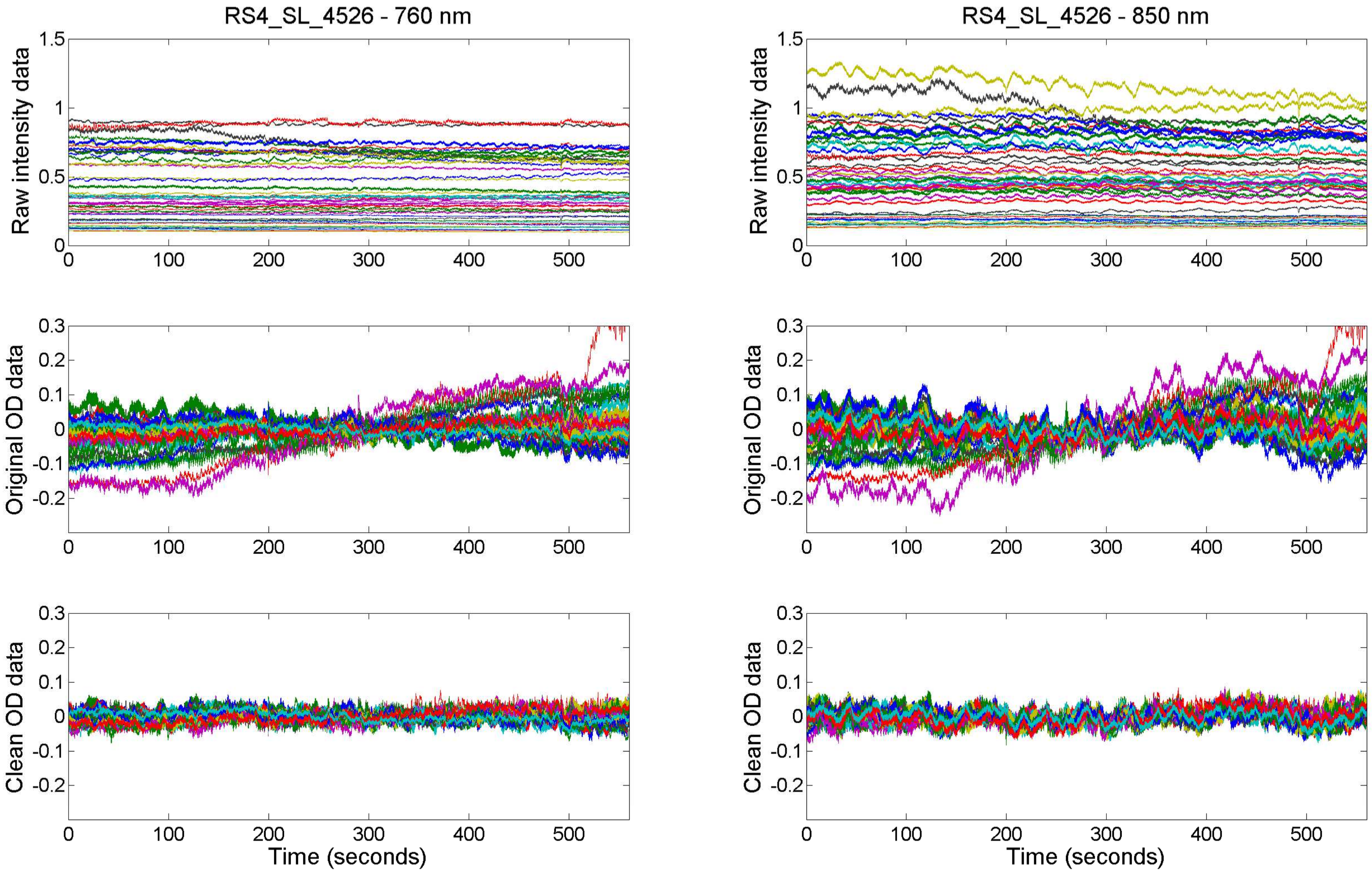

RS4_SL_4526 - 760 nm
RS4_SL_4526 - 850 nm
Raw intensity data
Original OD data
Clean OD data
Time (seconds)

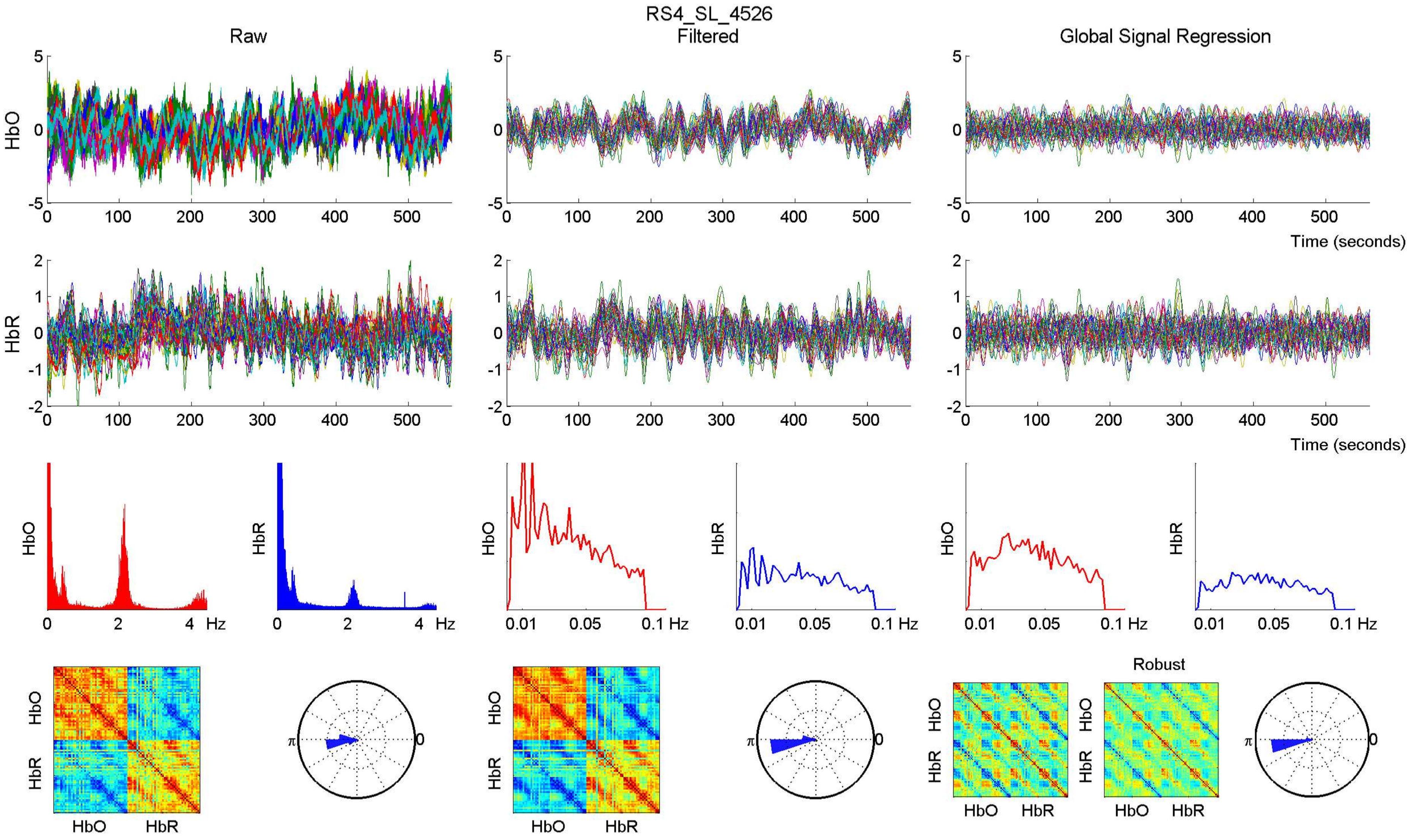
RS4_SL_4526
Raw
Filtered
Global Signal Regression
HbO
HbR
Time (seconds)
Hz
Robust

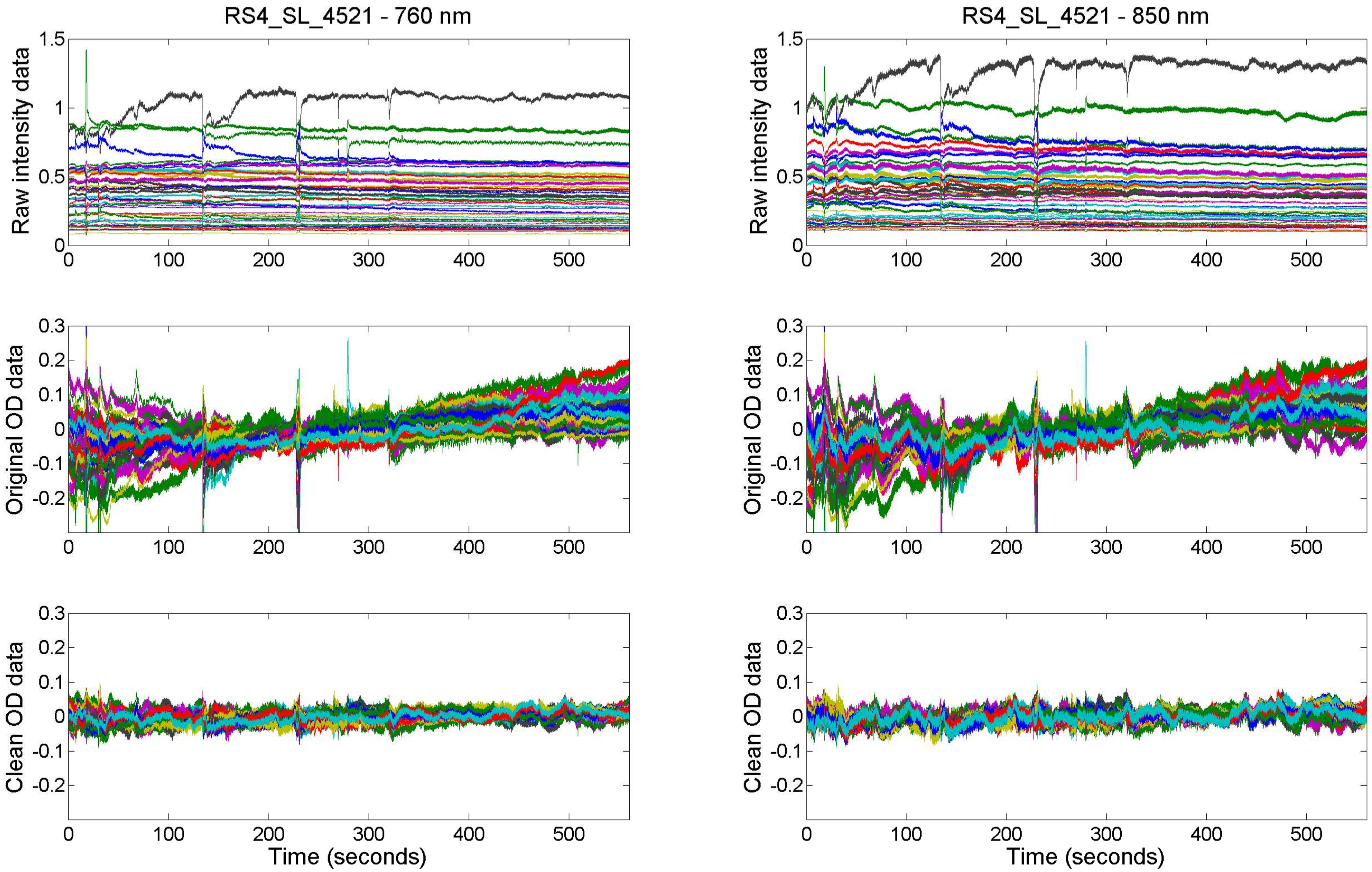

RS4_SL_4521 - 760 nm
RS4_SL_4521 - 850 nm
Raw intensity data
Original OD data
Clean OD data
Time (seconds)

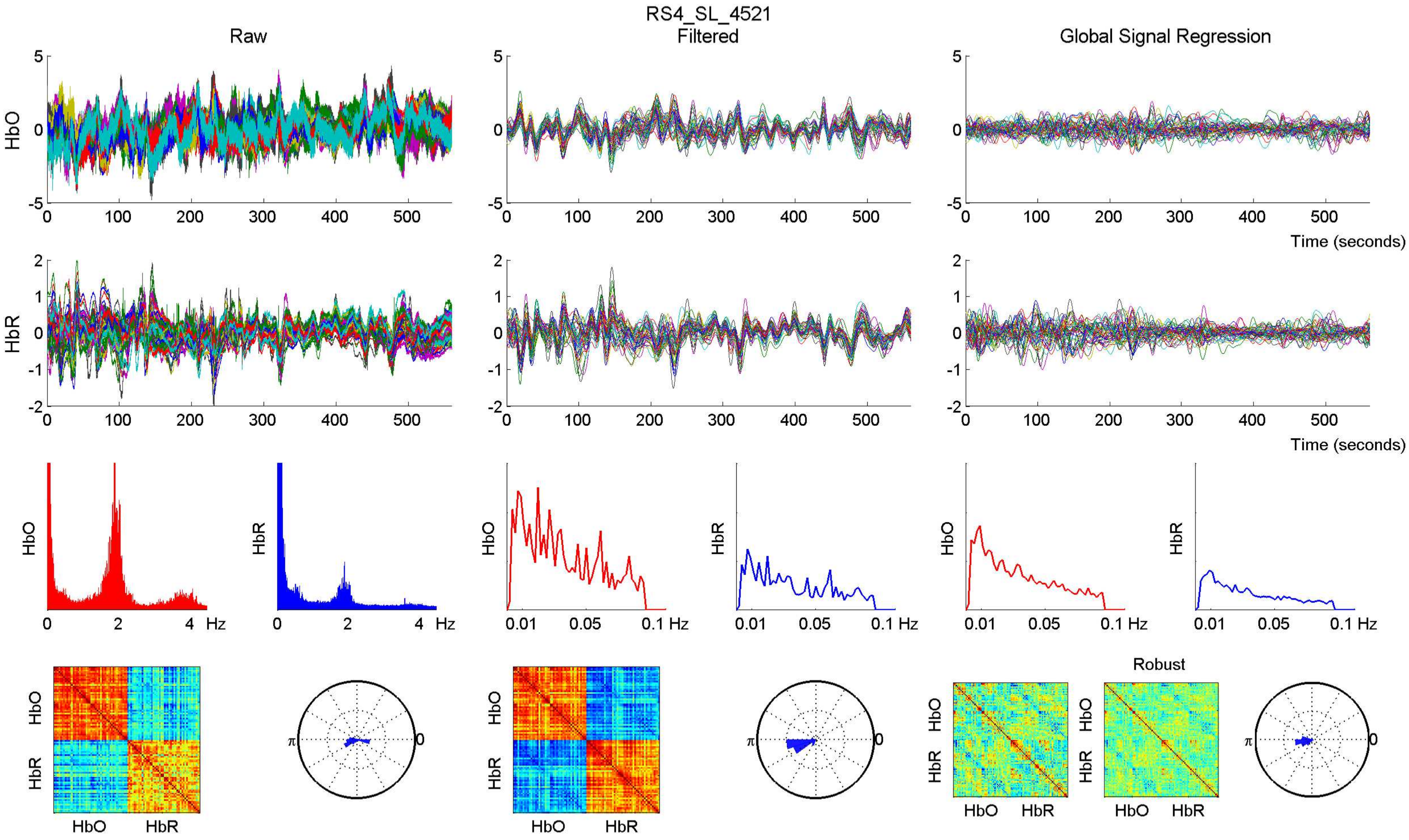

RS4_SL_4521
Raw
Filtered
Global Signal Regression
HbO
HbR
Time (seconds)
Hz
Robust
HbO
HbR

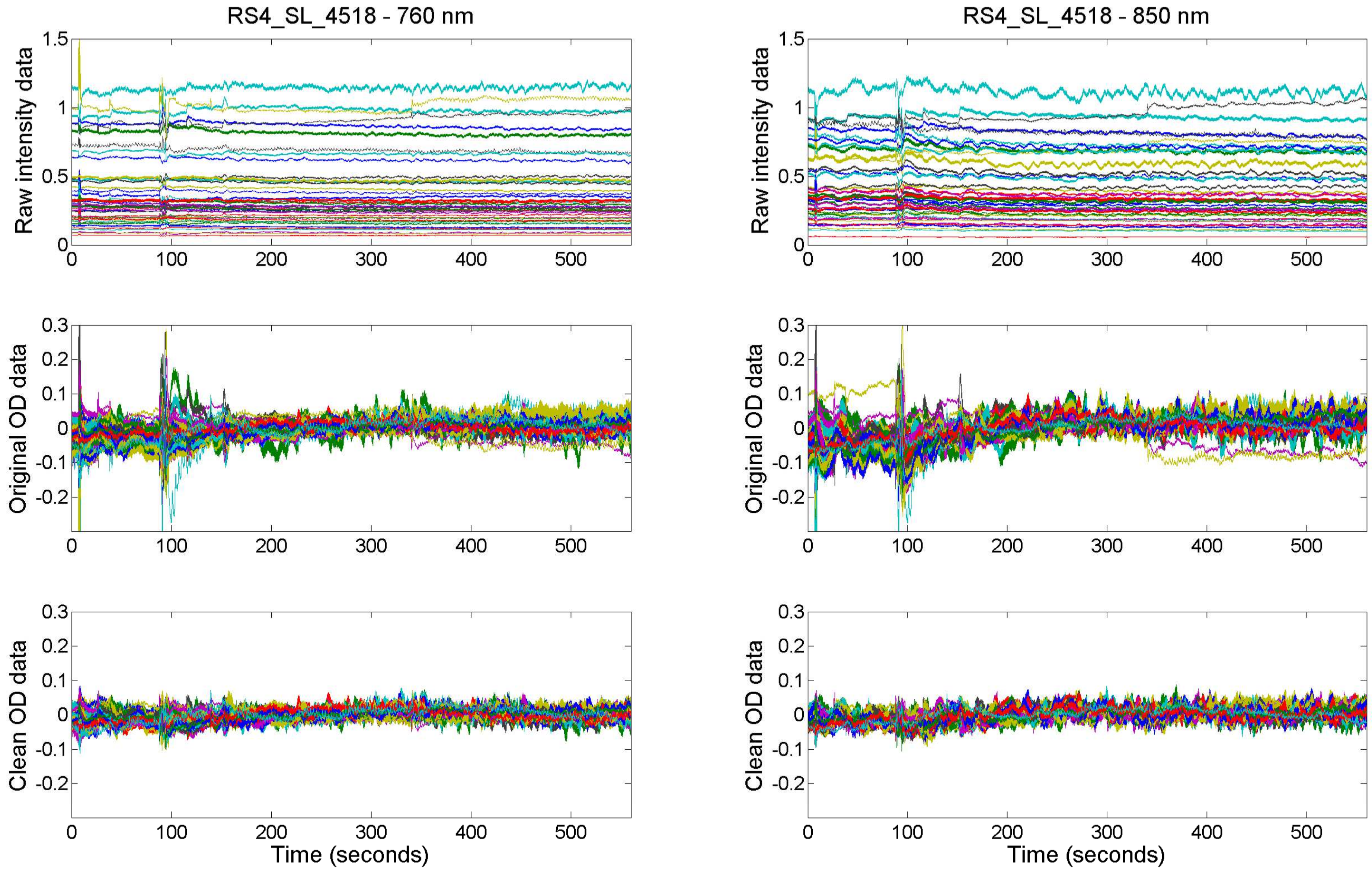
RS4_SL_4518 - 760 nm
RS4_SL_4518 - 850 nm
Raw intensity data
Original OD data
Clean OD data
Time (seconds)
0
0.5
1
1.5
0.3
0.2
0.1
-0.1
-0.2
100
200
300
400
500

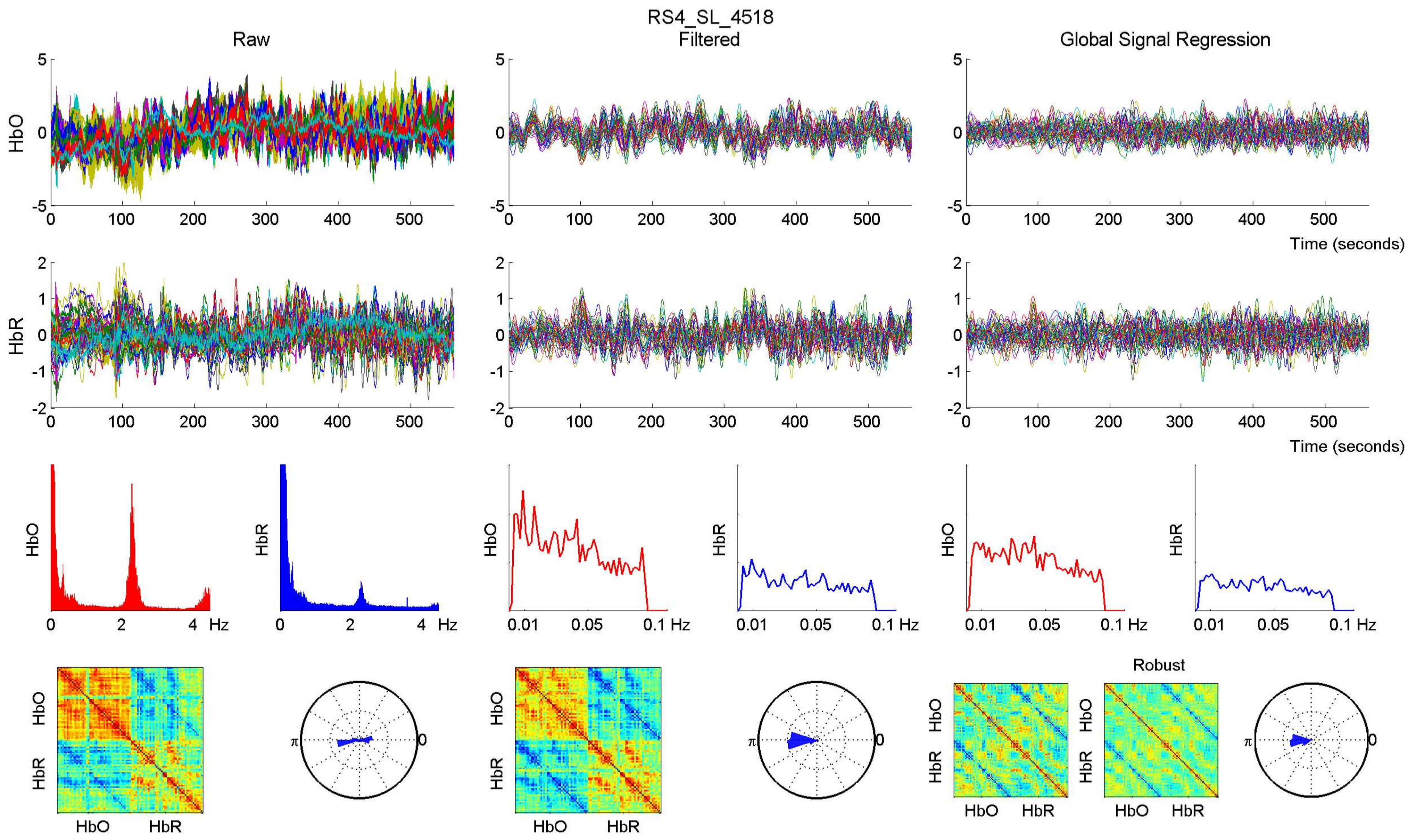
RS4_SL_4518
Raw
Filtered
Global Signal Regression
HbO
HbR
Time (seconds)
Hz
0.01
0.05
0.1 Hz
Robust
π
0

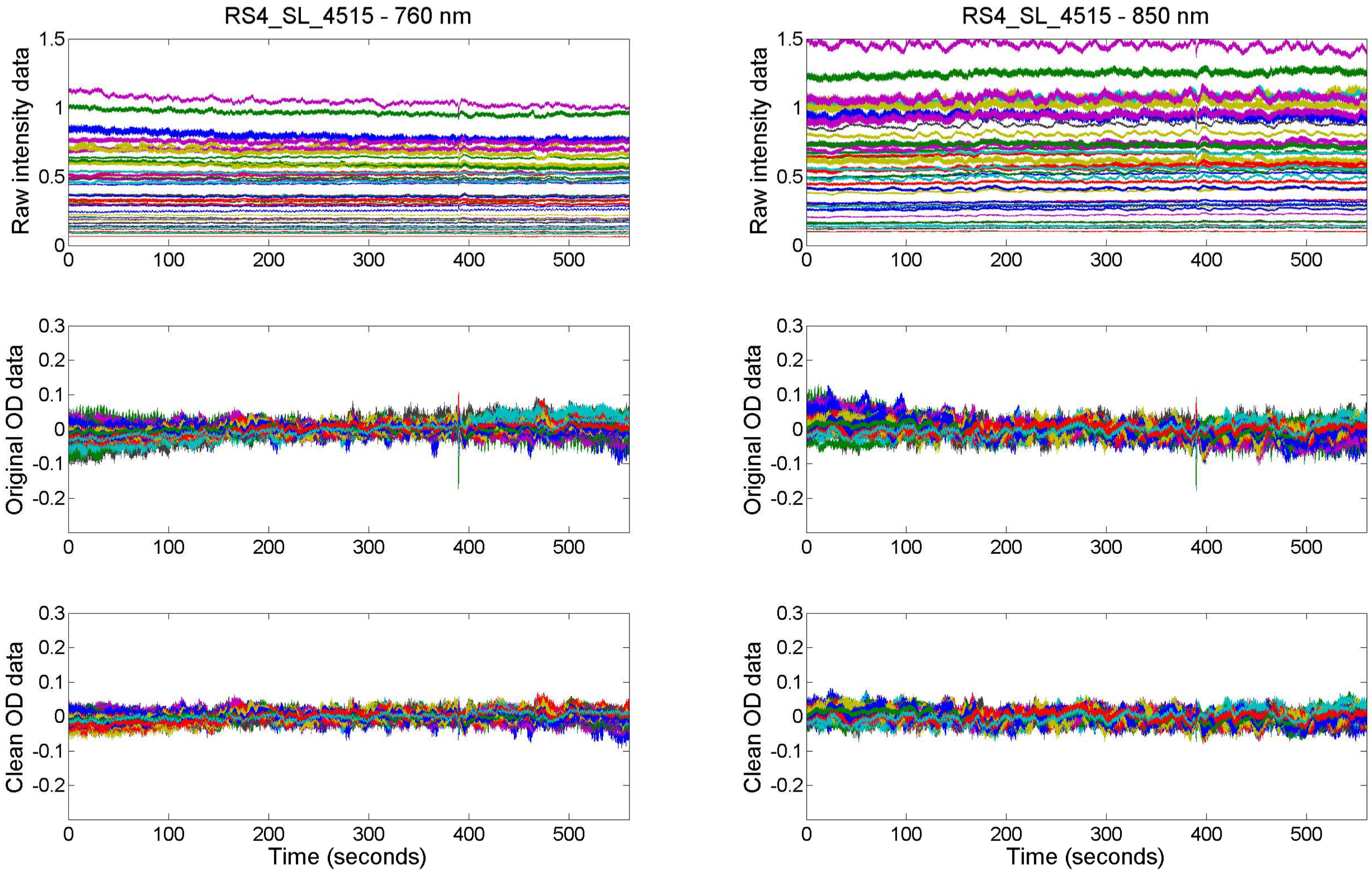

RS4_SL_4515 - 760 nm
RS4_SL_4515 - 850 nm
Raw intensity data
Original OD data
Clean OD data
Time (seconds)

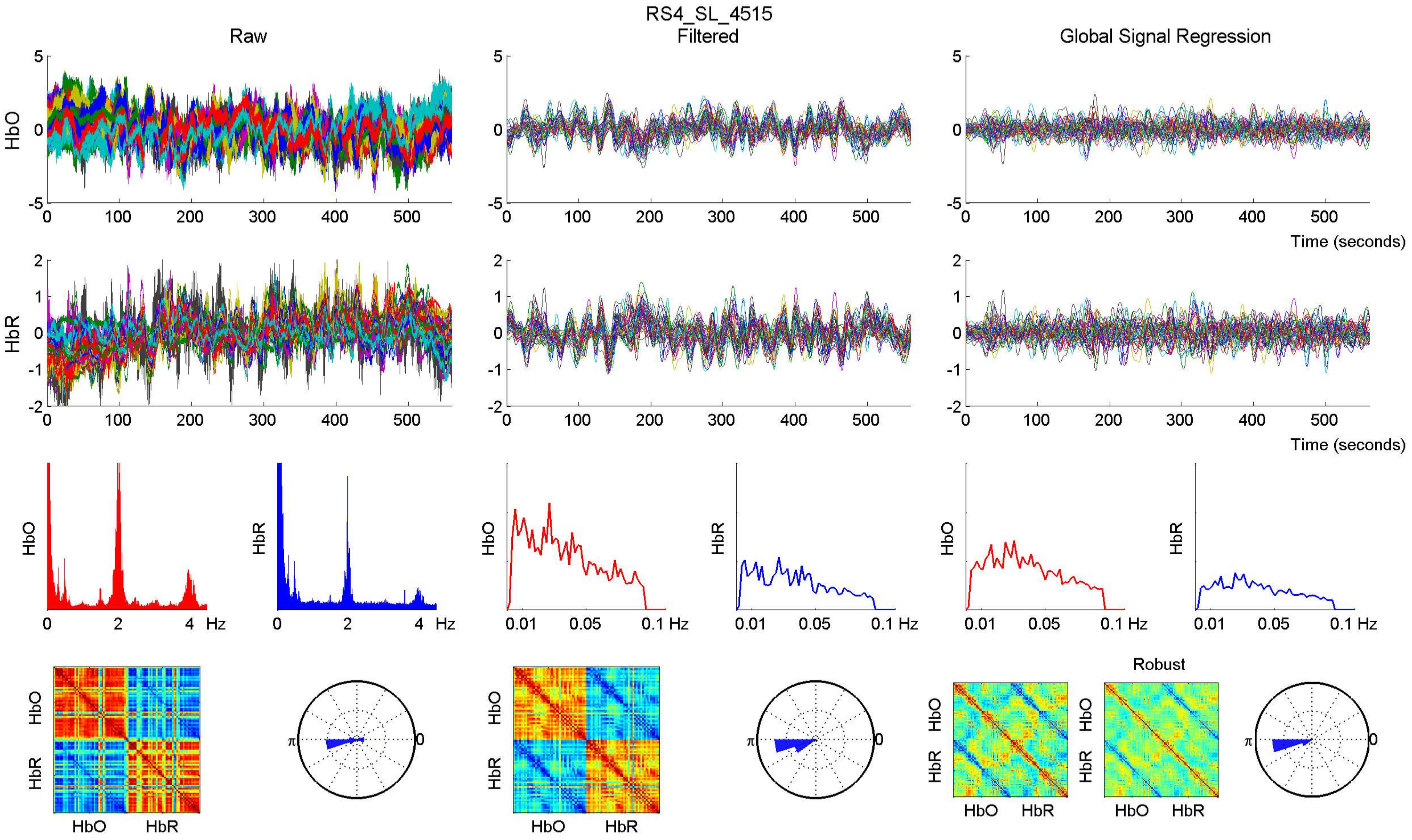
RS4_SL_4515
Raw
Filtered
Global Signal Regression
HbO
HbR
Time (seconds)
Hz
Robust

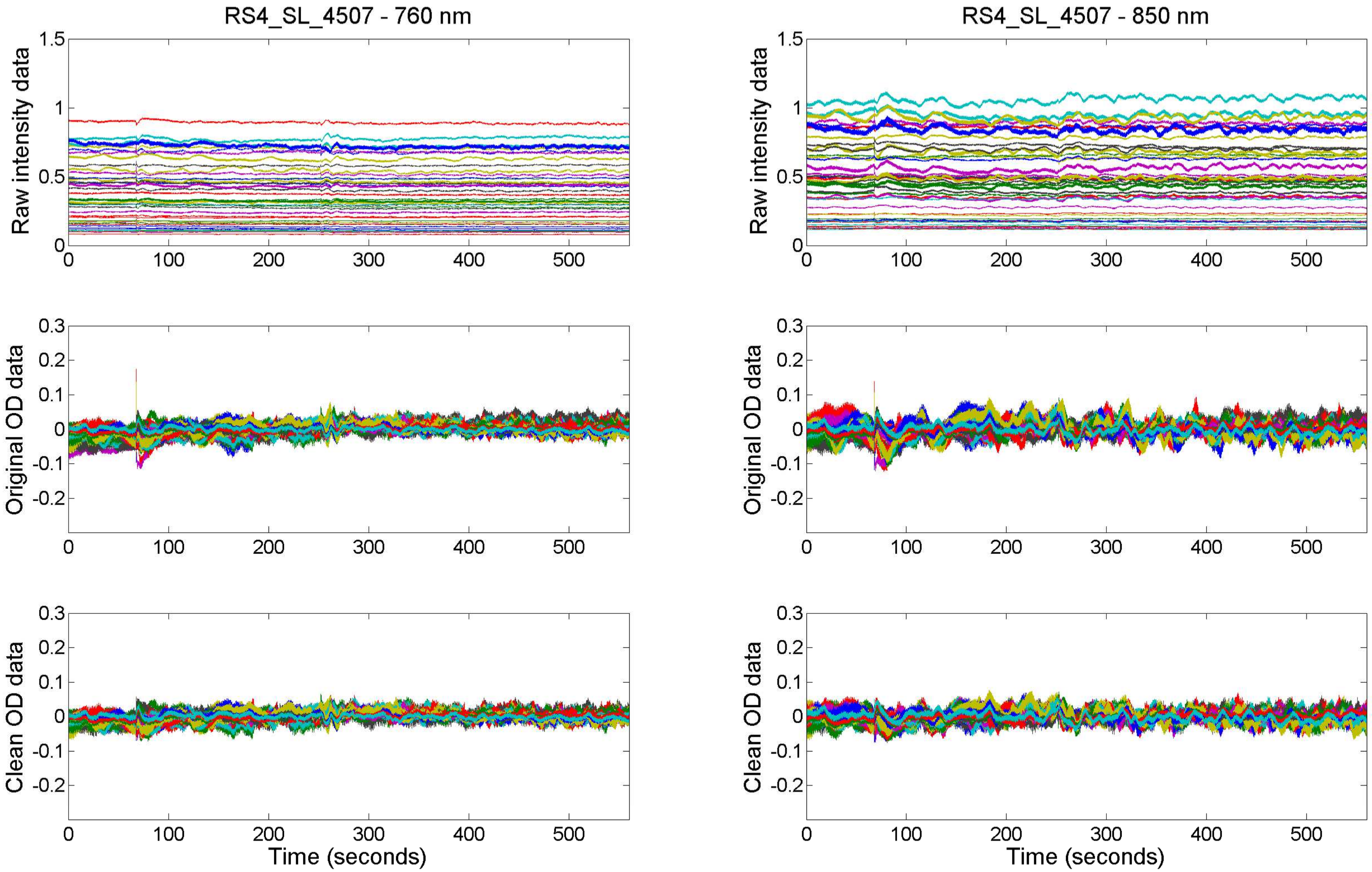
RS4_SL_4507 - 760 nm
RS4_SL_4507 - 850 nm
Raw intensity data
Original OD data
Clean OD data
Time (seconds)

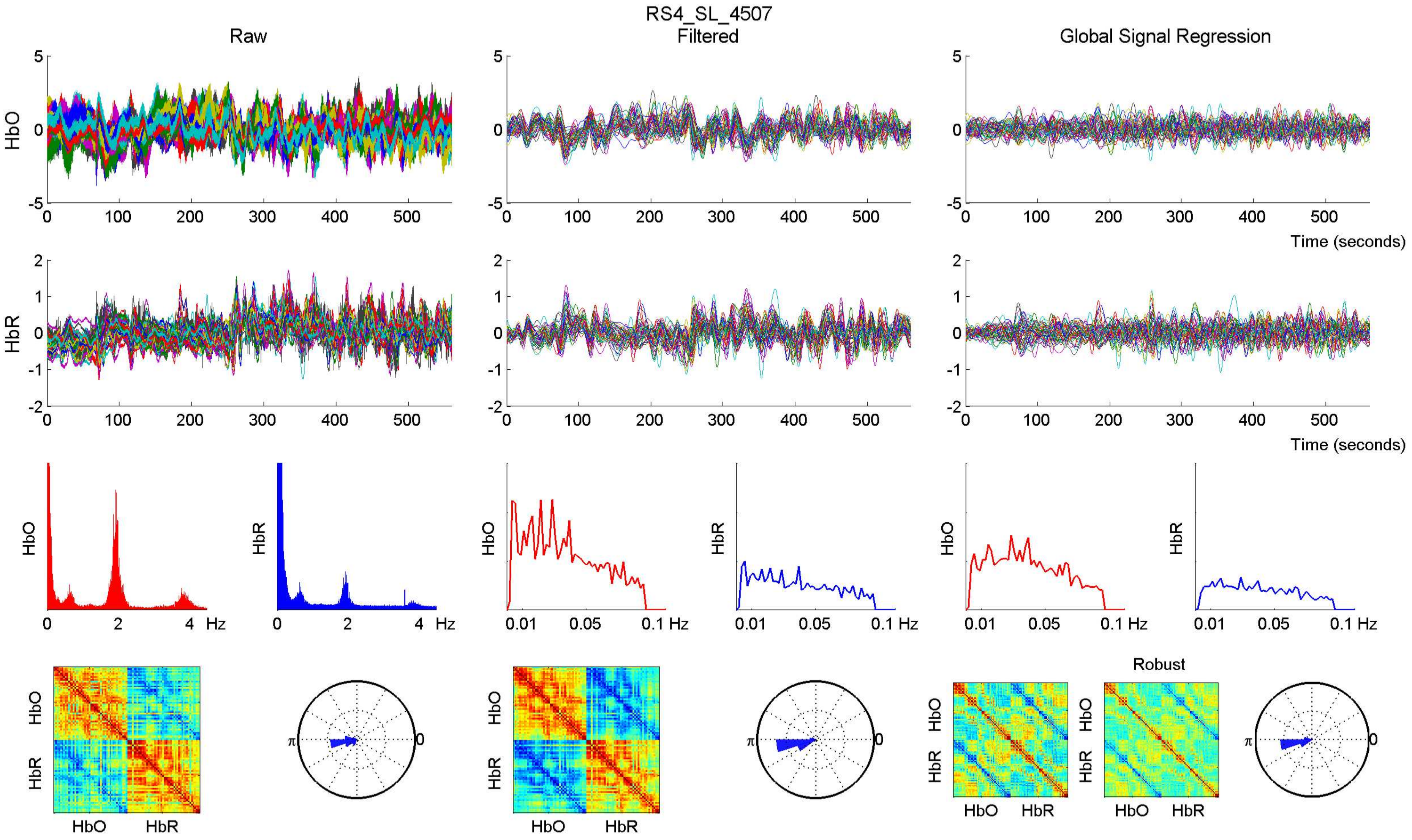

RS4_SL_4507
Raw
Filtered
Global Signal Regression
HbO
HbR
Time (seconds)
Hz
Robust
HbO
HbR

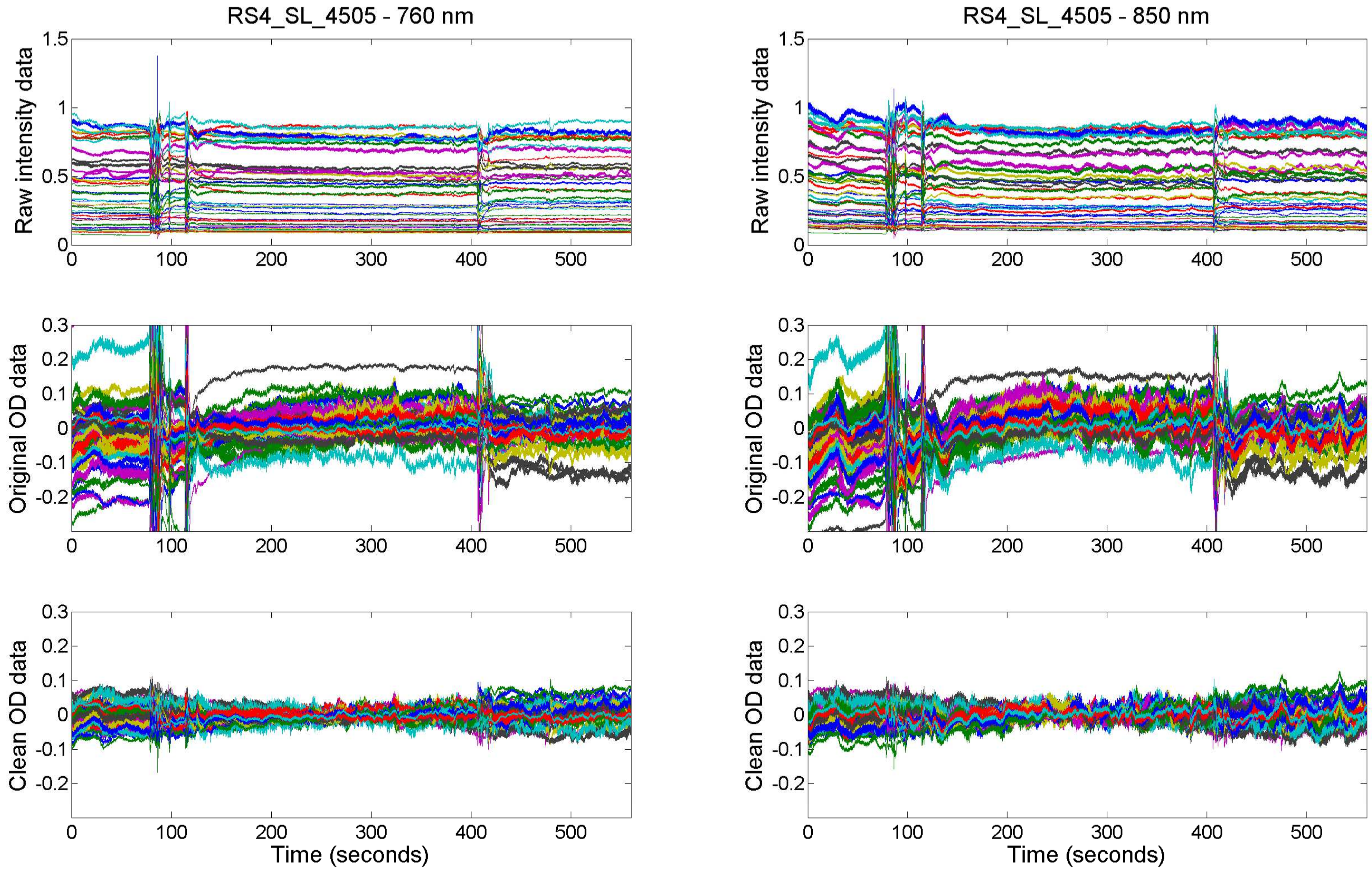

RS4_SL_4505 - 760 nm
RS4_SL_4505 - 850 nm
Raw intensity data
Original OD data
Clean OD data
Time (seconds)

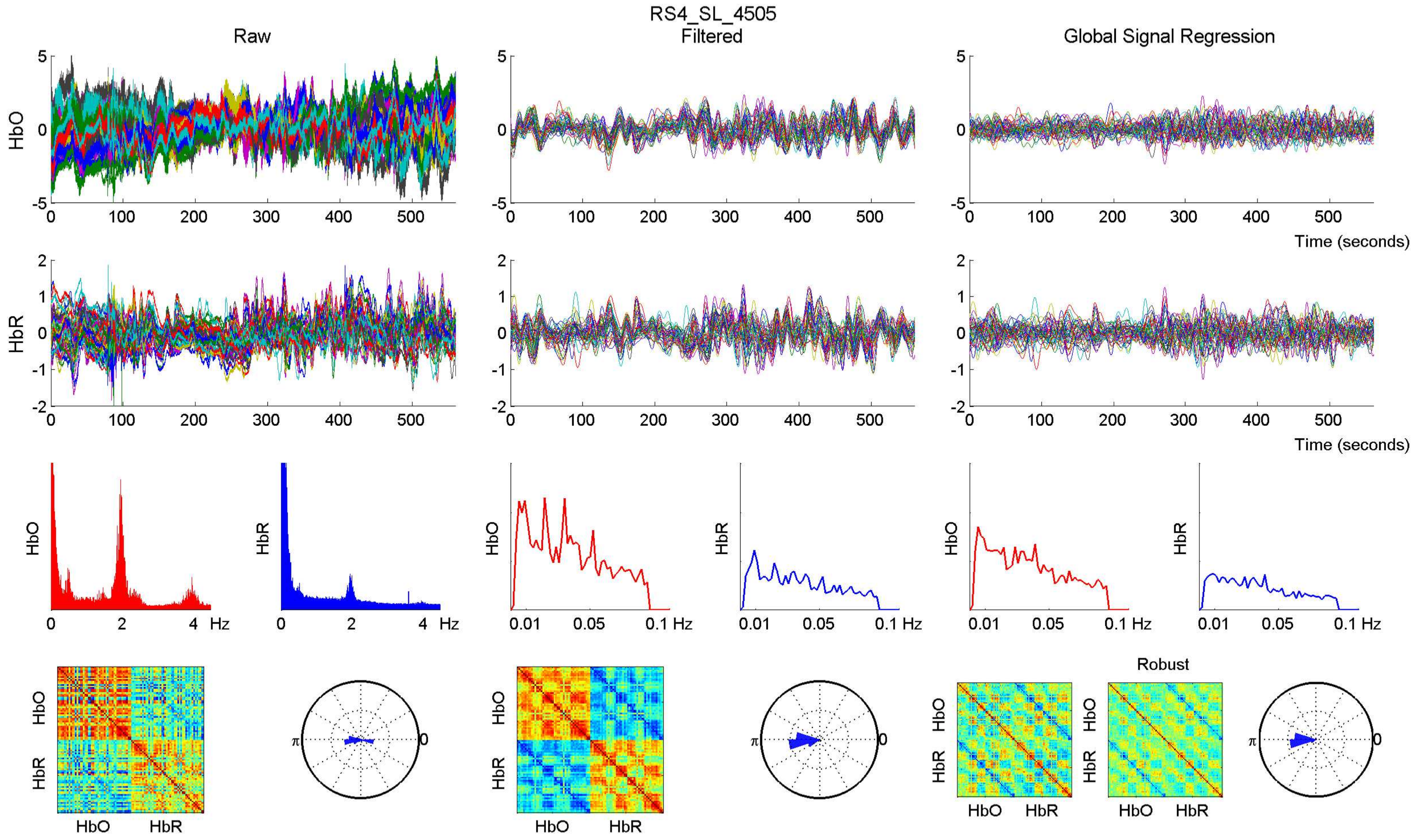

RS4_SL_4505
Raw
Filtered
Global Signal Regression
HbO
HbR
Time (seconds)
Hz
Robust

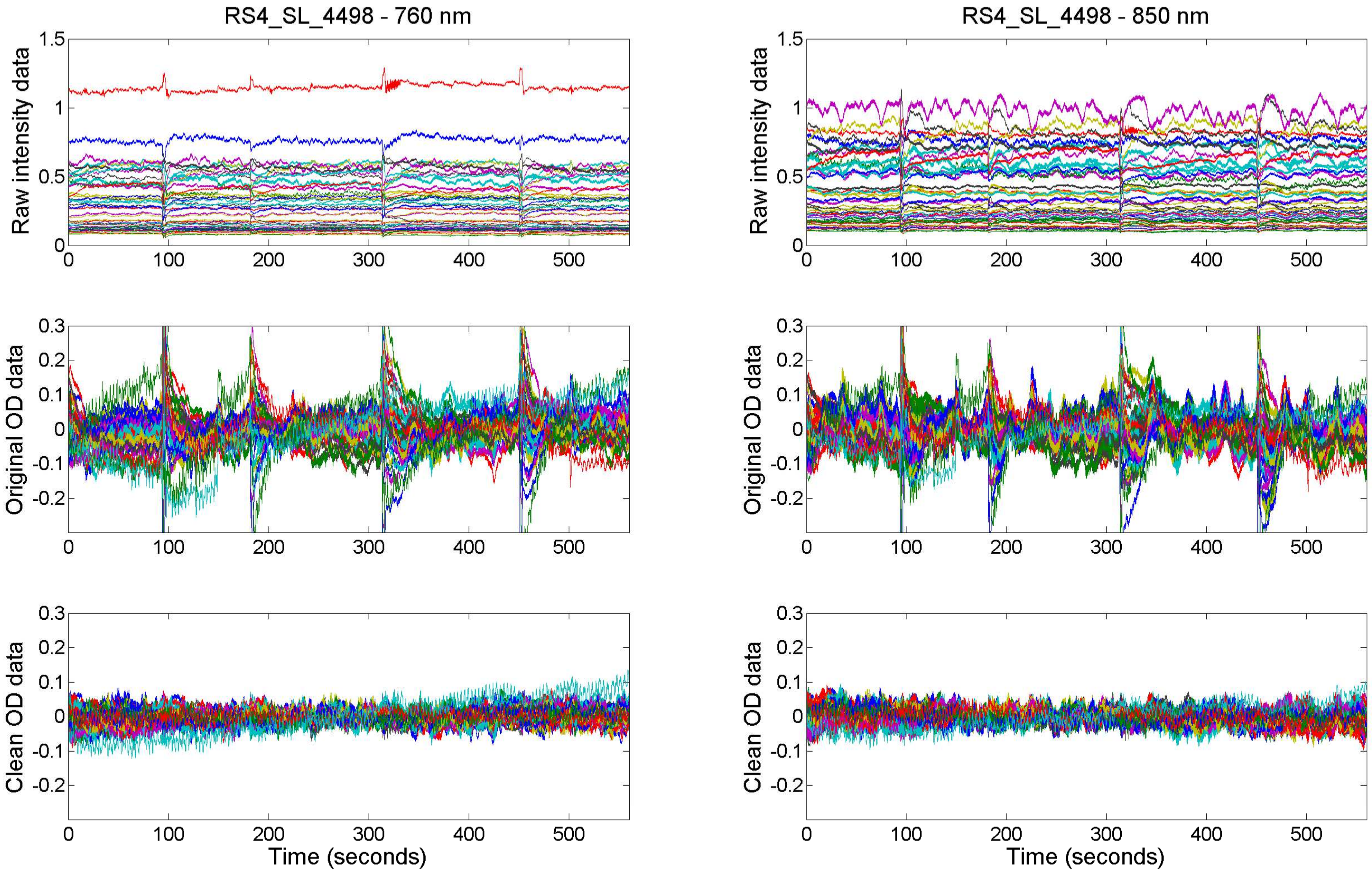

RS4_SL_4498 - 760 nm
RS4_SL_4498 - 850 nm
Raw intensity data
Original OD data
Clean OD data
Time (seconds)

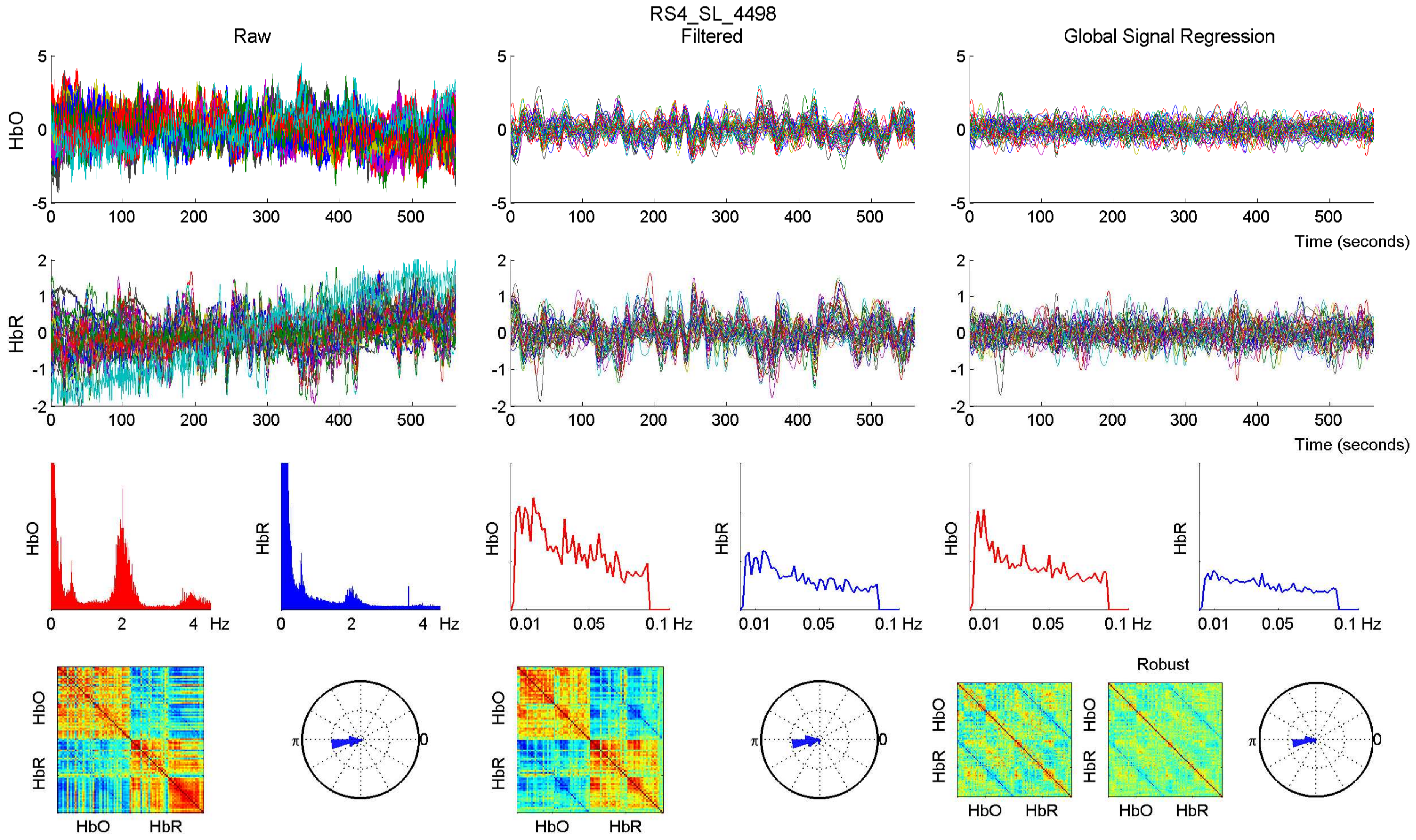

RS4_SL_4498
Raw
Filtered
Global Signal Regression
HbO
HbR
Time (seconds)
Hz
0.01
0.05
0.1 Hz
Robust
π
0

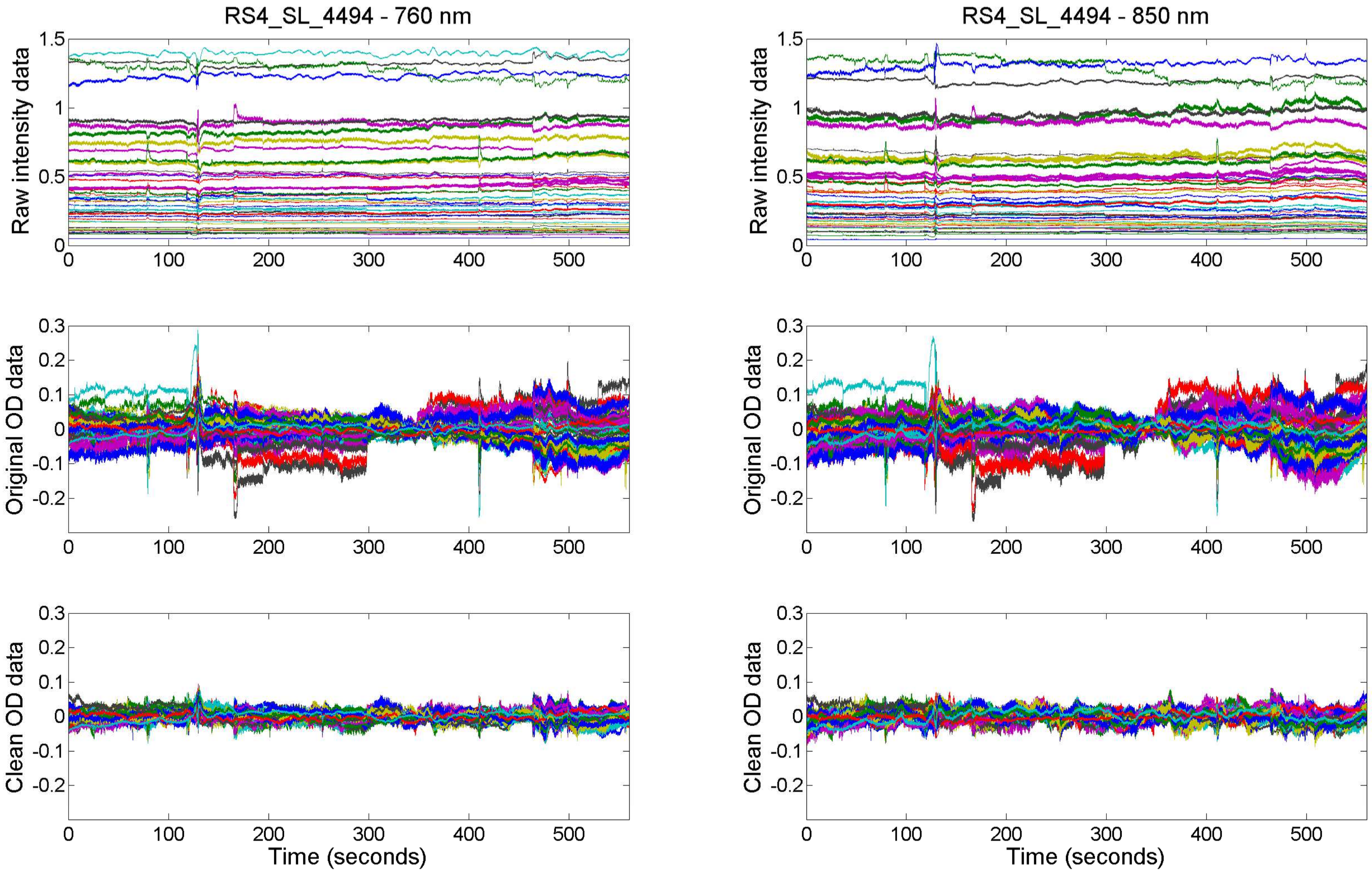

RS4_SL_4494 - 760 nm
RS4_SL_4494 - 850 nm
Raw intensity data
Original OD data
Clean OD data
Time (seconds)

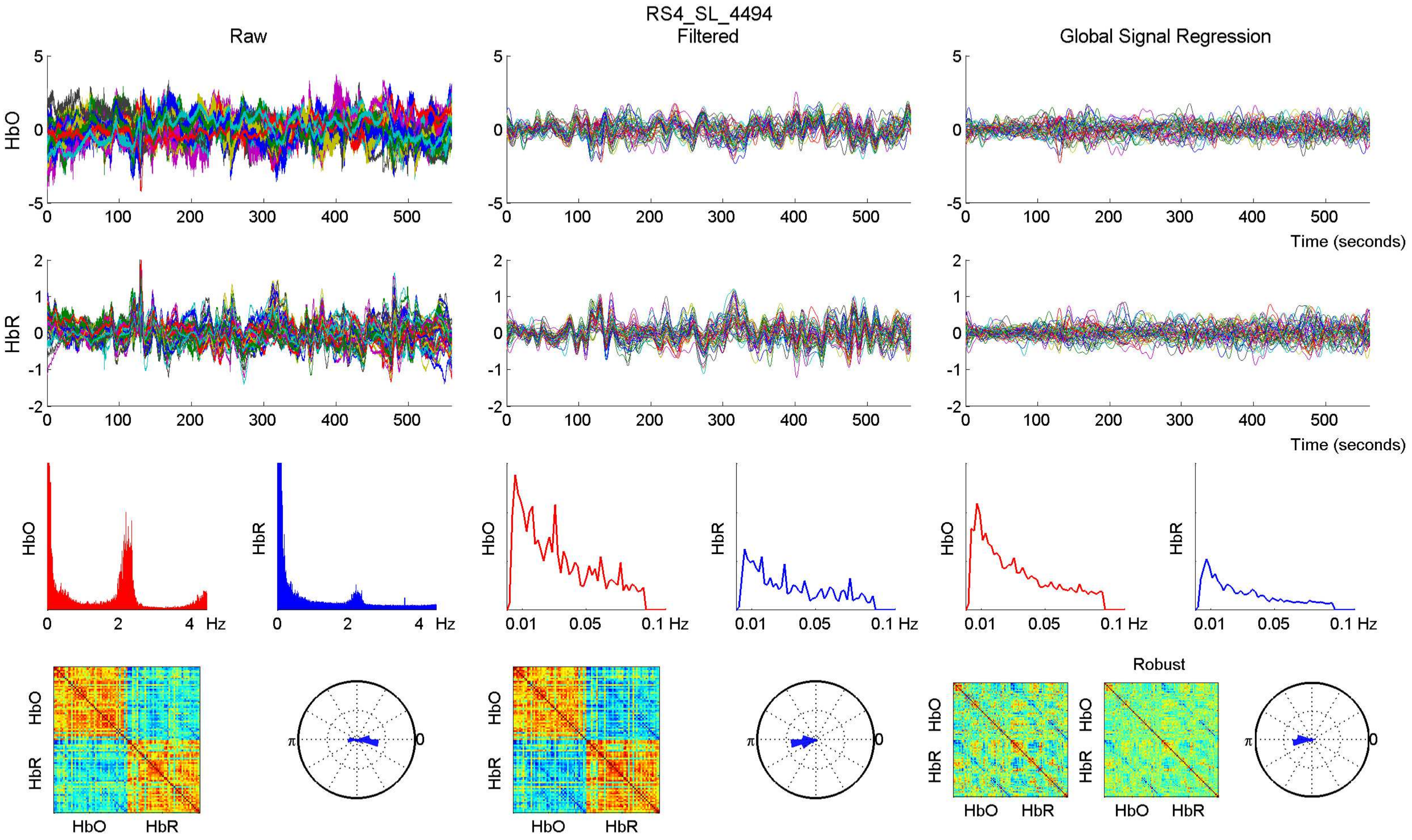

RS4_SL_4494
Raw
Filtered
Global Signal Regression
HbO
HbR
Time (seconds)
Hz
Robust

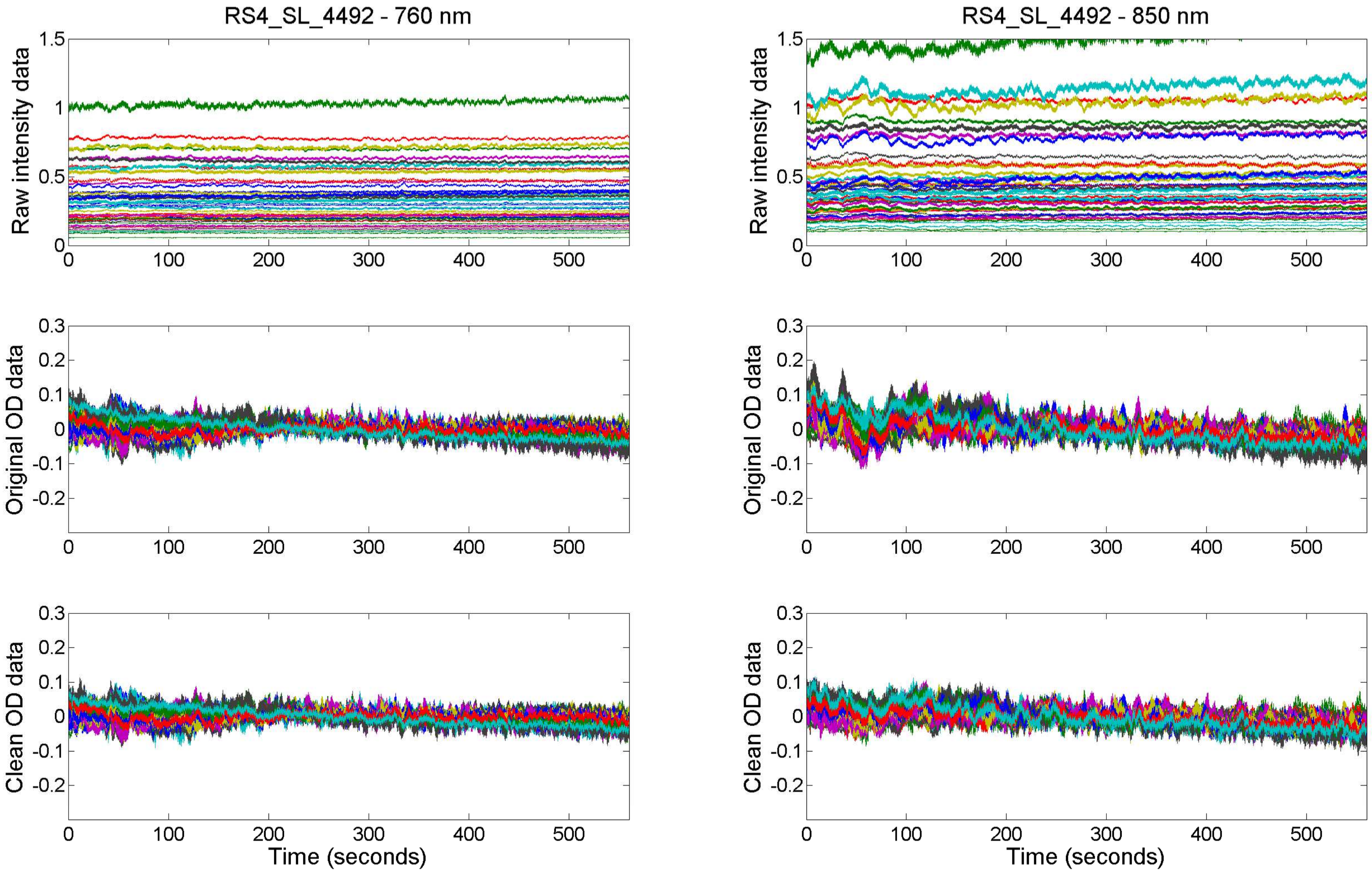

RS4_SL_4492 - 760 nm
RS4_SL_4492 - 850 nm
Raw intensity data
Original OD data
Clean OD data
Time (seconds)

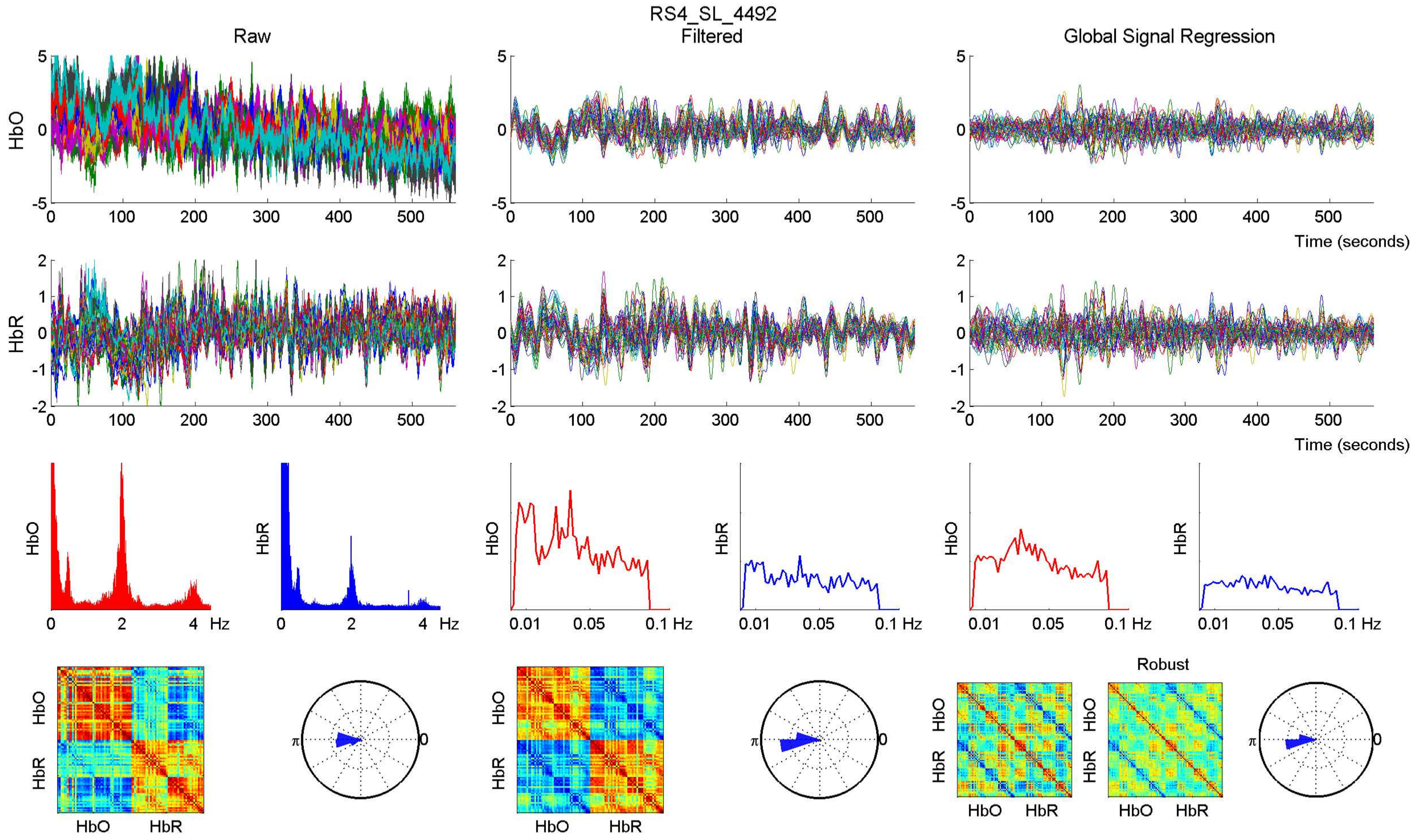
RS4_SL_4492
Raw
Filtered
Global Signal Regression
HbO
HbR
Time (seconds)
Hz
Robust

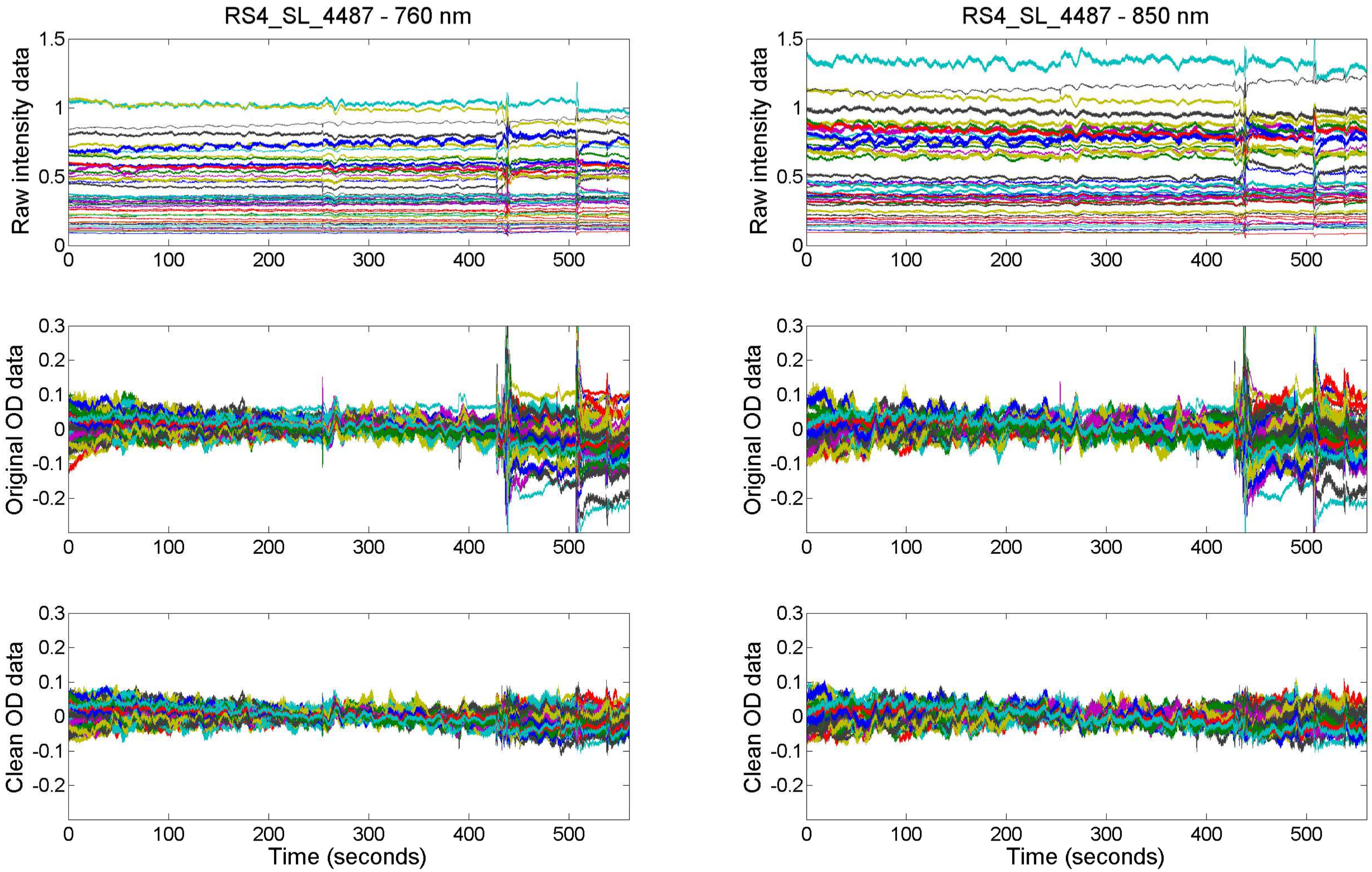

RS4_SL_4487 - 760 nm
RS4_SL_4487 - 850 nm
Raw intensity data
Original OD data
Clean OD data
Time (seconds)
0
100
200
300
400
500
1.5
1
0.5
0.3
0.2
0.1
-0.1
-0.2

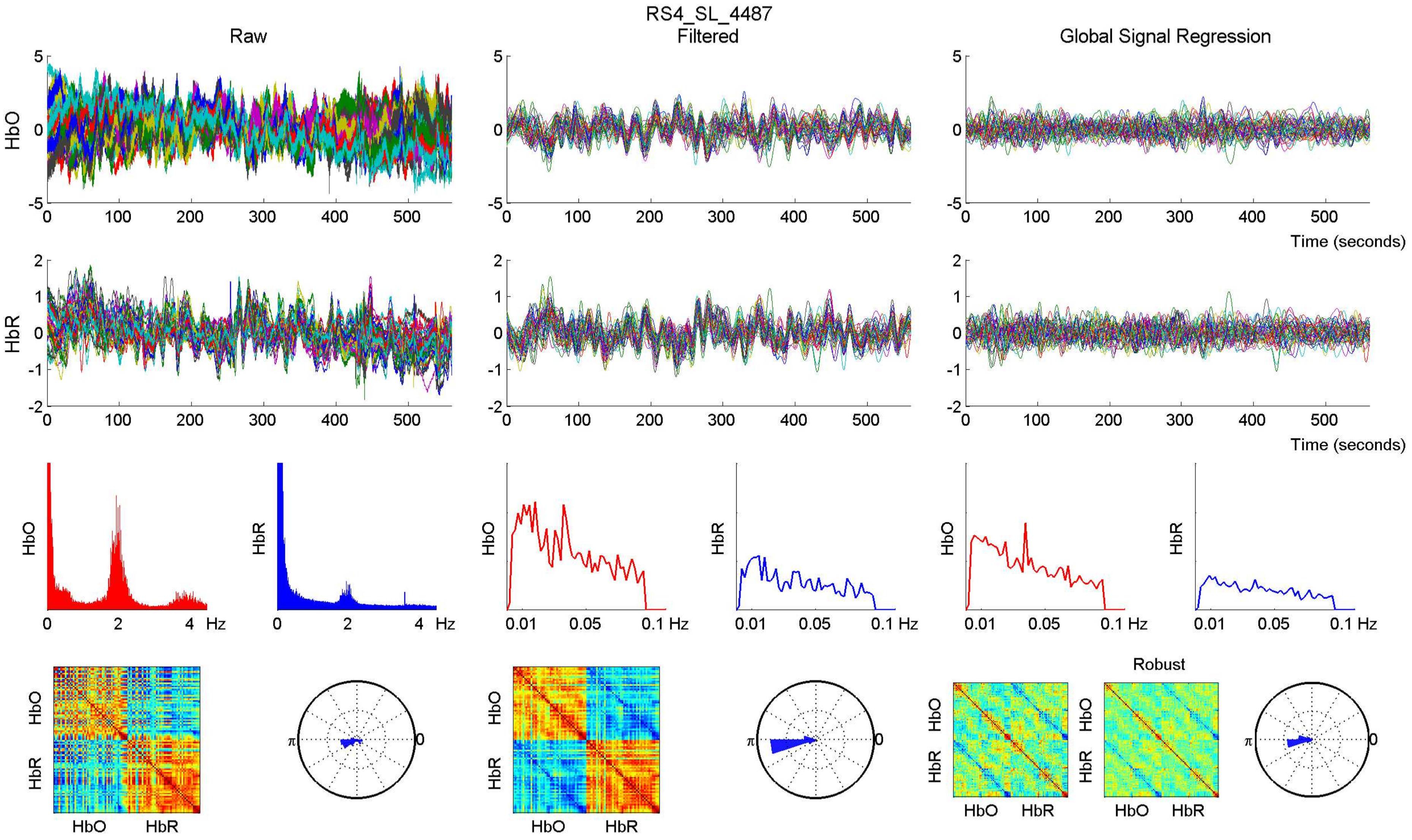

RS4_SL_4487
Raw
Filtered
Global Signal Regression
HbO
HbR
Time (seconds)
Hz
0.01
0.05
0.1 Hz
Robust
π
0

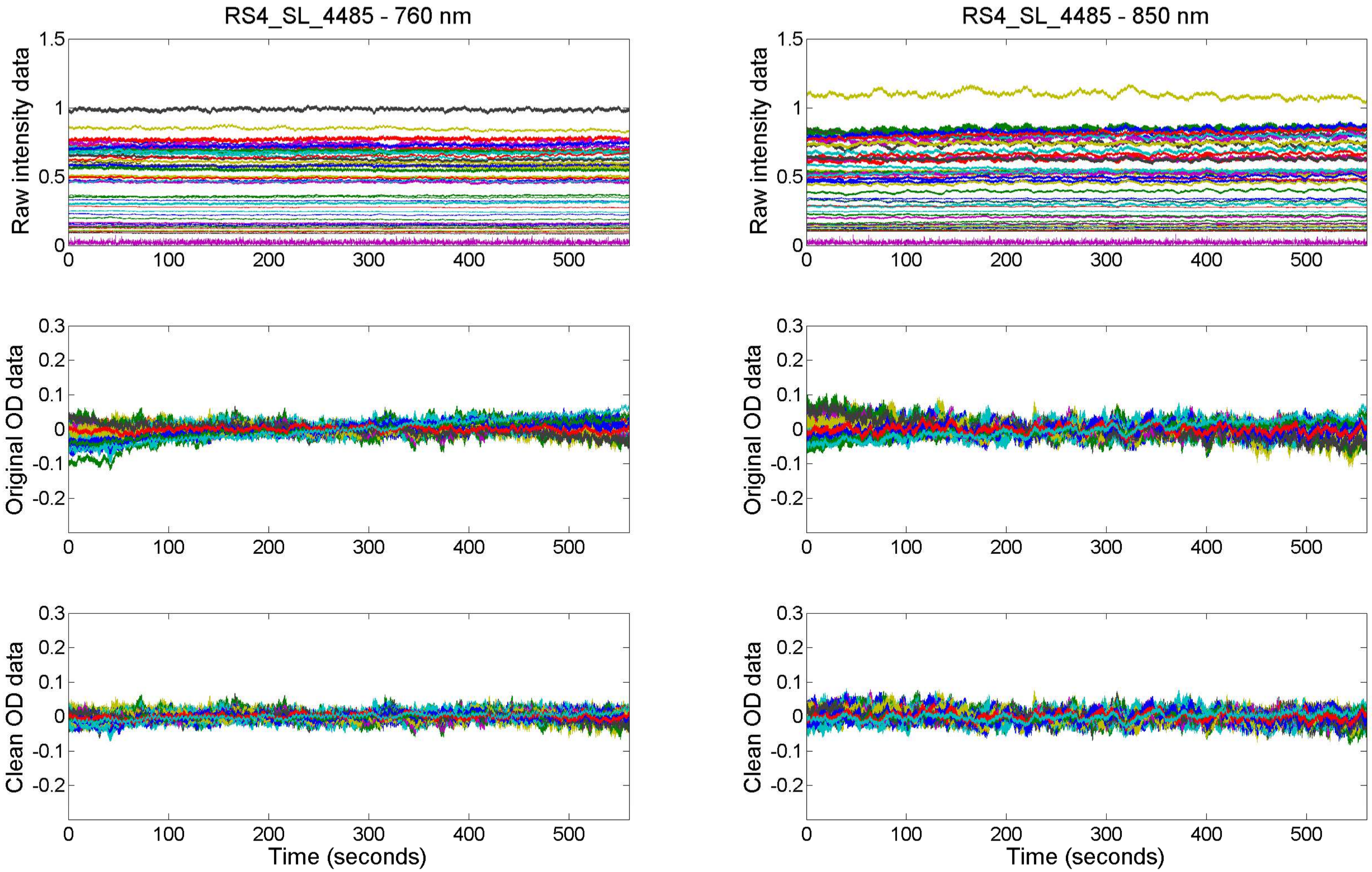

RS4_SL_4485 - 760 nm
RS4_SL_4485 - 850 nm
Raw intensity data
Original OD data
Clean OD data
Time (seconds)

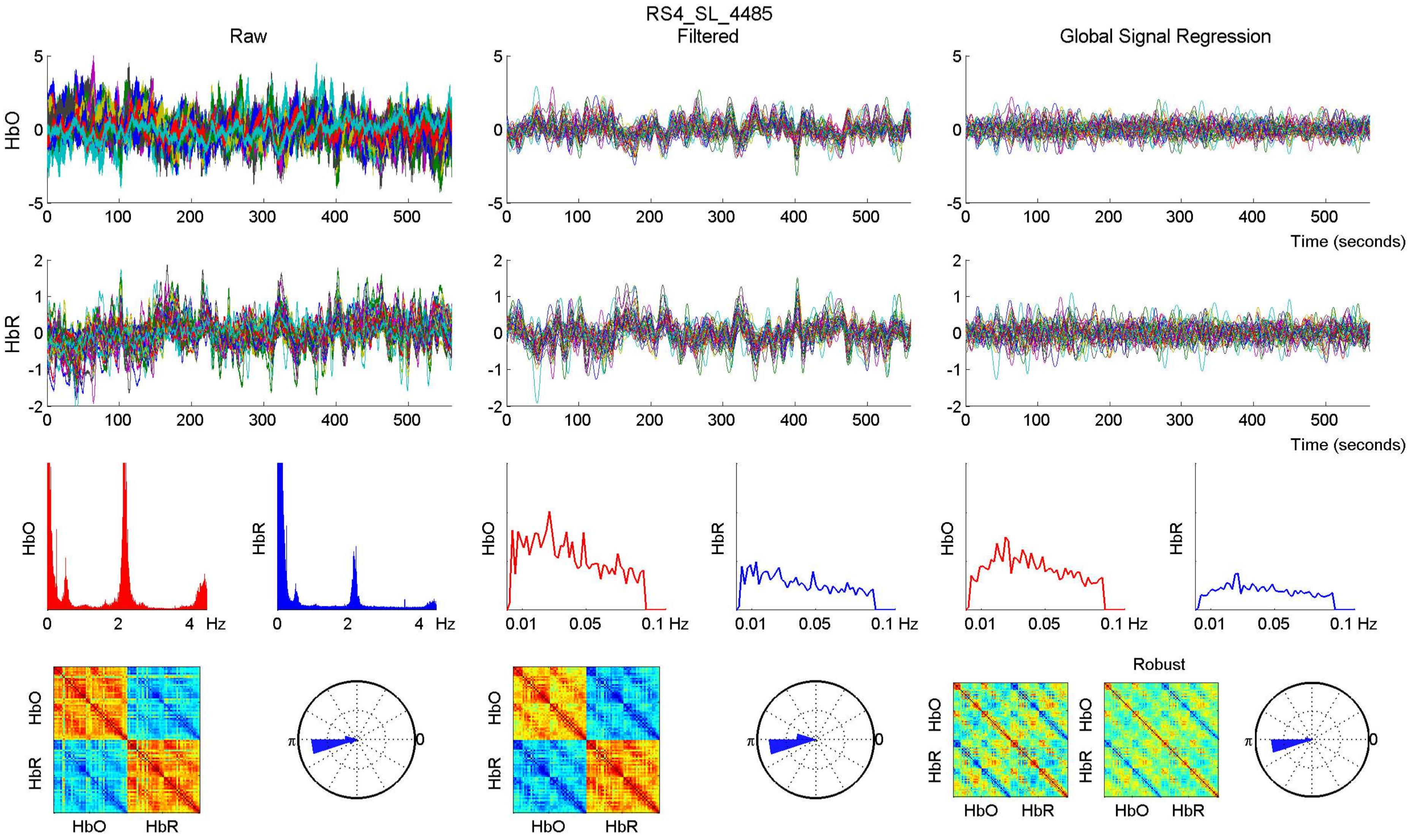

RS4_SL_4485
Raw
Filtered
Global Signal Regression
HbO
HbR
Time (seconds)
Hz
Robust
HbO
HbR

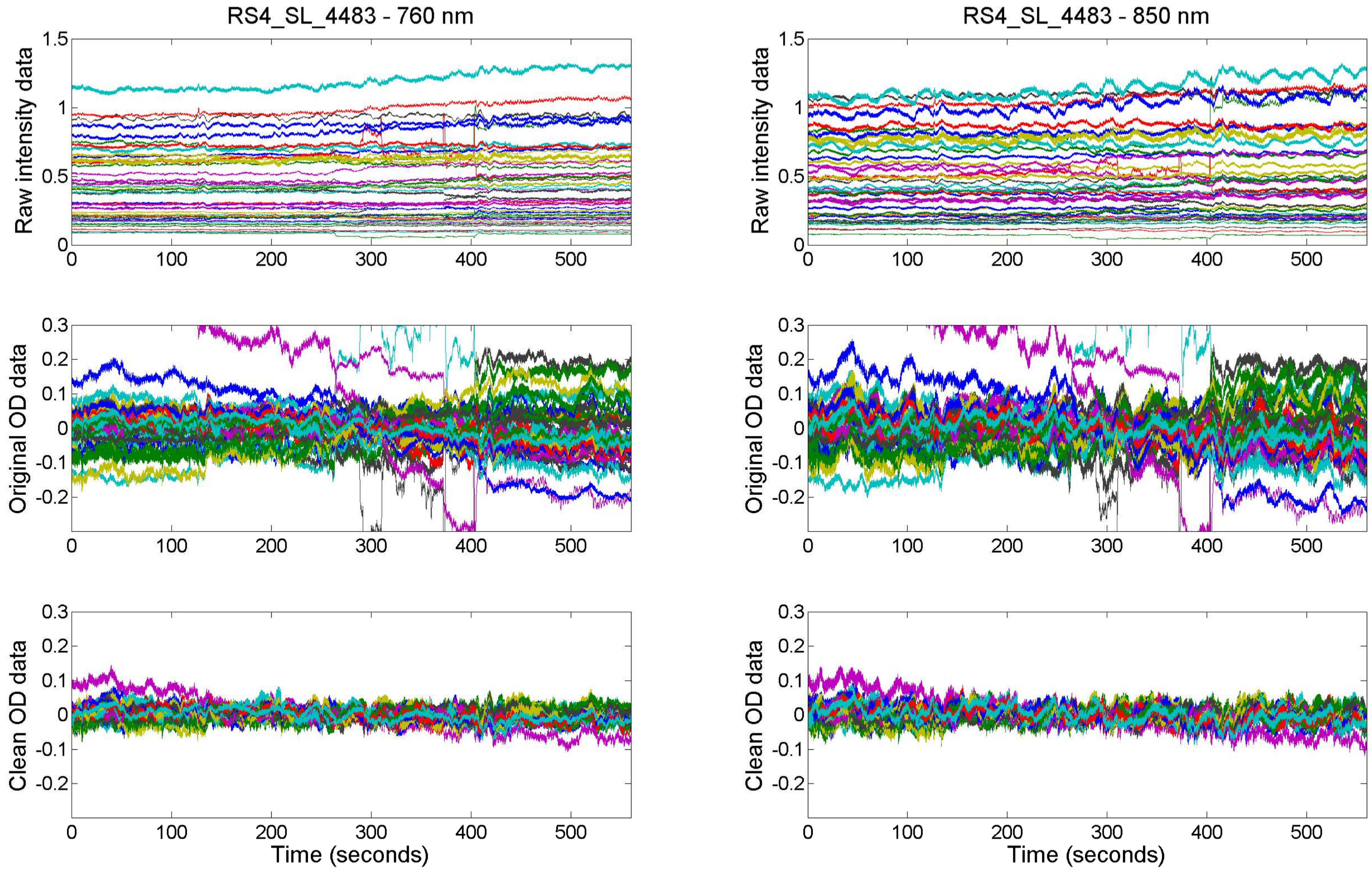

RS4_SL_4483 - 760 nm
RS4_SL_4483 - 850 nm
Raw intensity data
Original OD data
Clean OD data
Time (seconds)

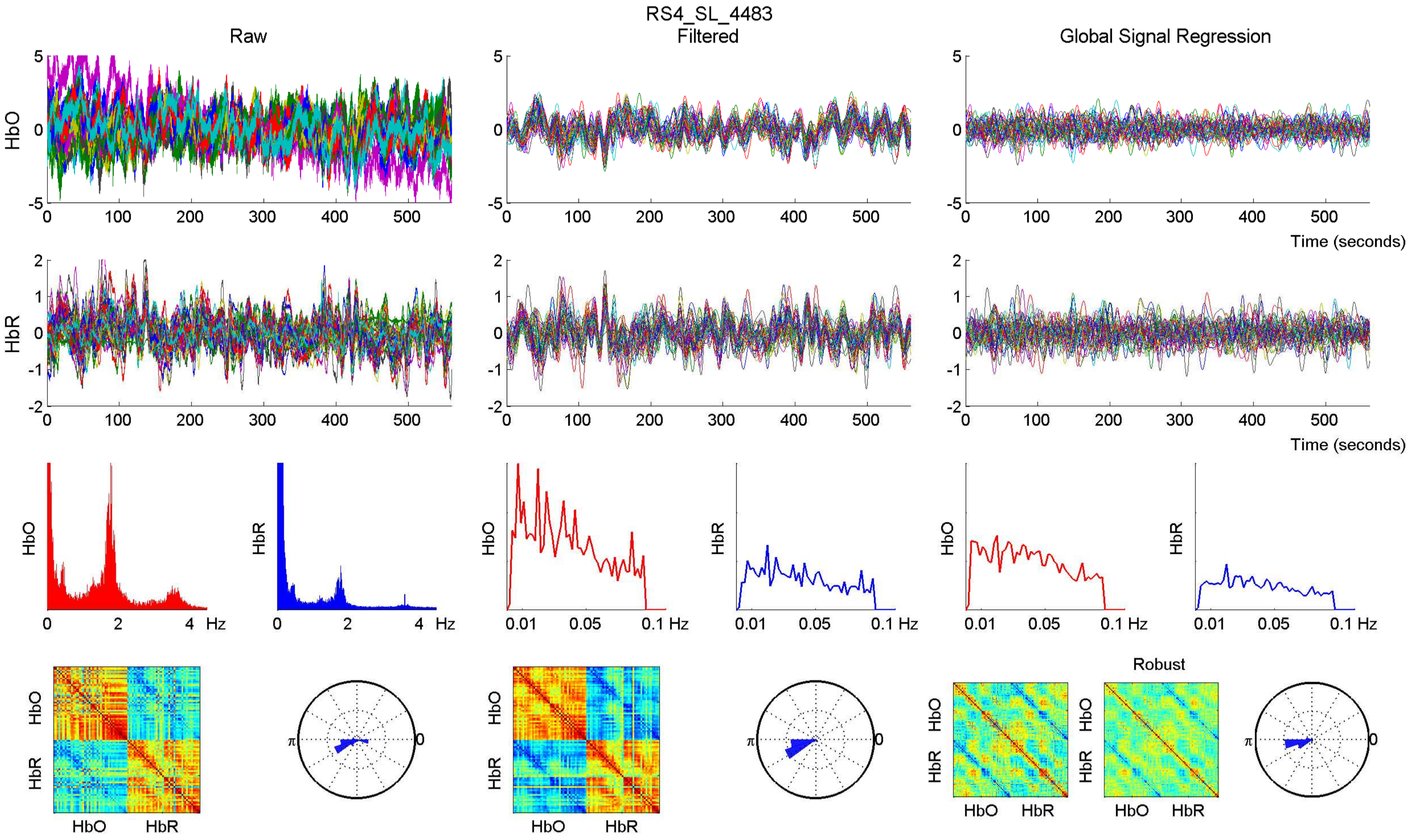

RS4_SL_4483
Raw
Filtered
Global Signal Regression
HbO
HbR
Time (seconds)
Hz
0.01
0.05
0.1 Hz
Robust
π
0

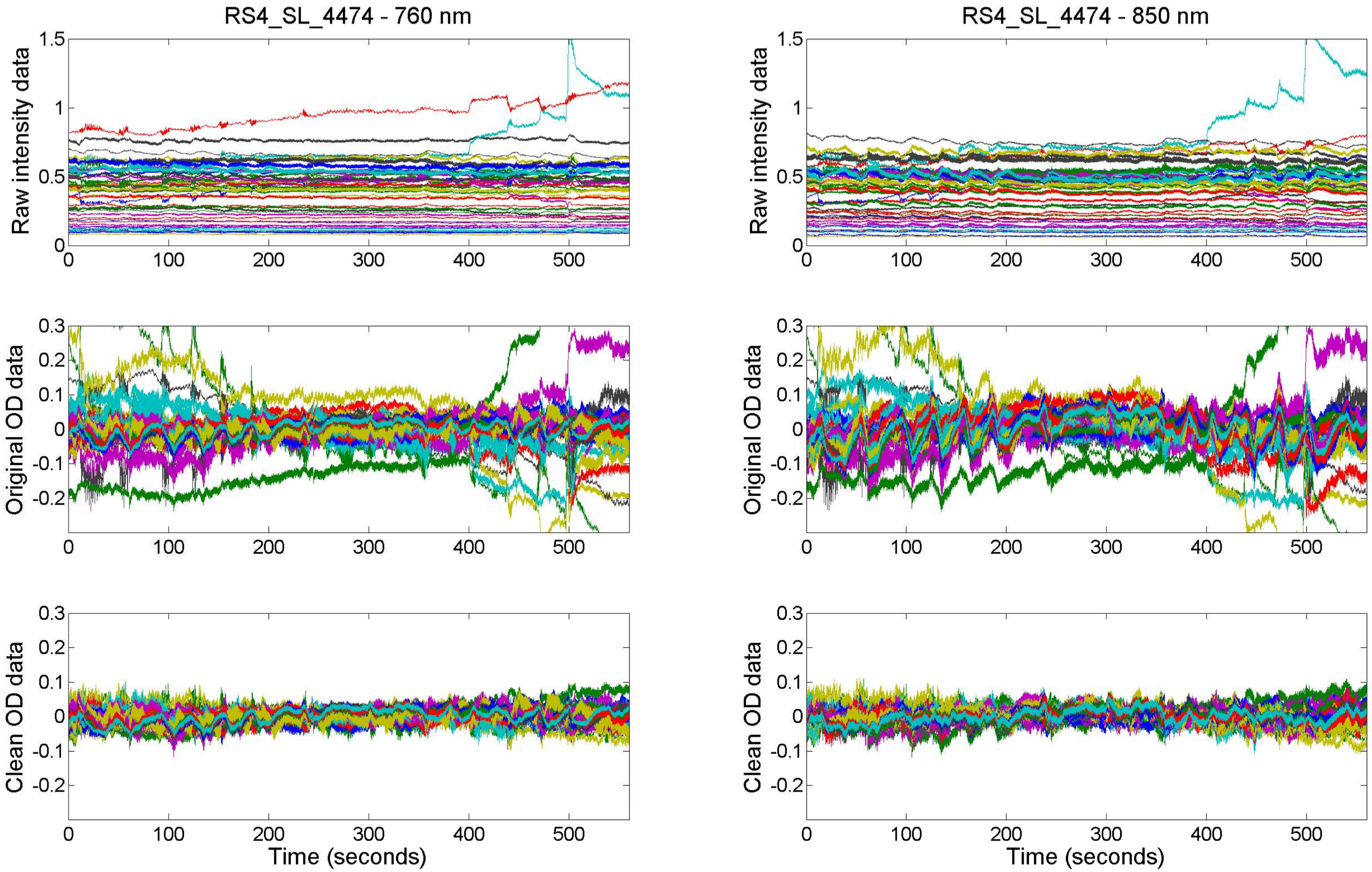

RS4_SL_4474 - 760 nm
RS4_SL_4474 - 850 nm
Raw intensity data
Original OD data
Clean OD data
Time (seconds)

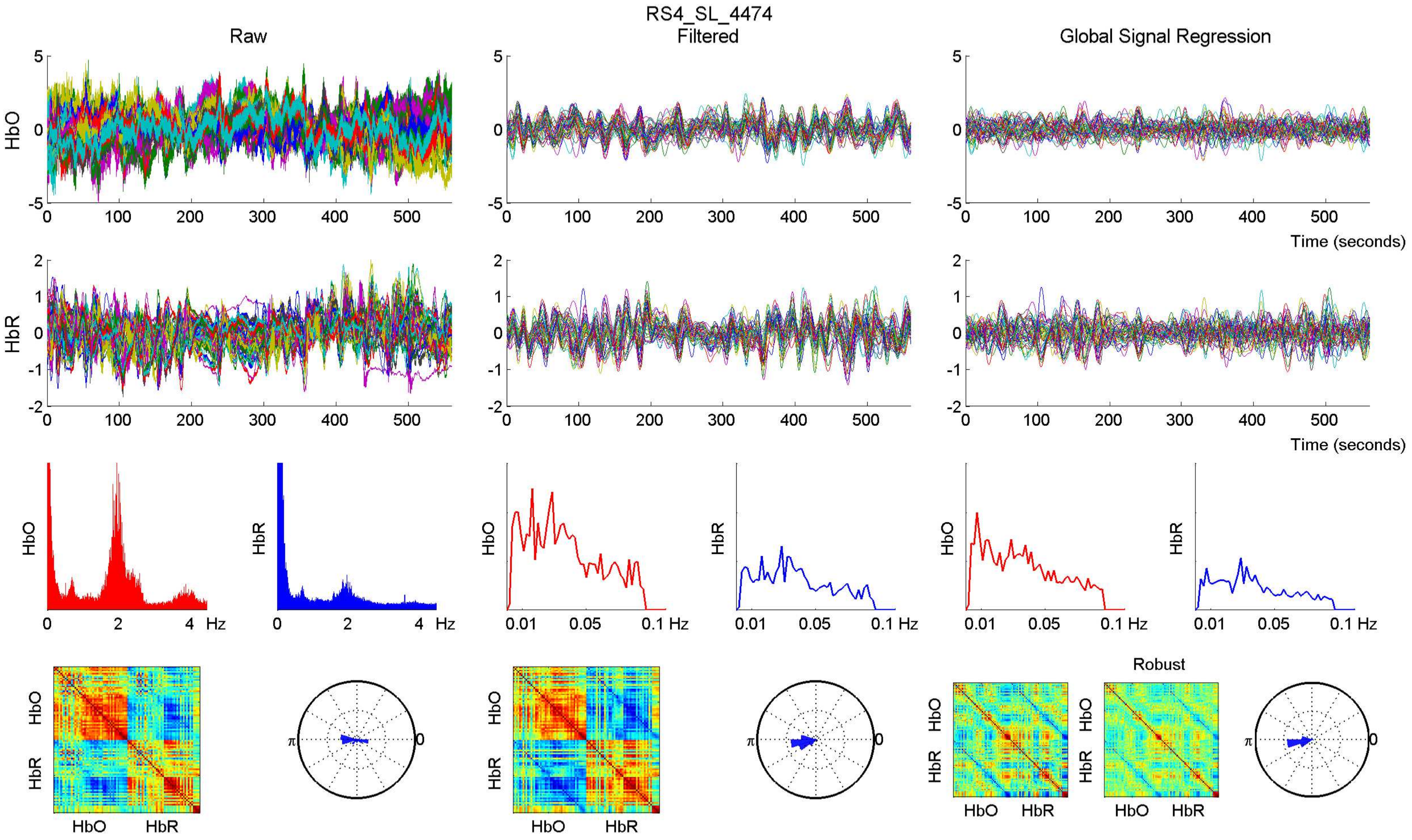
RS4_SL_4474
Raw
Filtered
Global Signal Regression
HbO
HbR
Time (seconds)
Hz
Robust

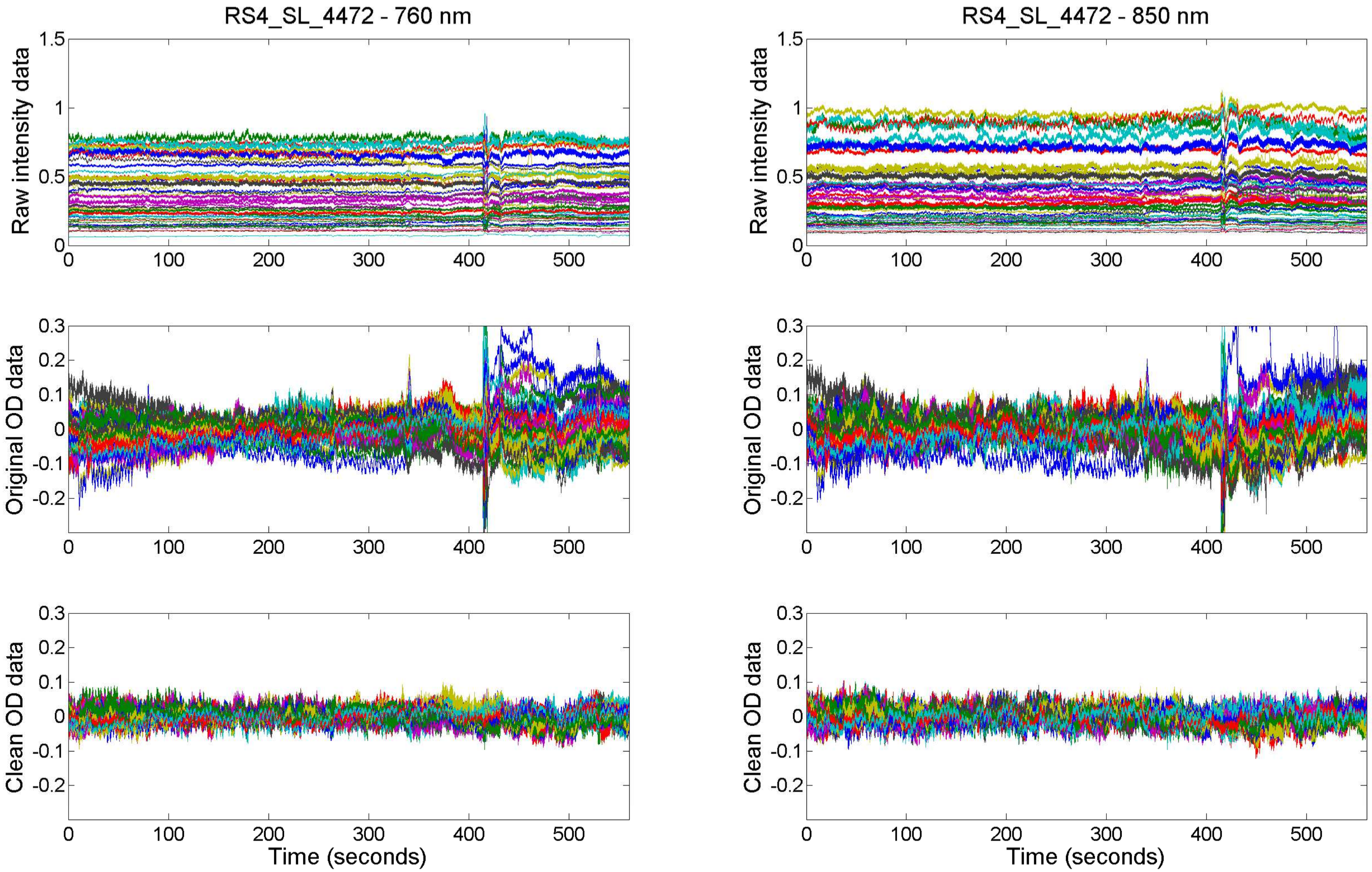

RS4_SL_4472 - 760 nm
RS4_SL_4472 - 850 nm
Raw intensity data
Original OD data
Clean OD data
Time (seconds)

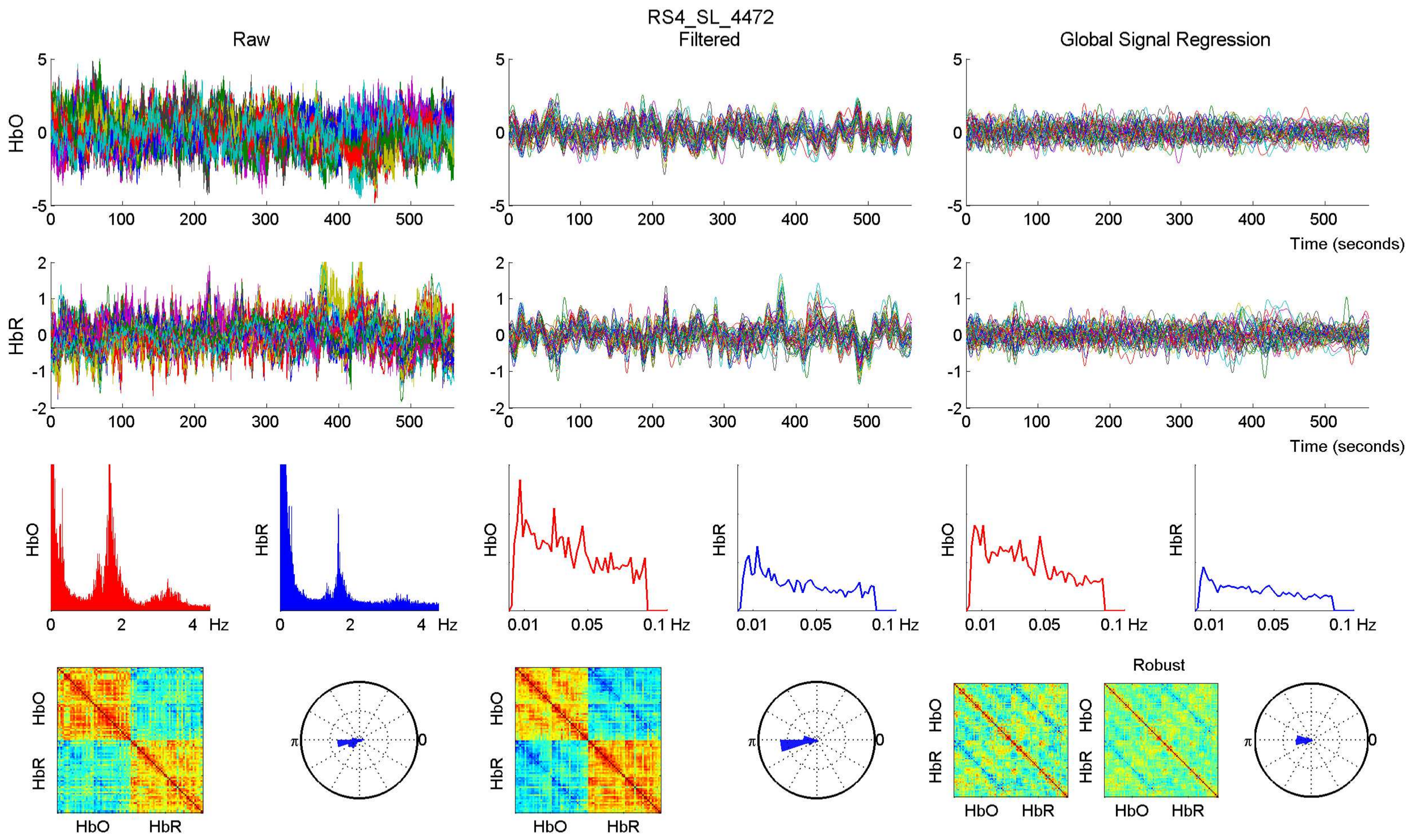
RS4_SL_4472
Raw
Filtered
Global Signal Regression
HbO
HbR
Time (seconds)
Hz
Robust

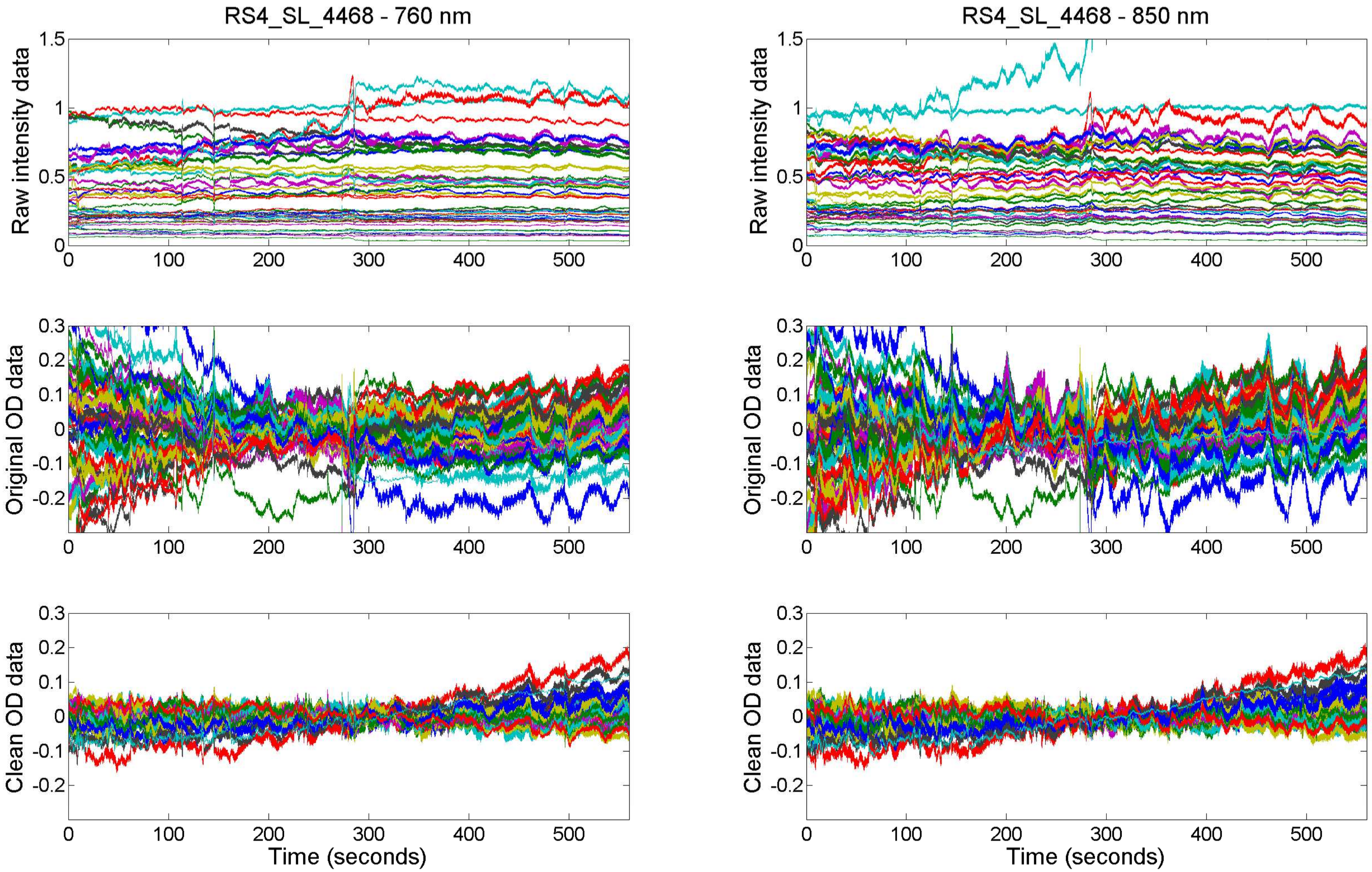

RS4_SL_4468 - 760 nm
RS4_SL_4468 - 850 nm
Raw intensity data
Original OD data
Clean OD data
Time (seconds)

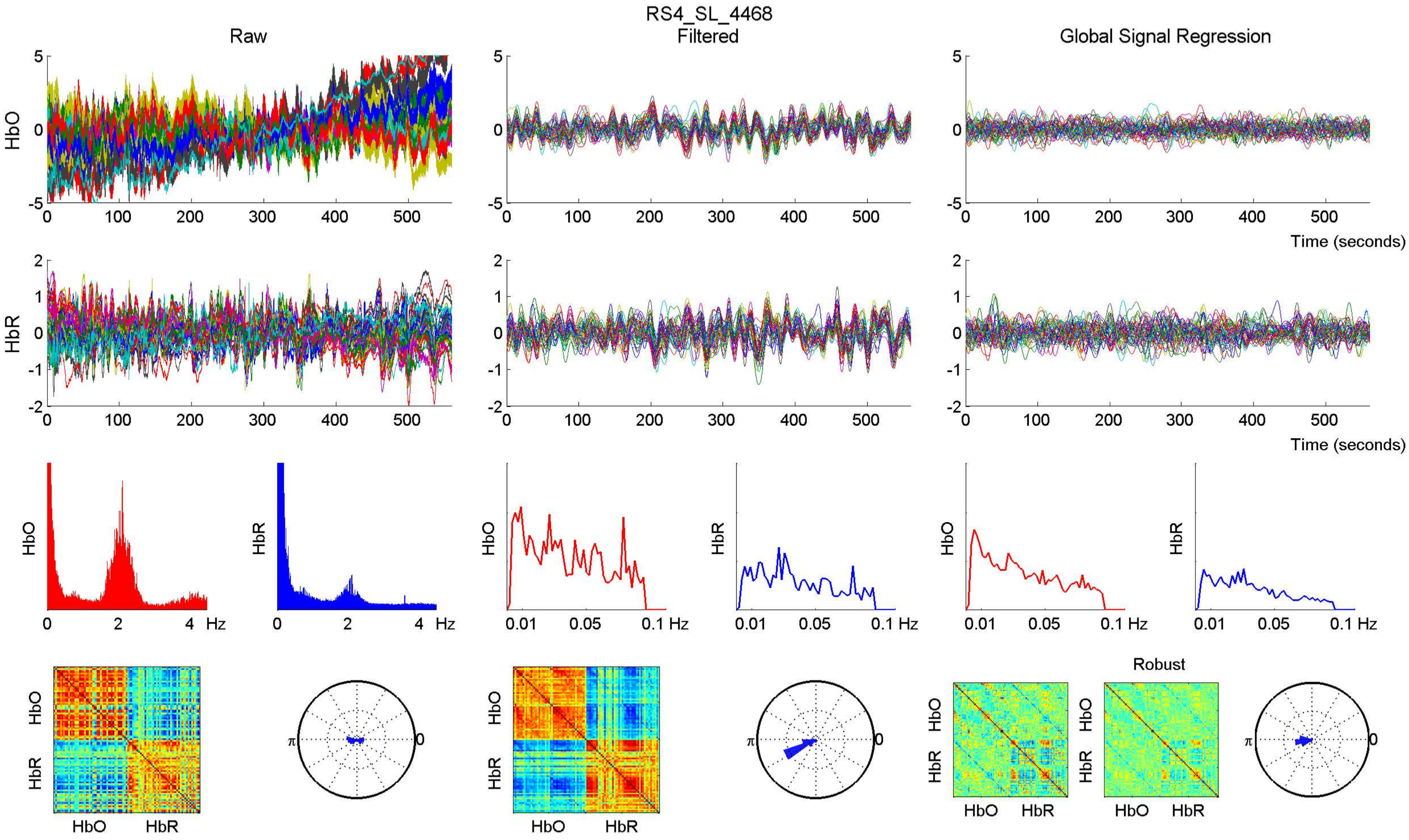

RS4_SL_4468
Raw
Filtered
Global Signal Regression
HbO
HbR
Time (seconds)
Hz
Robust

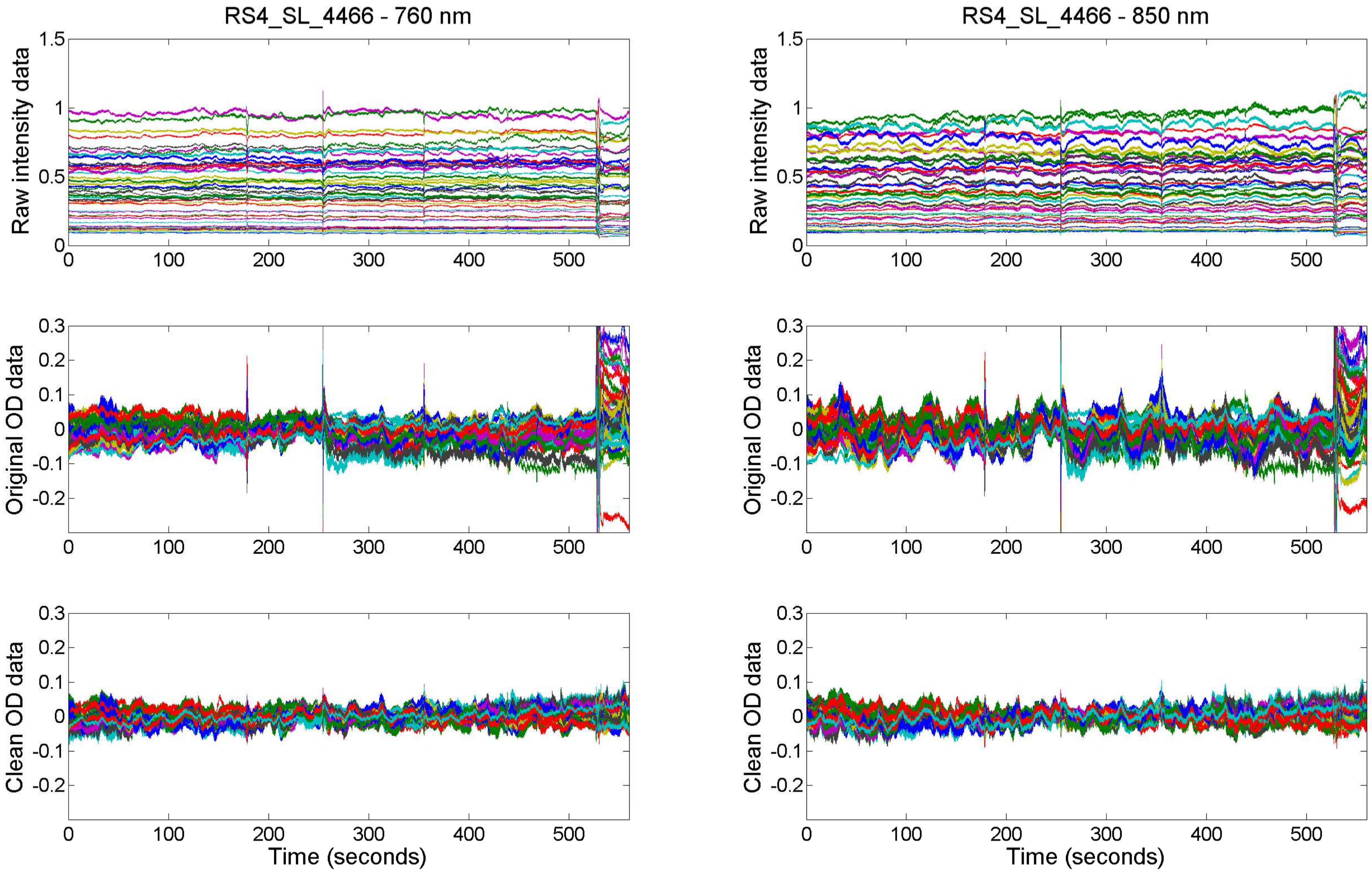
RS4_SL_4466 - 760 nm
RS4_SL_4466 - 850 nm
Raw intensity data
Original OD data
Clean OD data
Time (seconds)
0
100
200
300
400
500
1.5
1
0.5
0.3
0.2
0.1
-0.1
-0.2

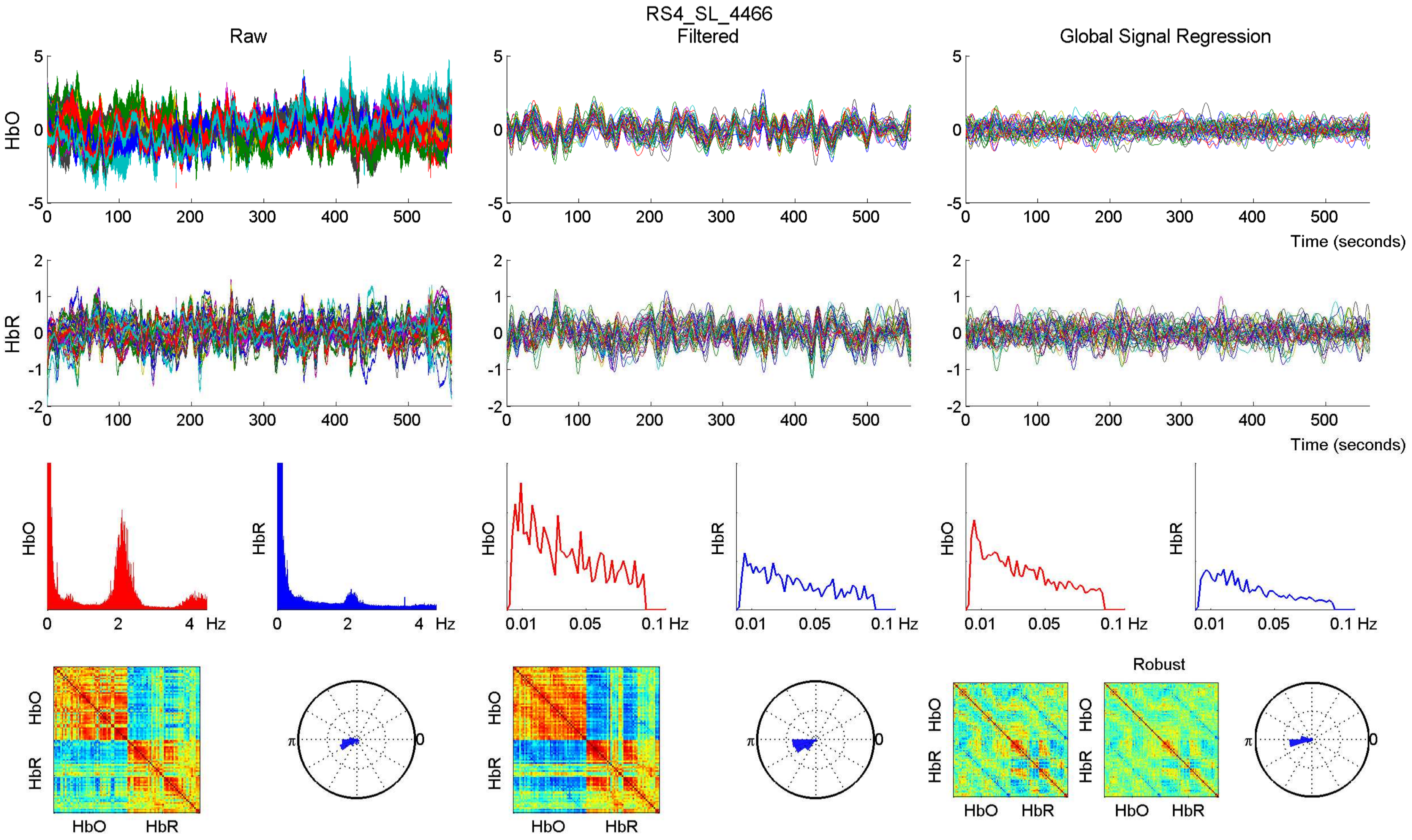

RS4_SL_4466
Raw
Filtered
Global Signal Regression
HbO
HbR
Time (seconds)
Hz
Robust

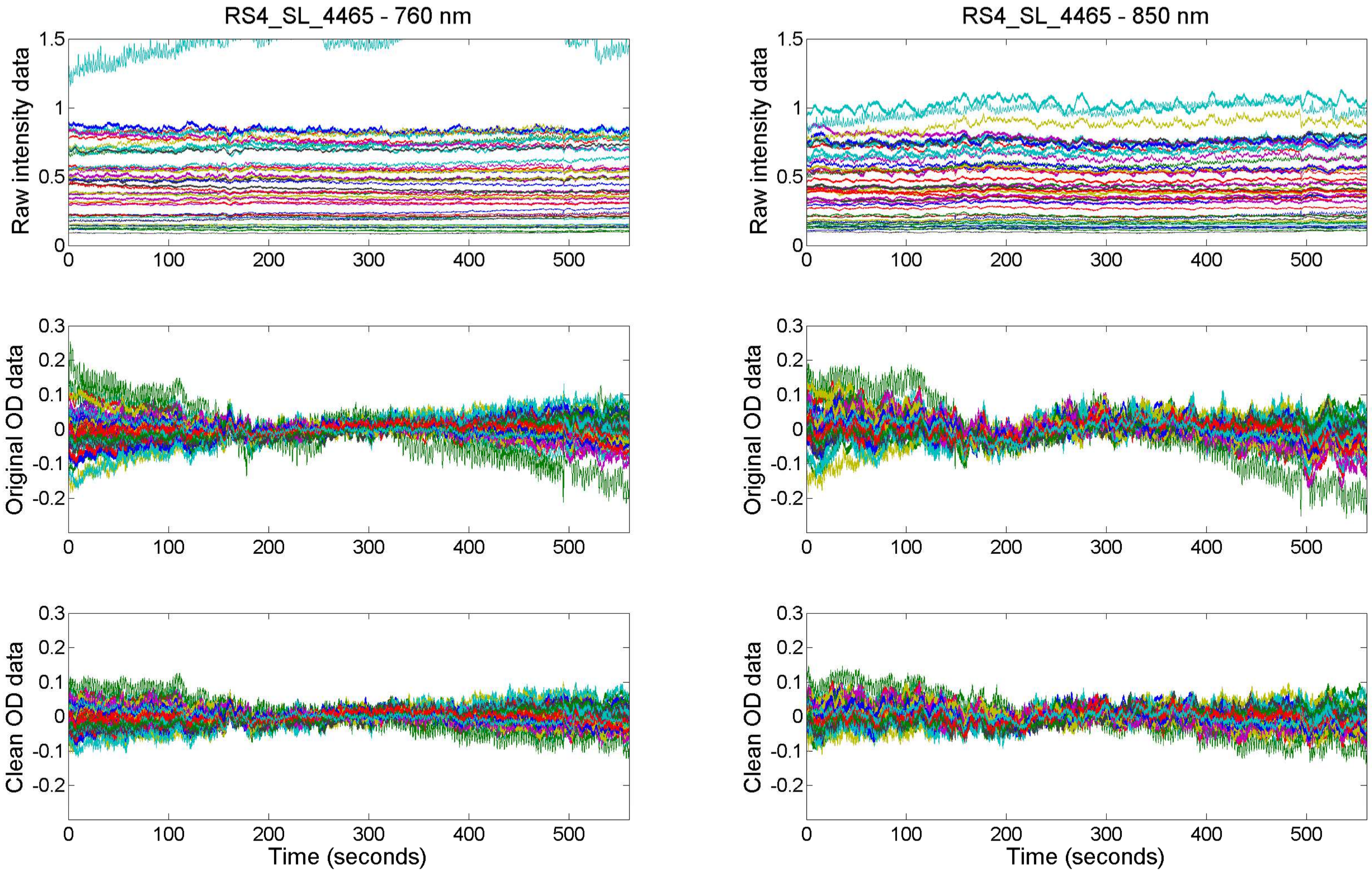

RS4_SL_4465 - 760 nm
RS4_SL_4465 - 850 nm
Raw intensity data
Original OD data
Clean OD data
Time (seconds)

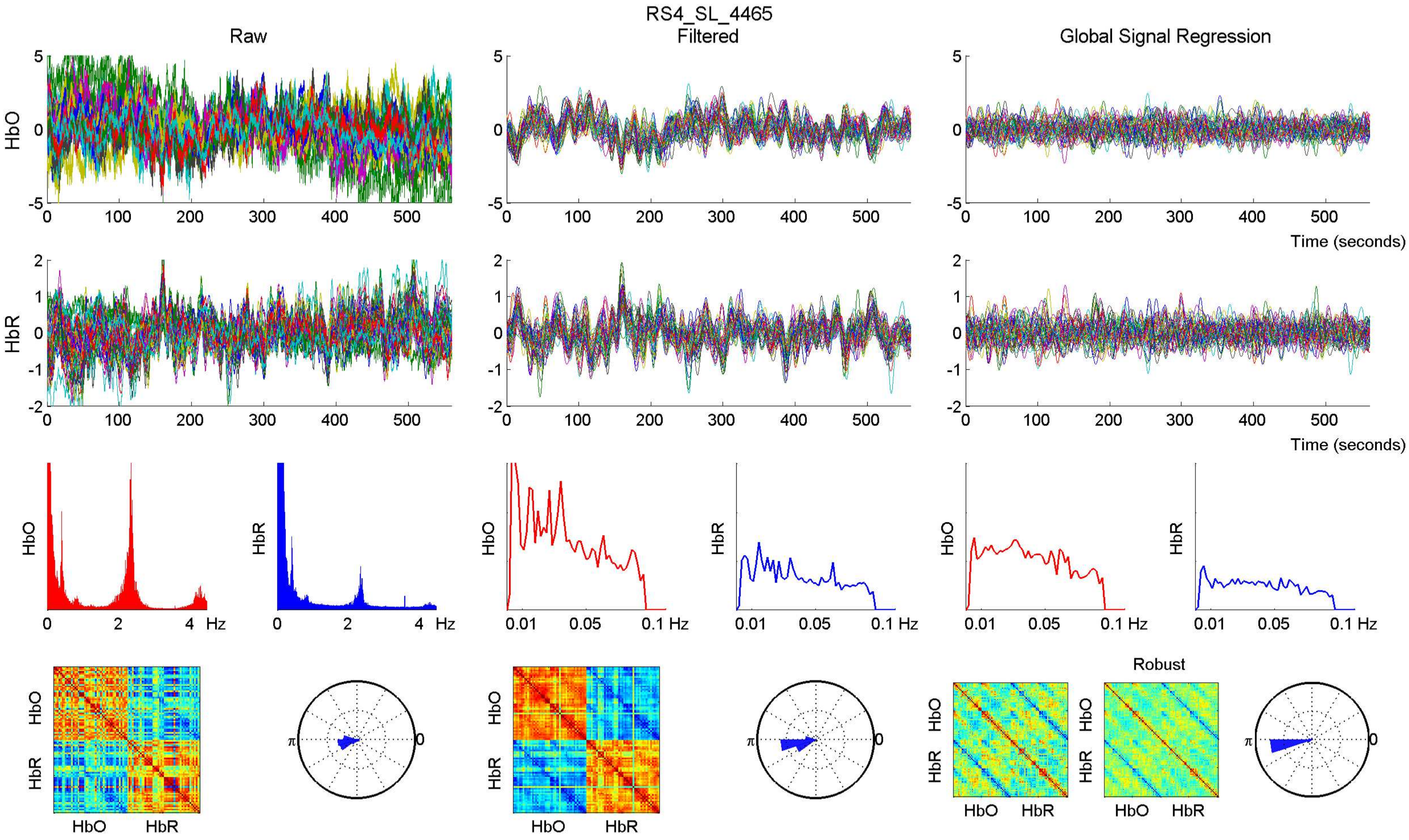

RS4_SL_4465
Raw
Filtered
Global Signal Regression
HbO
HbR
Time (seconds)
Hz
Robust

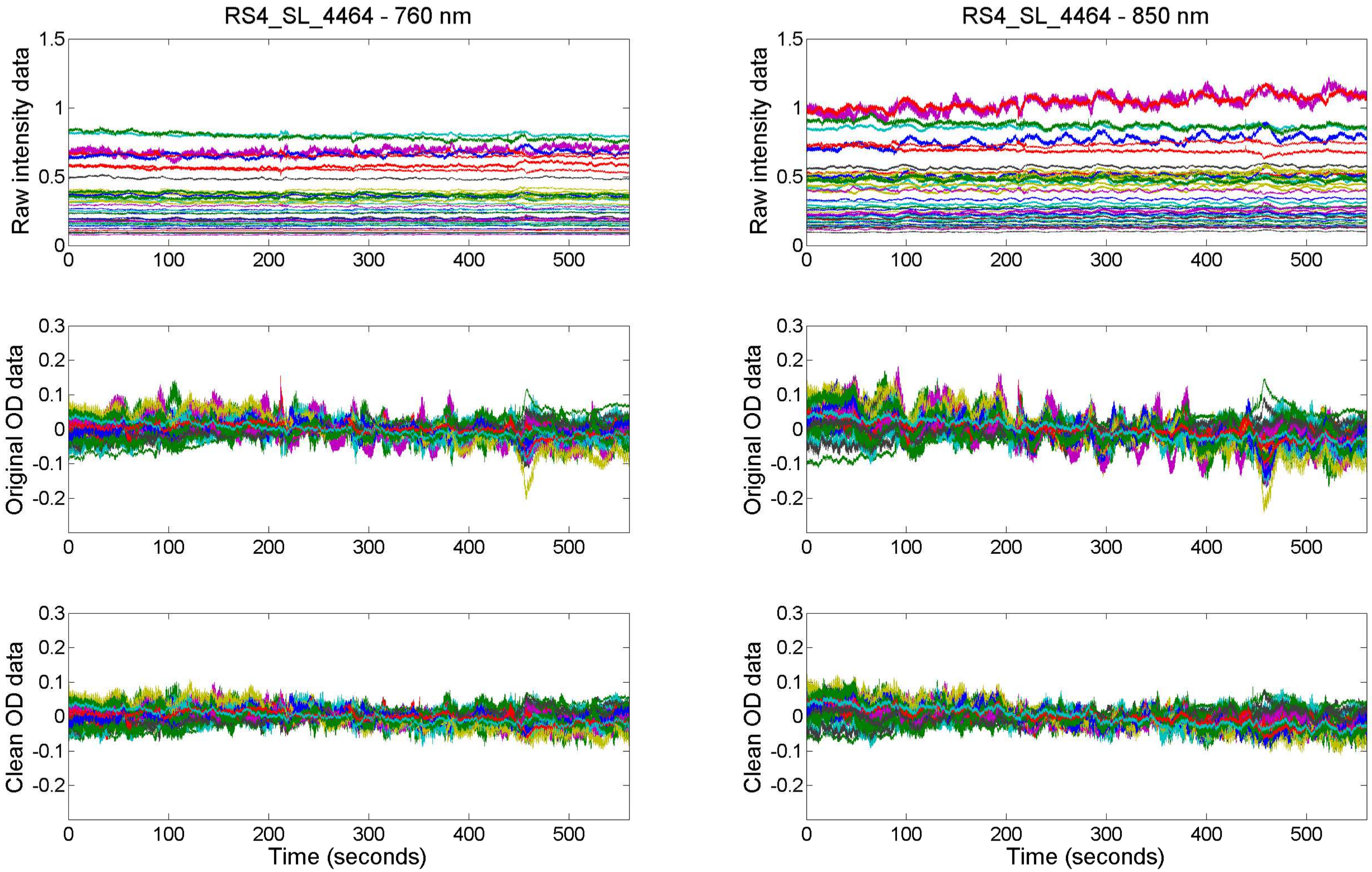

RS4_SL_4464 - 760 nm
RS4_SL_4464 - 850 nm
Raw intensity data
Original OD data
Clean OD data
Time (seconds)

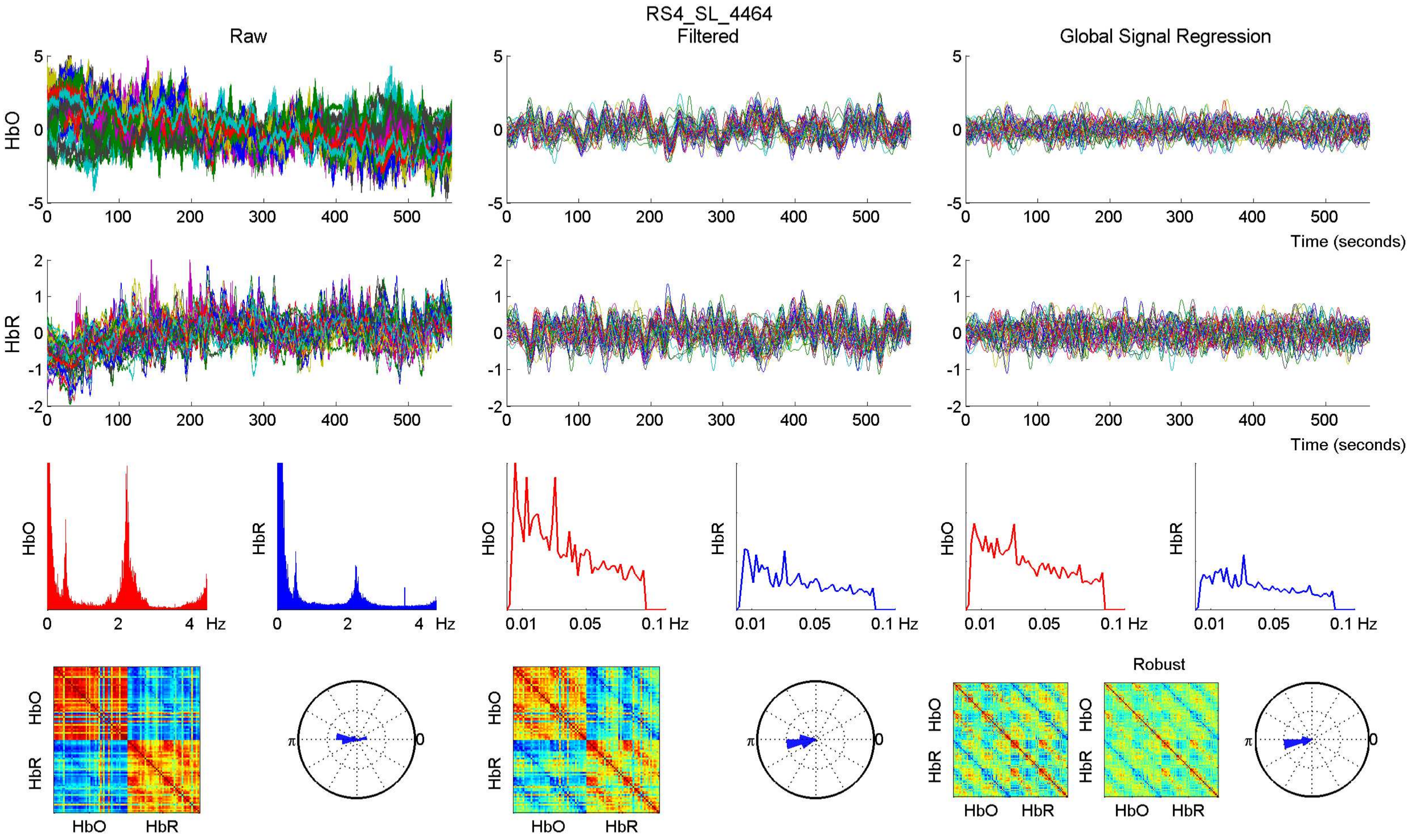

RS4_SL_4464
Raw
Filtered
Global Signal Regression
HbO
HbR
Time (seconds)
Hz
0.01
0.05
0.1 Hz
Robust
π
0

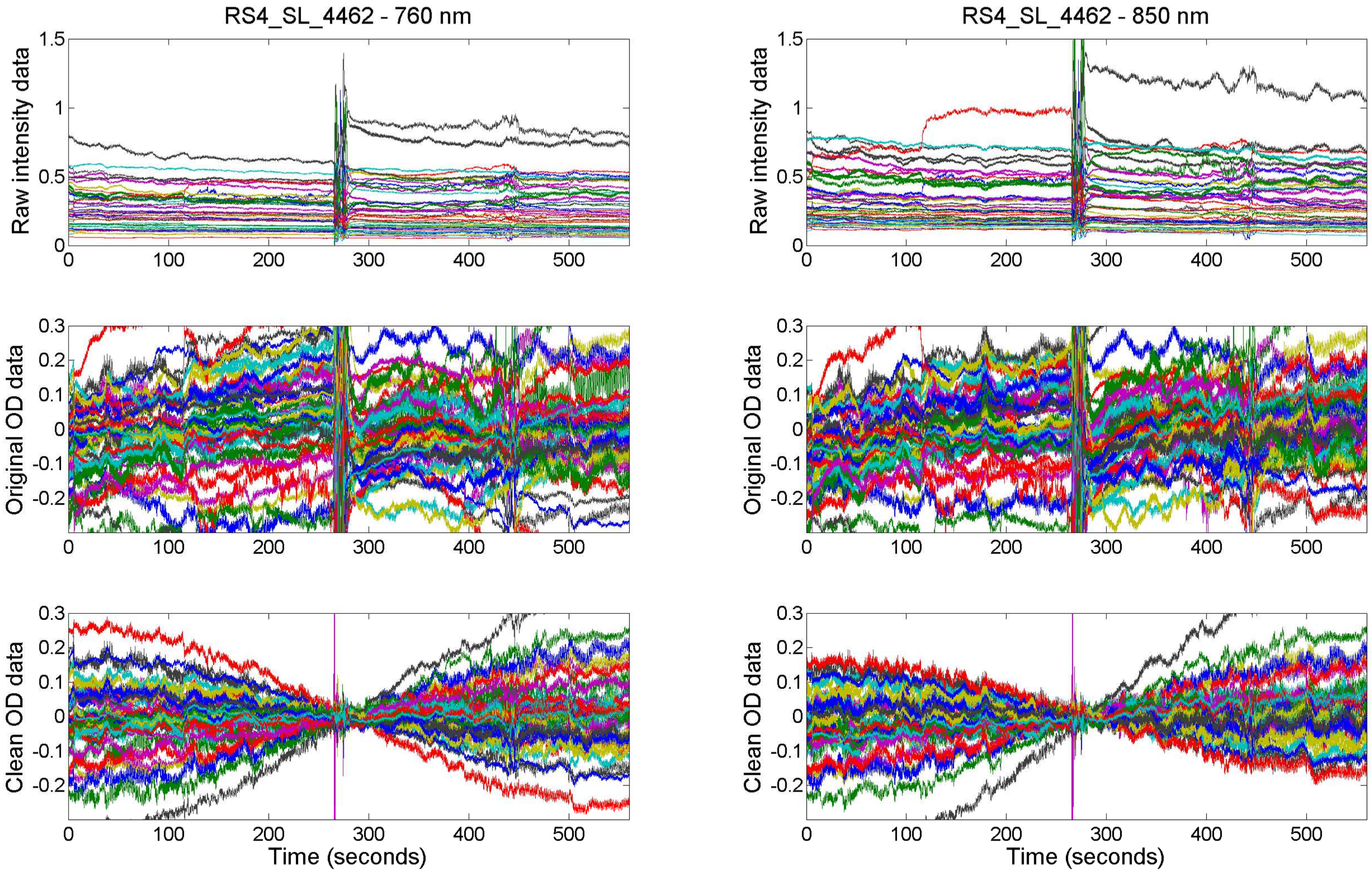
RS4_SL_4462 - 760 nm
RS4_SL_4462 - 850 nm
Raw intensity data
Original OD data
Clean OD data
Time (seconds)

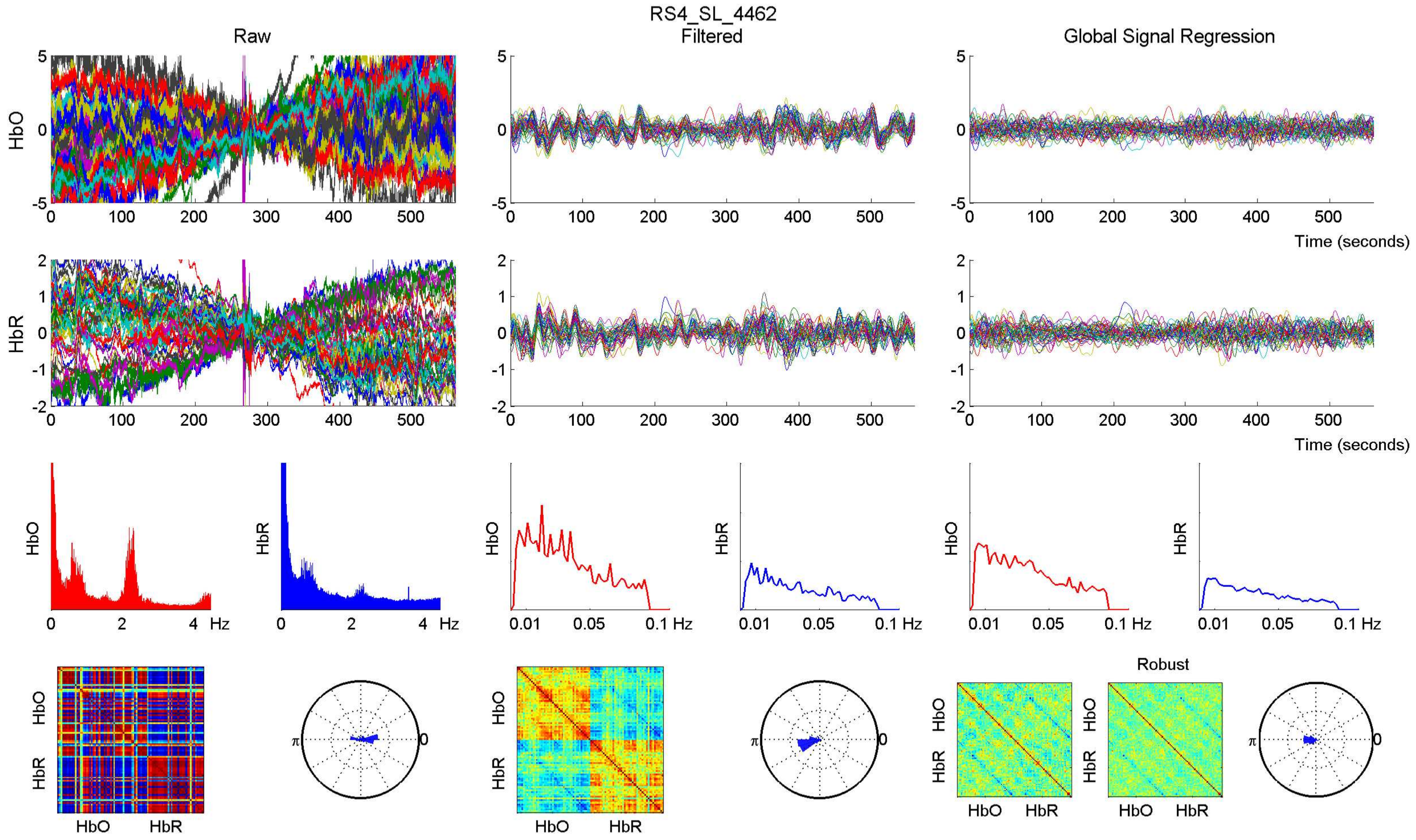

RS4_SL_4462
Raw
Filtered
Global Signal Regression
HbO
HbR
Time (seconds)
Hz
Robust

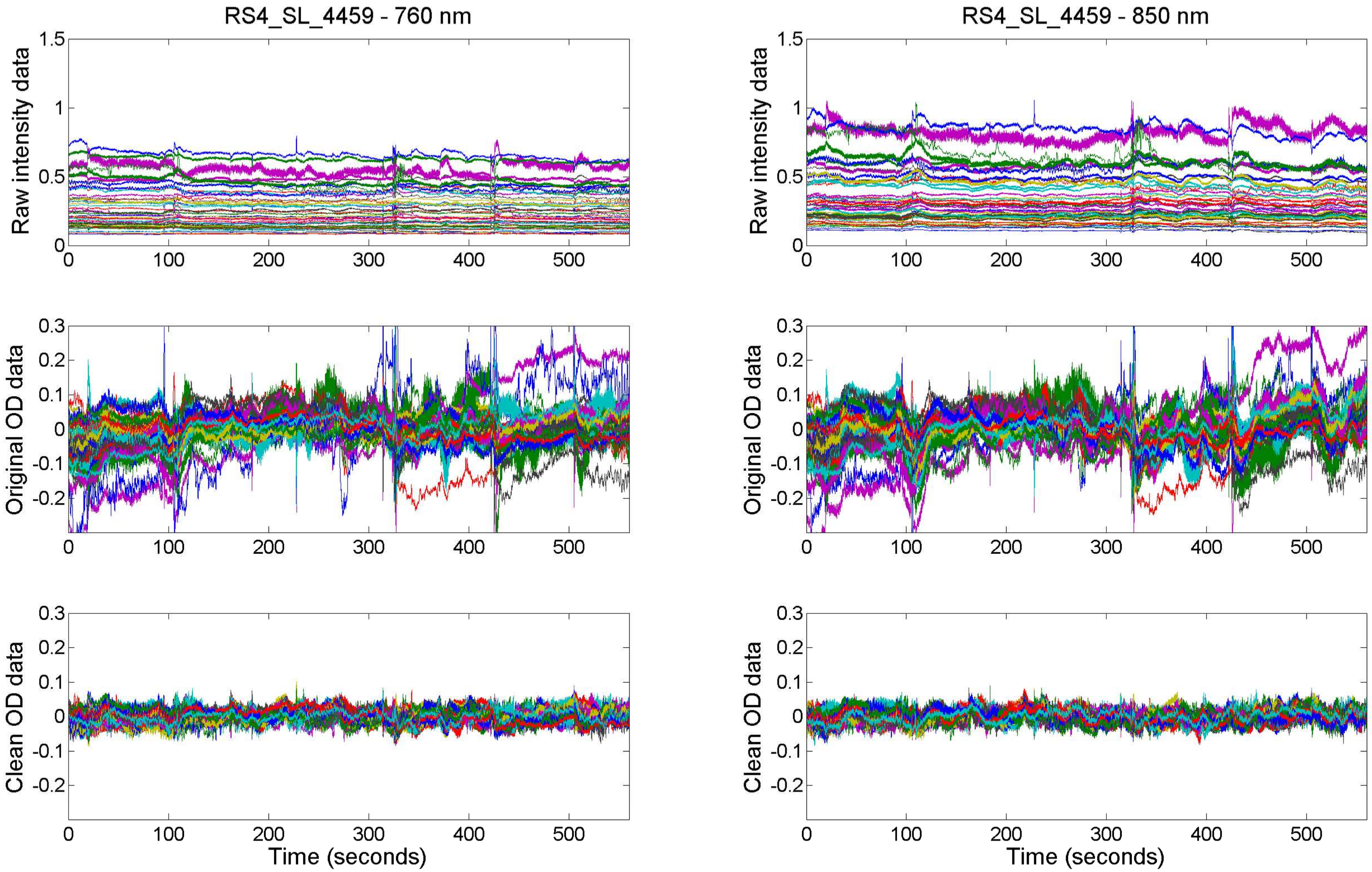

RS4_SL_4459 - 760 nm
RS4_SL_4459 - 850 nm
Raw intensity data
Original OD data
Clean OD data
Time (seconds)

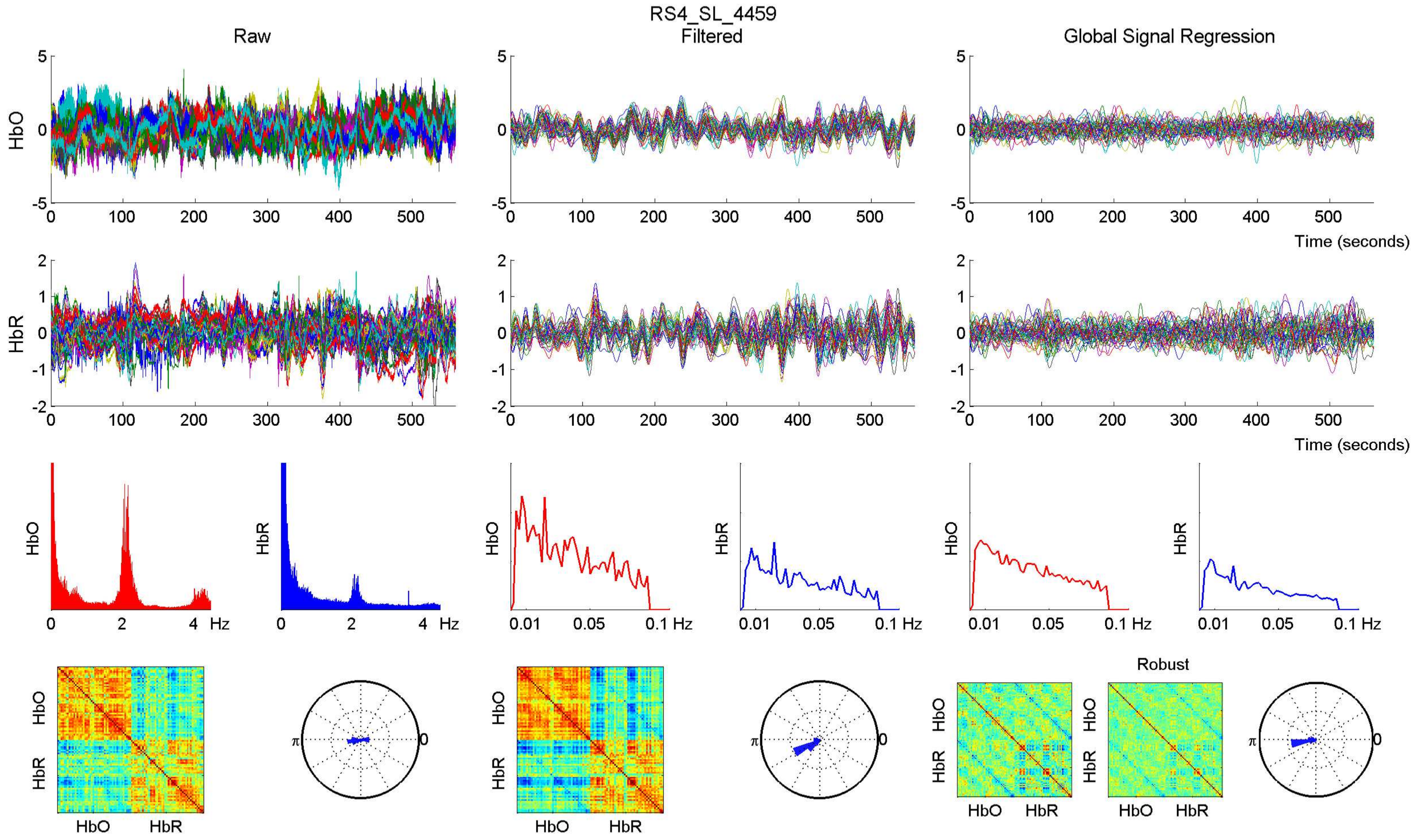
RS4_SL_4459
Raw
Filtered
Global Signal Regression
HbO
HbR
Time (seconds)
Hz
Robust
HbO
HbR

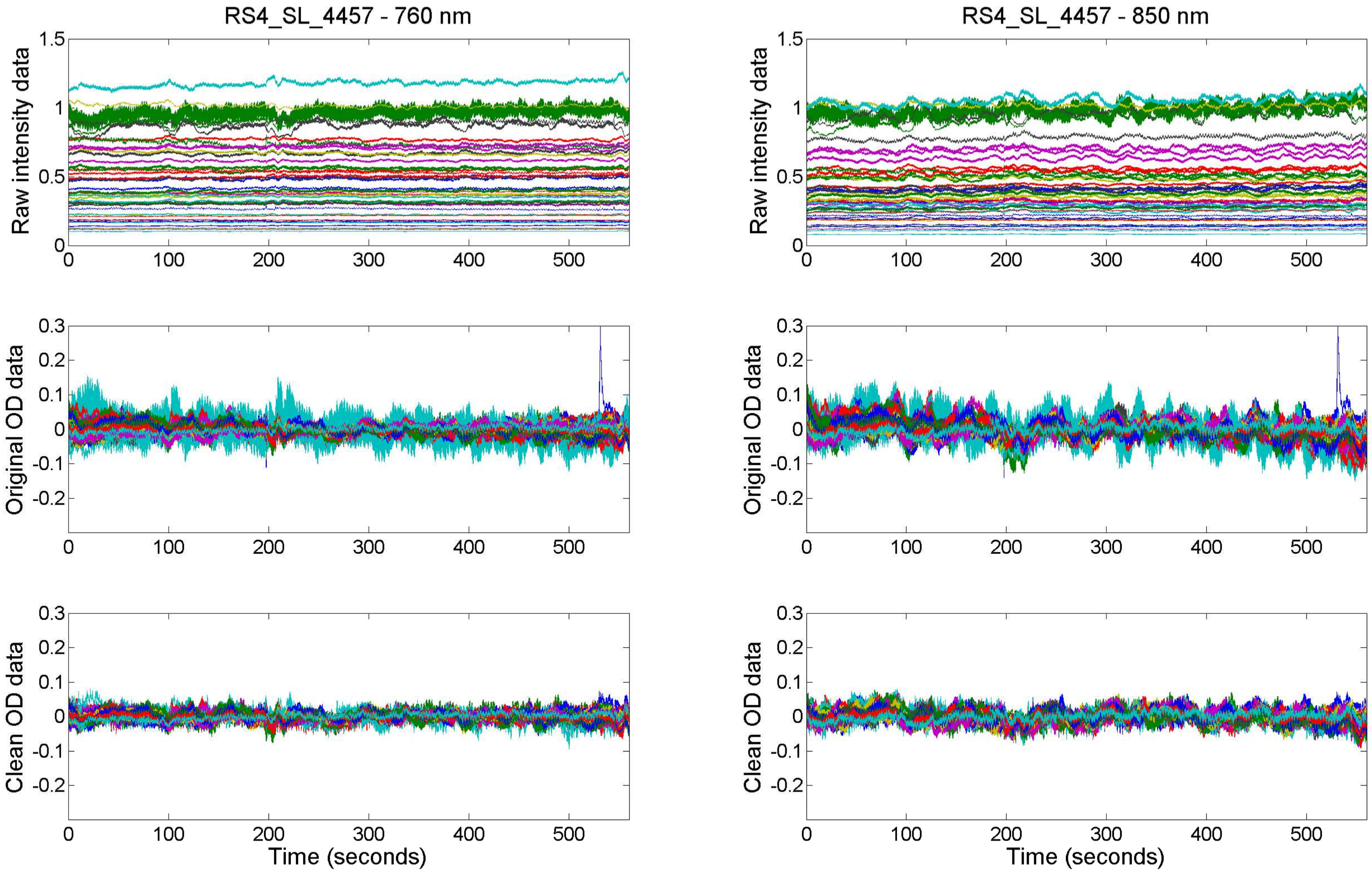

RS4_SL_4457 - 760 nm
RS4_SL_4457 - 850 nm
Raw intensity data
Original OD data
Clean OD data
Time (seconds)

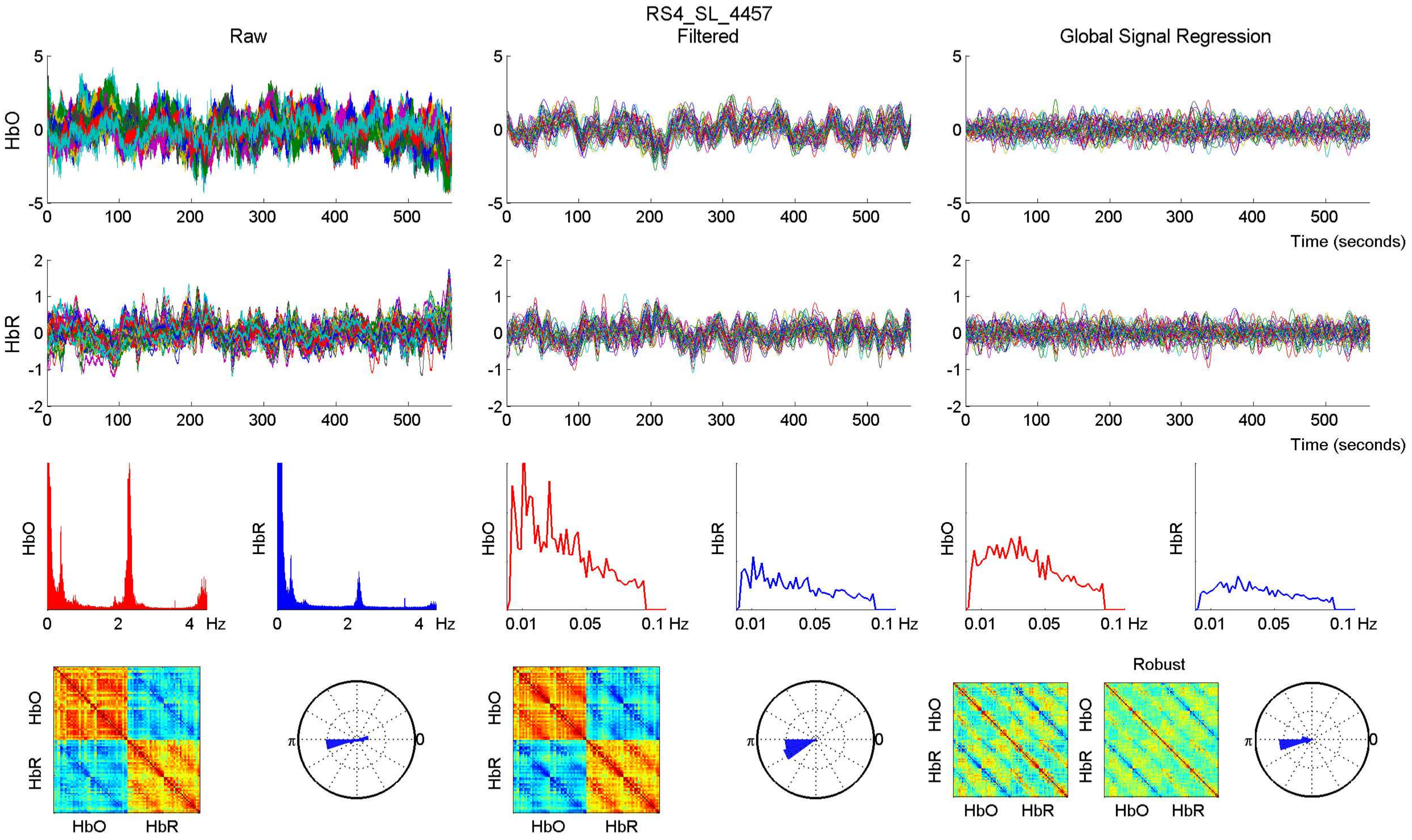

RS4_SL_4457
Raw
Filtered
Global Signal Regression
HbO
HbR
Time (seconds)
Hz
Robust

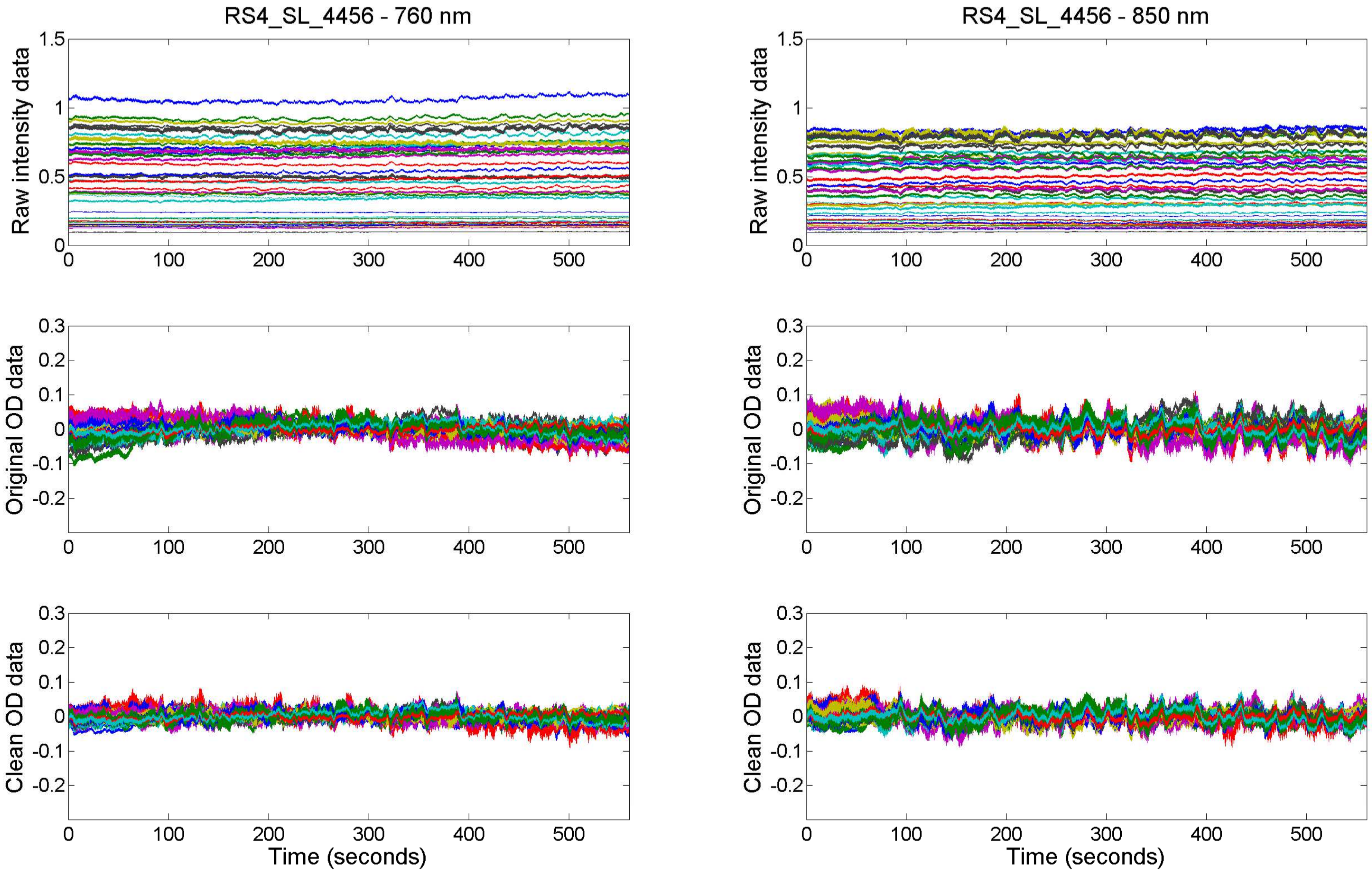

RS4_SL_4456 - 760 nm
RS4_SL_4456 - 850 nm
Raw intensity data
Original OD data
Clean OD data
Time (seconds)

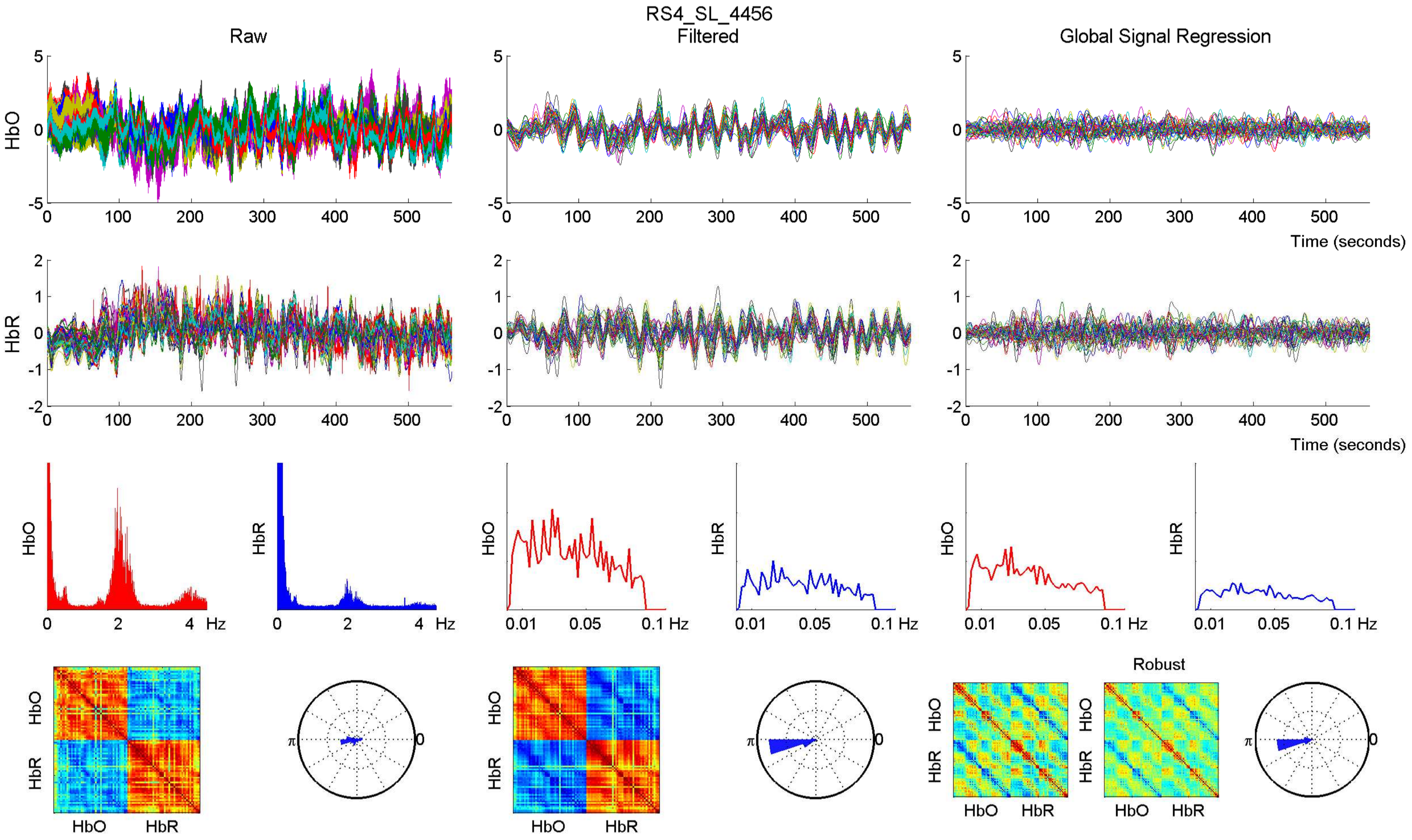

RS4_SL_4456
Raw
Filtered
Global Signal Regression
HbO
HbR
Time (seconds)
Hz
Robust

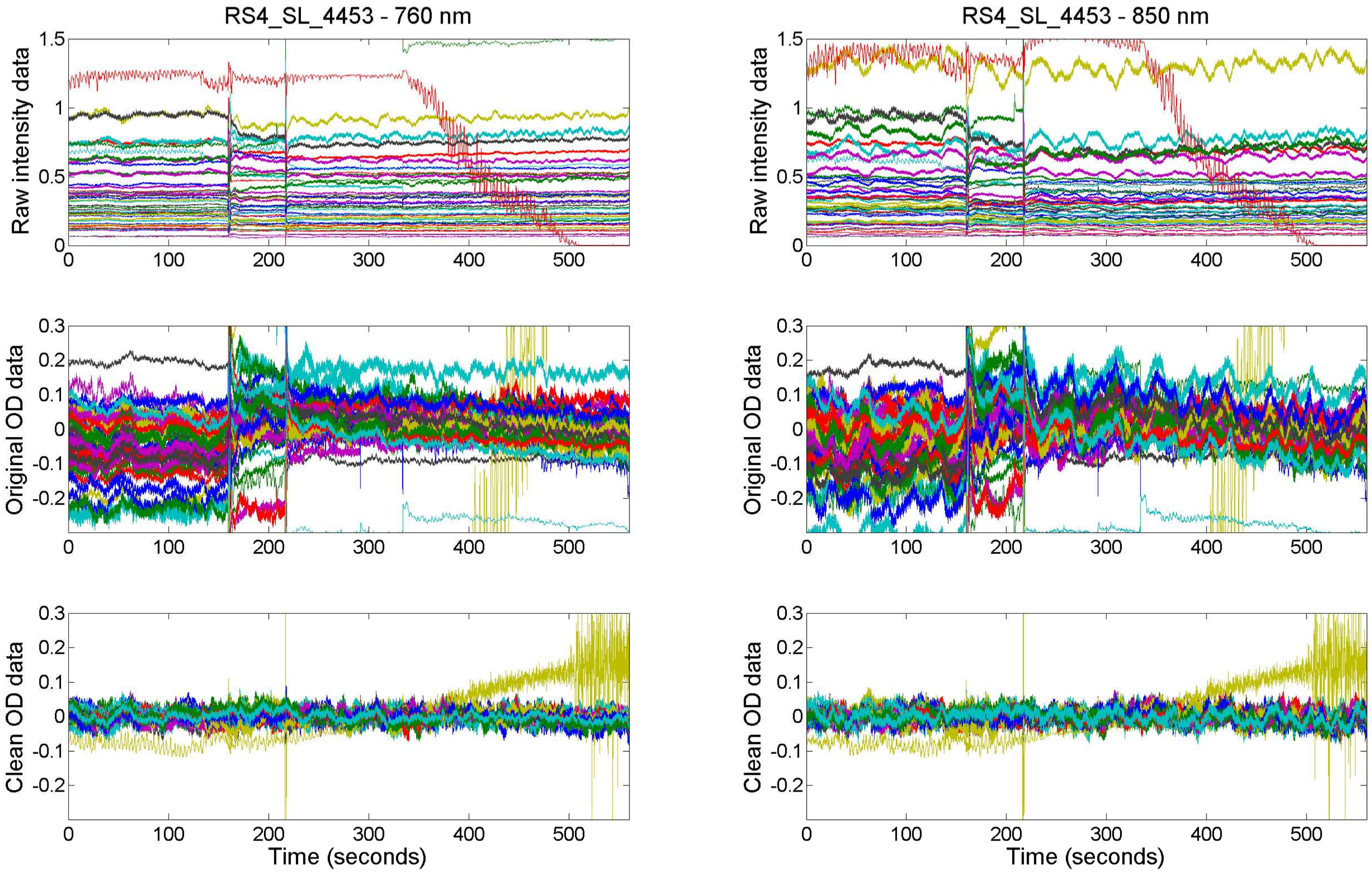

RS4_SL_4453 - 760 nm
RS4_SL_4453 - 850 nm
Raw intensity data
Original OD data
Clean OD data
Time (seconds)

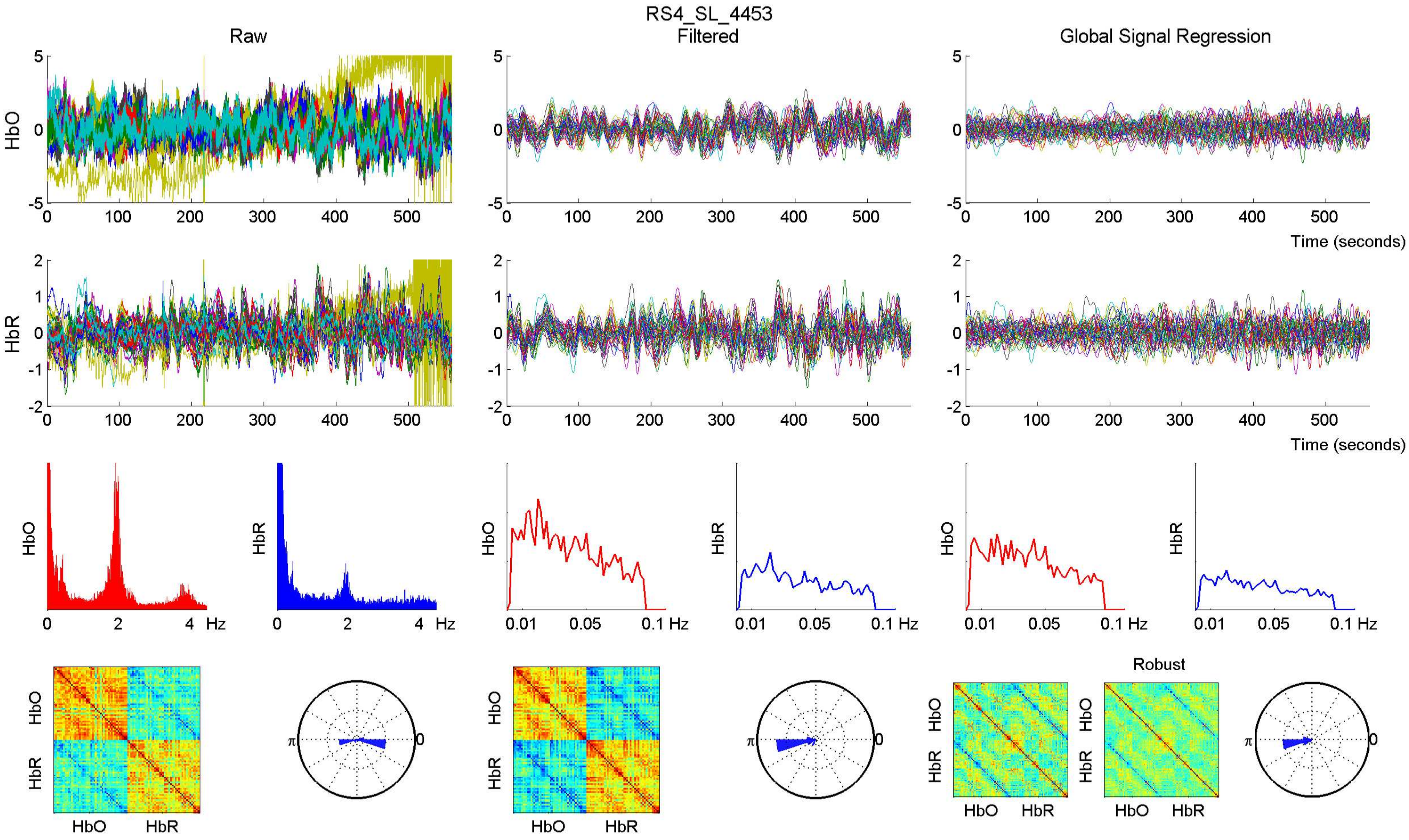
RS4_SL_4453
Raw
Filtered
Global Signal Regression
HbO
HbR
Time (seconds)
Hz
0.01
0.05
0.1 Hz
Robust
HbO
HbR
π
0

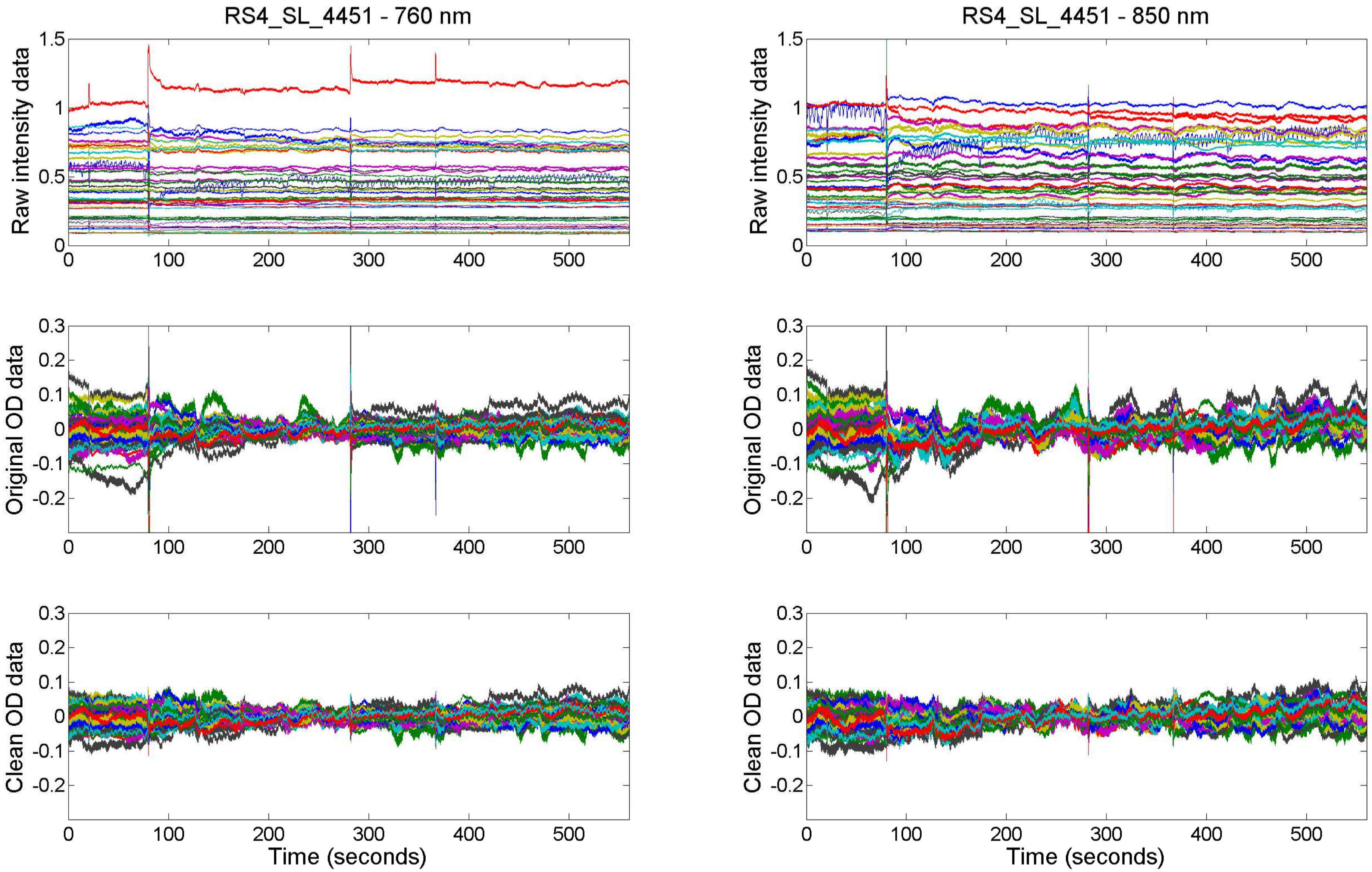

RS4_SL_4451 - 760 nm
RS4_SL_4451 - 850 nm
Raw intensity data
Original OD data
Clean OD data
Time (seconds)

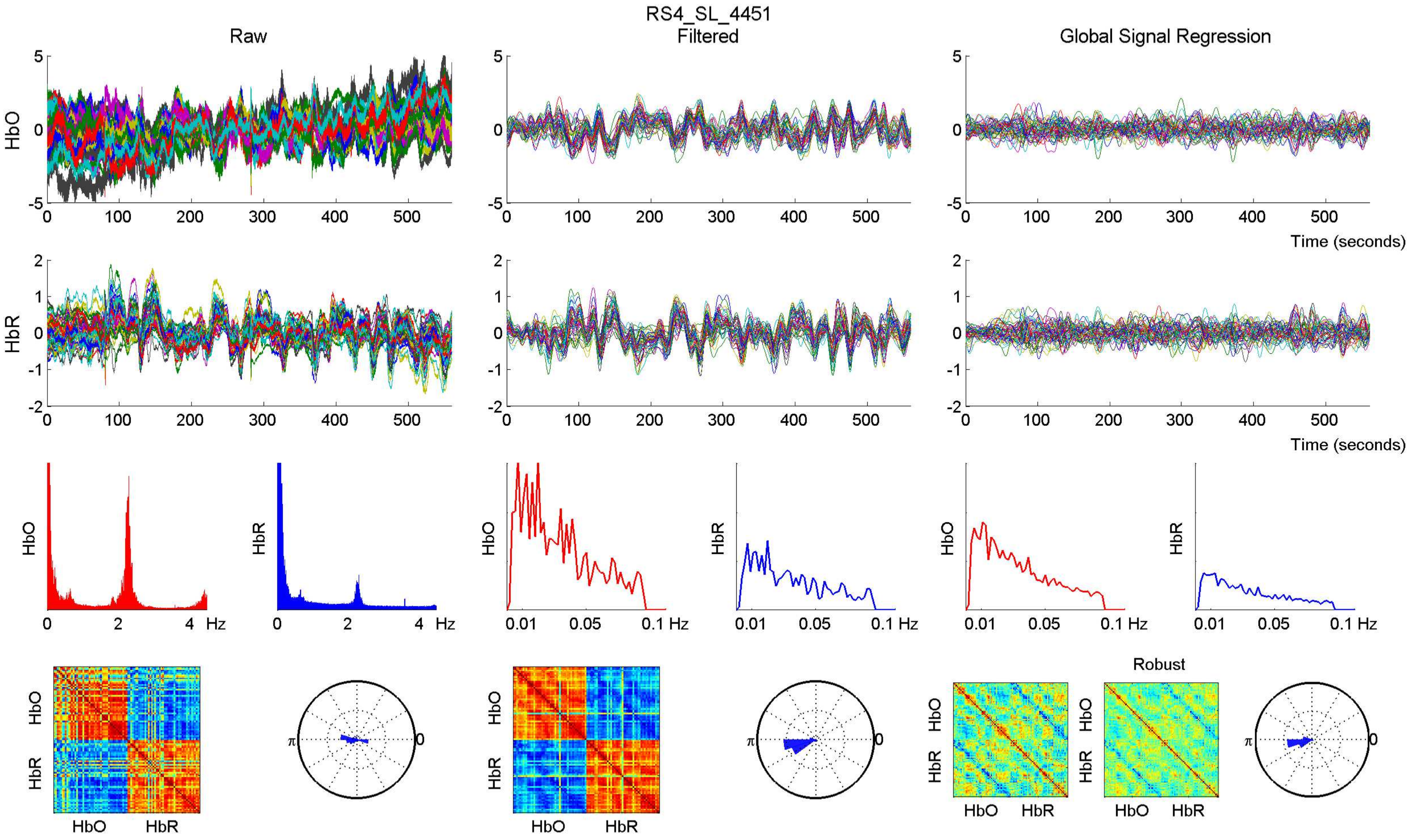

RS4_SL_4451
Raw
Filtered
Global Signal Regression
HbO
HbR
Time (seconds)
Hz
Robust

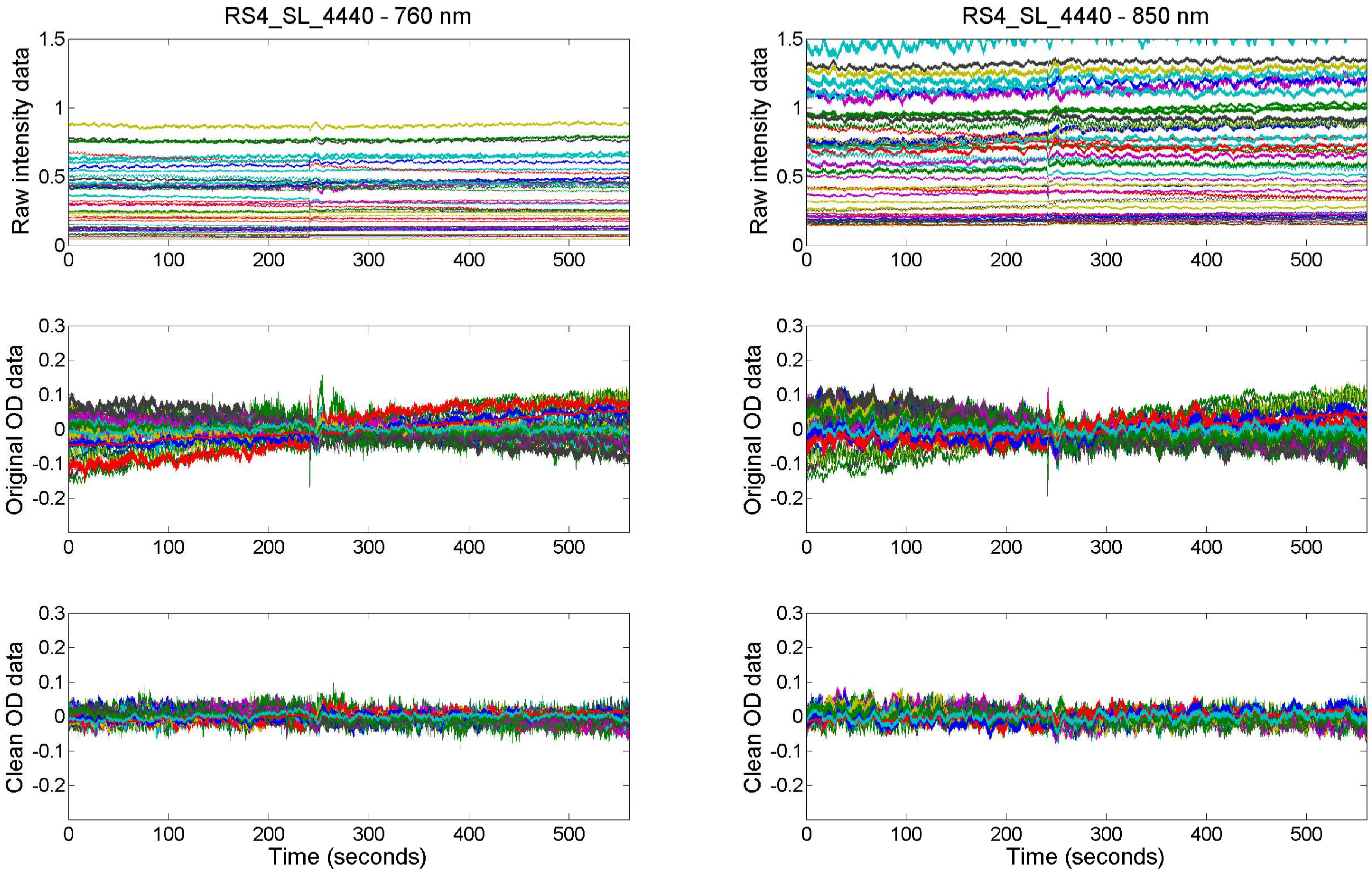

RS4_SL_4440 - 760 nm
RS4_SL_4440 - 850 nm
Raw intensity data
Original OD data
Clean OD data
Time (seconds)

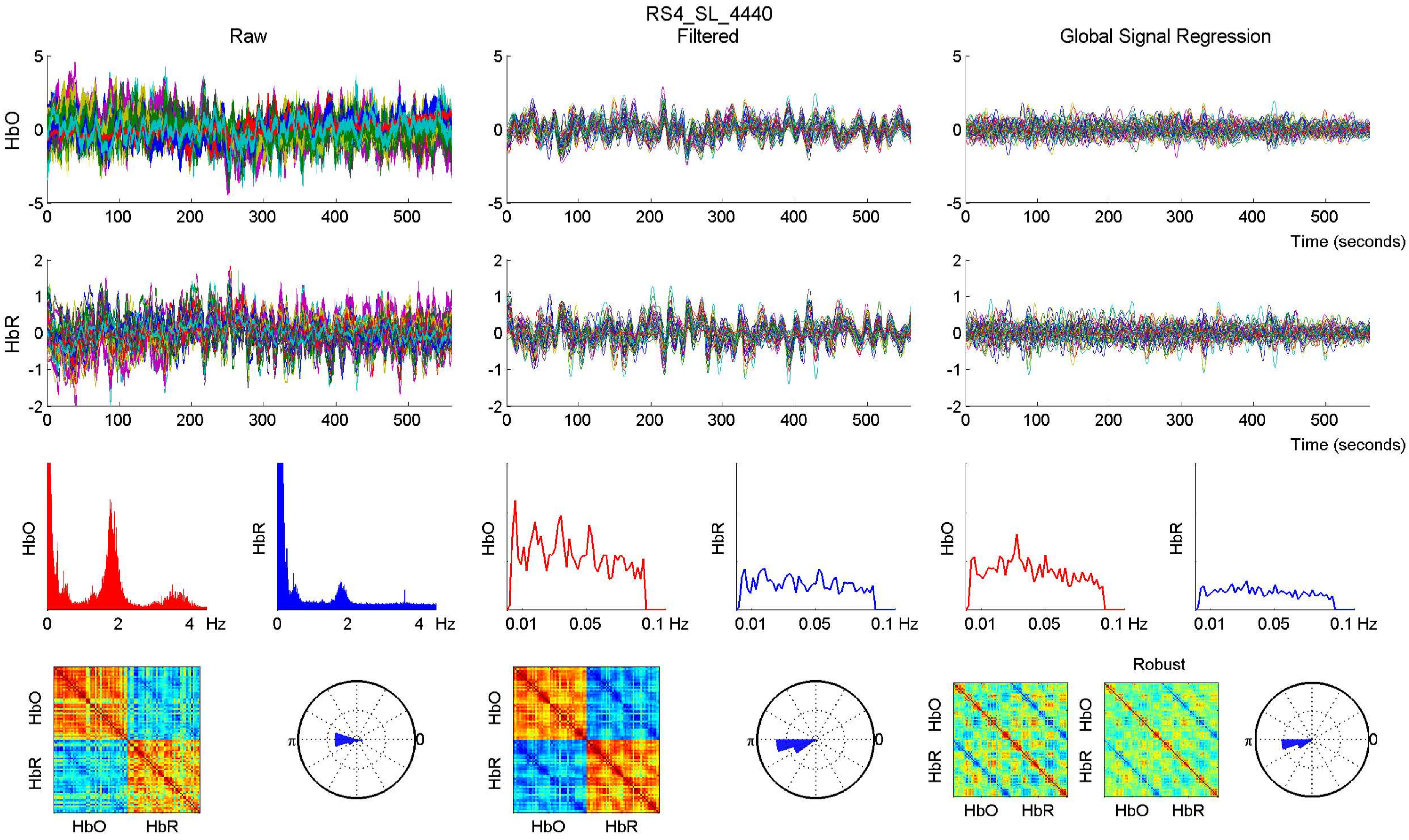
RS4_SL_4440
Raw
Filtered
Global Signal Regression
HbO
HbR
Time (seconds)
Hz
0.01
0.05
0.1 Hz
Robust
π
0

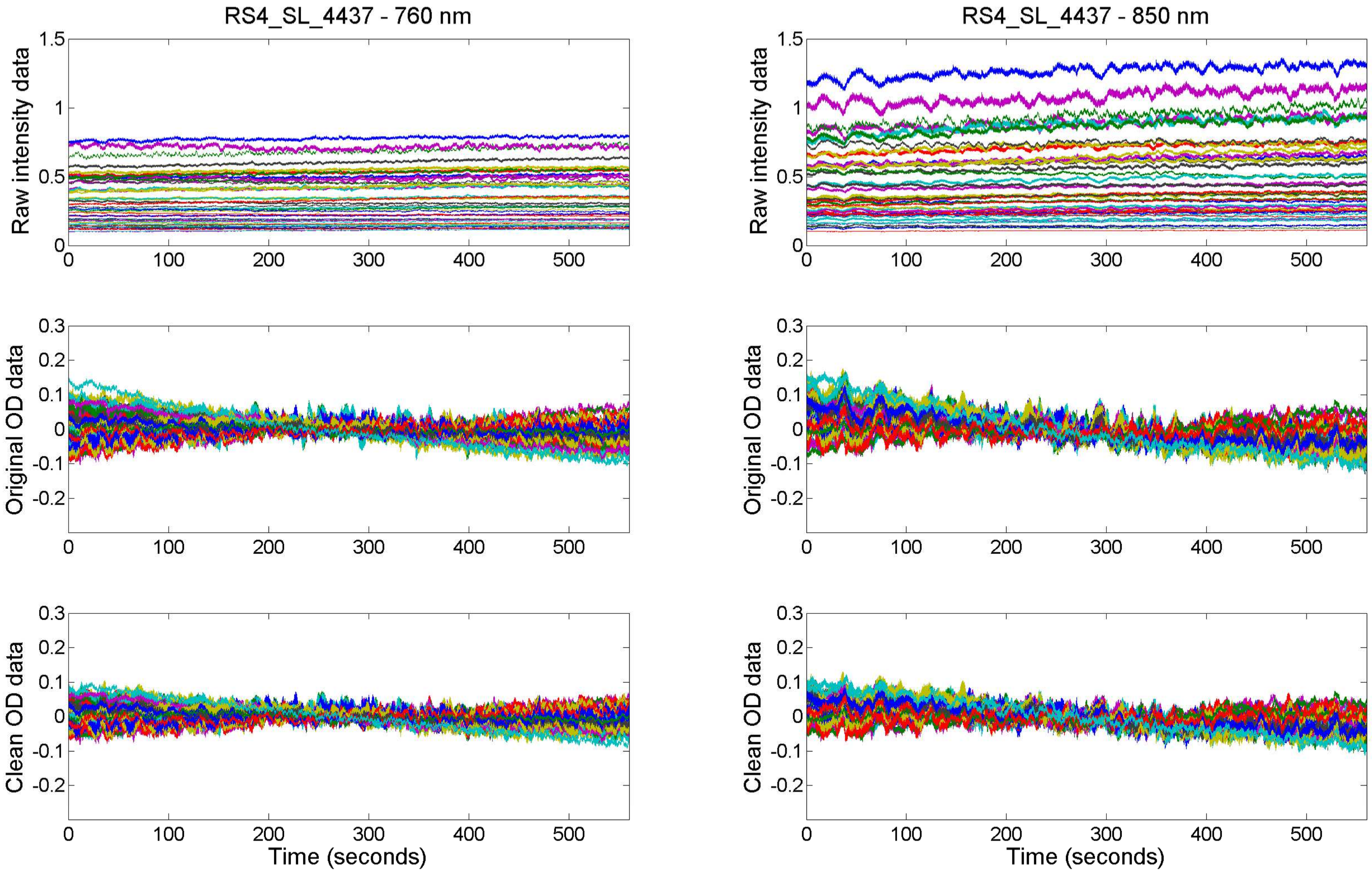

RS4_SL_4437 - 760 nm
RS4_SL_4437 - 850 nm
Raw intensity data
Original OD data
Clean OD data
Time (seconds)

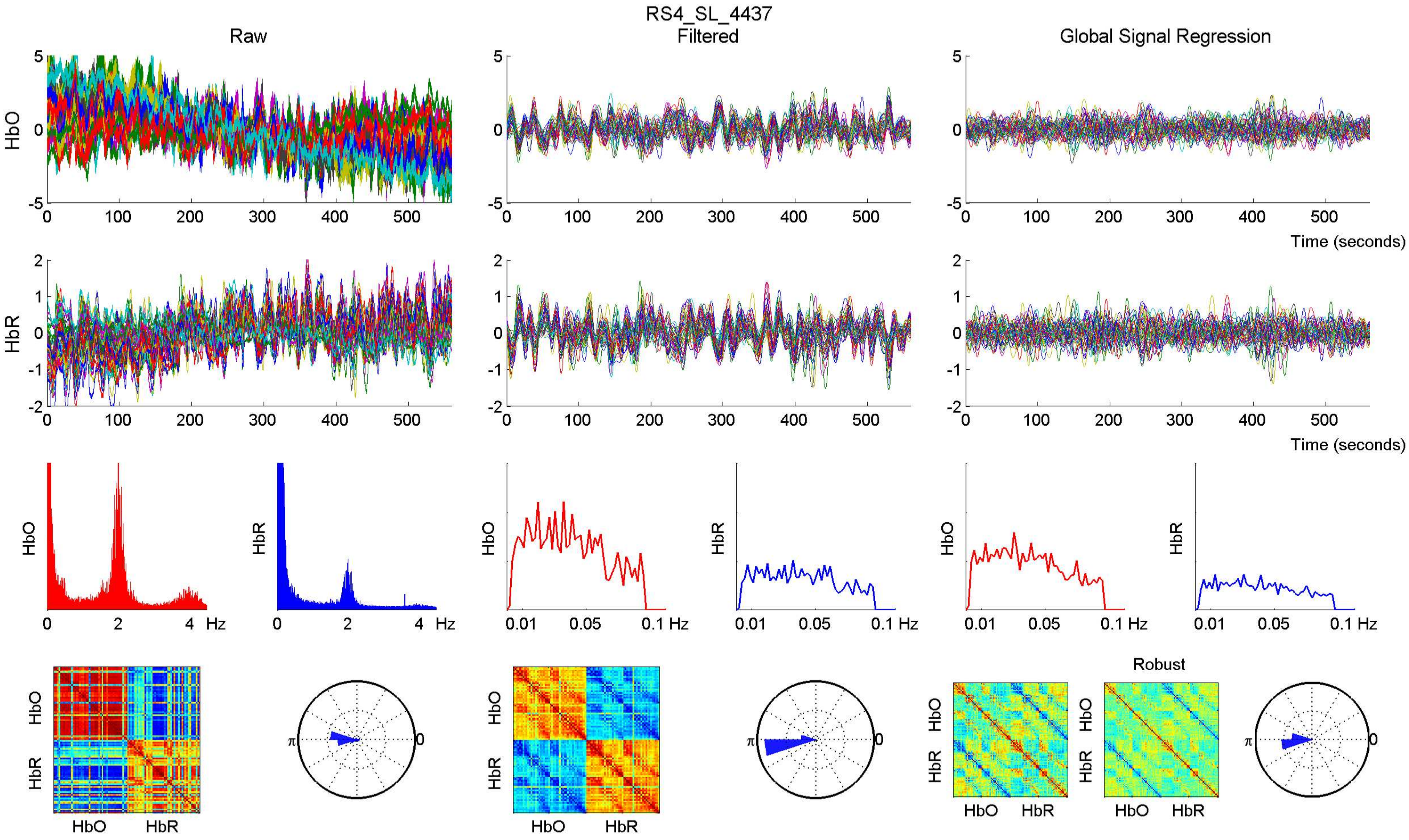

RS4_SL_4437
Raw
Filtered
Global Signal Regression
HbO
HbR
Time (seconds)
Hz
Robust
HbO
HbR

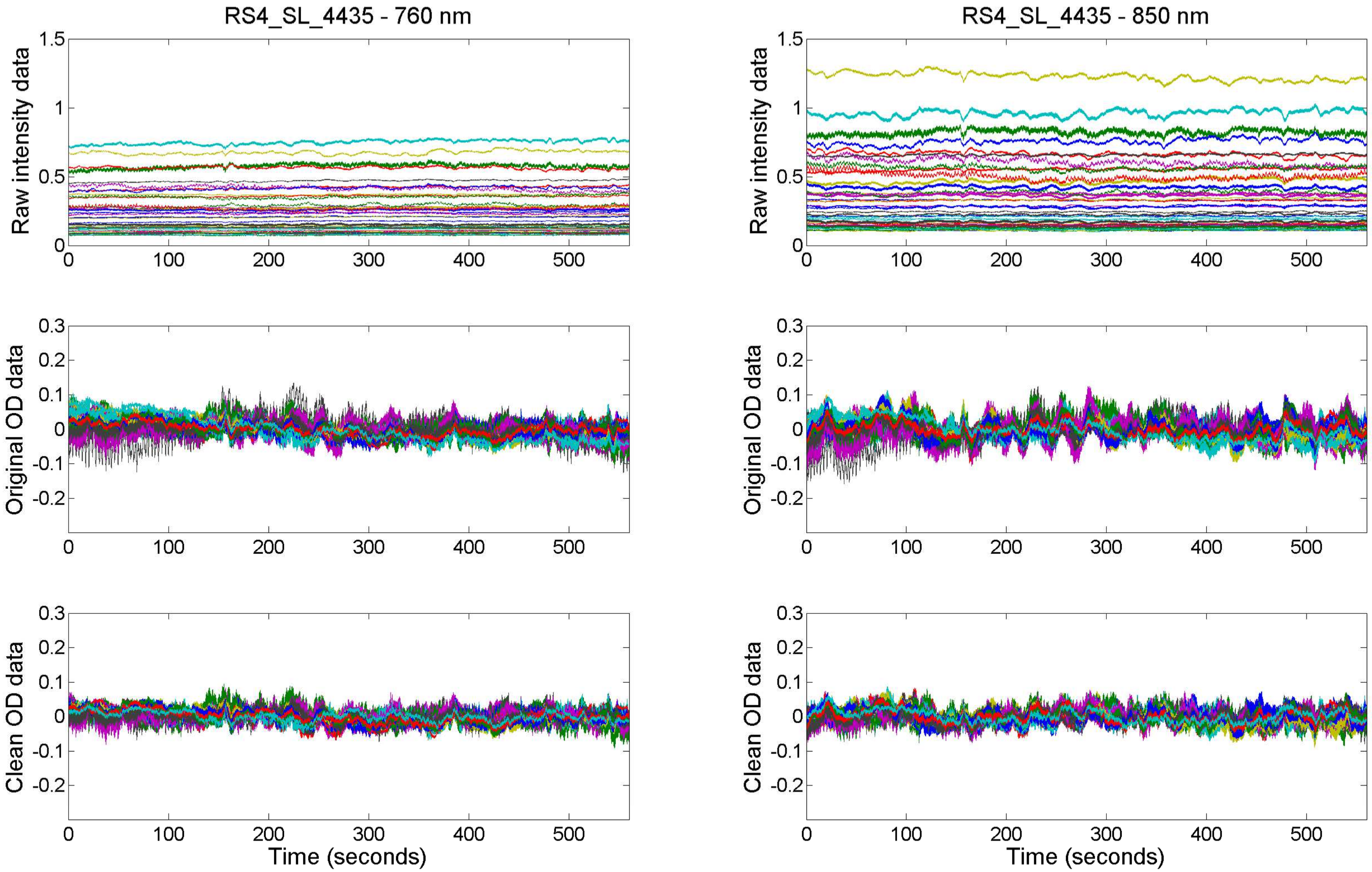

RS4_SL_4435 - 760 nm
RS4_SL_4435 - 850 nm
Raw intensity data
Original OD data
Clean OD data
Time (seconds)

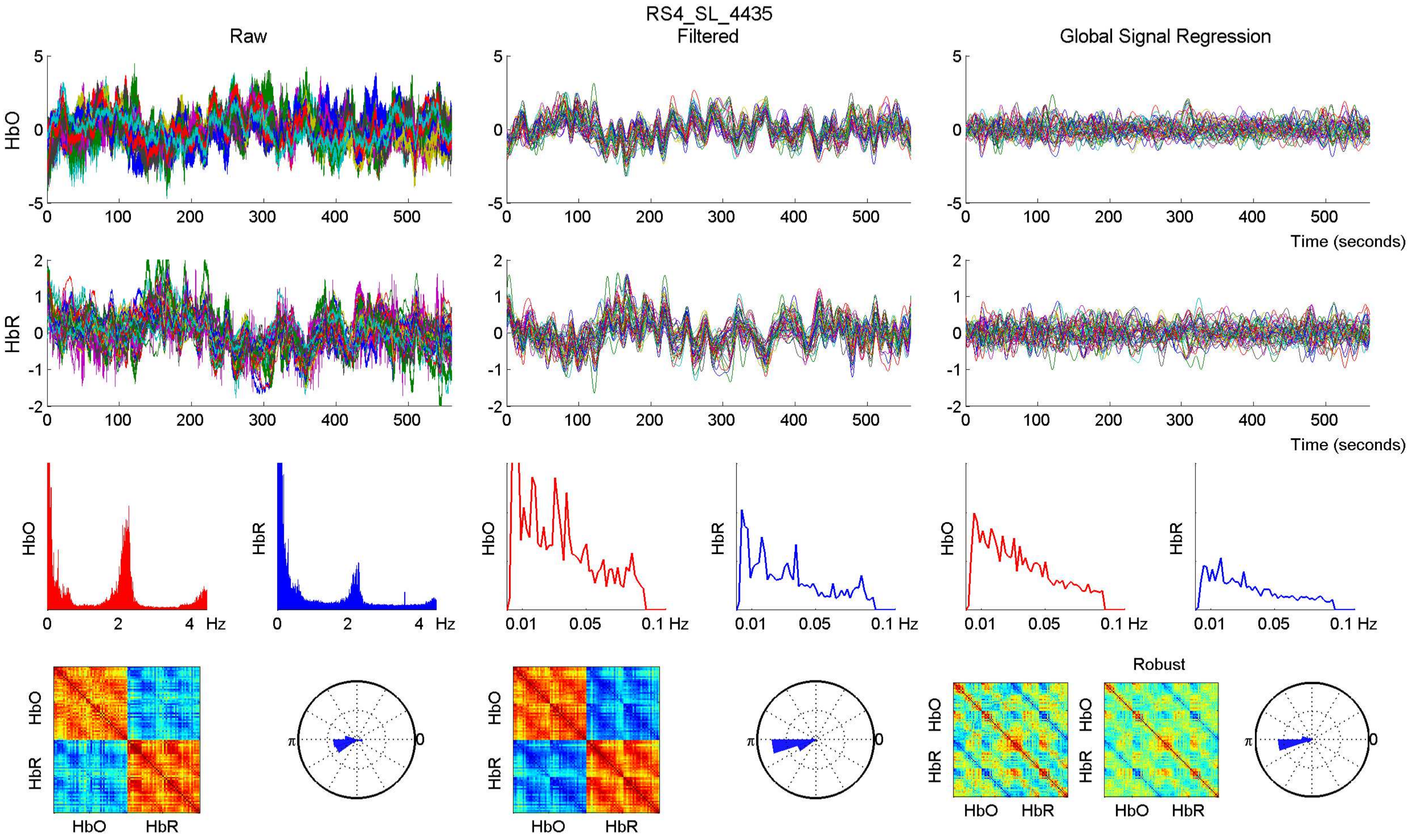

RS4_SL_4435
Raw
Filtered
Global Signal Regression
HbO
HbR
Time (seconds)
Hz
Robust
HbO
HbR

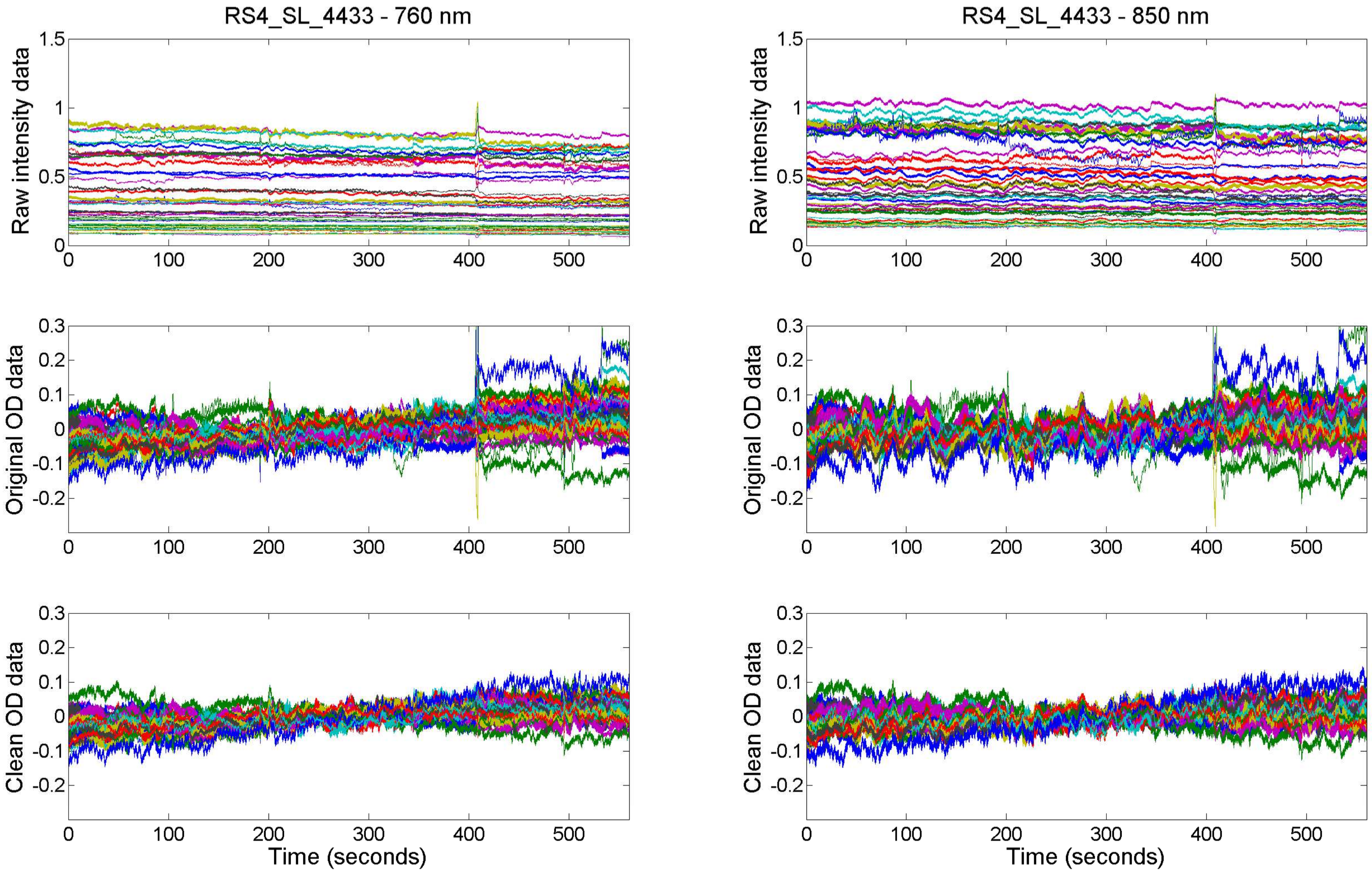

RS4_SL_4433 - 760 nm
RS4_SL_4433 - 850 nm
Raw intensity data
Original OD data
Clean OD data
Time (seconds)

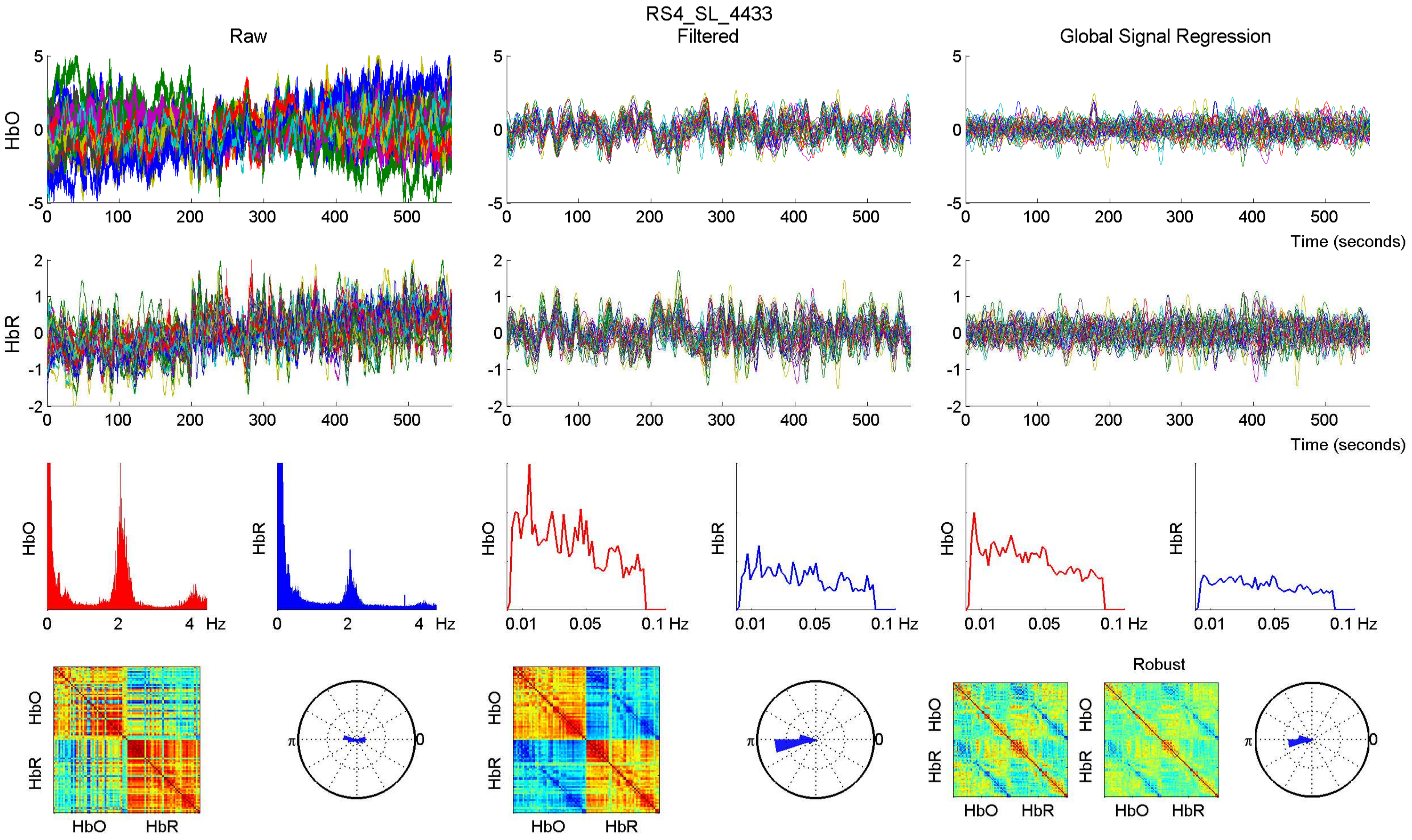
RS4_SL_4433
Raw
Filtered
Global Signal Regression
HbO
HbR
Time (seconds)
Hz
Robust
HbO
HbR

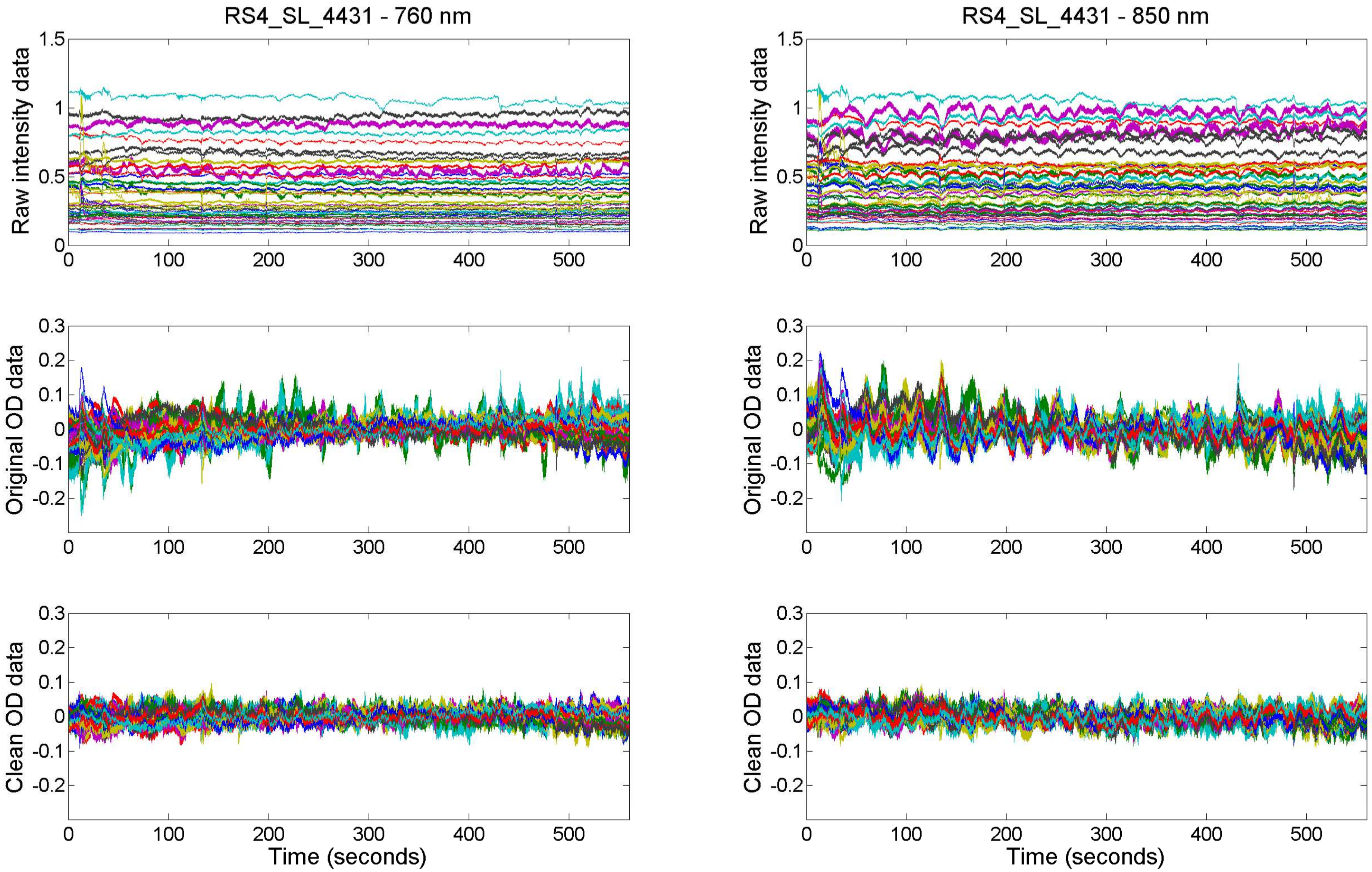
RS4_SL_4431 - 760 nm
RS4_SL_4431 - 850 nm
Raw intensity data
Original OD data
Clean OD data
Time (seconds)

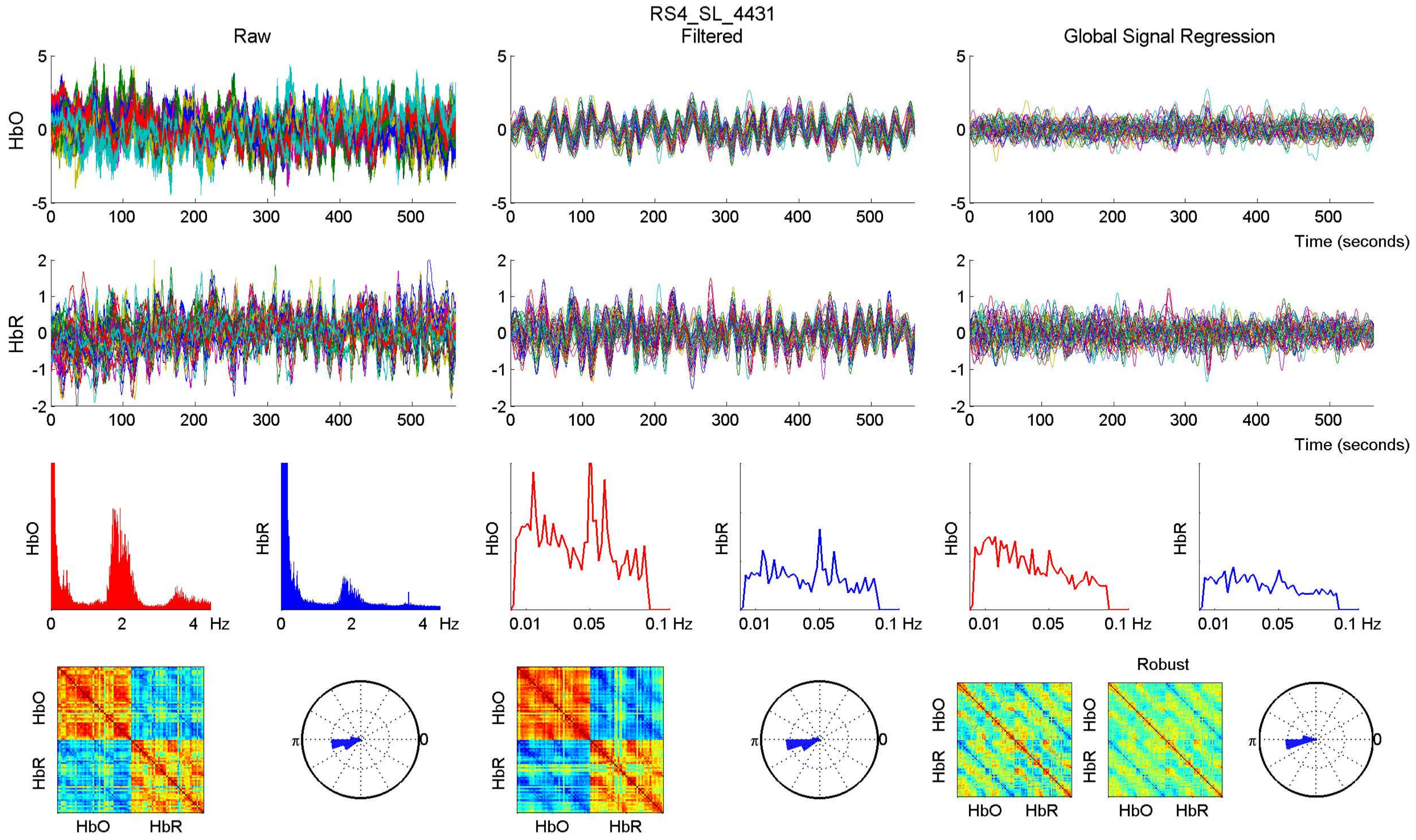

RS4_SL_4431
Raw
Filtered
Global Signal Regression
HbO
HbR
Time (seconds)
Hz
Robust

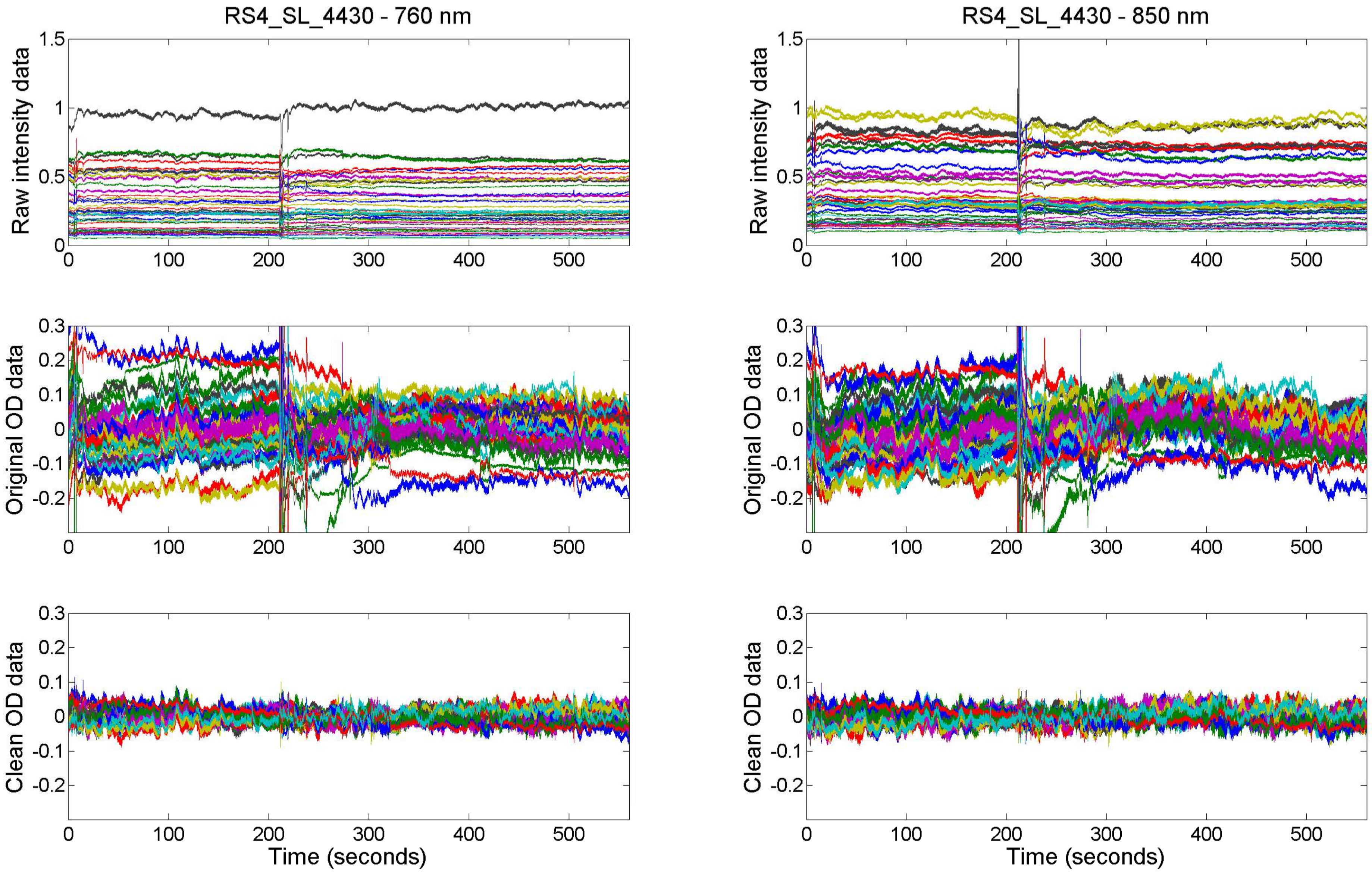

RS4_SL_4430 - 760 nm
RS4_SL_4430 - 850 nm
Raw intensity data
Original OD data
Clean OD data
Time (seconds)

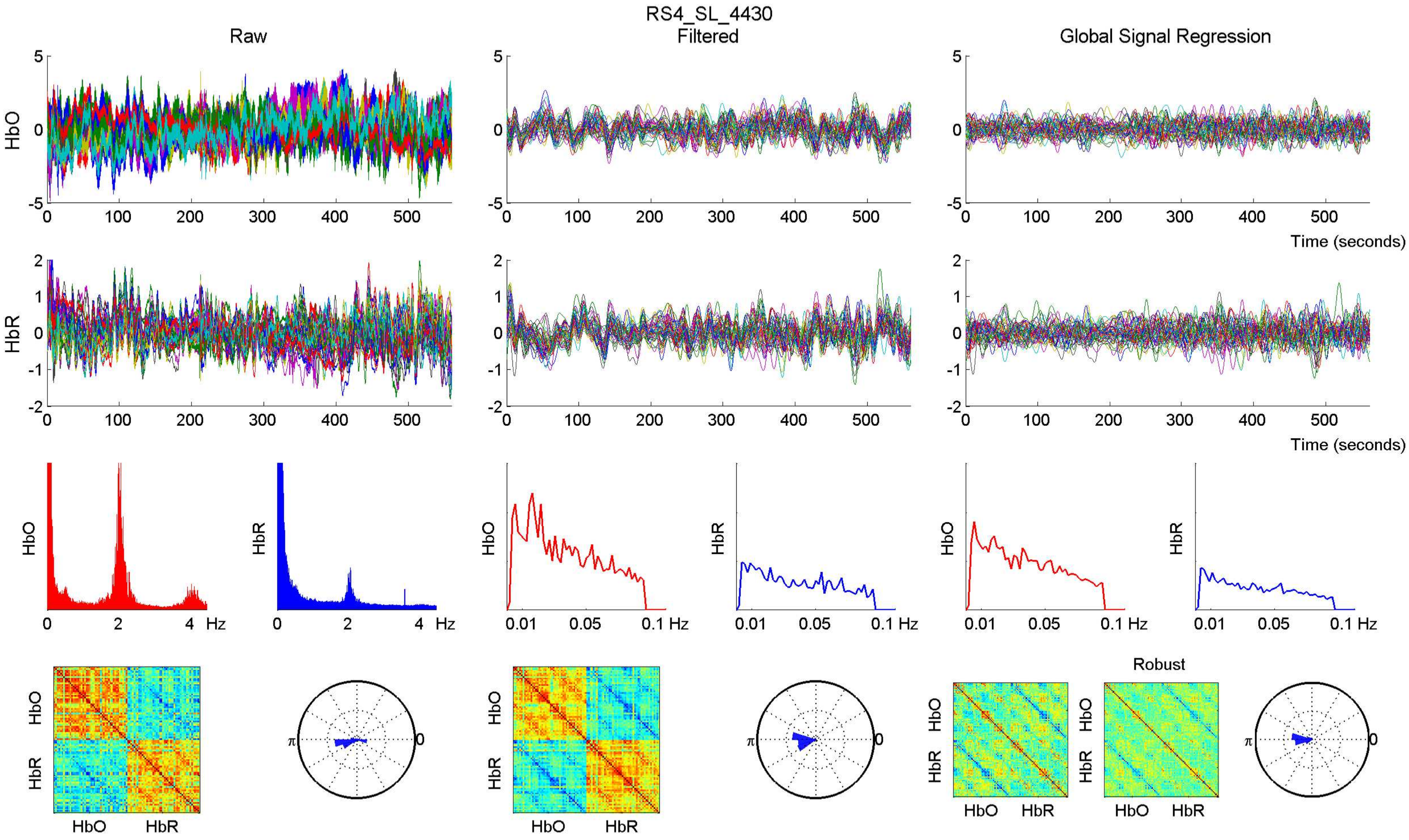

RS4_SL_4430
Raw
Filtered
Global Signal Regression
HbO
HbR
Time (seconds)
Hz
Robust

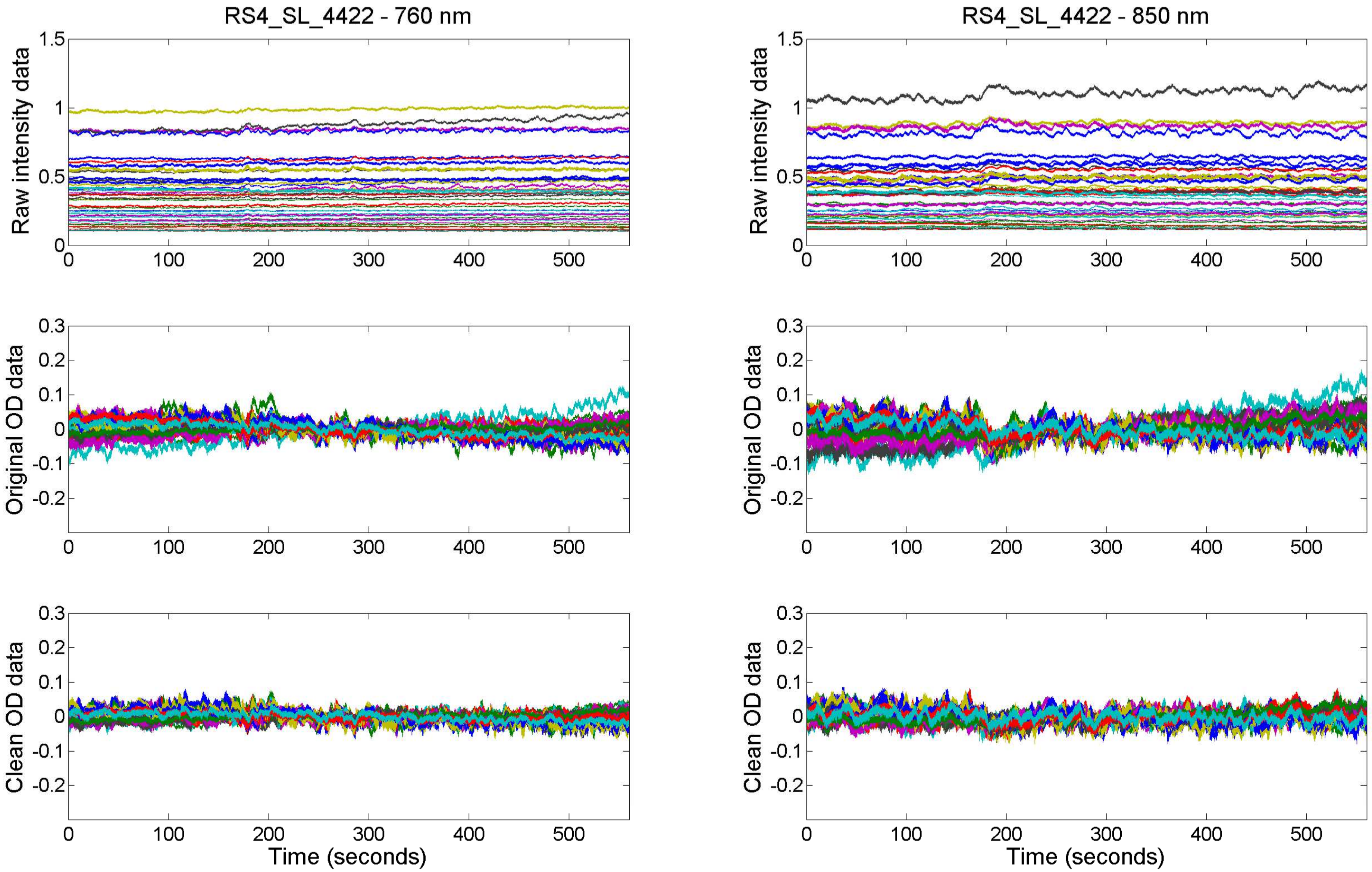

RS4_SL_4422 - 760 nm
RS4_SL_4422 - 850 nm
Raw intensity data
Original OD data
Clean OD data
Time (seconds)

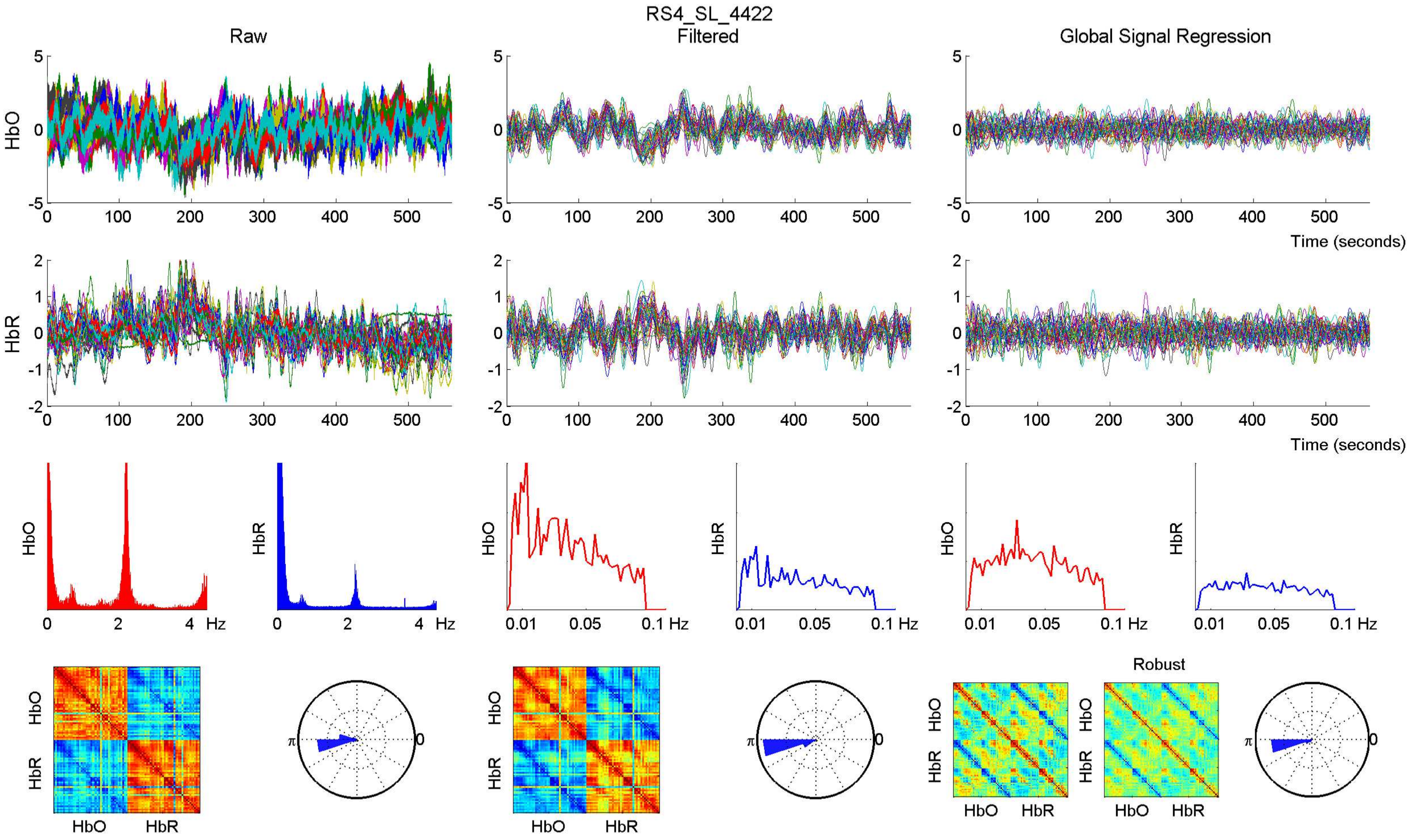

RS4_SL_4422
Raw
Filtered
Global Signal Regression
HbO
HbR
Time (seconds)
Hz
Robust
HbO
HbR

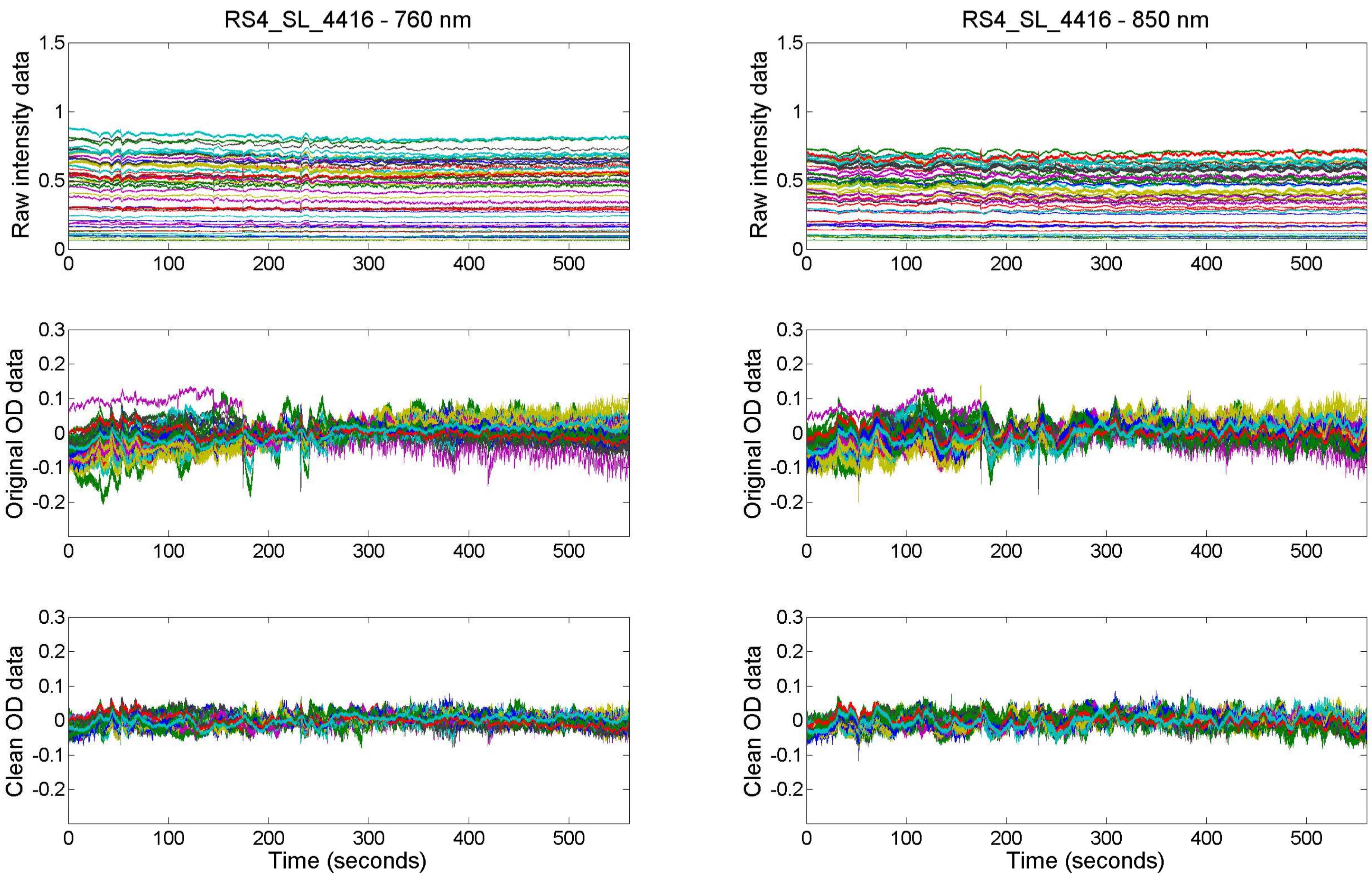
RS4_SL_4416 - 760 nm
RS4_SL_4416 - 850 nm
Raw intensity data
Original OD data
Clean OD data
Time (seconds)
Time (seconds)

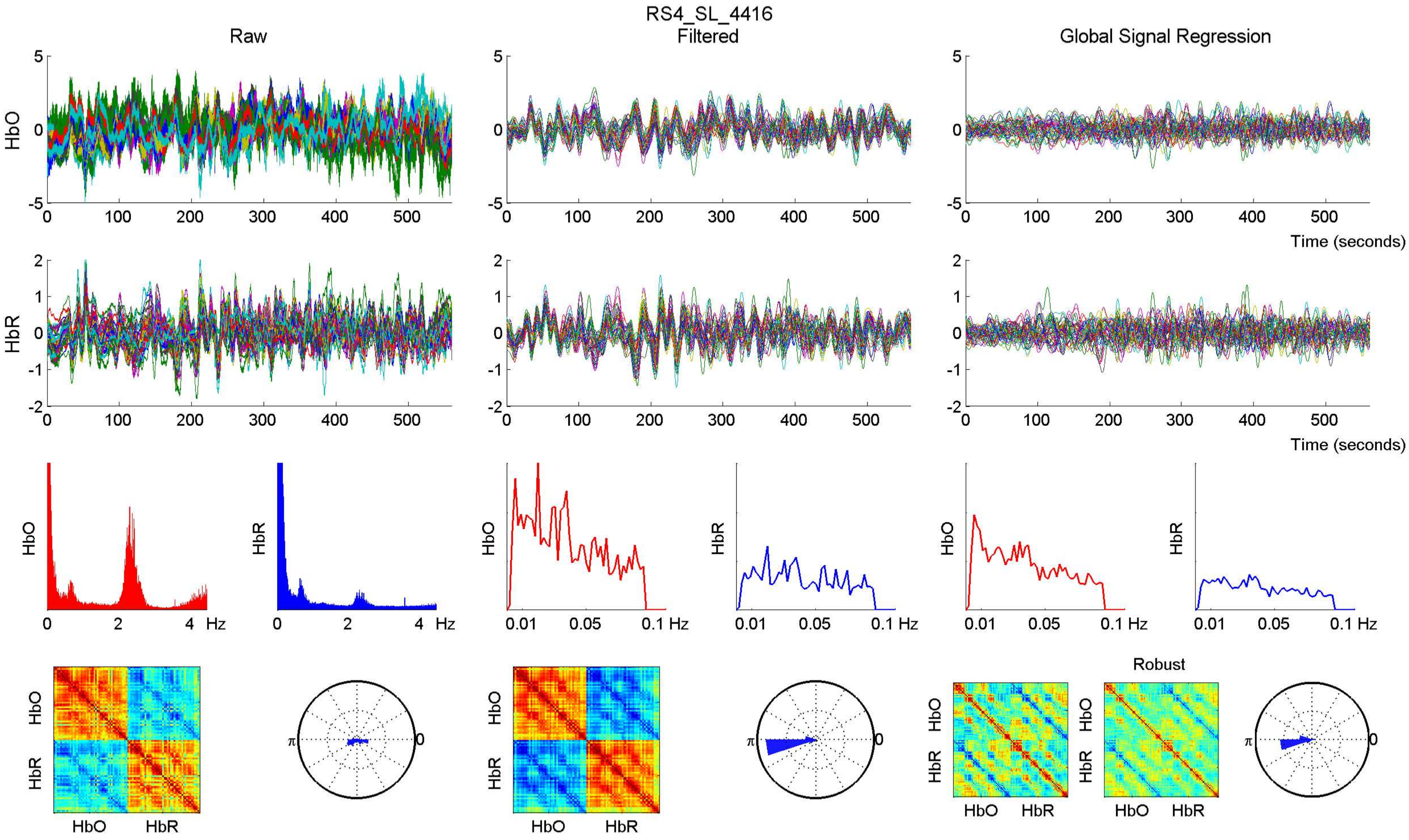
RS4_SL_4416
Raw
Filtered
Global Signal Regression
HbO
HbR
Time (seconds)
Hz
Robust
HbO
HbR

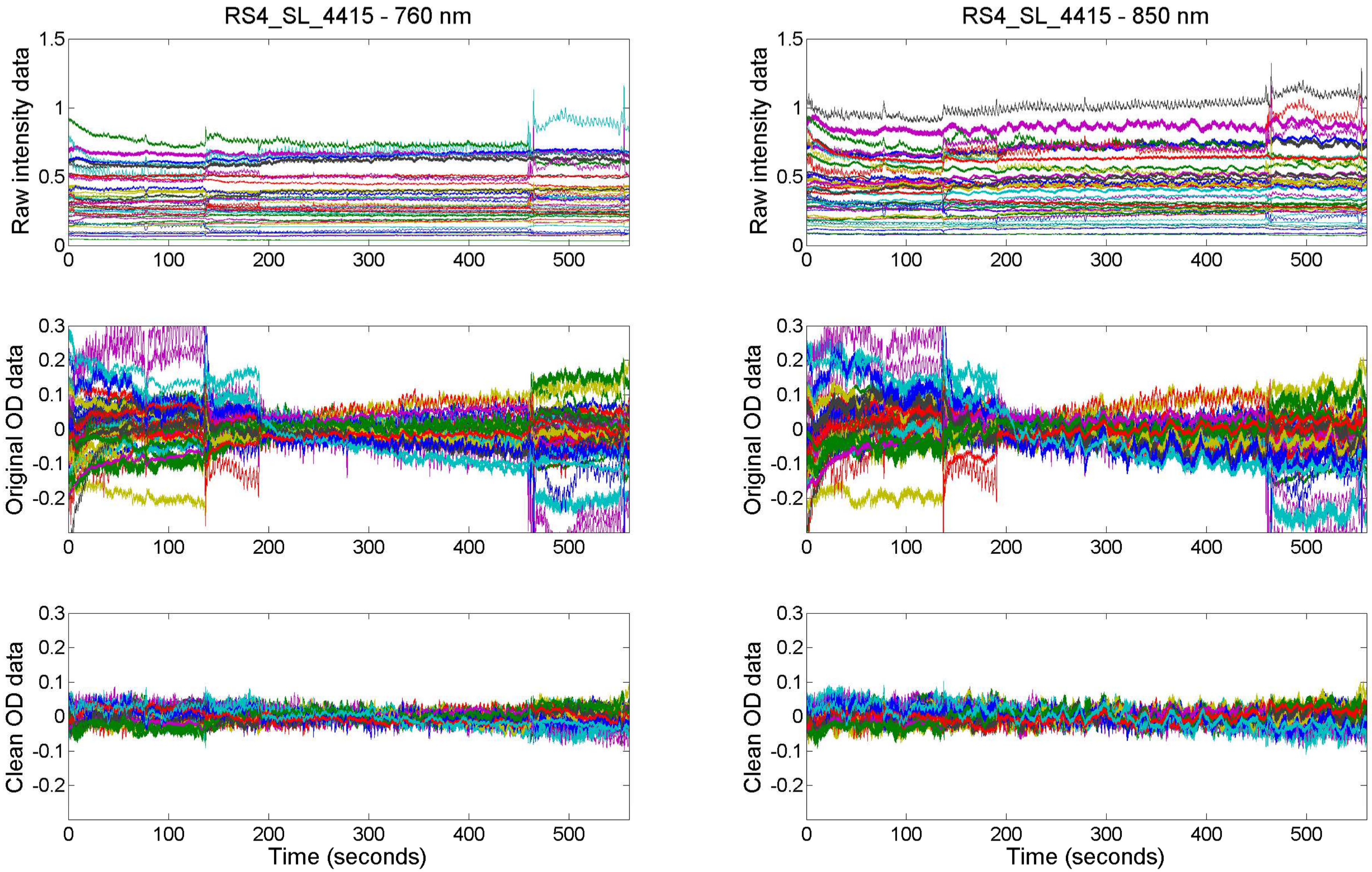

RS4_SL_4415 - 760 nm
RS4_SL_4415 - 850 nm
Raw intensity data
Original OD data
Clean OD data
Time (seconds)

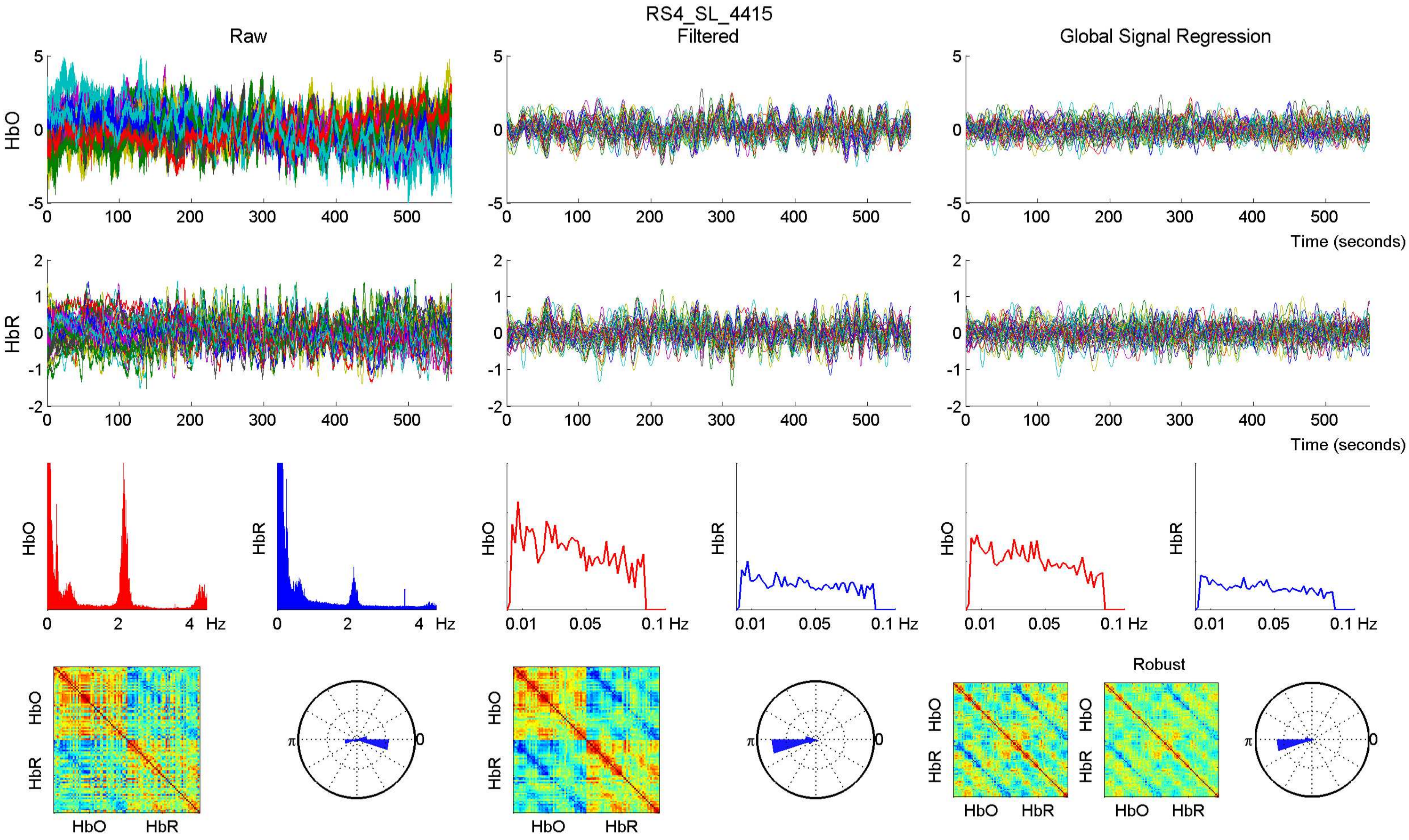

RS4_SL_4415
Raw
Filtered
Global Signal Regression
HbO
HbR
Time (seconds)
Hz
0.01 0.05 0.1 Hz
Robust
HbO HbR
π
0

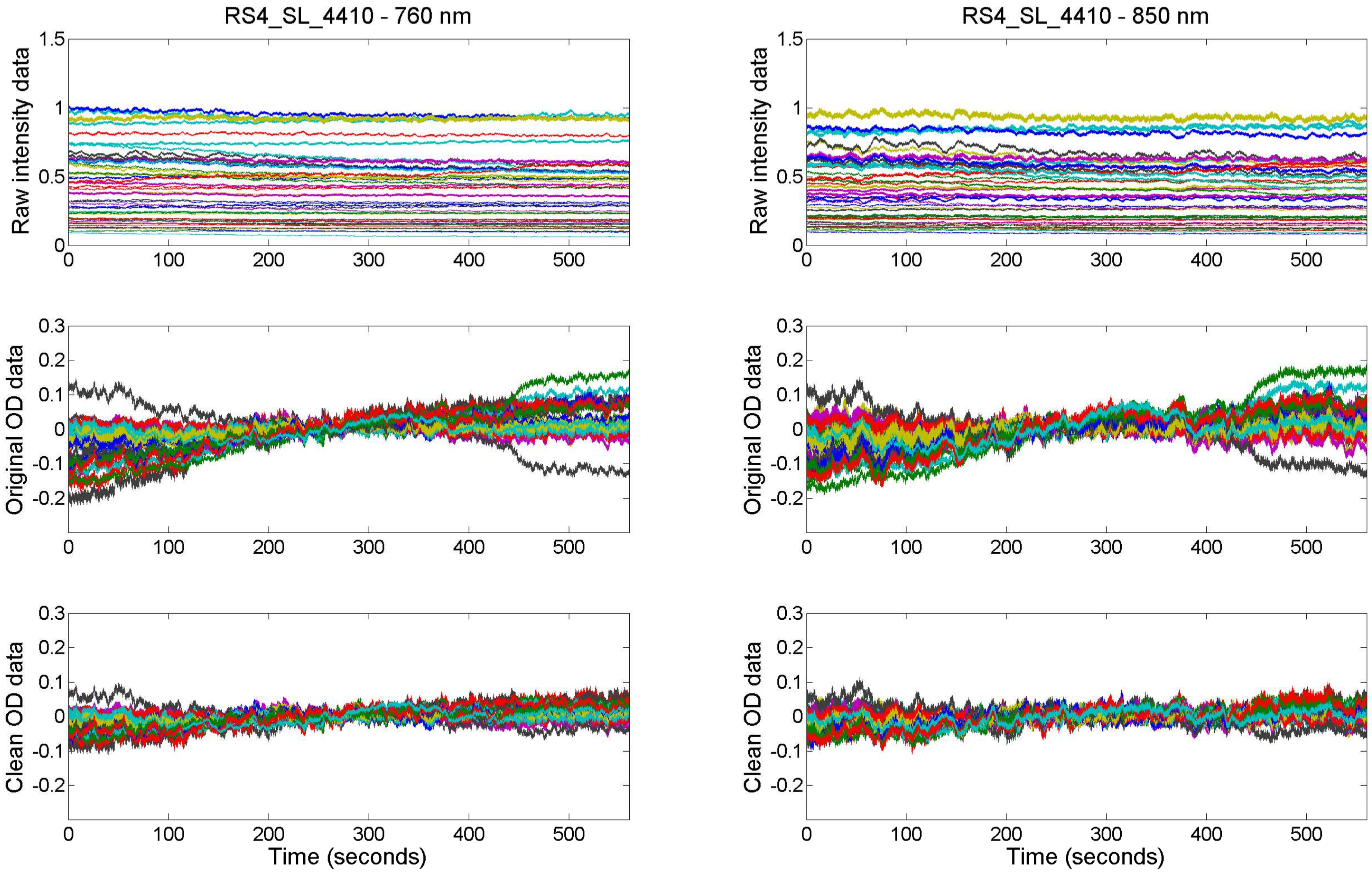

RS4_SL_4410 - 760 nm
RS4_SL_4410 - 850 nm
Raw intensity data
Original OD data
Clean OD data
Time (seconds)

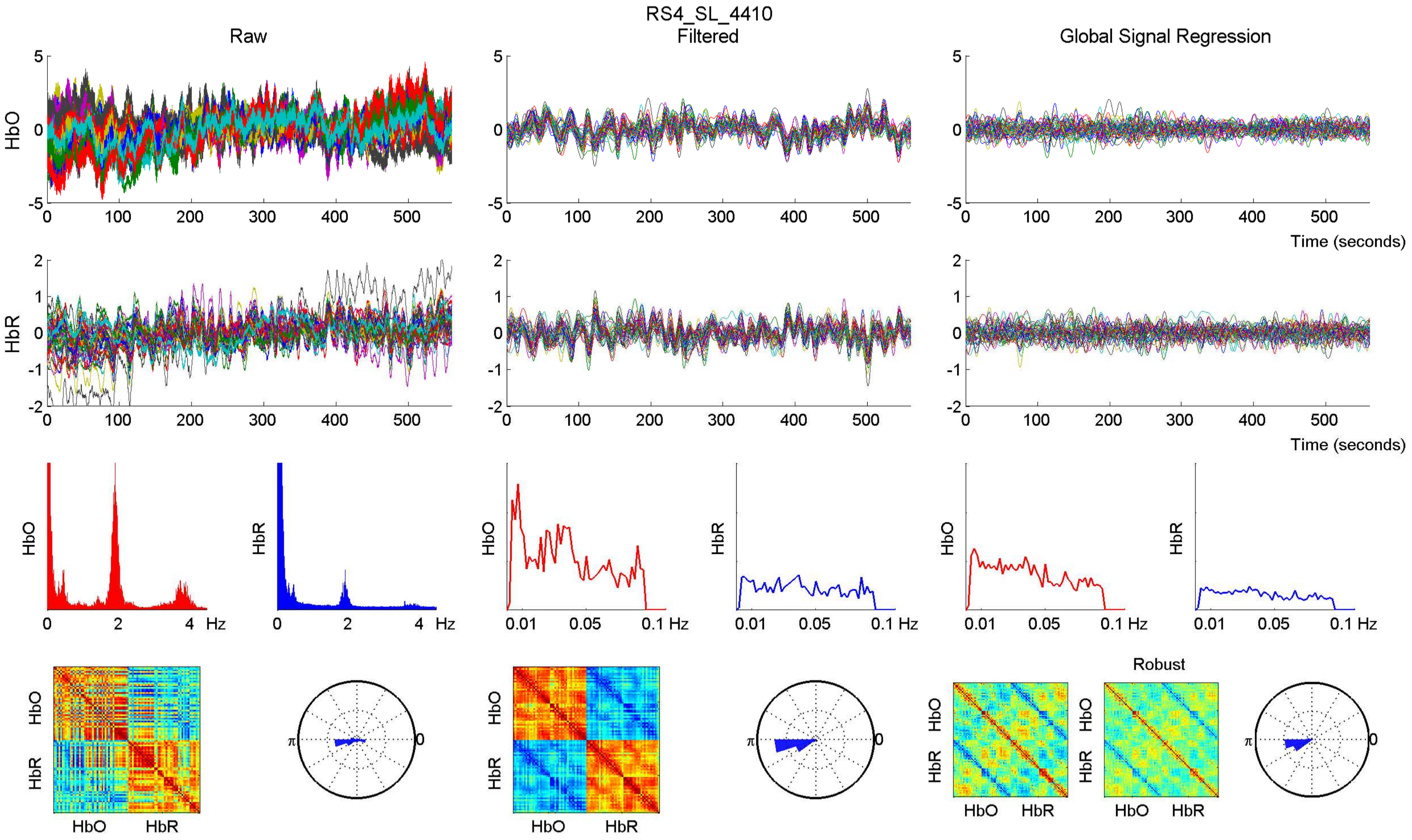

RS4_SL_4410
Raw
Filtered
Global Signal Regression
HbO
HbR
Time (seconds)
Hz
Robust

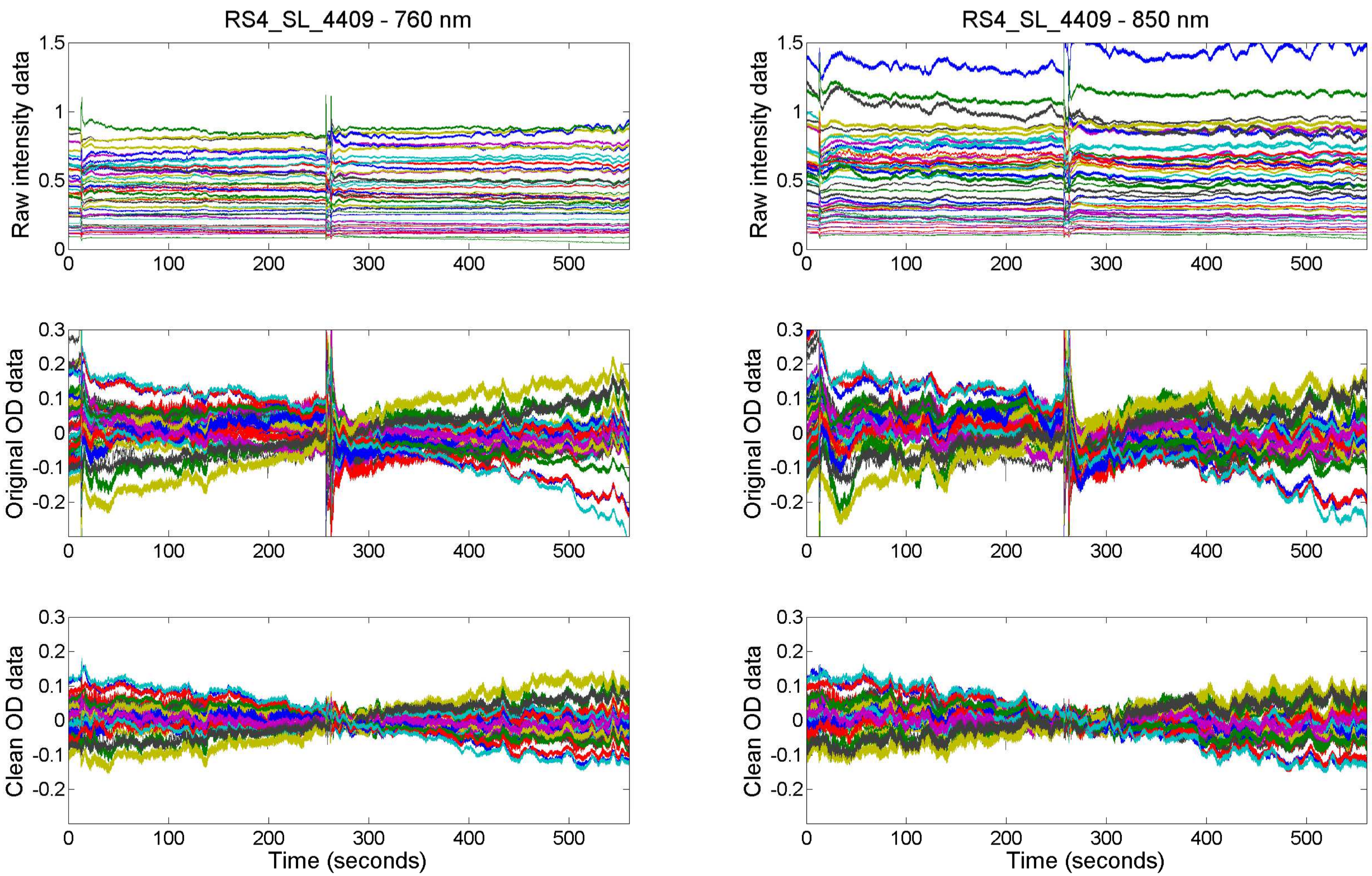

RS4_SL_4409 - 760 nm
RS4_SL_4409 - 850 nm
Raw intensity data
Original OD data
Clean OD data
Time (seconds)

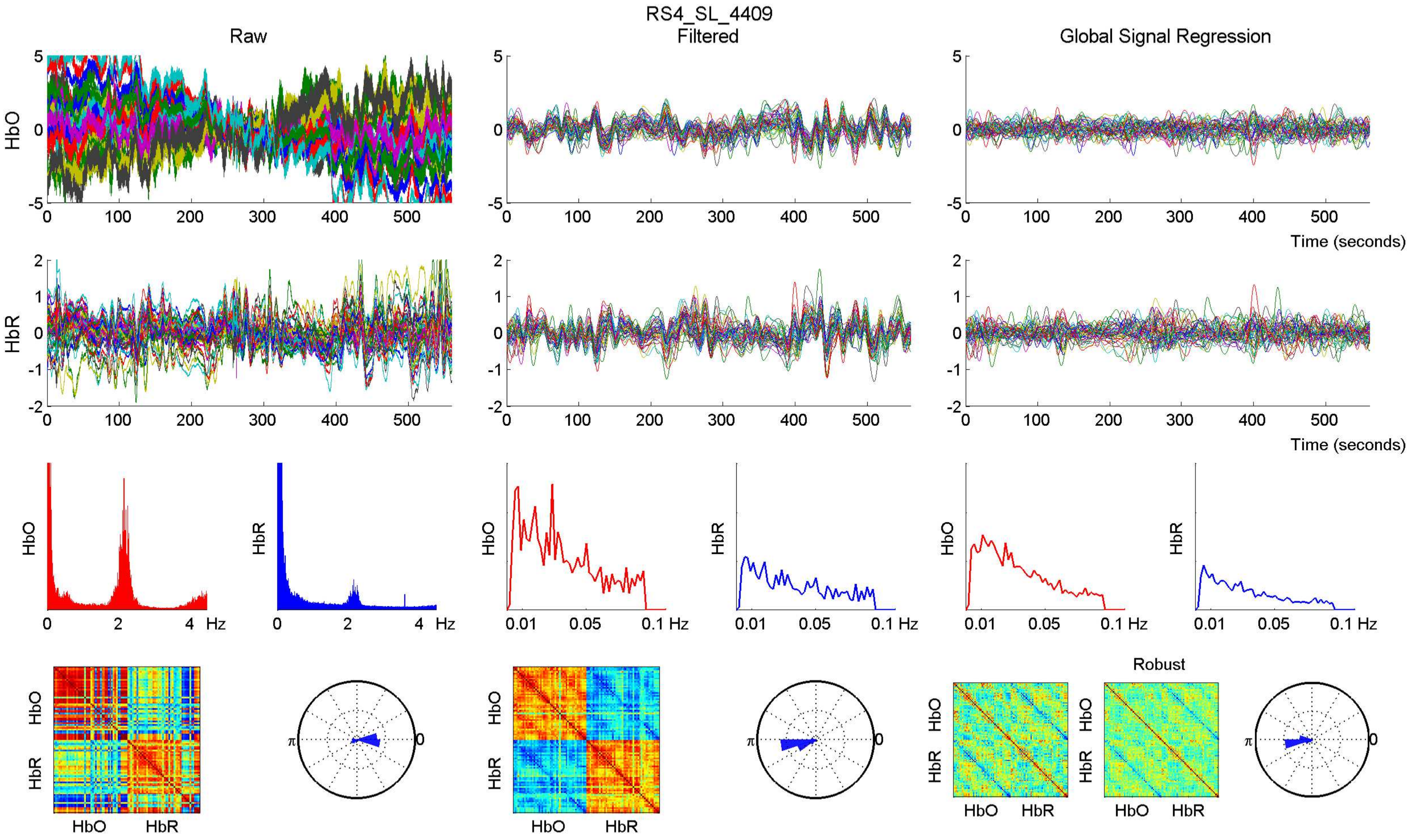

RS4_SL_4409
Raw
Filtered
Global Signal Regression
HbO
HbR
Time (seconds)
Hz
0.01
0.05
0.1 Hz
Robust
π
0

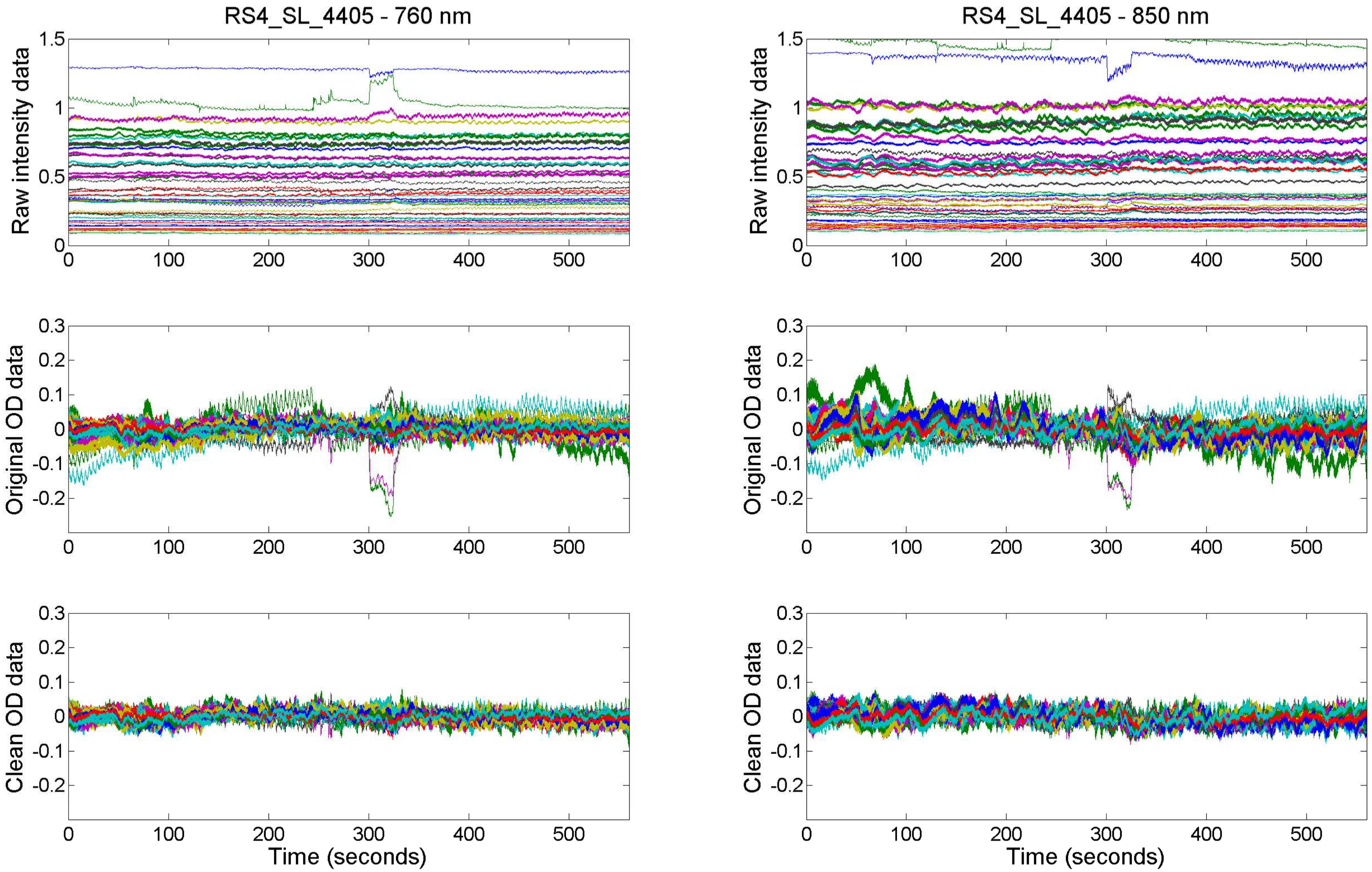
RS4_SL_4405 - 760 nm
RS4_SL_4405 - 850 nm
Raw intensity data
Original OD data
Clean OD data
Time (seconds)

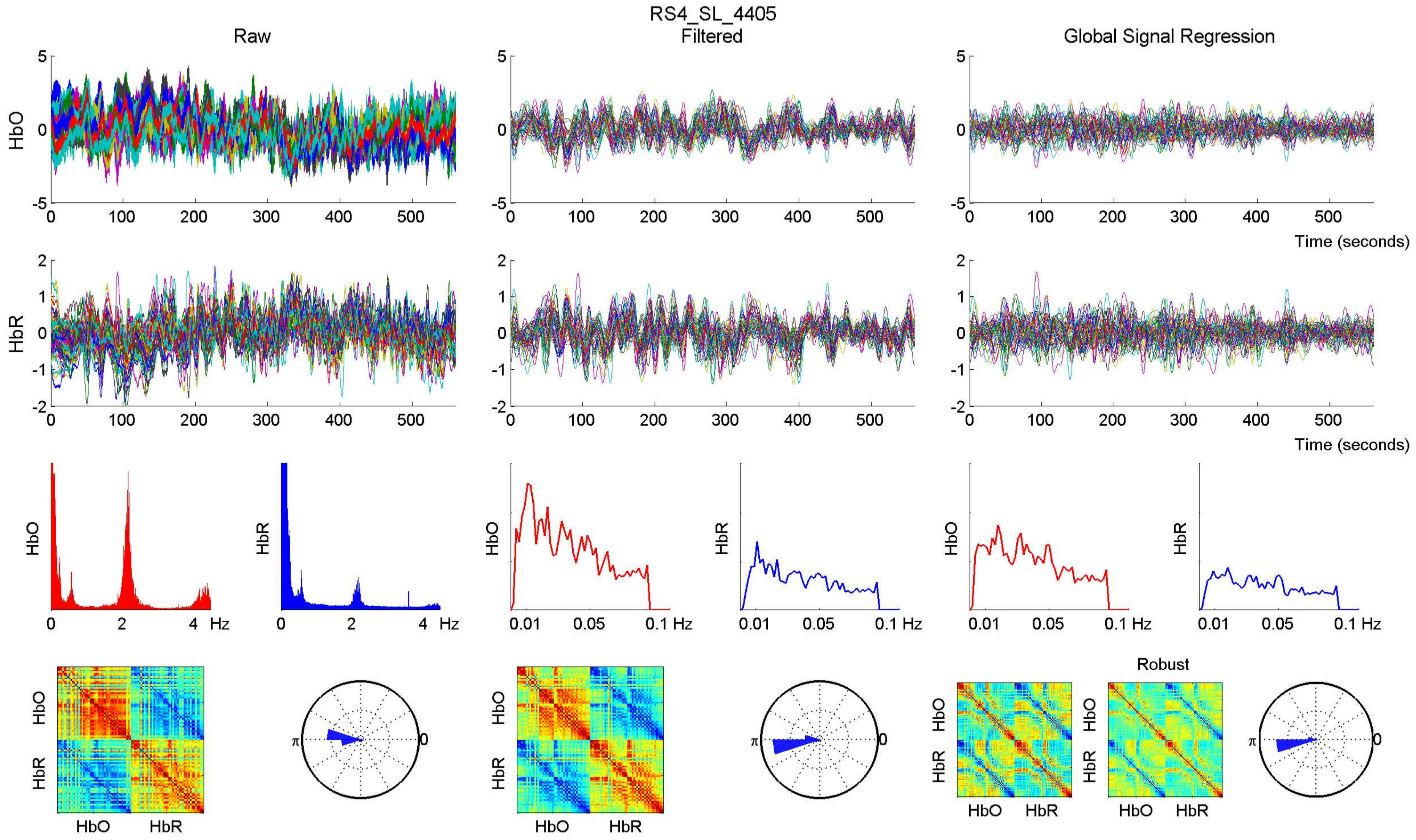

RS4_SL_4405
Raw
Filtered
Global Signal Regression
HbO
HbR
Time (seconds)
Hz
Robust

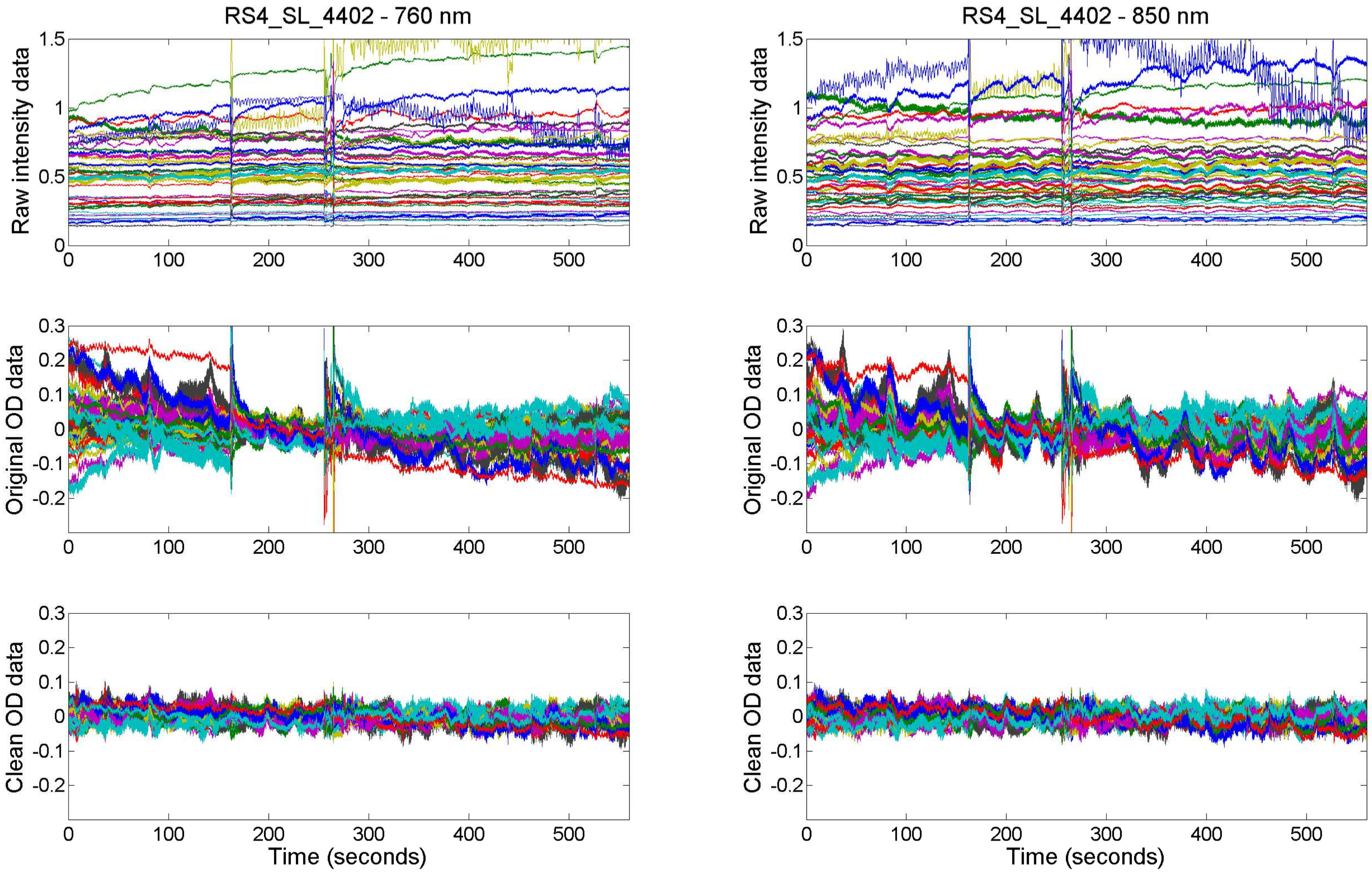
RS4_SL_4402 - 760 nm
RS4_SL_4402 - 850 nm
Raw intensity data
Original OD data
Clean OD data
Time (seconds)
0
100
200
300
400
500
1.5
1
0.5
0.3
0.2
0.1
-0.1
-0.2

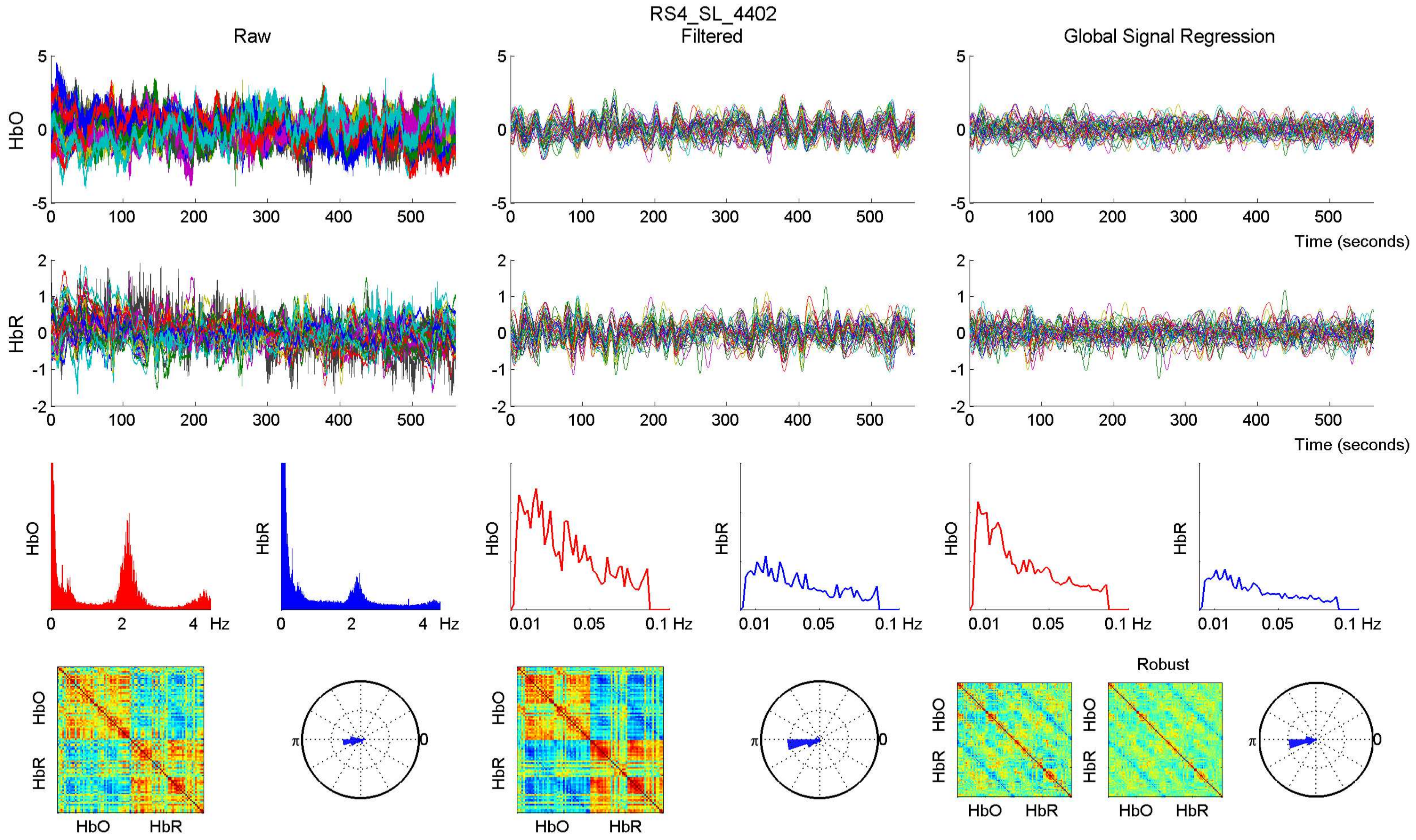

RS4_SL_4402
Raw
Filtered
Global Signal Regression
HbO
HbR
Time (seconds)
Hz
0.01
0.05
0.1 Hz
Robust
π
0

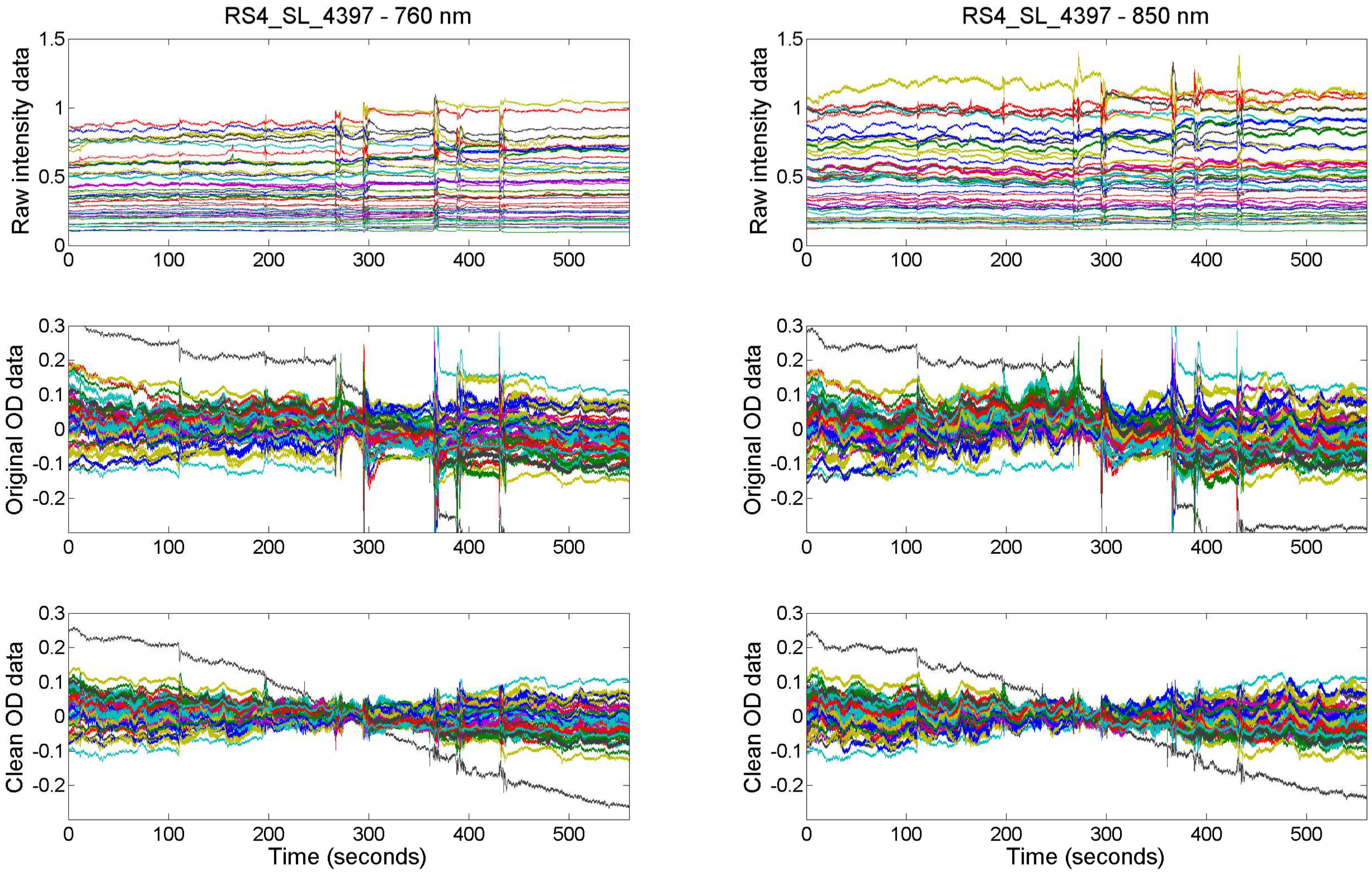
RS4_SL_4397 - 760 nm
RS4_SL_4397 - 850 nm
Raw intensity data
Original OD data
Clean OD data
Time (seconds)

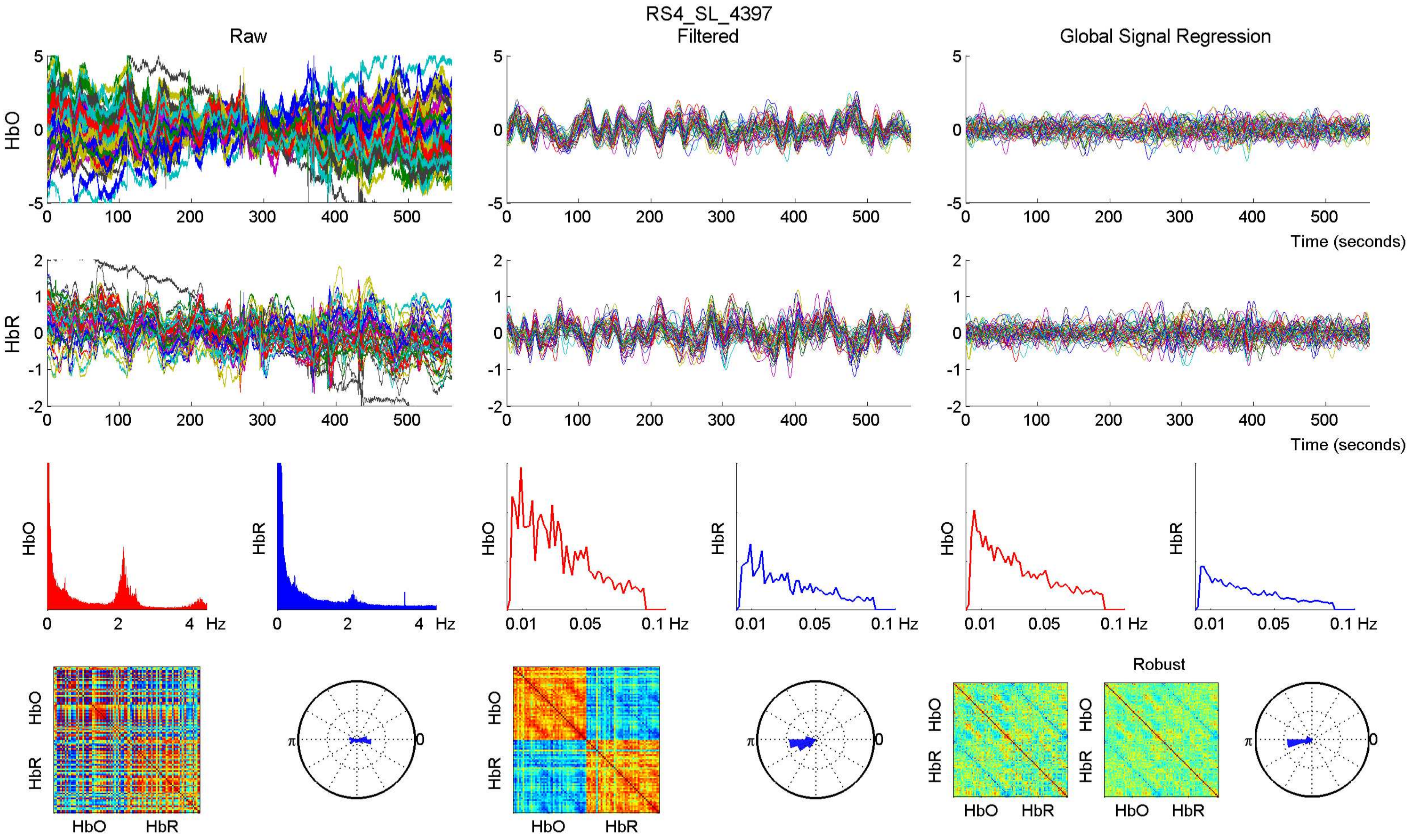

RS4_SL_4397
Raw
Filtered
Global Signal Regression
HbO
HbR
Time (seconds)
Hz
Robust

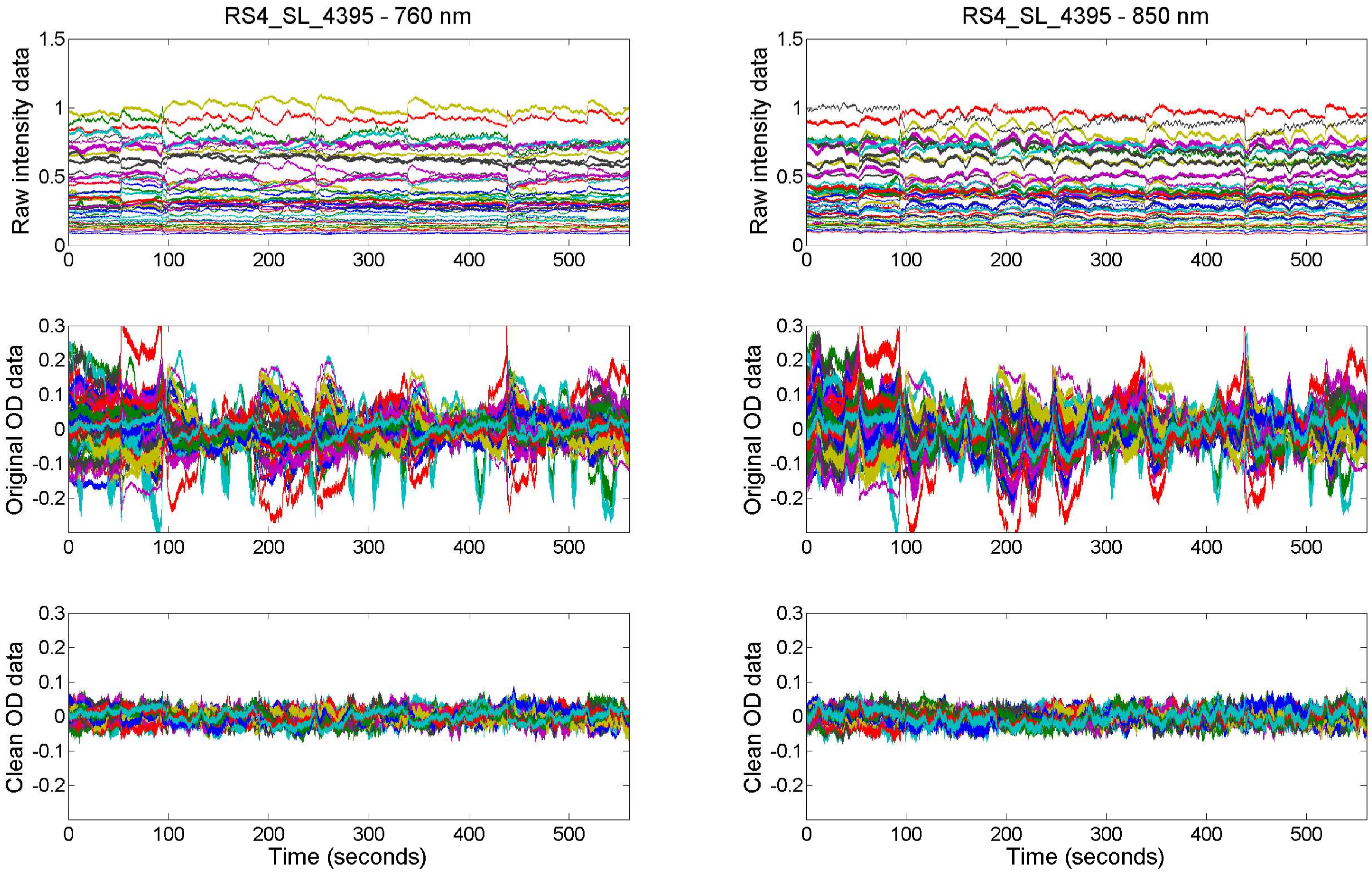

RS4_SL_4395 - 760 nm
RS4_SL_4395 - 850 nm
Raw intensity data
Original OD data
Clean OD data
Time (seconds)

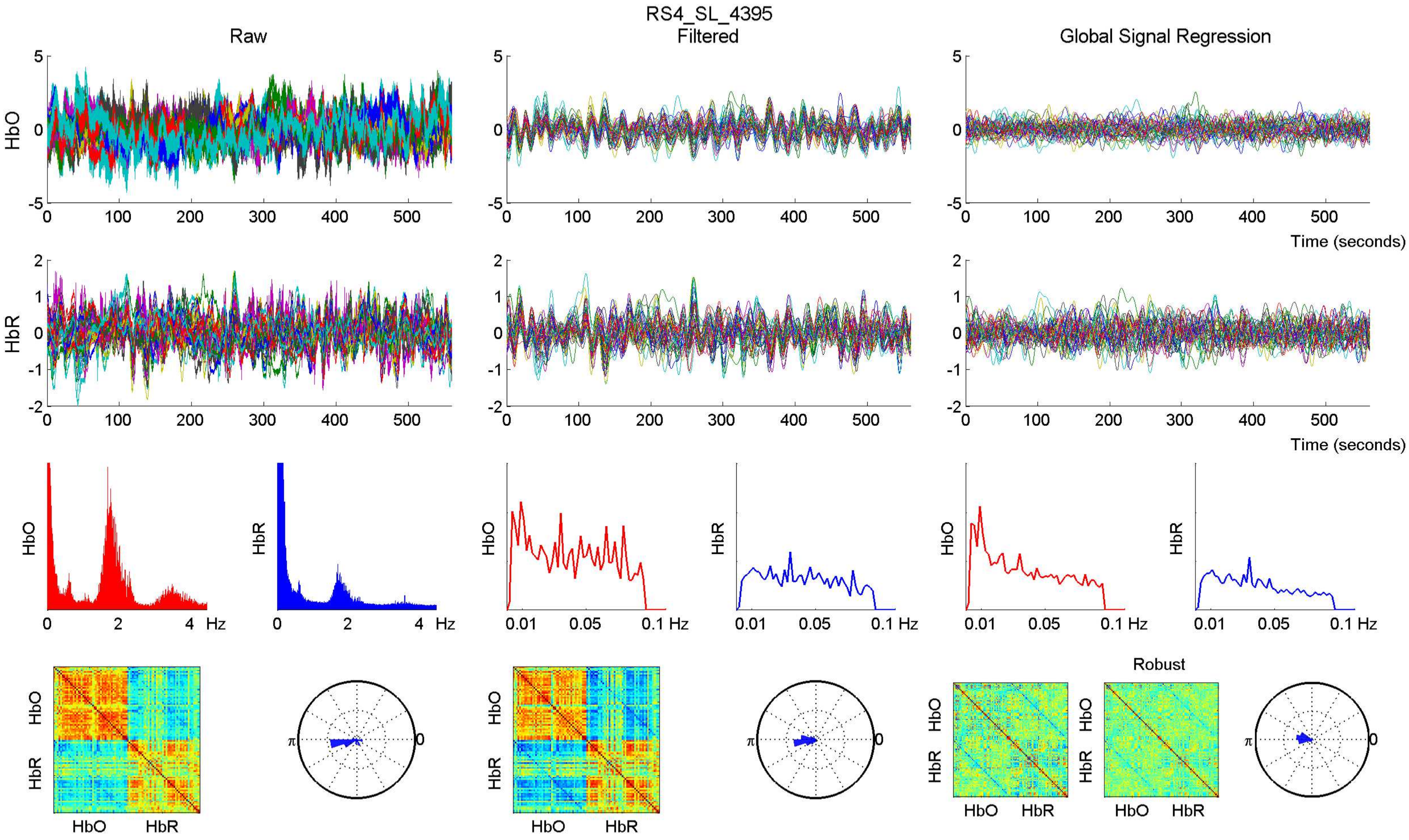
RS4_SL_4395
Raw
Filtered
Global Signal Regression
HbO
HbR
Time (seconds)
Hz
Robust

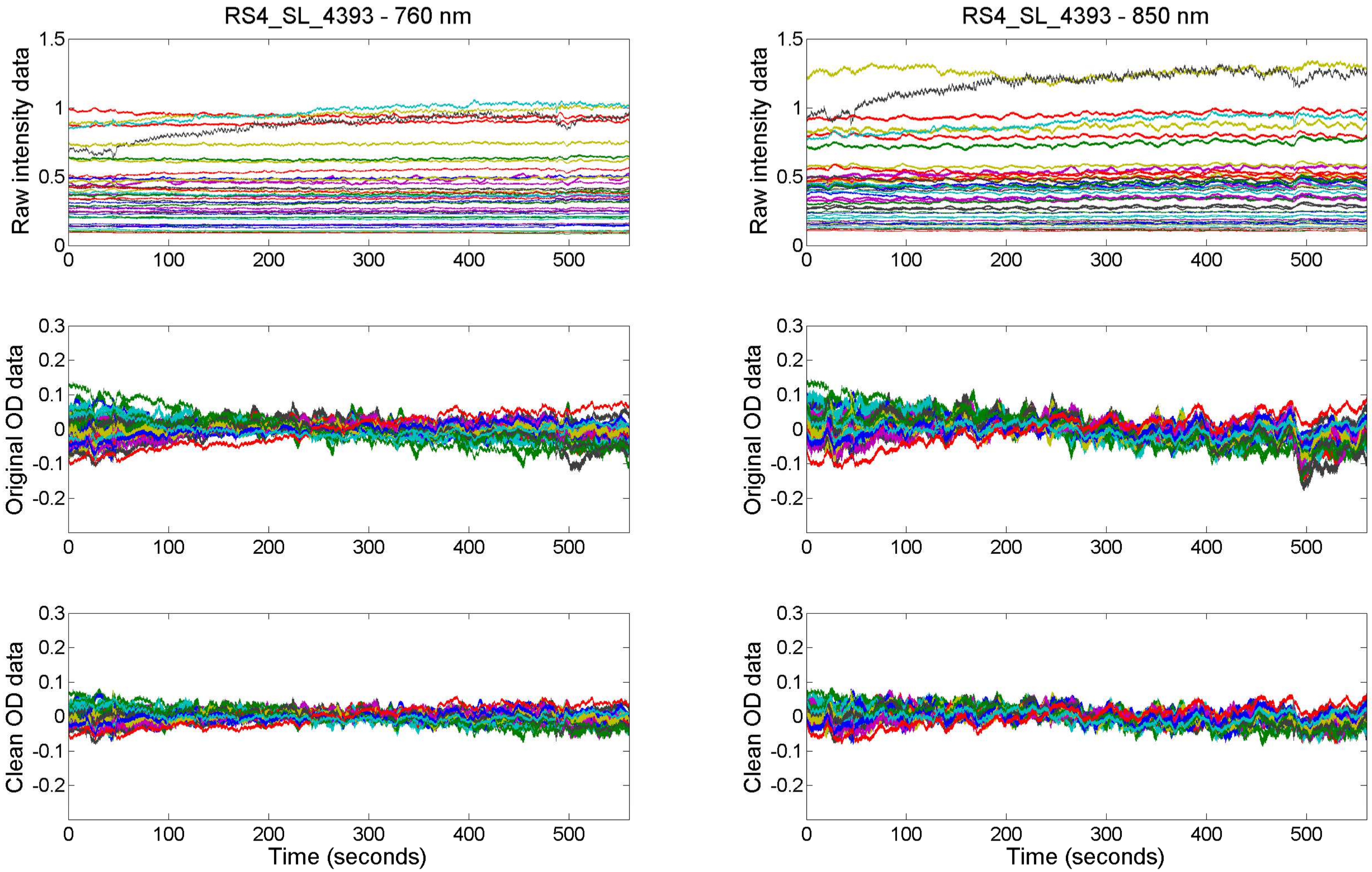

RS4_SL_4393 - 760 nm
RS4_SL_4393 - 850 nm
Raw intensity data
Original OD data
Clean OD data
Time (seconds)

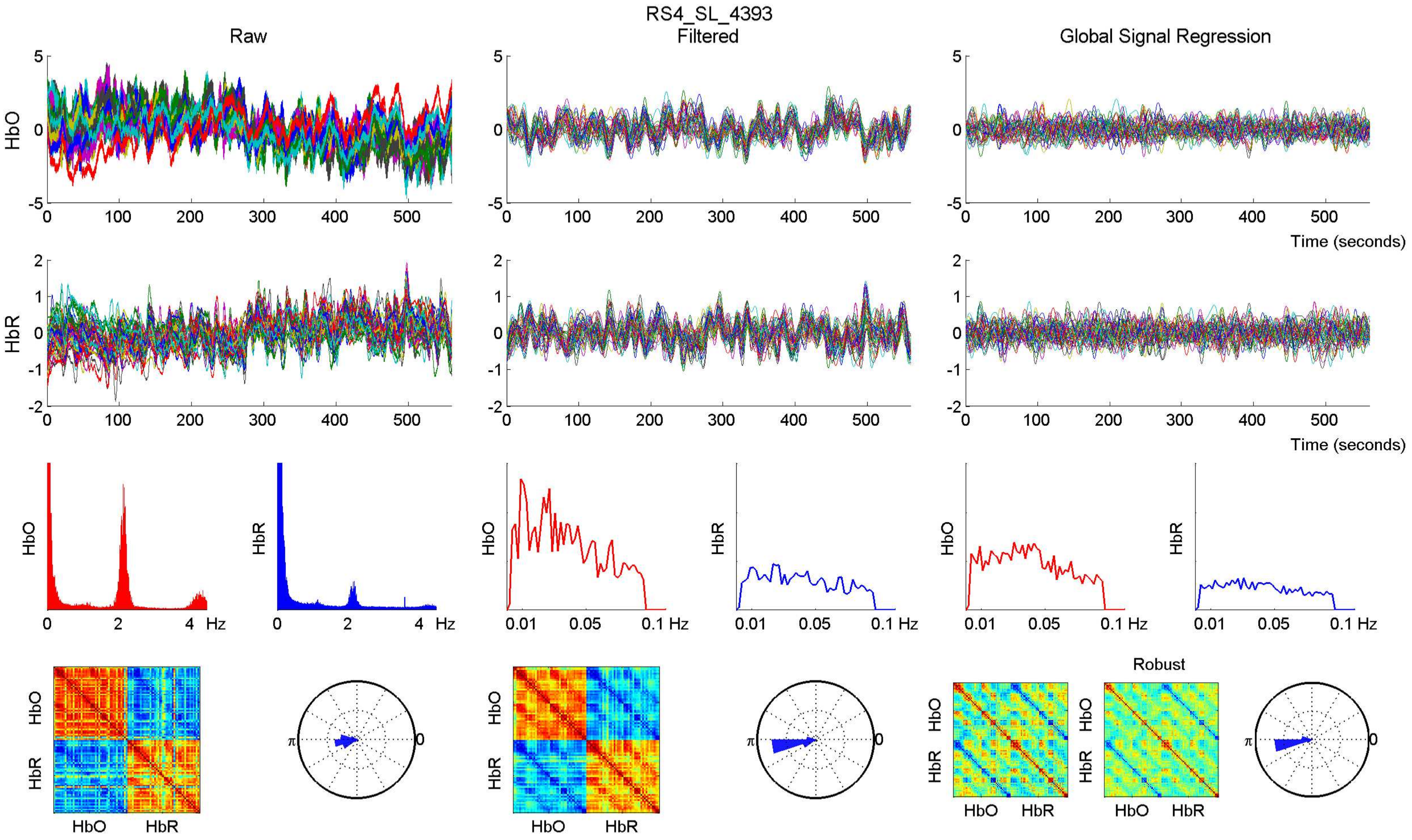

RS4_SL_4393
Raw
Filtered
Global Signal Regression
HbO
HbR
Time (seconds)
Hz
Robust

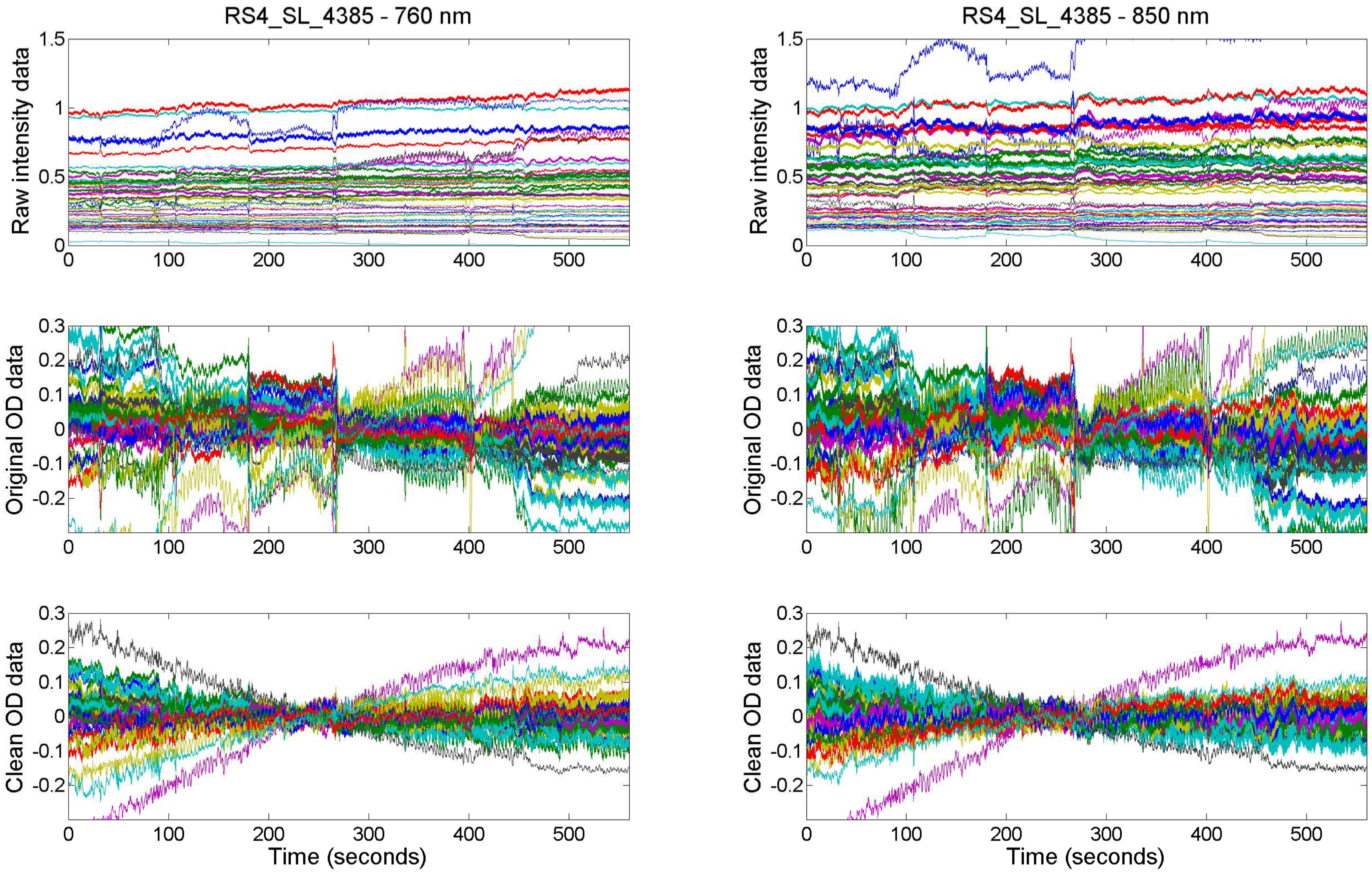

RS4_SL_4385 - 760 nm
RS4_SL_4385 - 850 nm
Raw intensity data
Original OD data
Clean OD data
Time (seconds)

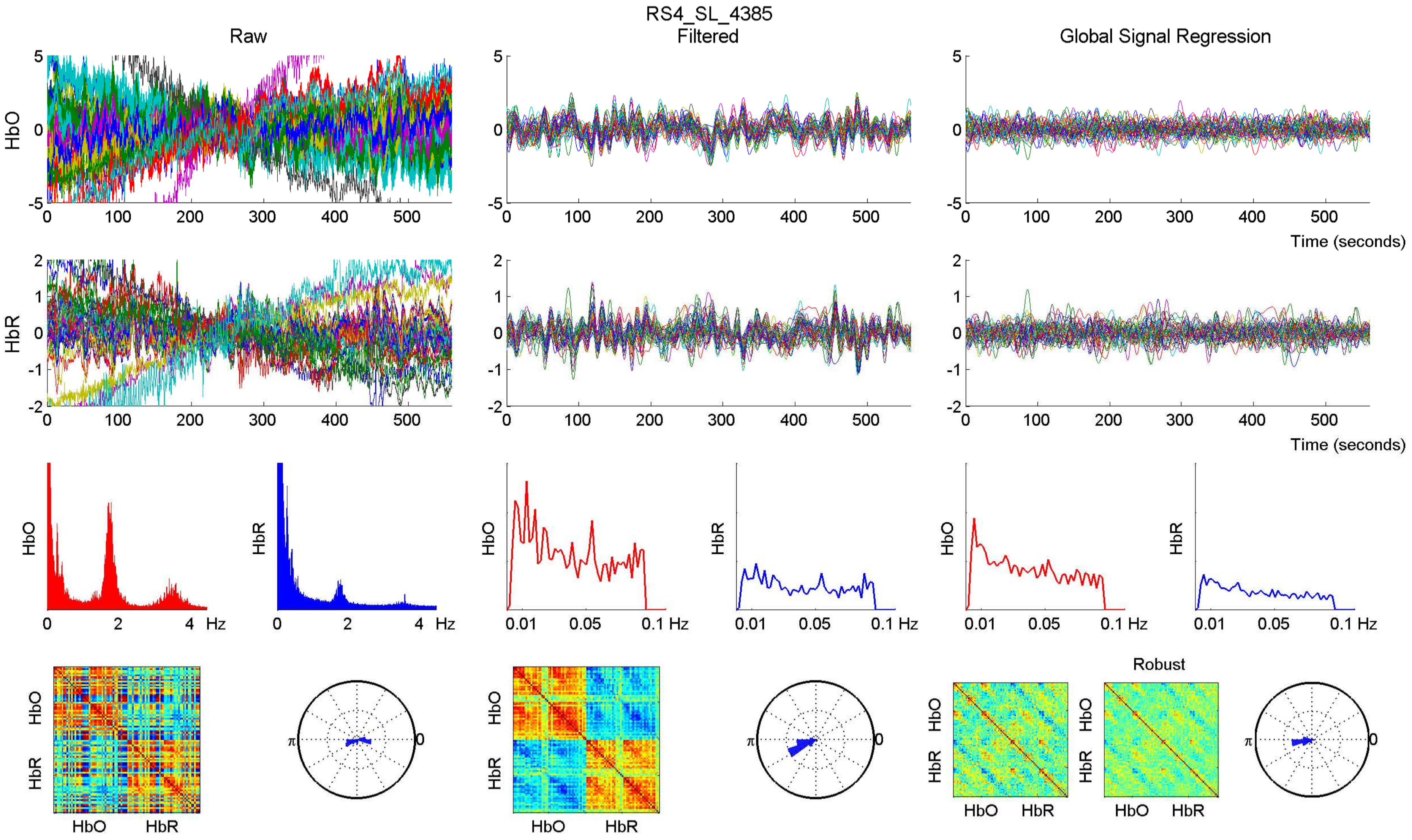
RS4_SL_4385
Raw
Filtered
Global Signal Regression
HbO
HbR
Time (seconds)
Hz
Robust

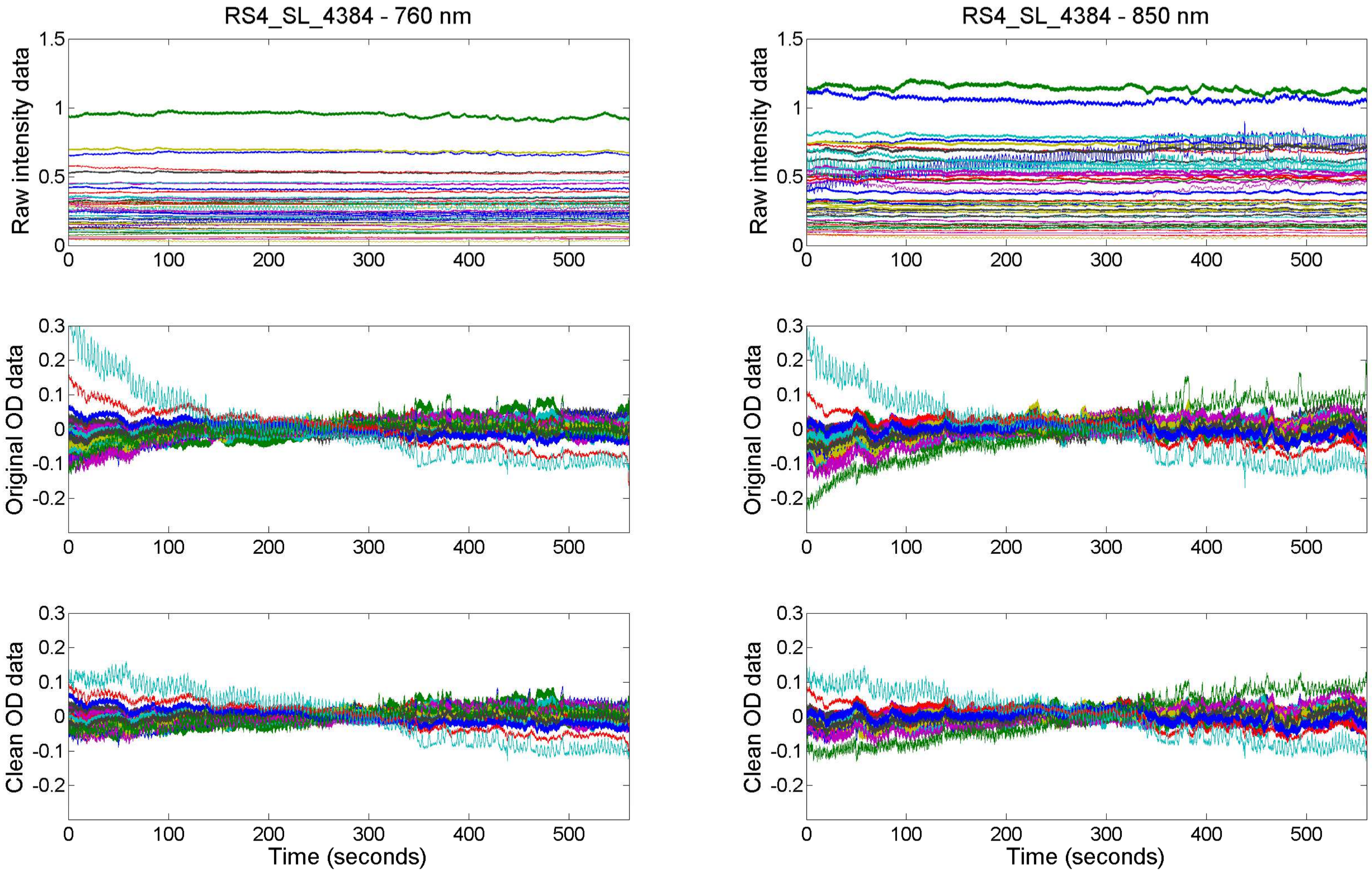
RS4_SL_4384 - 760 nm
RS4_SL_4384 - 850 nm
Raw intensity data
Original OD data
Clean OD data
Time (seconds)

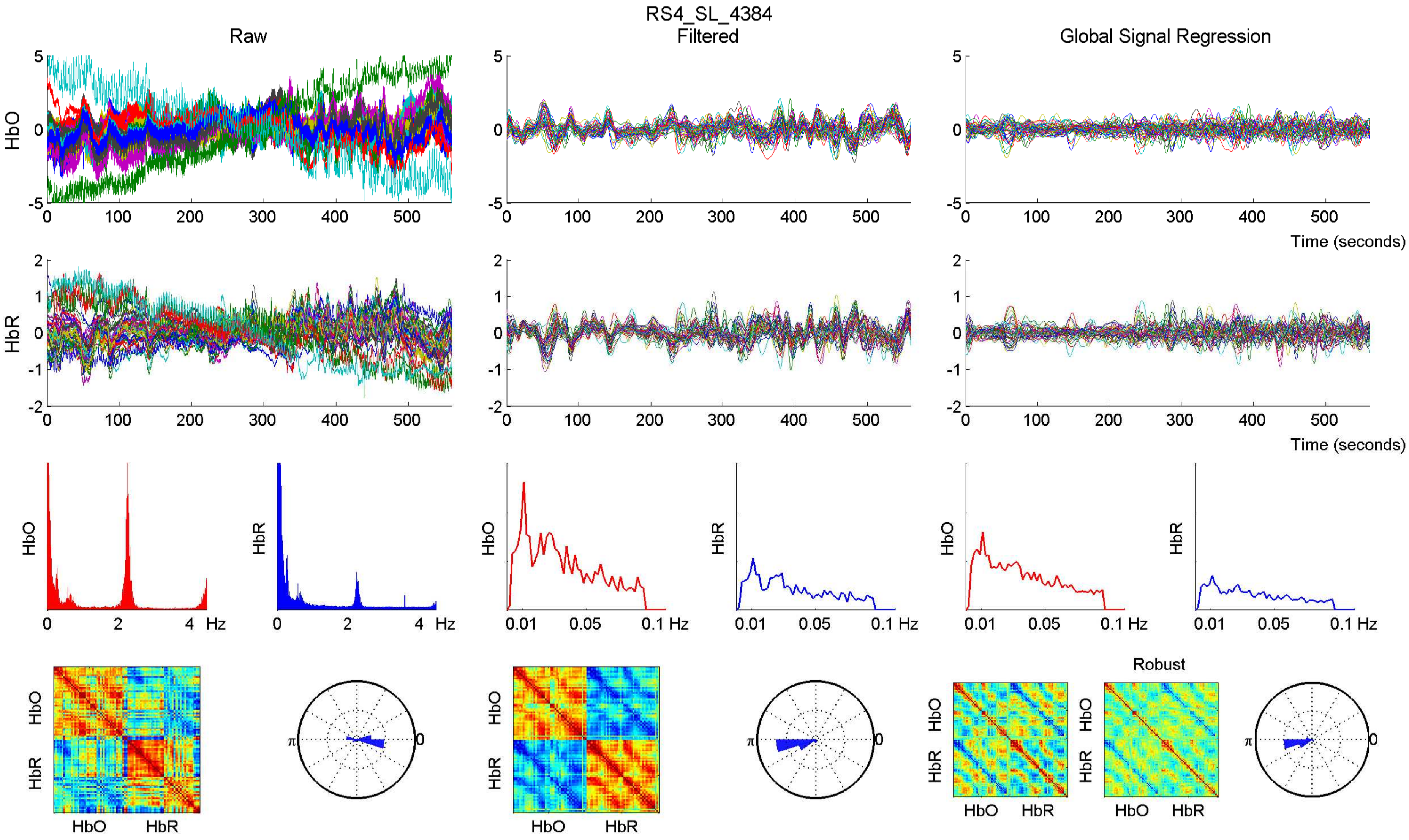

RS4_SL_4384
Raw
Filtered
Global Signal Regression
HbO
HbR
Time (seconds)
Hz
Robust
HbO HbR

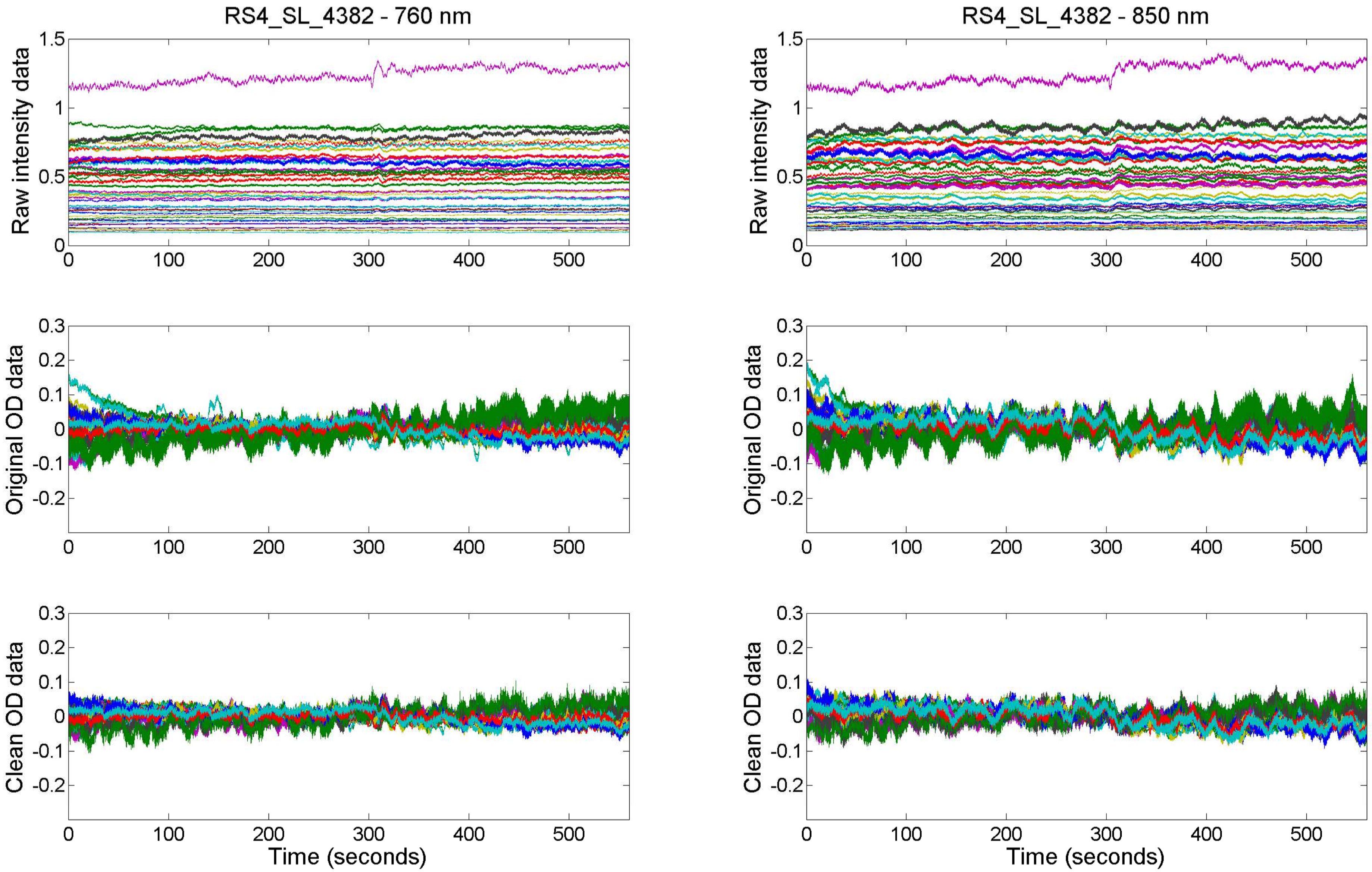

RS4_SL_4382 - 760 nm
RS4_SL_4382 - 850 nm
Raw intensity data
Original OD data
Clean OD data
Time (seconds)

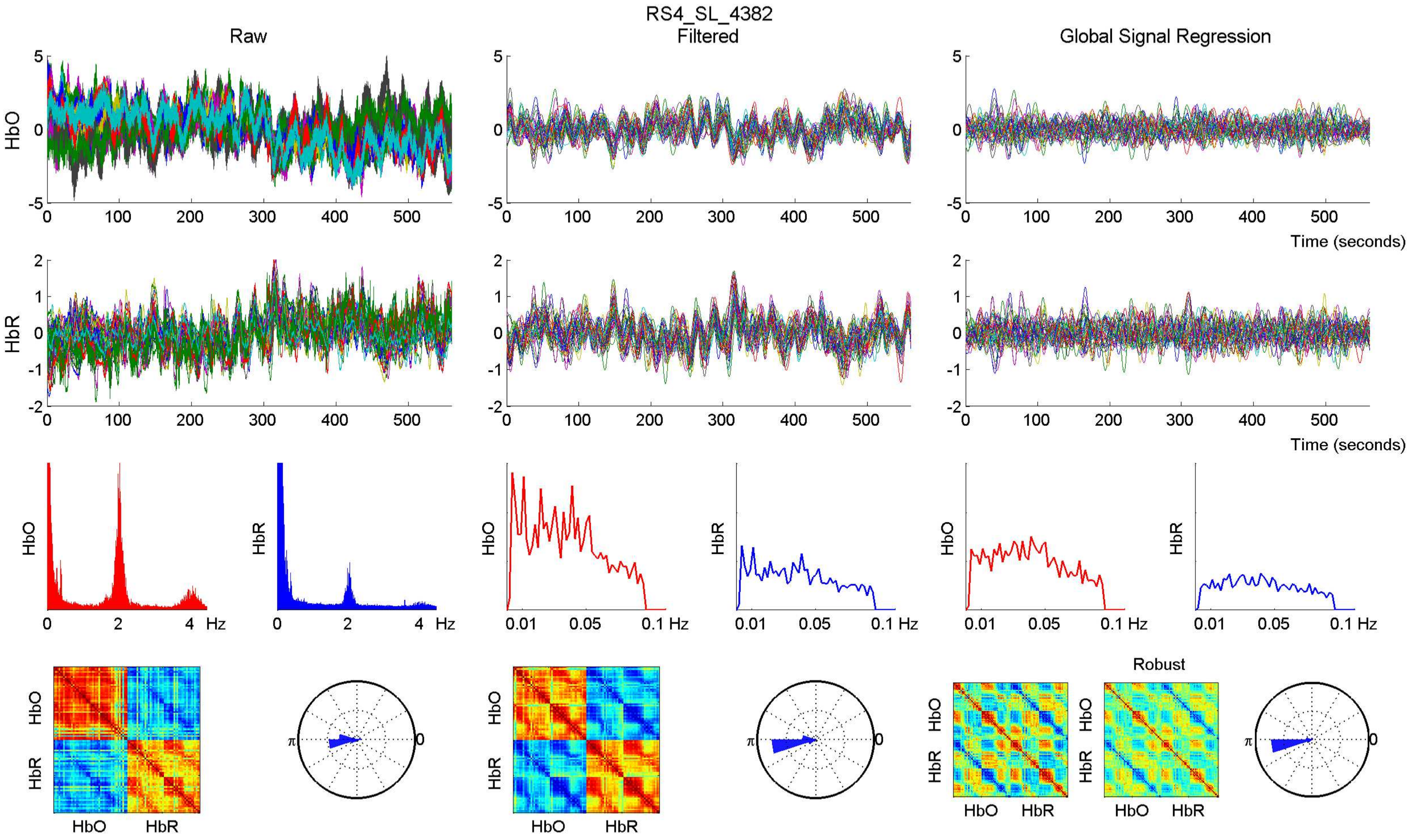

RS4_SL_4382
Raw
Filtered
Global Signal Regression
HbO
HbR
Time (seconds)
Hz
0.01 0.05 0.1 Hz
Robust

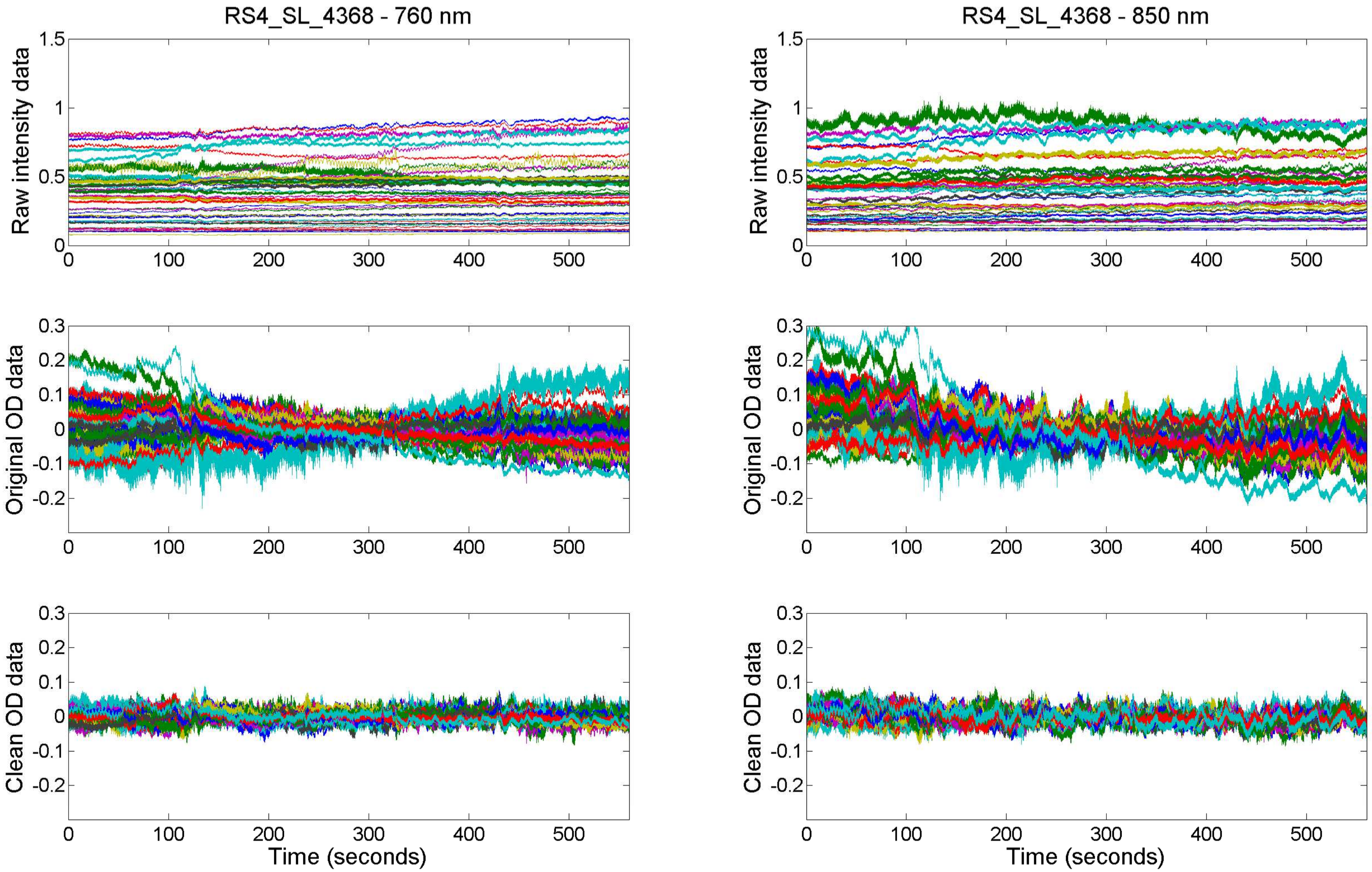
RS4_SL_4368 - 760 nm
RS4_SL_4368 - 850 nm
Raw intensity data
Original OD data
Clean OD data
Time (seconds)
0
100
200
300
400
500
1.5
1
0.5
0.3
0.2
0.1
-0.1
-0.2

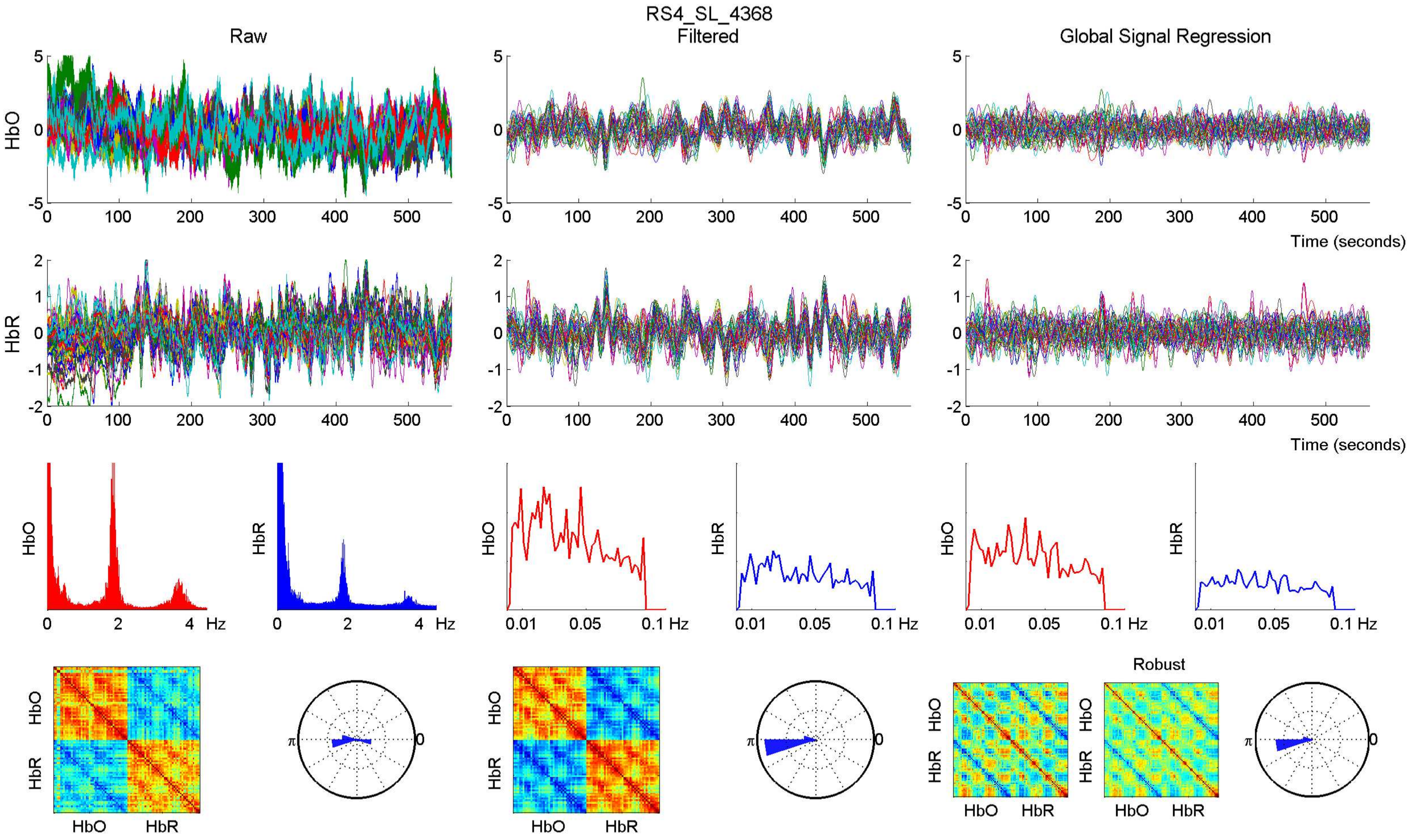

RS4_SL_4368
Raw
Filtered
Global Signal Regression
HbO
HbR
Time (seconds)
Hz
Robust
HbO
HbR

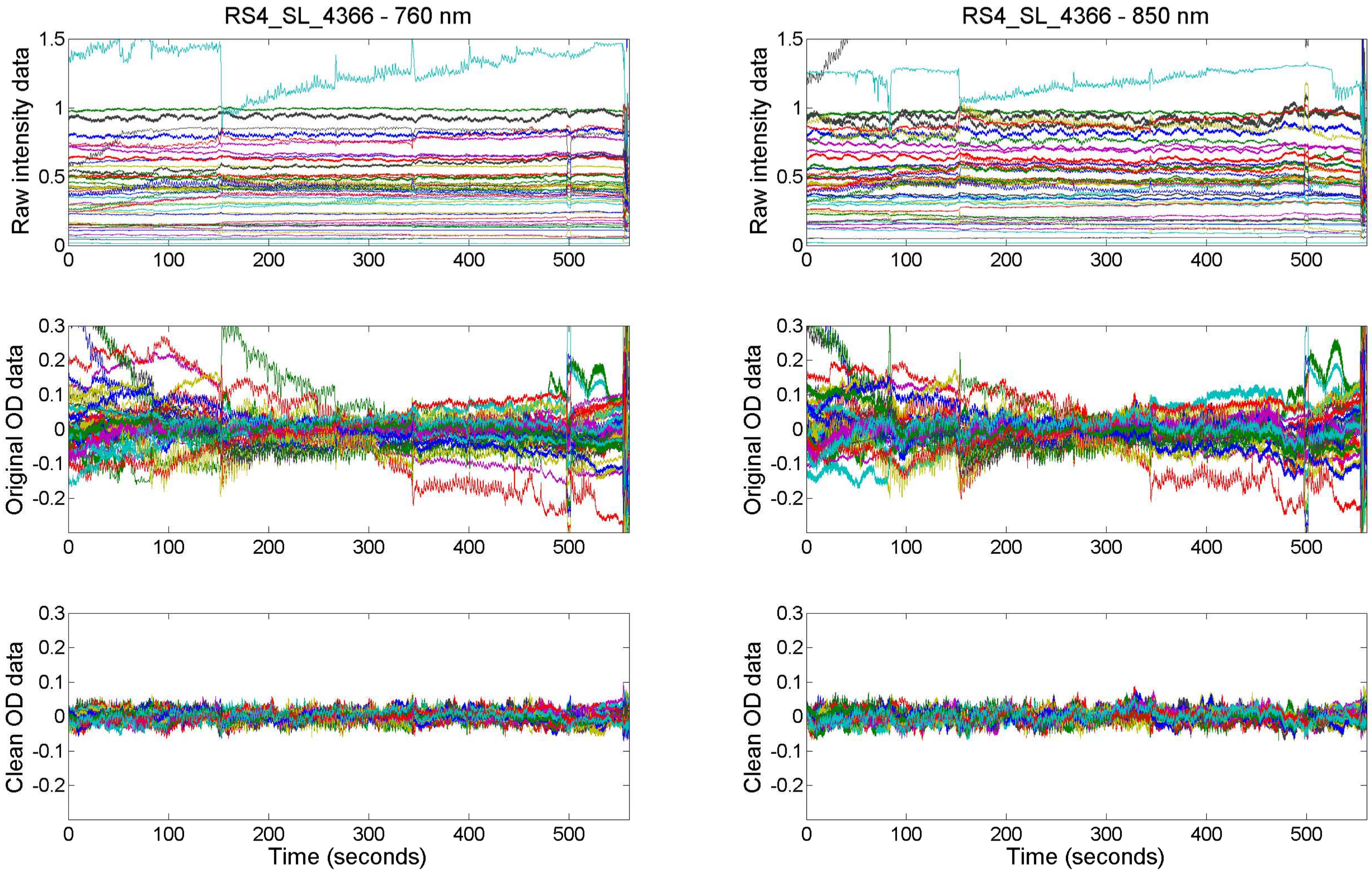

RS4_SL_4366 - 760 nm
RS4_SL_4366 - 850 nm
Raw intensity data
Original OD data
Clean OD data
Time (seconds)

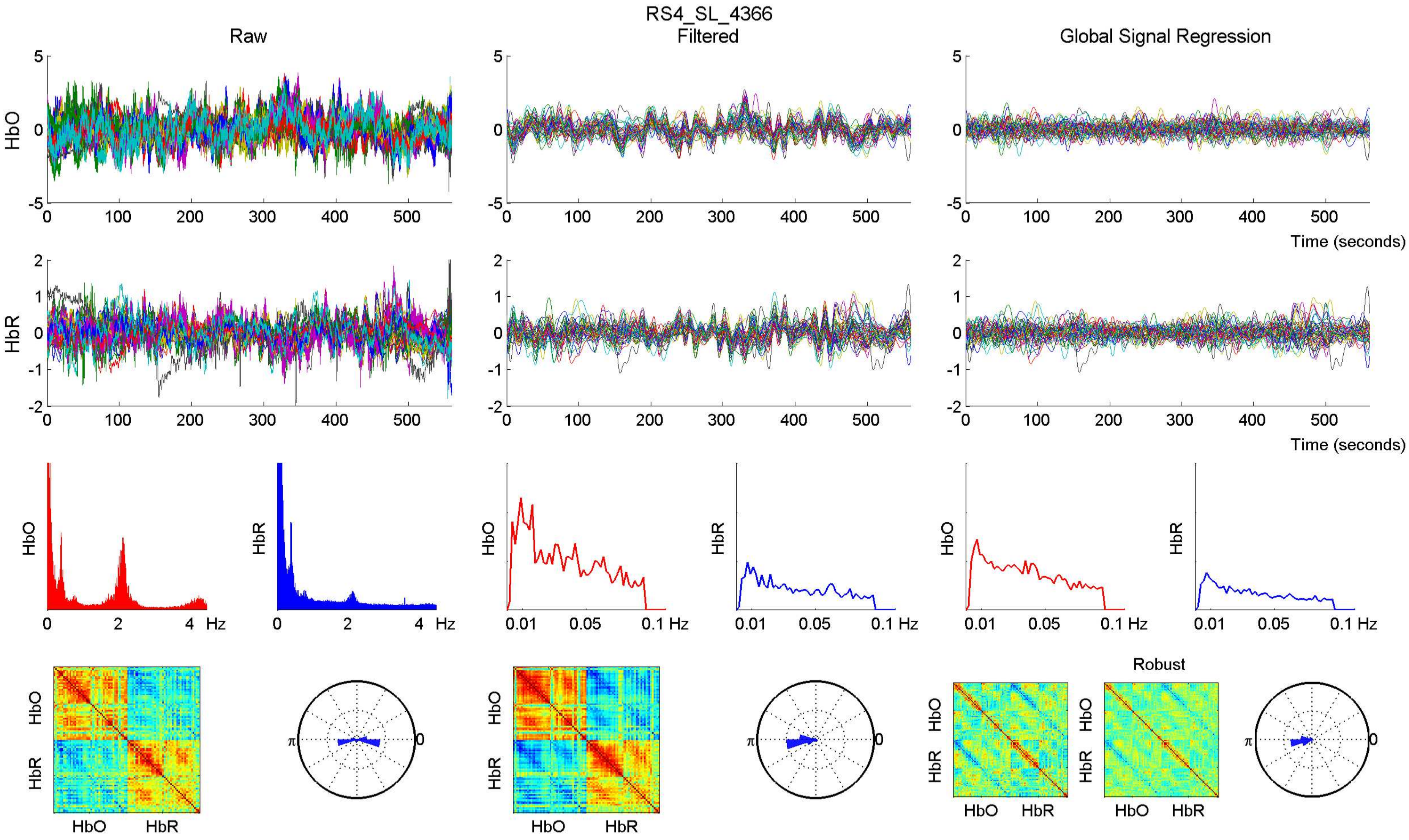
RS4_SL_4366
Raw
Filtered
Global Signal Regression
HbO
HbR
Time (seconds)
Hz
Robust

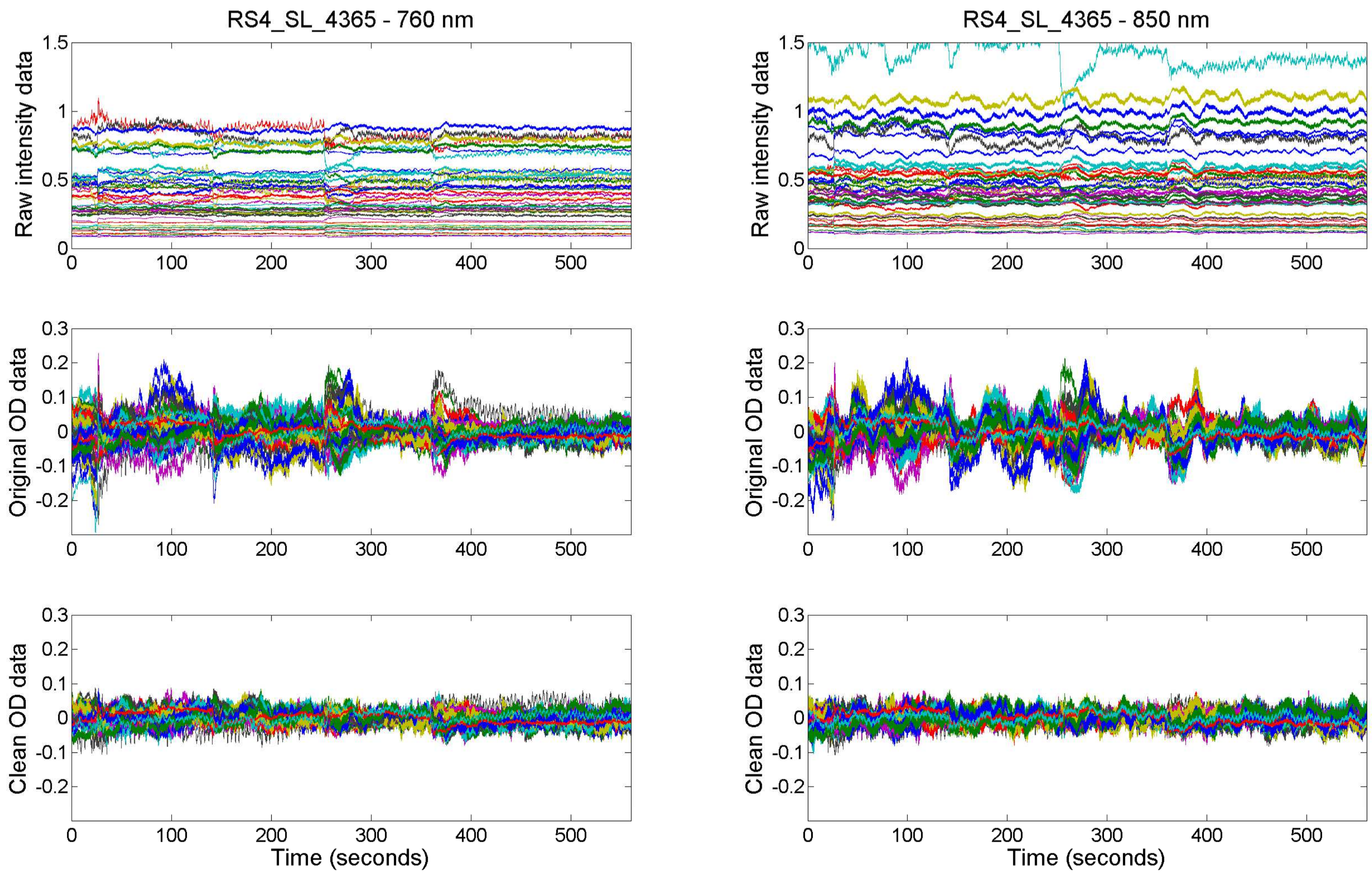

RS4_SL_4365 - 760 nm
RS4_SL_4365 - 850 nm
Raw intensity data
Original OD data
Clean OD data
Time (seconds)

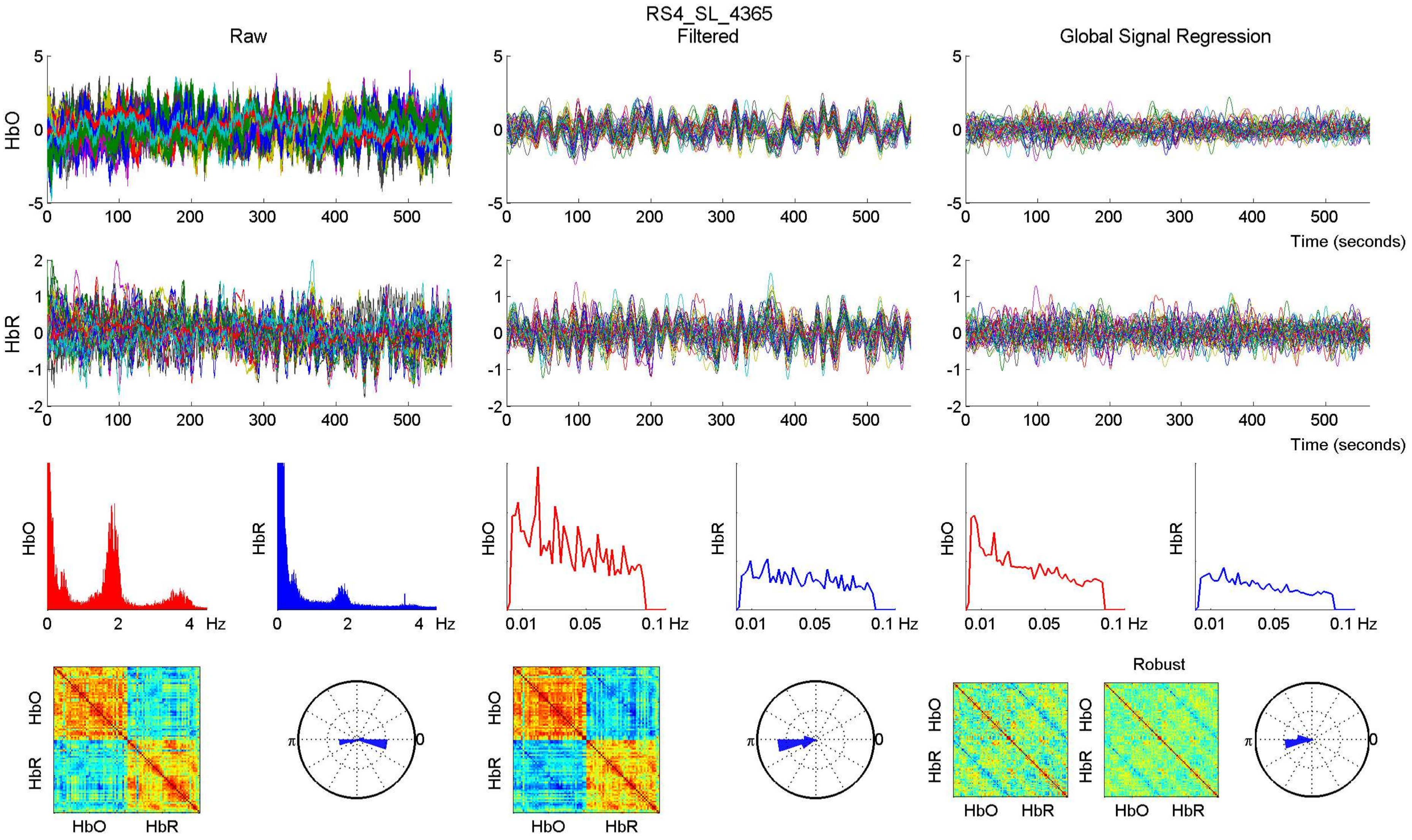

RS4_SL_4365
Raw
Filtered
Global Signal Regression
HbO
HbR
Time (seconds)
Hz
Robust

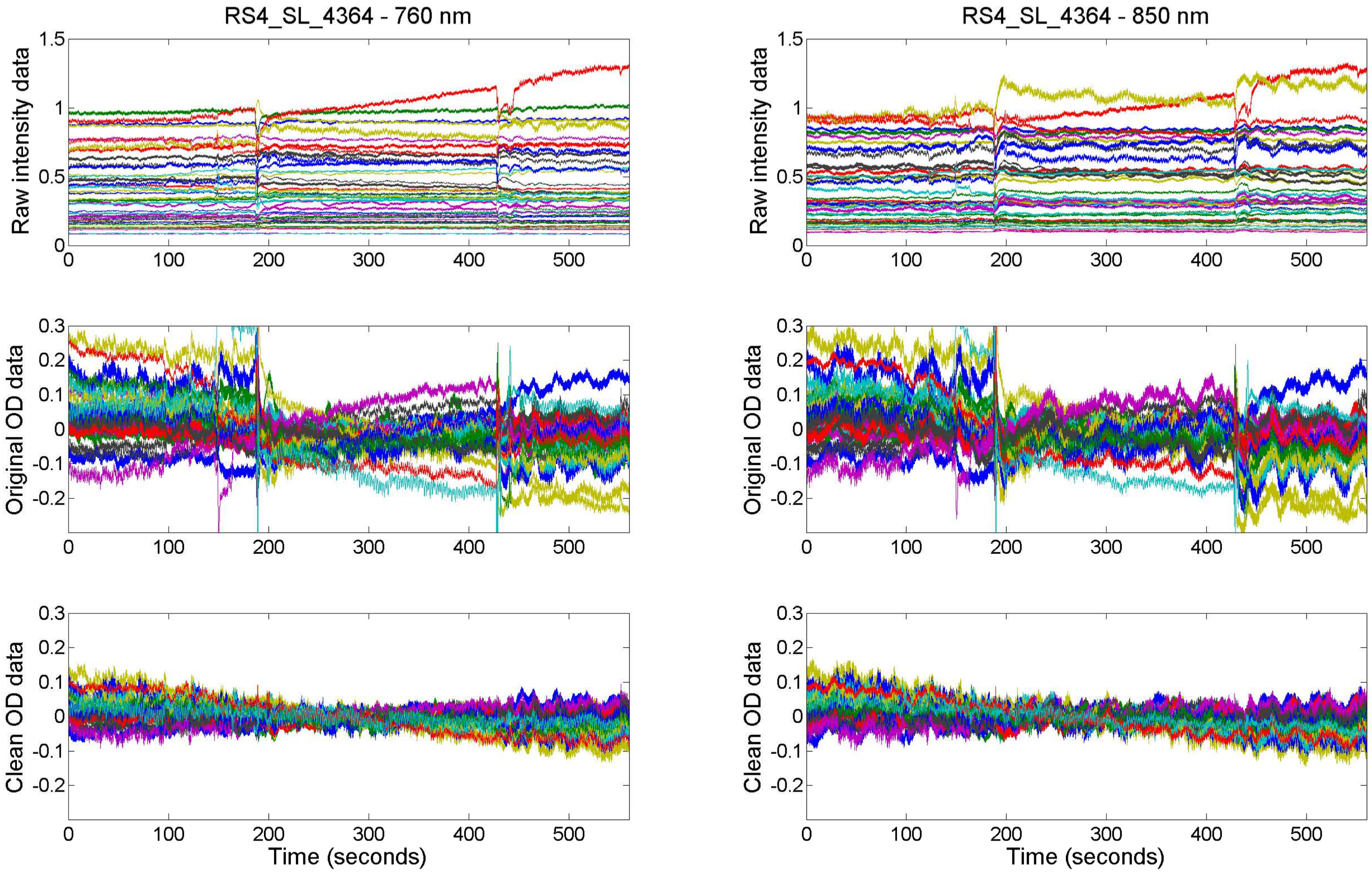

RS4_SL_4364 - 760 nm
RS4_SL_4364 - 850 nm
Raw intensity data
Original OD data
Clean OD data
Time (seconds)

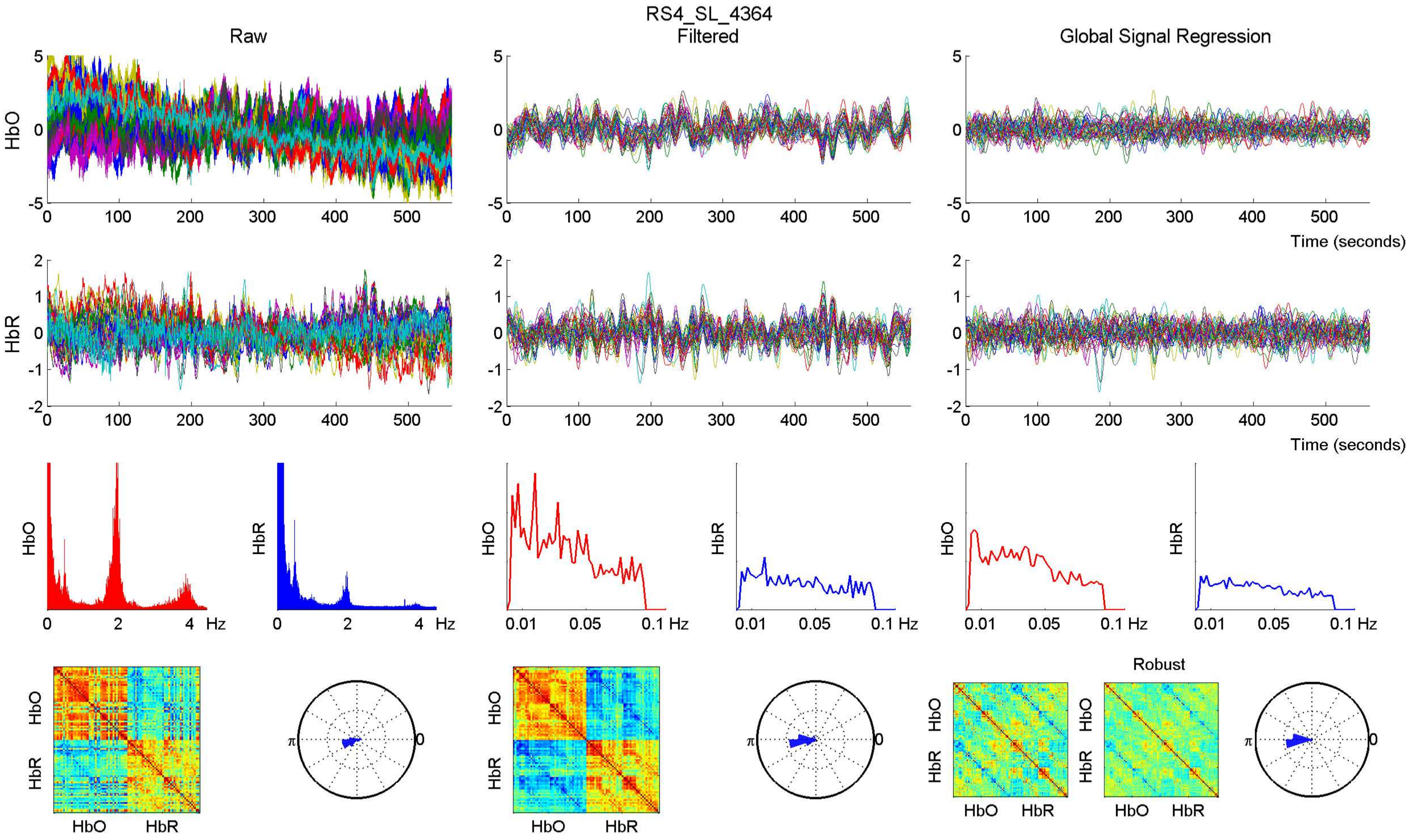

RS4_SL_4364
Raw
Filtered
Global Signal Regression
HbO
HbR
Time (seconds)
Hz
0.01
0.05
0.1 Hz
Robust
π
0

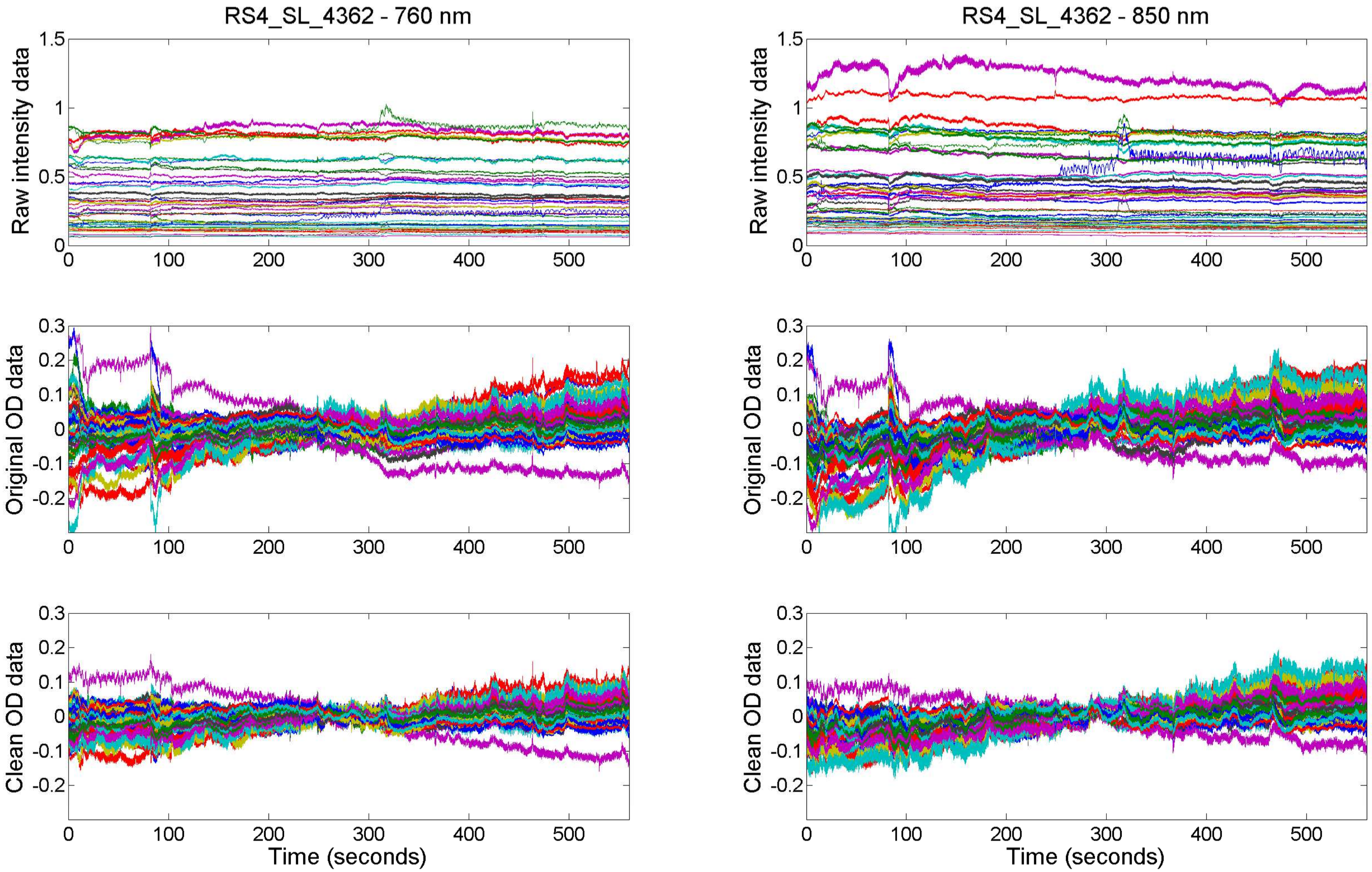
RS4_SL_4362 - 760 nm
RS4_SL_4362 - 850 nm
Raw intensity data
Original OD data
Clean OD data
Time (seconds)
0
100
200
300
400
500
1.5
1
0.5
0.3
0.2
0.1
-0.1
-0.2

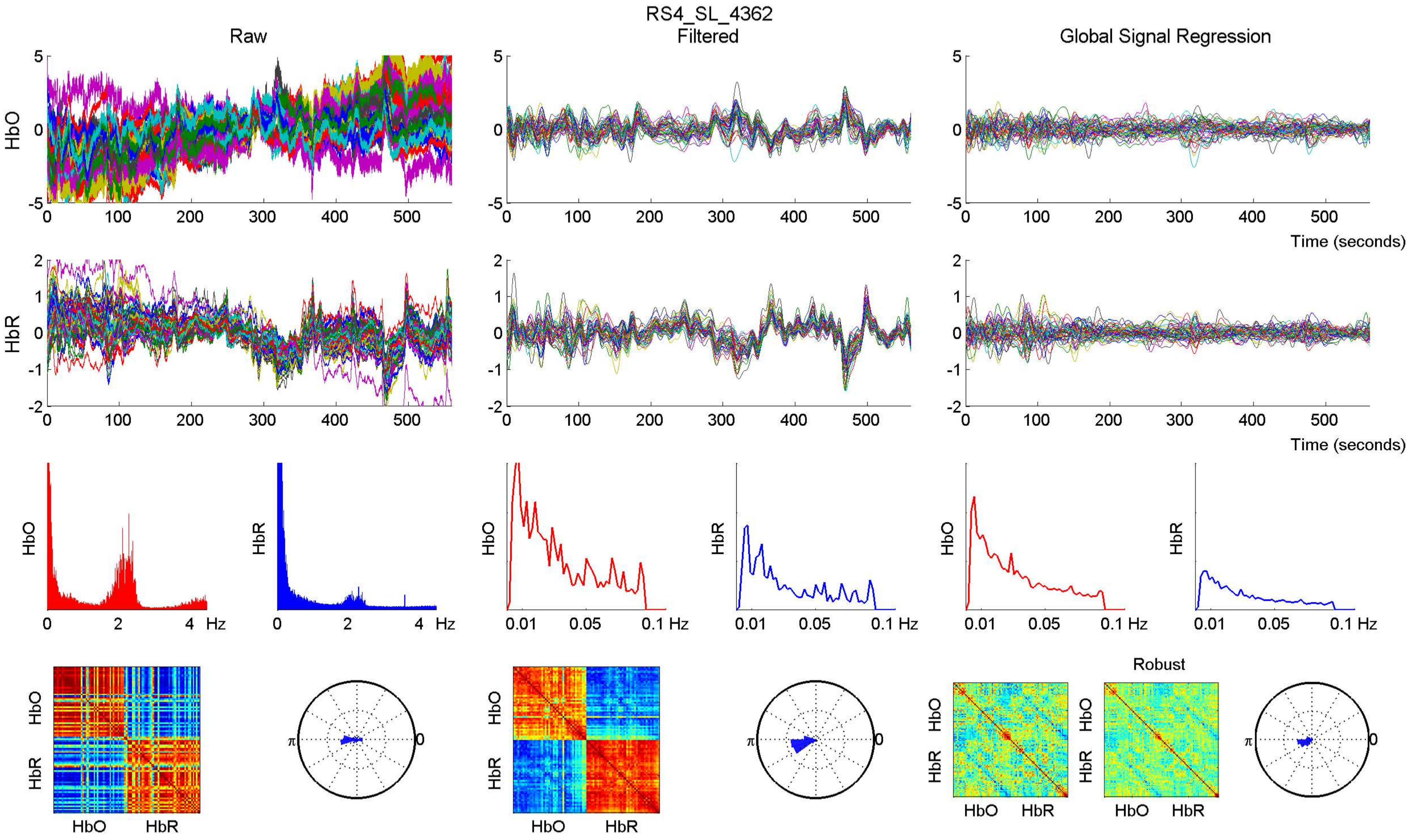

RS4_SL_4362
Raw
Filtered
Global Signal Regression
HbO
HbR
Time (seconds)
Hz
Robust
HbO
HbR

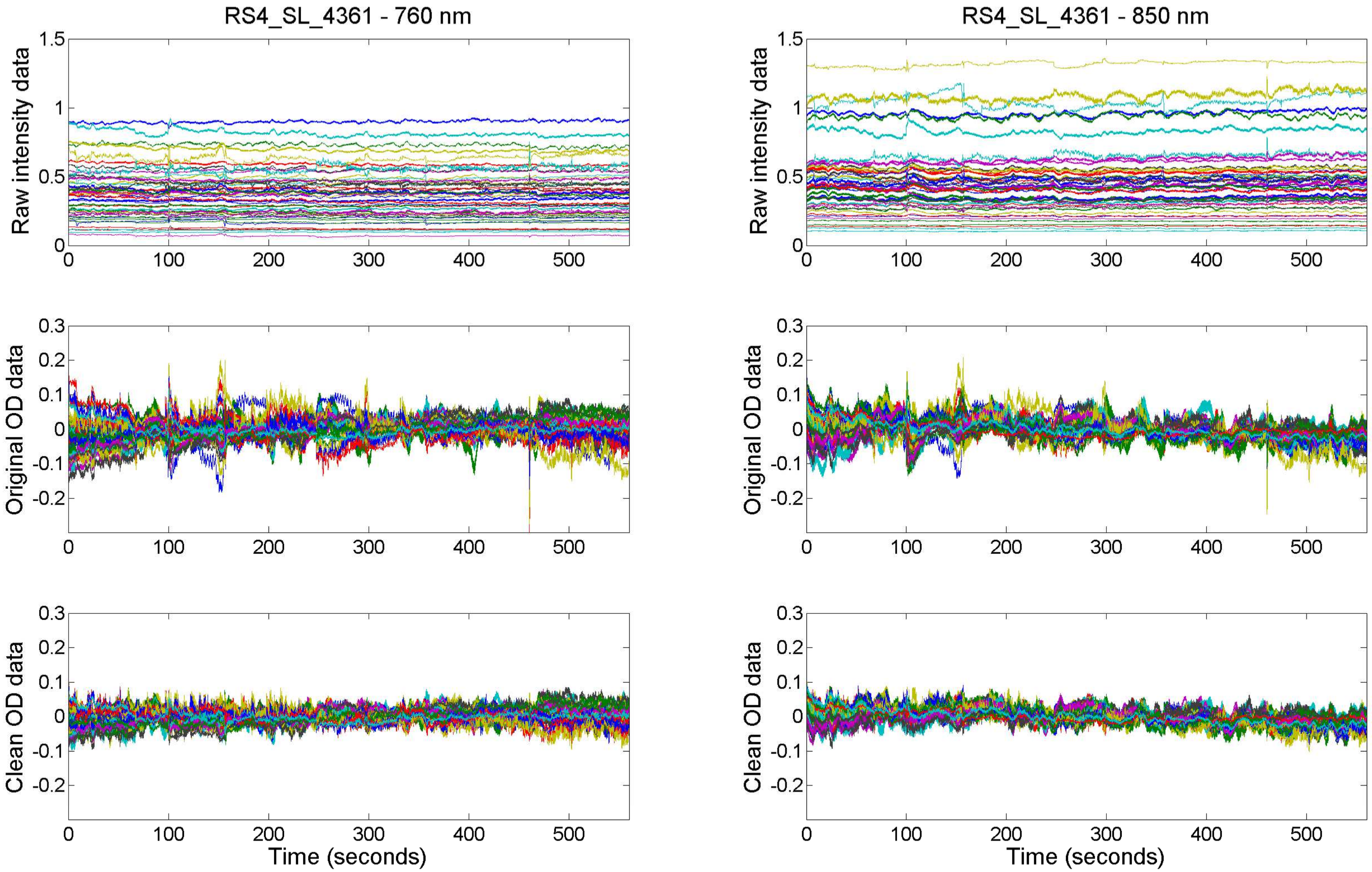

RS4_SL_4361 - 760 nm
RS4_SL_4361 - 850 nm
Raw intensity data
Original OD data
Clean OD data
Time (seconds)

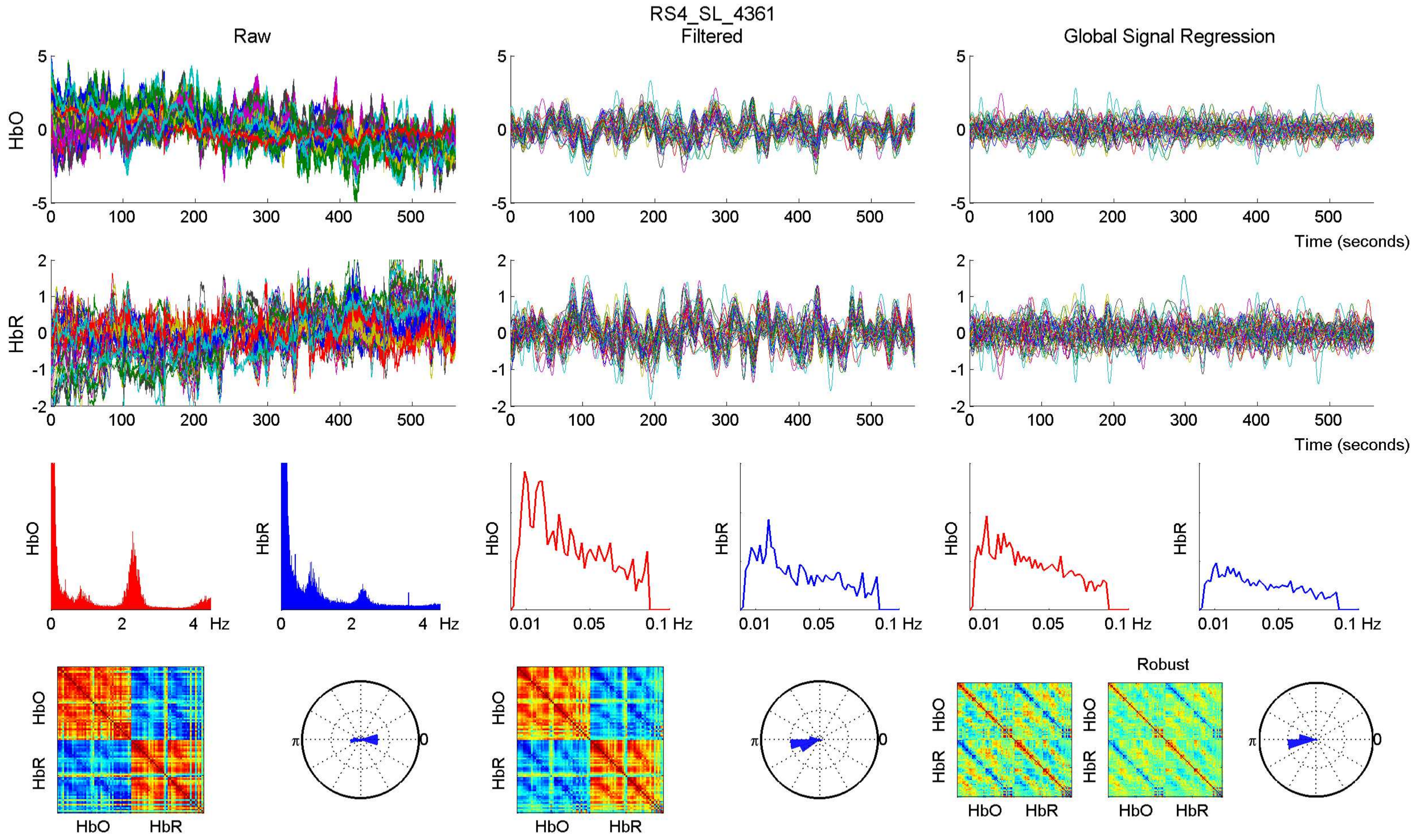

RS4_SL_4361
Raw
Filtered
Global Signal Regression
HbO
HbR
Time (seconds)
Hz
0.01
0.05
0.1 Hz
Robust
π
0

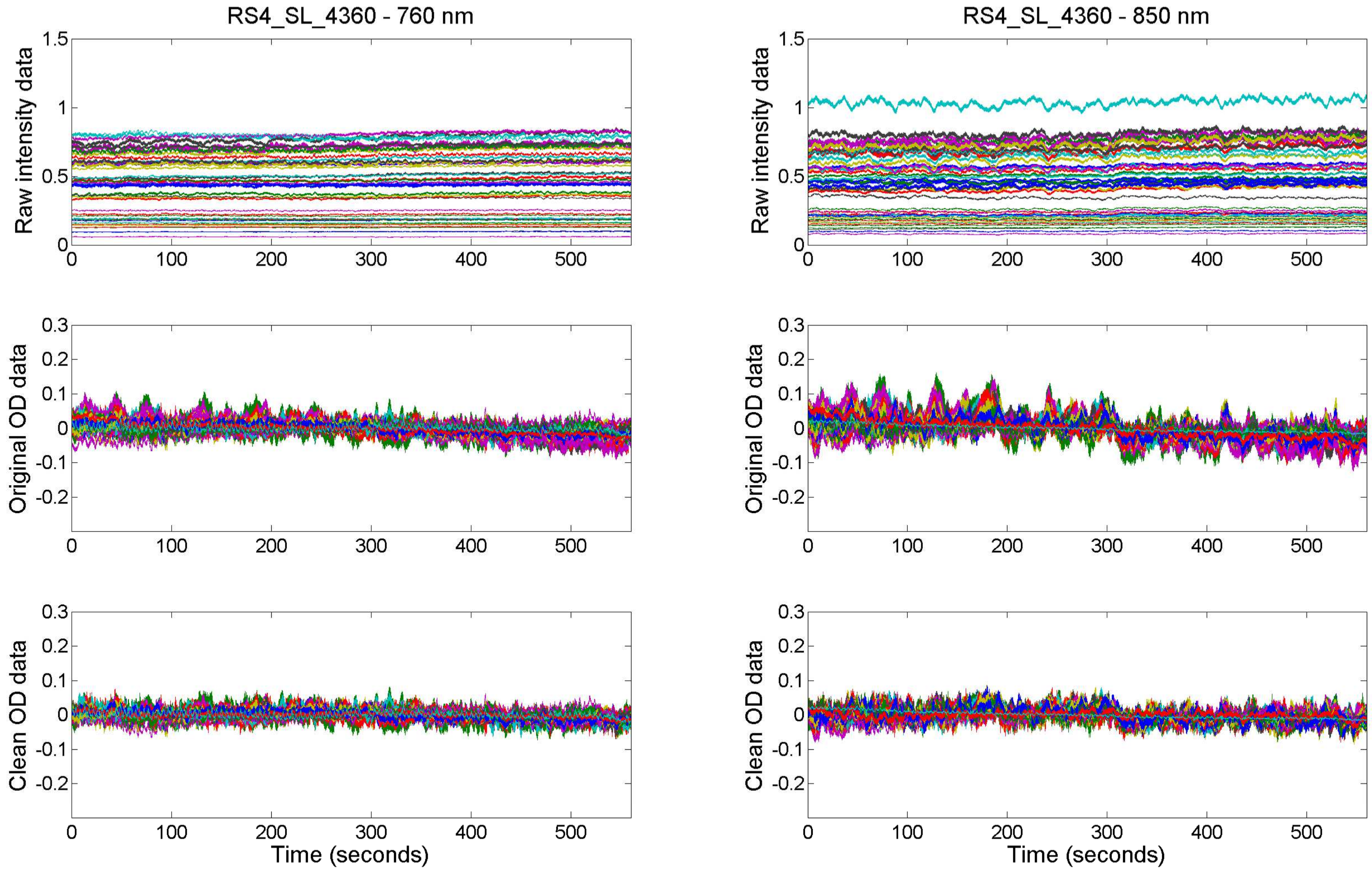

RS4_SL_4360 - 760 nm
RS4_SL_4360 - 850 nm
Raw intensity data
Original OD data
Clean OD data
Time (seconds)

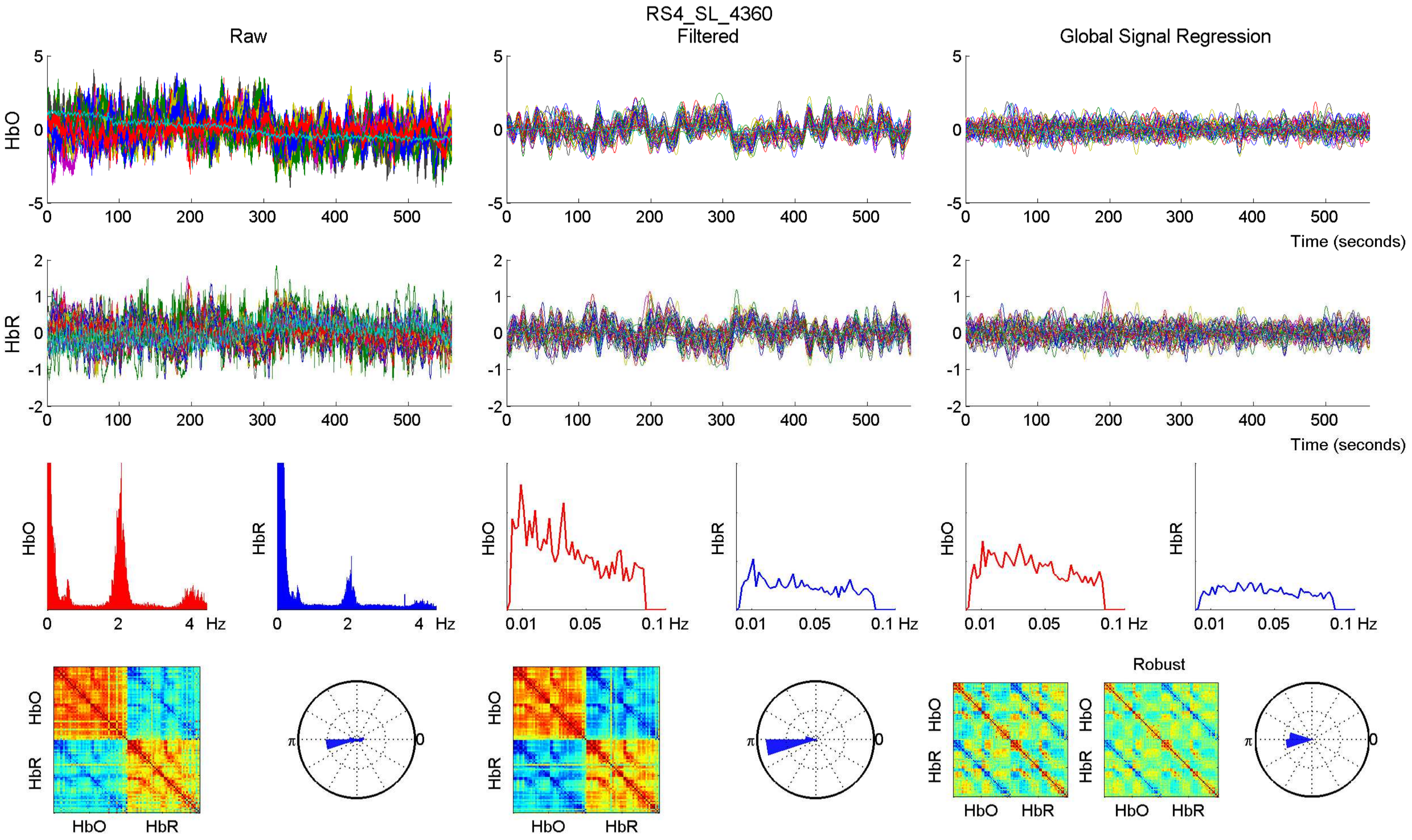

RS4_SL_4360
Raw
Filtered
Global Signal Regression
HbO
HbR
Time (seconds)
Time (seconds)
Hz
0.01
0.05
0.1 Hz
Robust
HbO
HbR
π
0

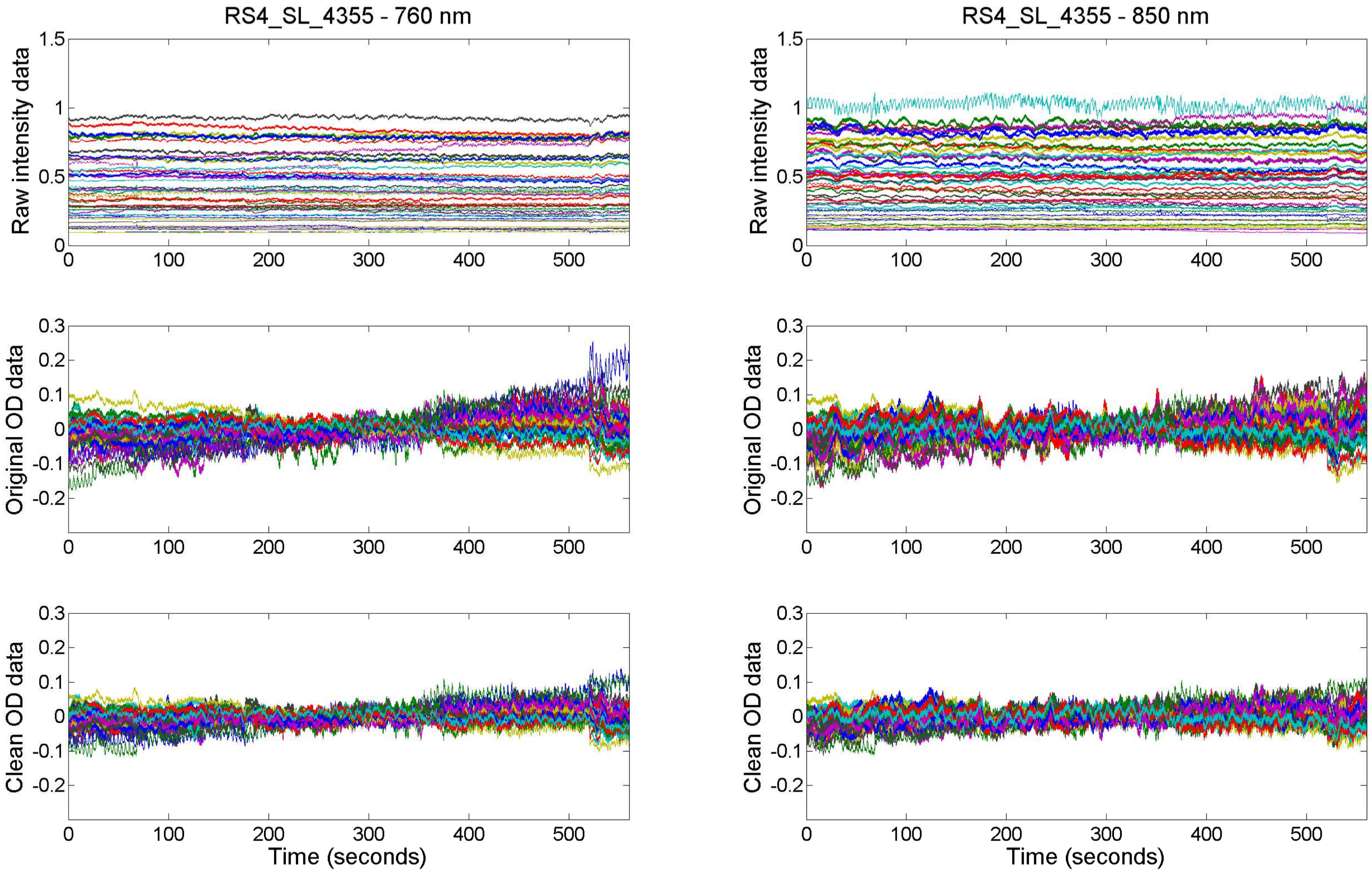

RS4_SL_4355 - 760 nm
RS4_SL_4355 - 850 nm
Raw intensity data
Original OD data
Clean OD data
Time (seconds)

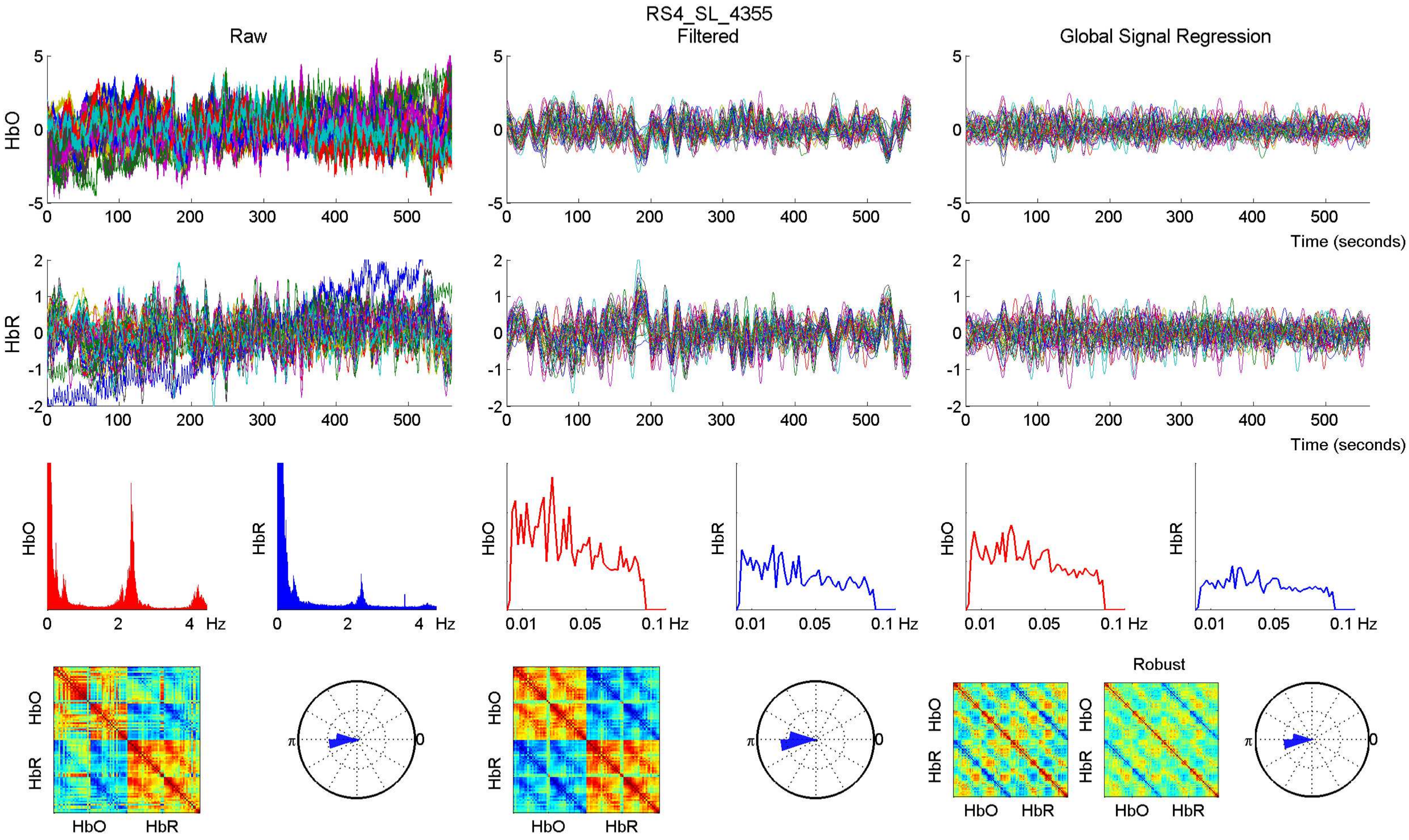

RS4_SL_4355
Raw
Filtered
Global Signal Regression
HbO
HbR
Time (seconds)
Hz
Robust

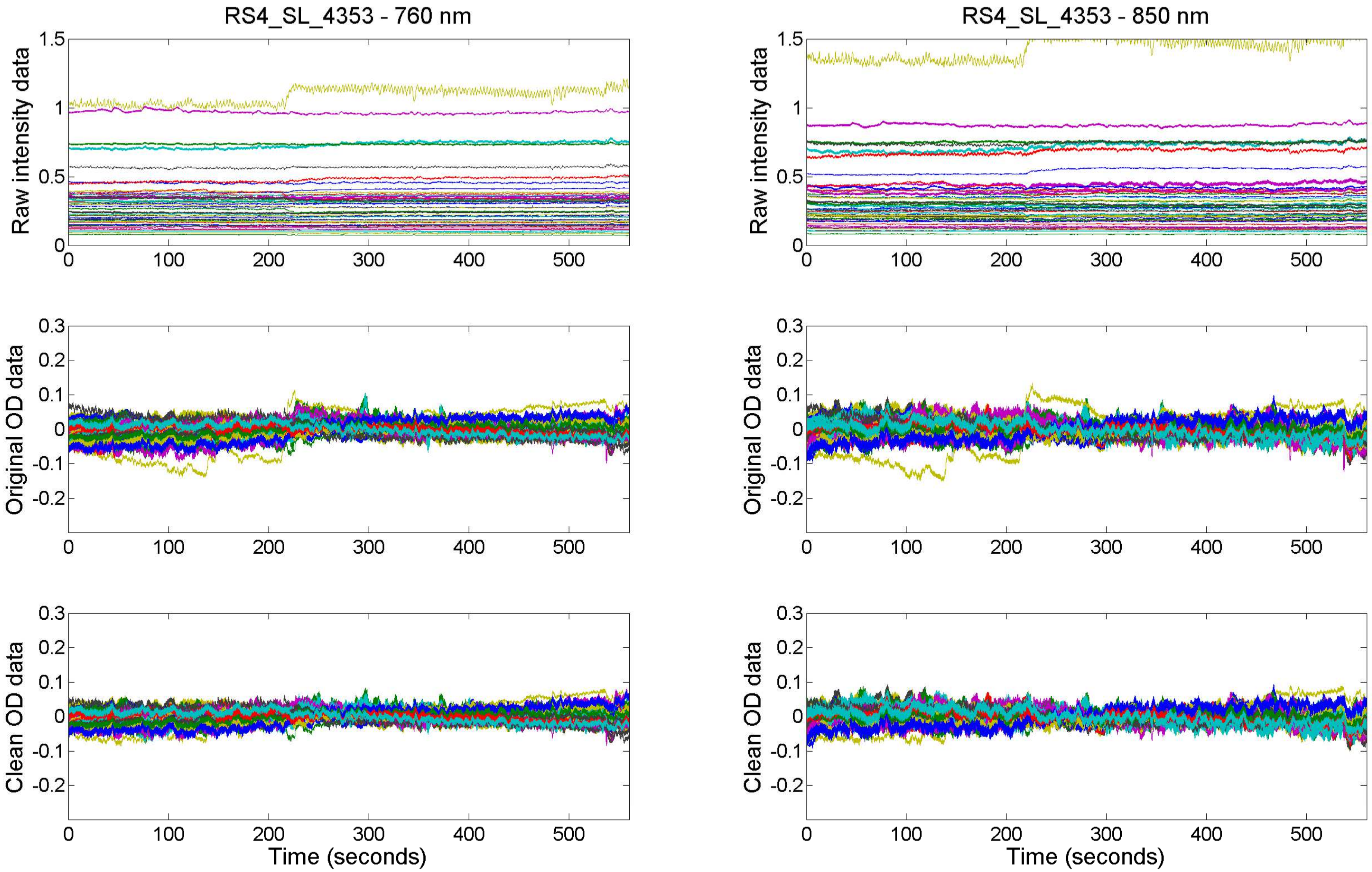

RS4_SL_4353 - 760 nm
RS4_SL_4353 - 850 nm
Raw intensity data
Original OD data
Clean OD data
Time (seconds)

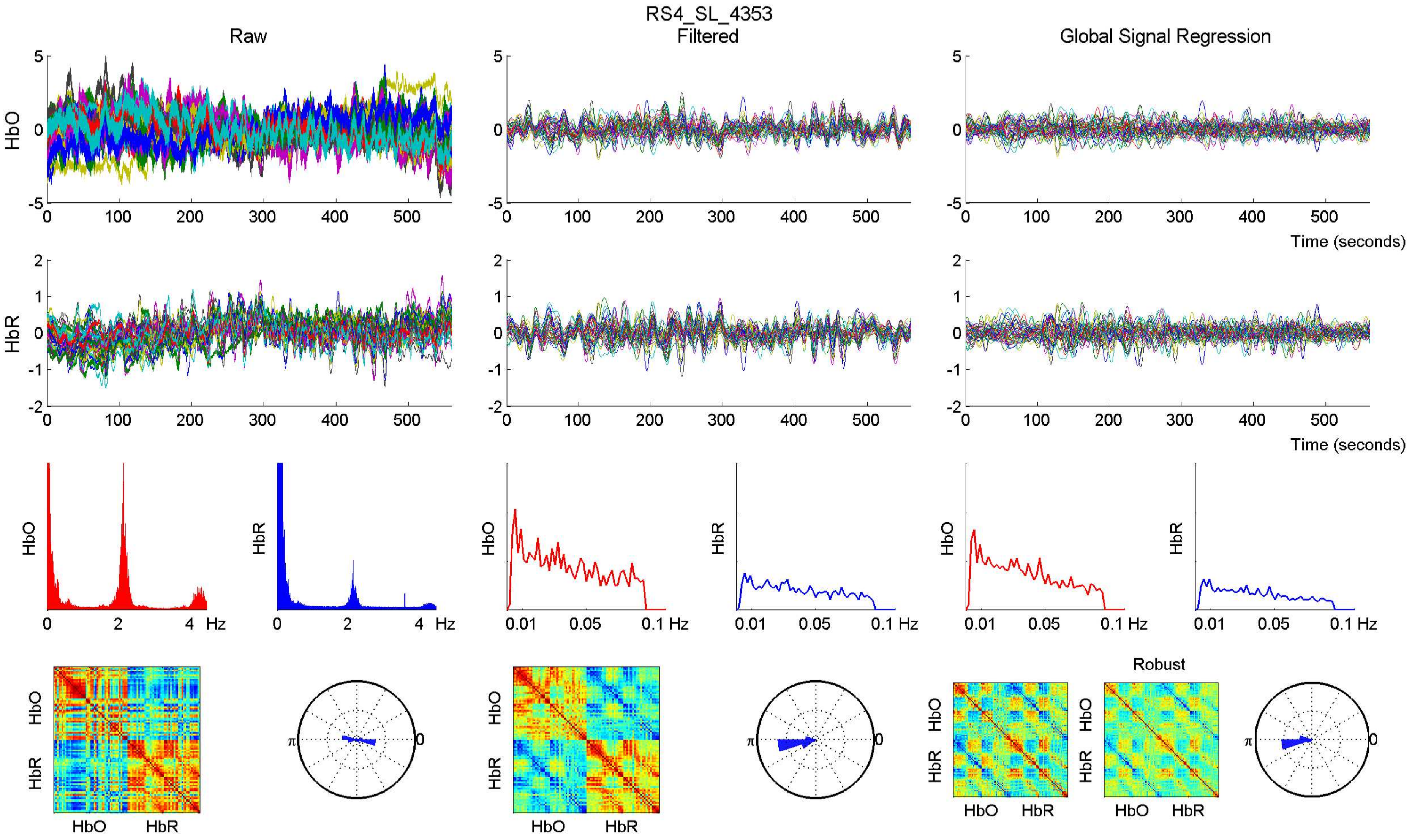
RS4_SL_4353
Raw
Filtered
Global Signal Regression
HbO
HbR
Time (seconds)
Hz
Robust

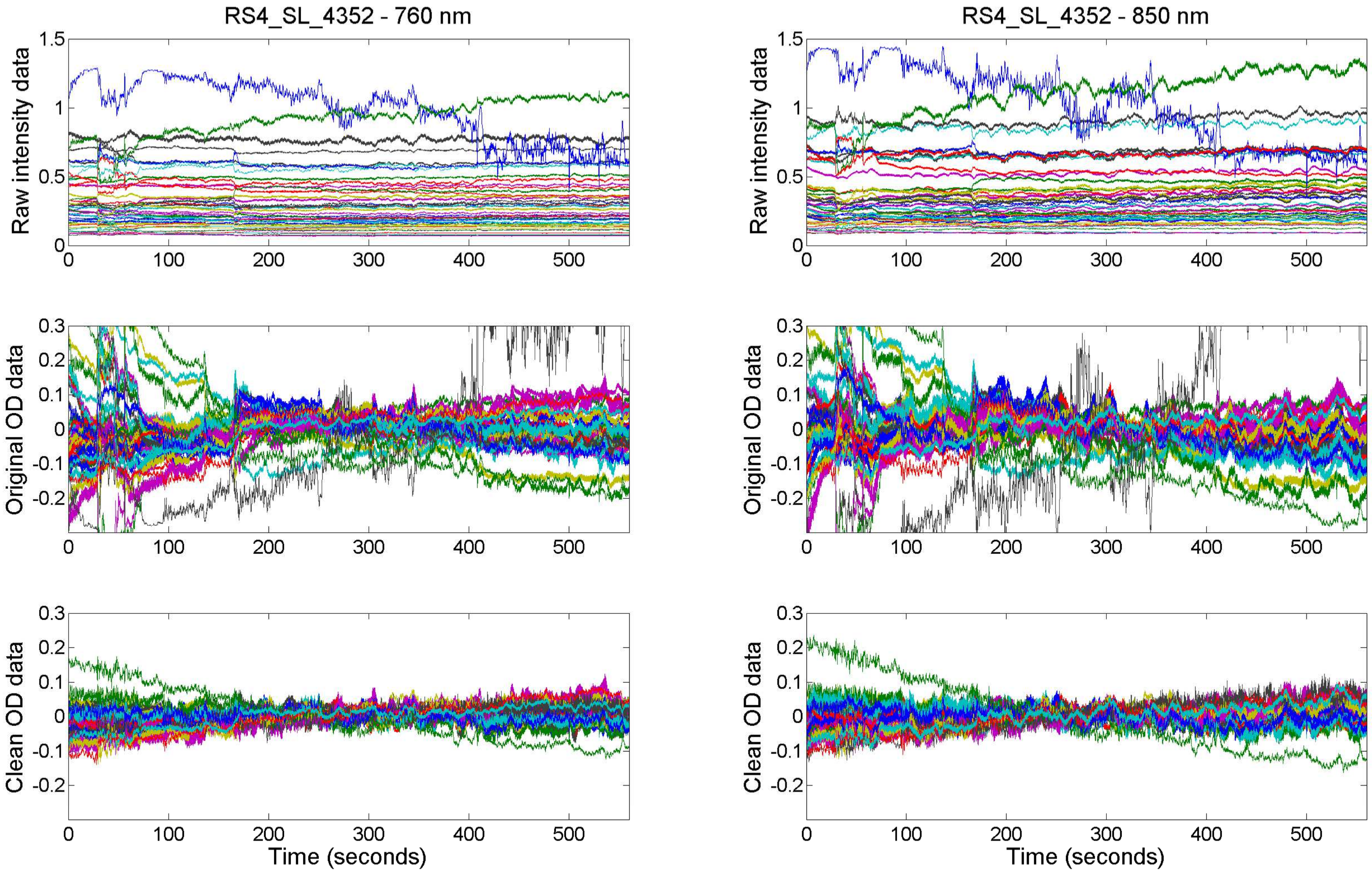
RS4_SL_4352 - 760 nm
RS4_SL_4352 - 850 nm
Raw intensity data
Original OD data
Clean OD data
Time (seconds)
0
100
200
300
400
500
1.5
1
0.5
0.3
0.2
0.1
-0.1
-0.2

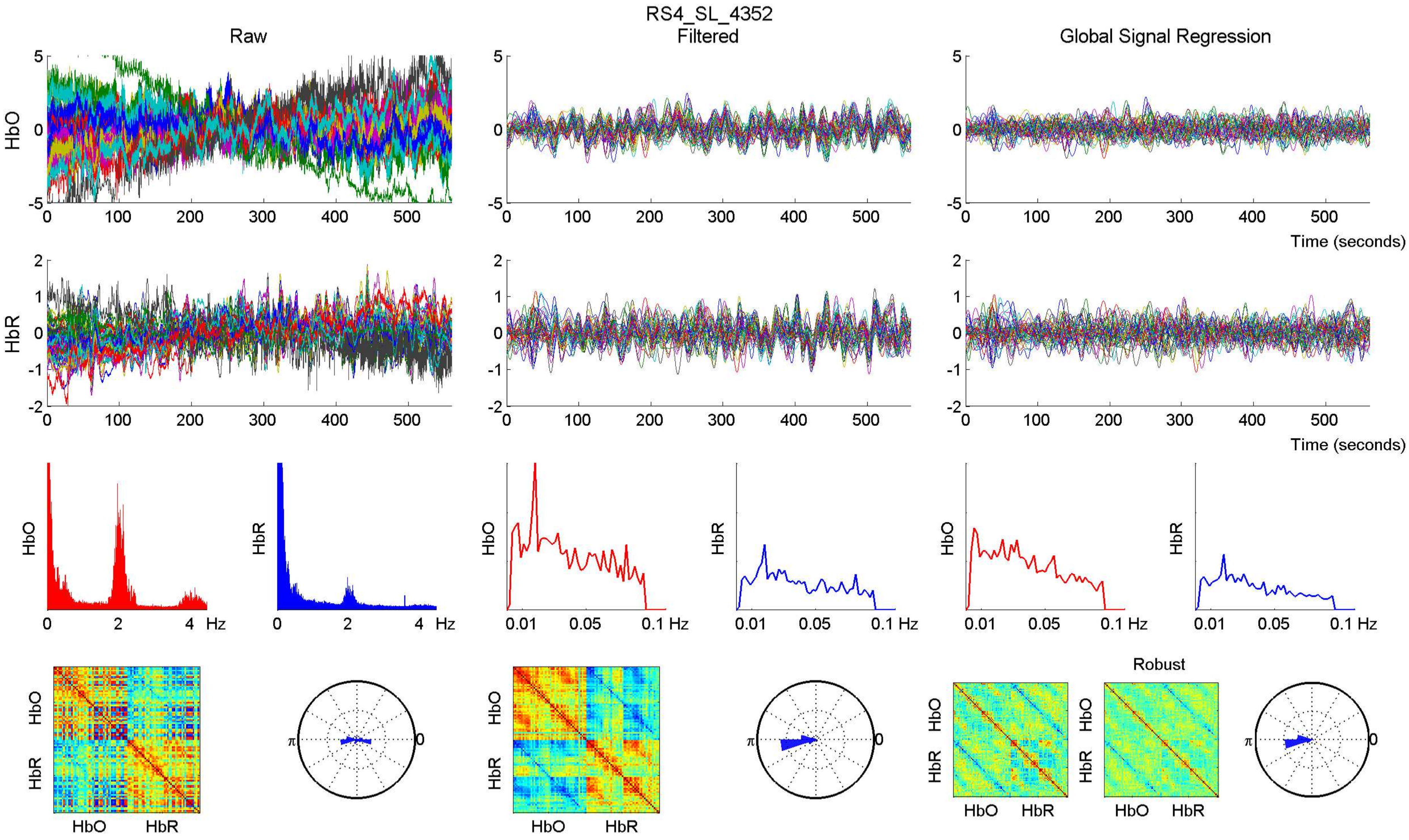

RS4_SL_4352
Raw
Filtered
Global Signal Regression
HbO
HbR
Time (seconds)
Hz
Robust

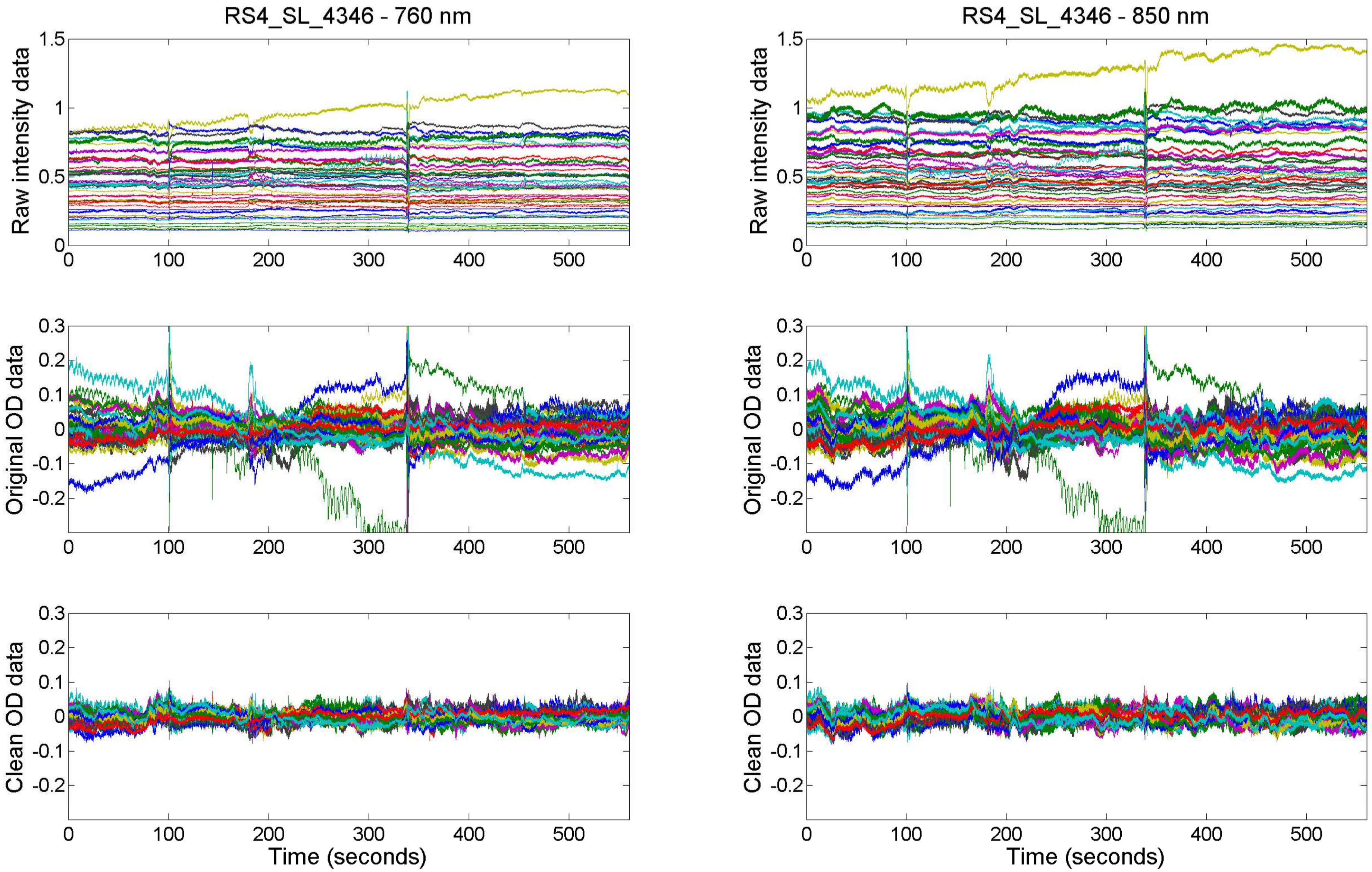

RS4_SL_4346 - 760 nm
RS4_SL_4346 - 850 nm
Raw intensity data
Original OD data
Clean OD data
Time (seconds)

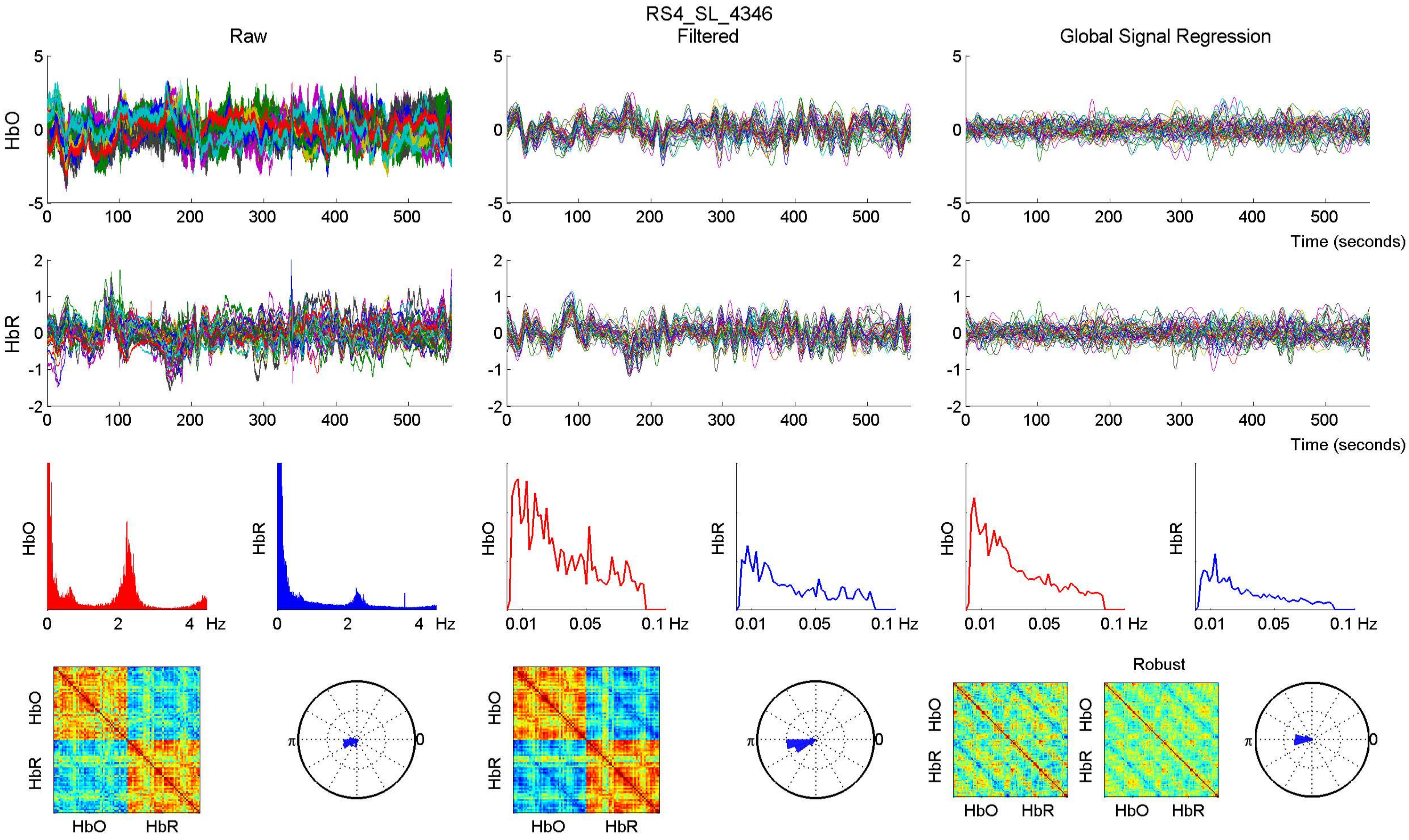

RS4_SL_4346
Raw
Filtered
Global Signal Regression
HbO
HbR
Time (seconds)
Hz
0.01
0.05
0.1 Hz
Robust
π
0
HbO
HbR

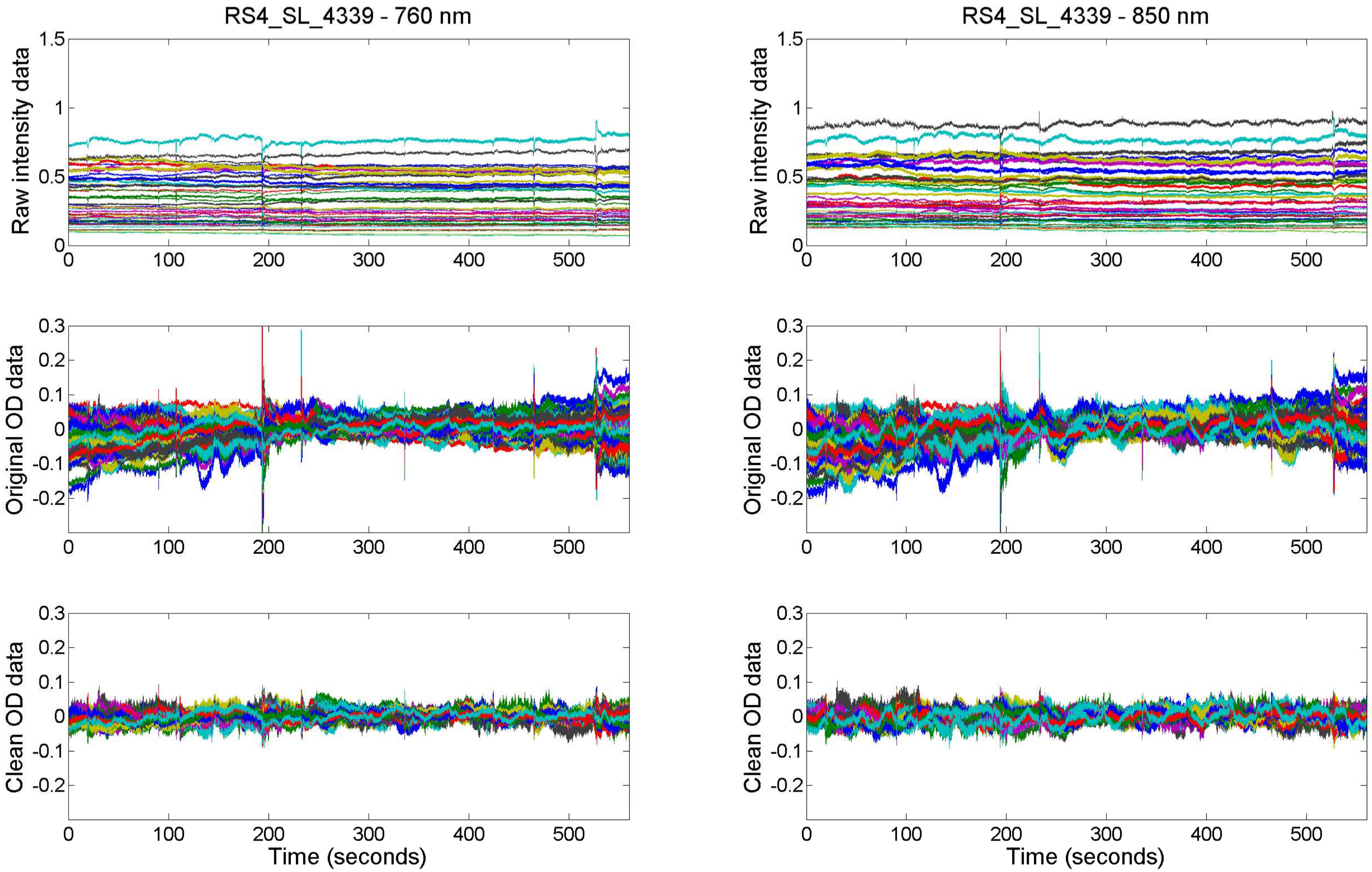
RS4_SL_4339 - 760 nm
RS4_SL_4339 - 850 nm
Raw intensity data
Original OD data
Clean OD data
Time (seconds)
Time (seconds)

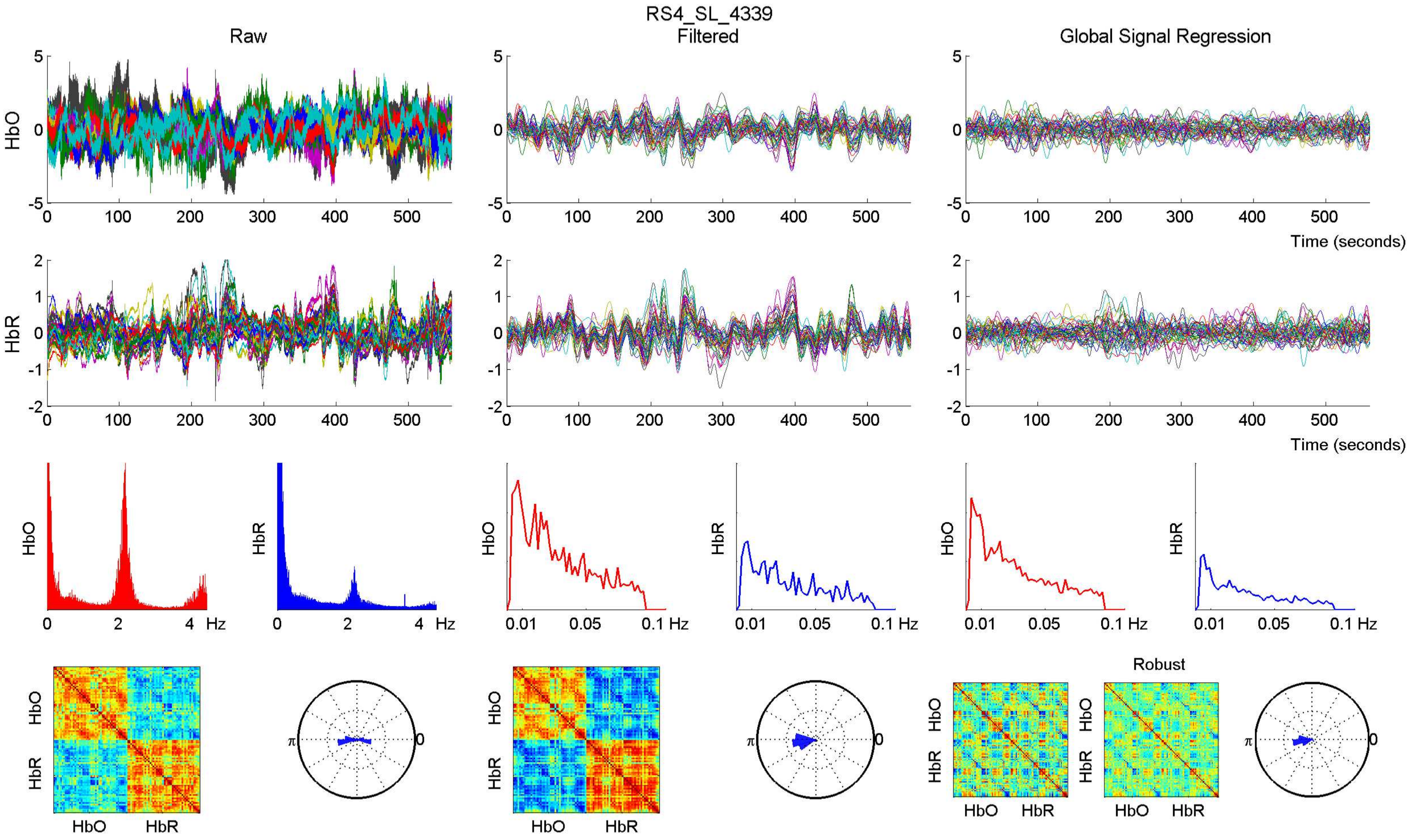
RS4_SL_4339
Raw
Filtered
Global Signal Regression
HbO
HbR
Time (seconds)
Hz
Robust

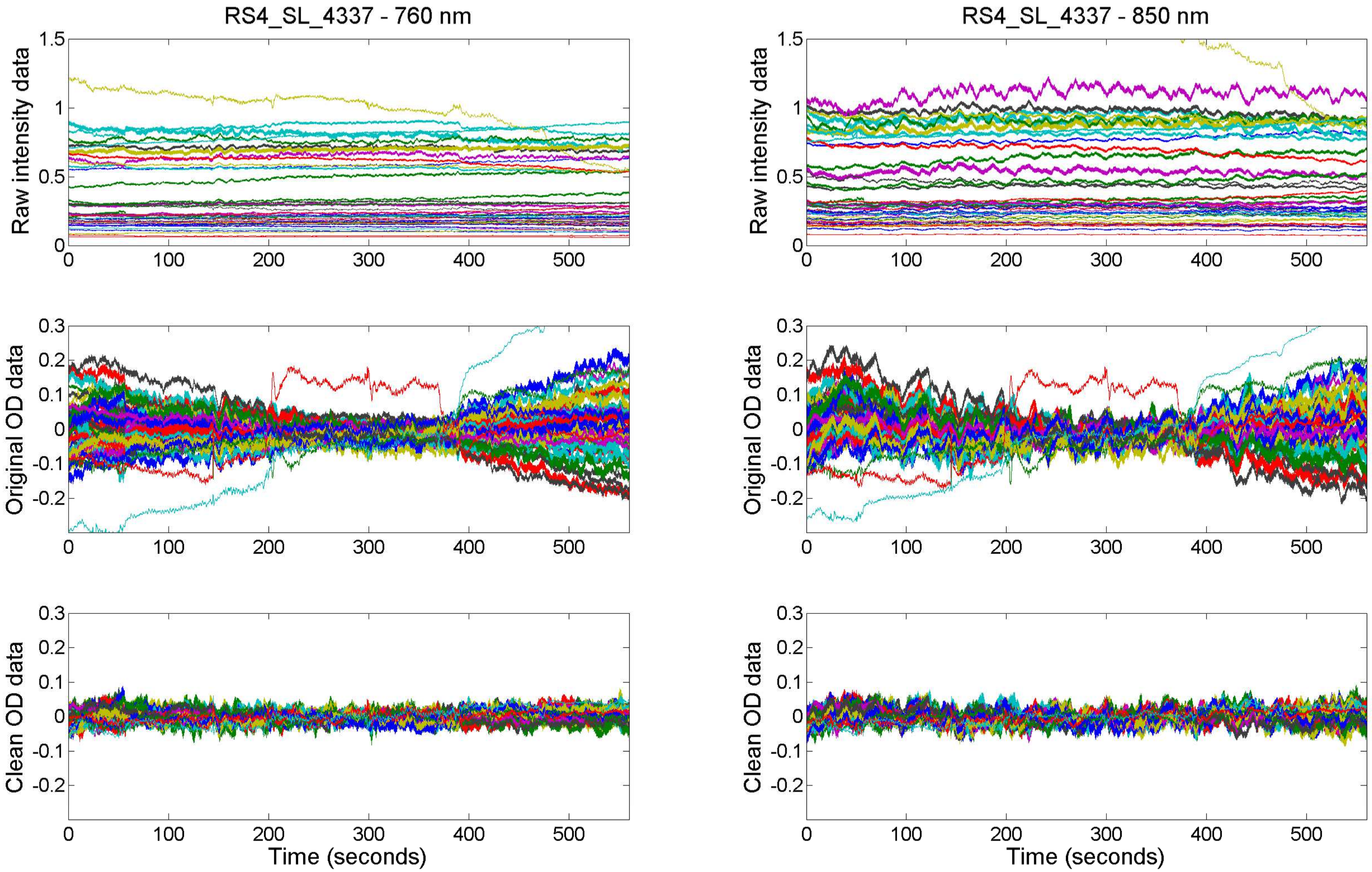
RS4_SL_4337 - 760 nm
RS4_SL_4337 - 850 nm
Raw intensity data
Original OD data
Clean OD data
Time (seconds)
0
100
200
300
400
500
1.5
1
0.5
0.3
0.2
0.1
-0.1
-0.2

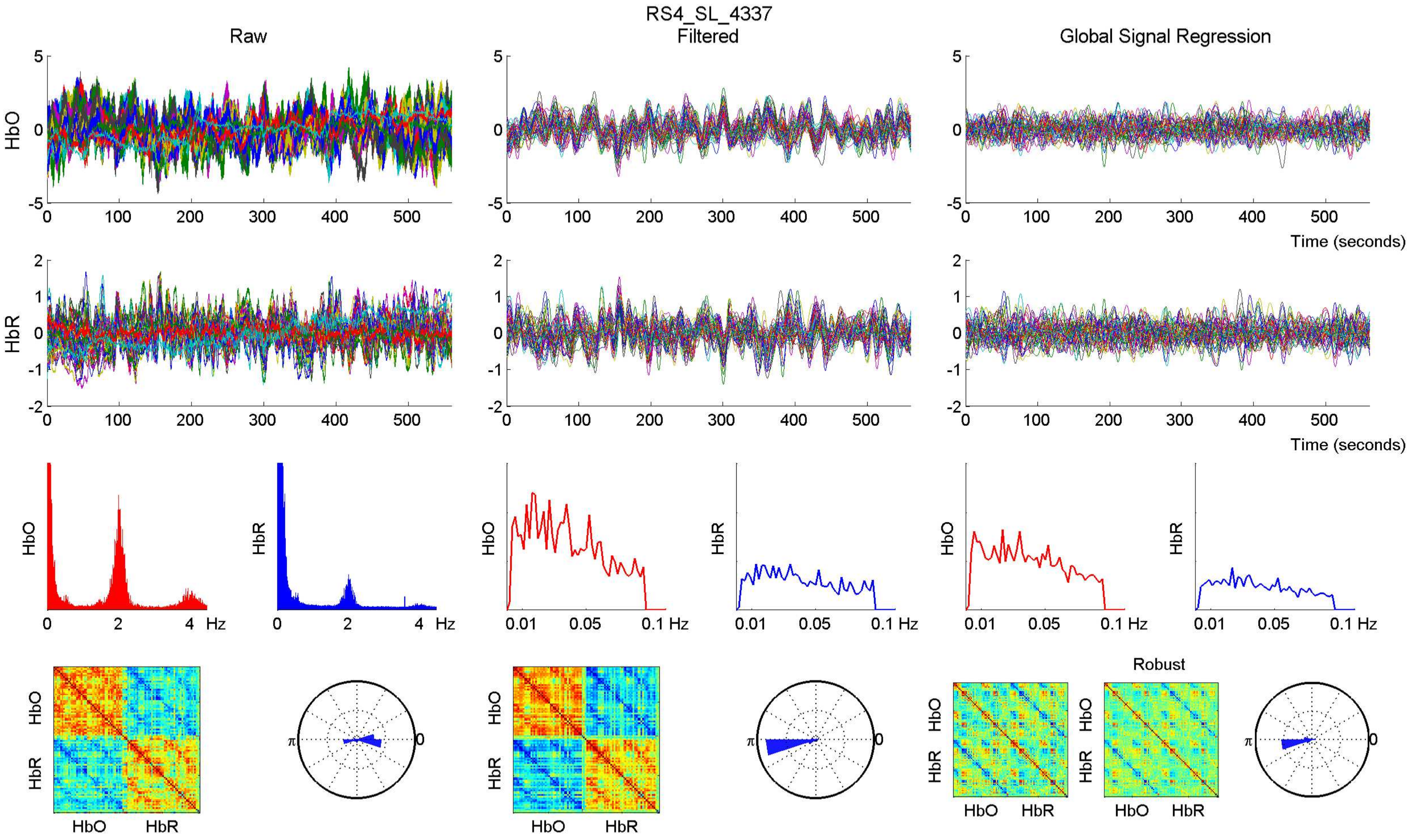

RS4_SL_4337
Raw
Filtered
Global Signal Regression
HbO
HbR
Time (seconds)
Hz
Robust
HbO
HbR

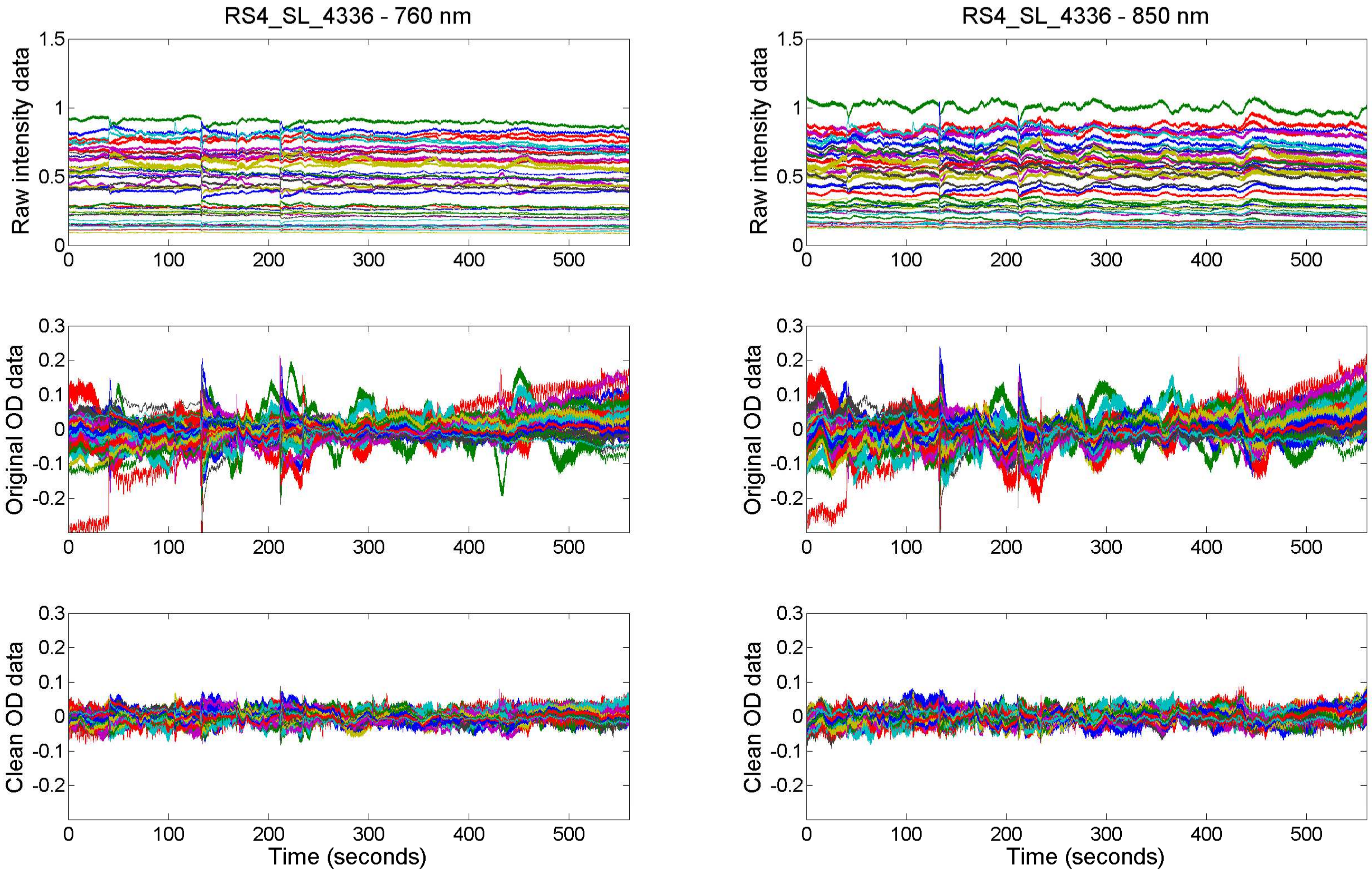
RS4_SL_4336 - 760 nm
RS4_SL_4336 - 850 nm
Raw intensity data
Original OD data
Clean OD data
Time (seconds)
0
100
200
300
400
500
1.5
1
0.5
0.3
0.2
0.1
-0.1
-0.2

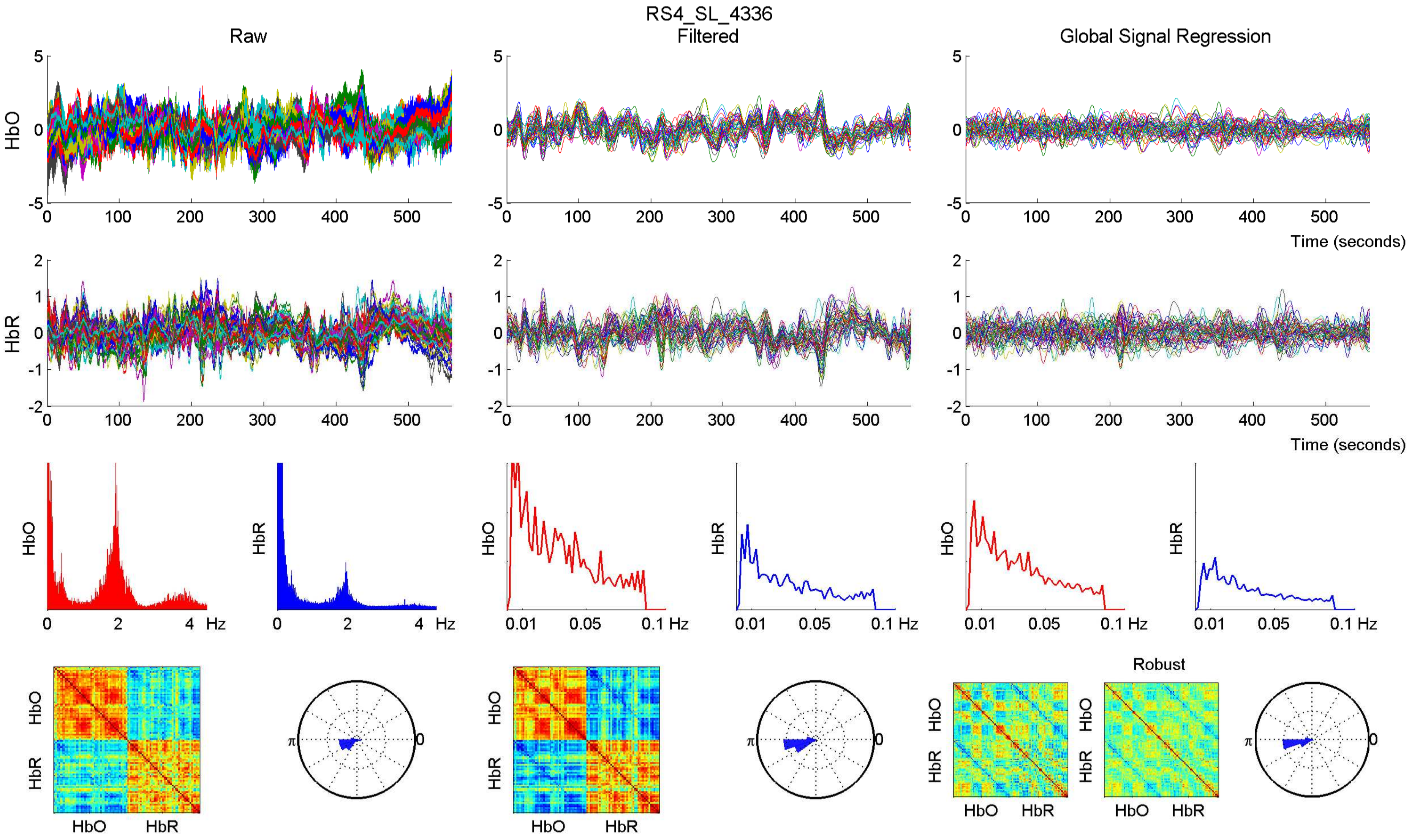

RS4_SL_4336
Raw
Filtered
Global Signal Regression
HbO
HbR
Time (seconds)
Hz
0.01
0.05
0.1 Hz
Robust
HbO
HbR
π
0

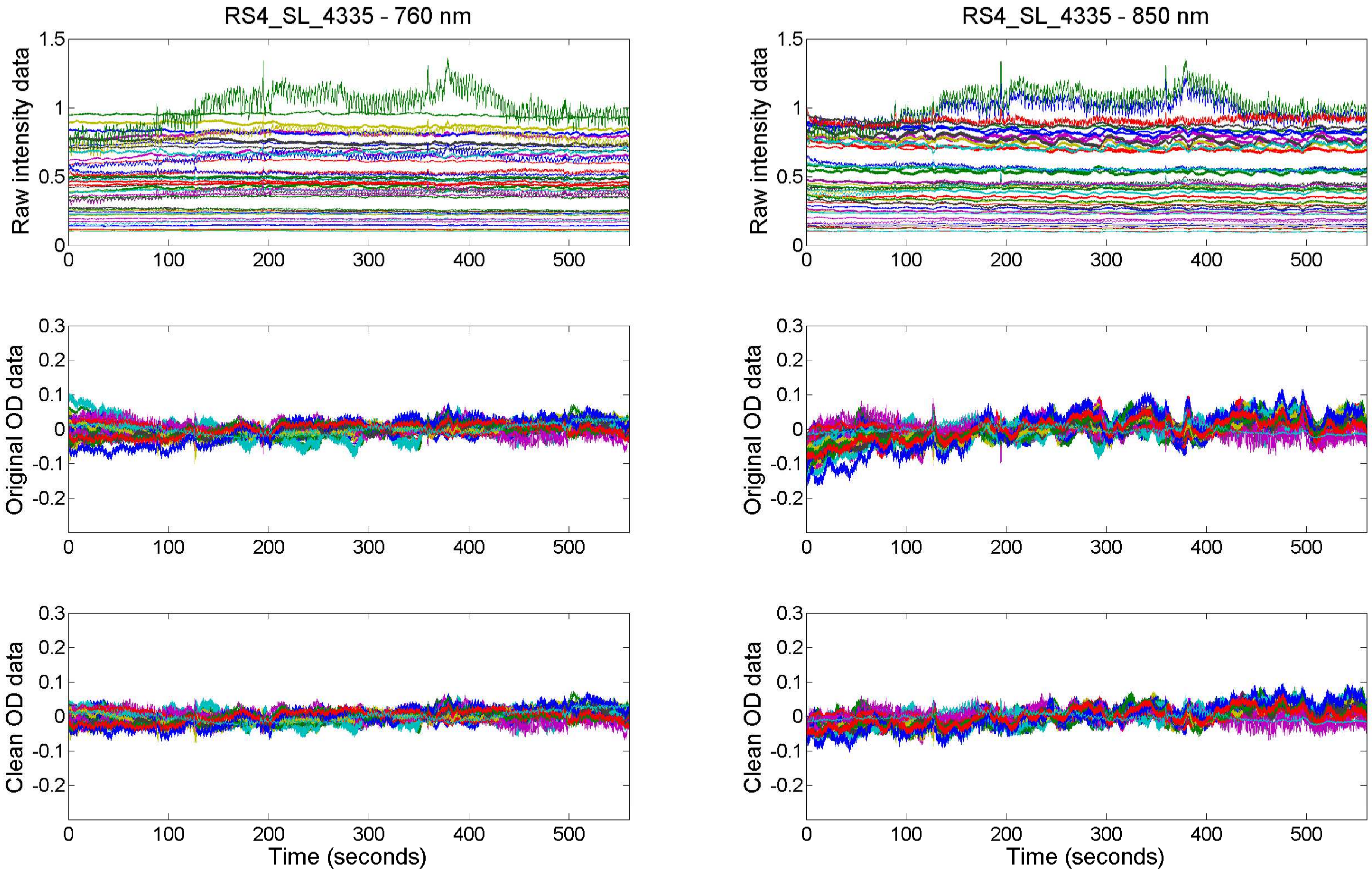
RS4_SL_4335 - 760 nm
RS4_SL_4335 - 850 nm
Raw intensity data
Original OD data
Clean OD data
Time (seconds)

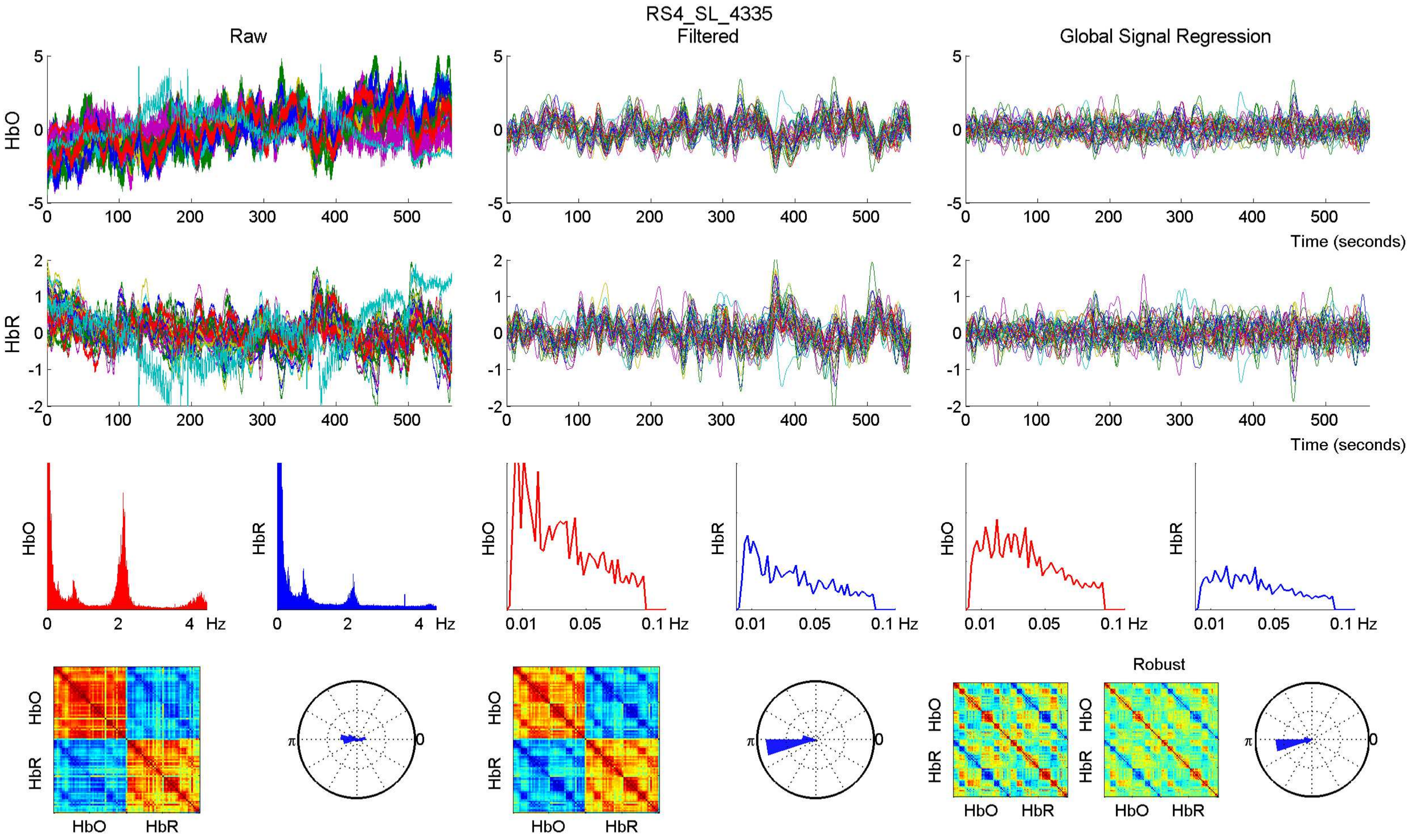
RS4_SL_4335
Raw
Filtered
Global Signal Regression
HbO
HbR
Time (seconds)
Hz
0.01
0.05
0.1 Hz
Robust
HbO
HbR
π
0

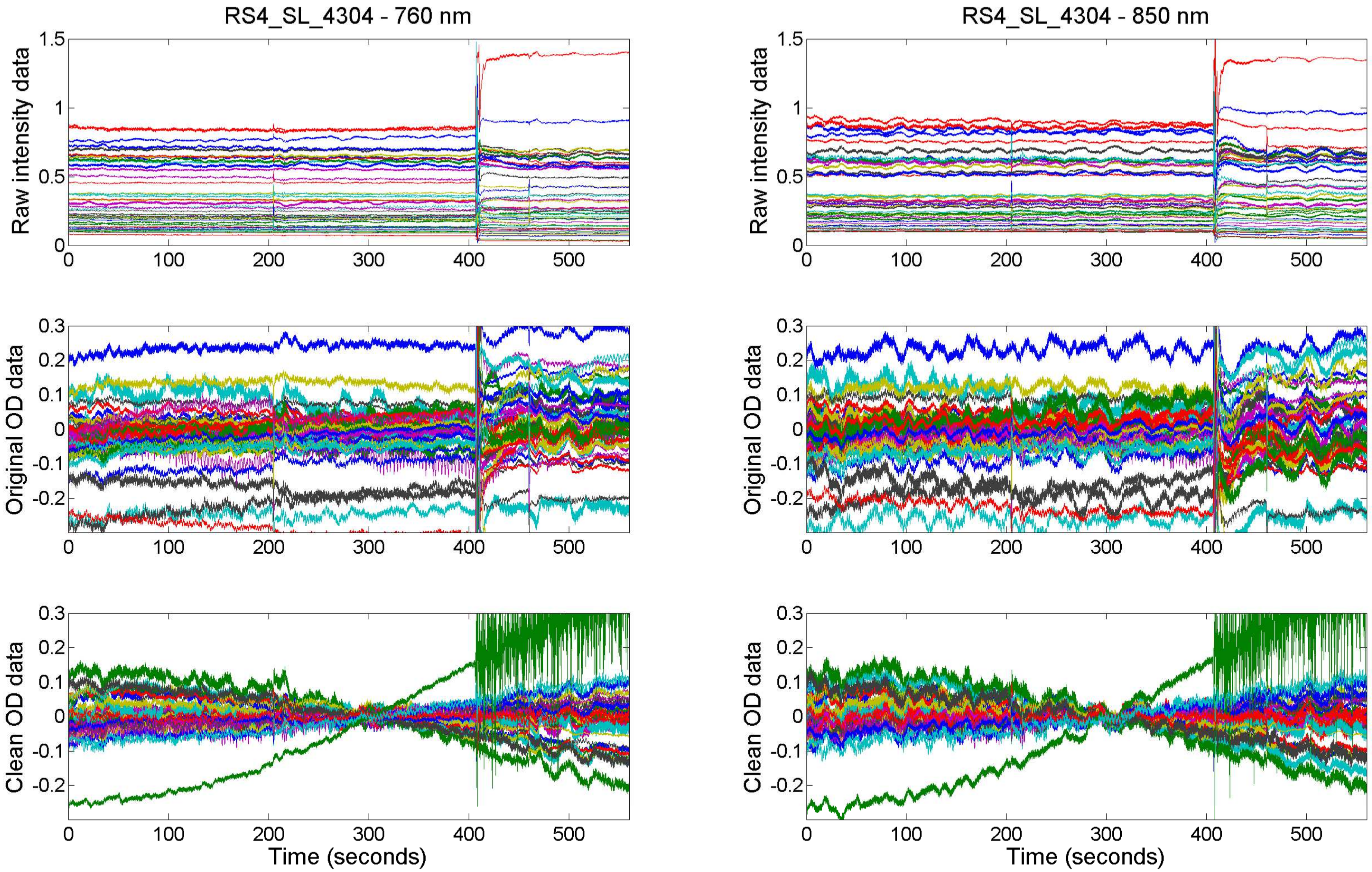

RS4_SL_4304 - 760 nm
RS4_SL_4304 - 850 nm
Raw intensity data
Original OD data
Clean OD data
Time (seconds)

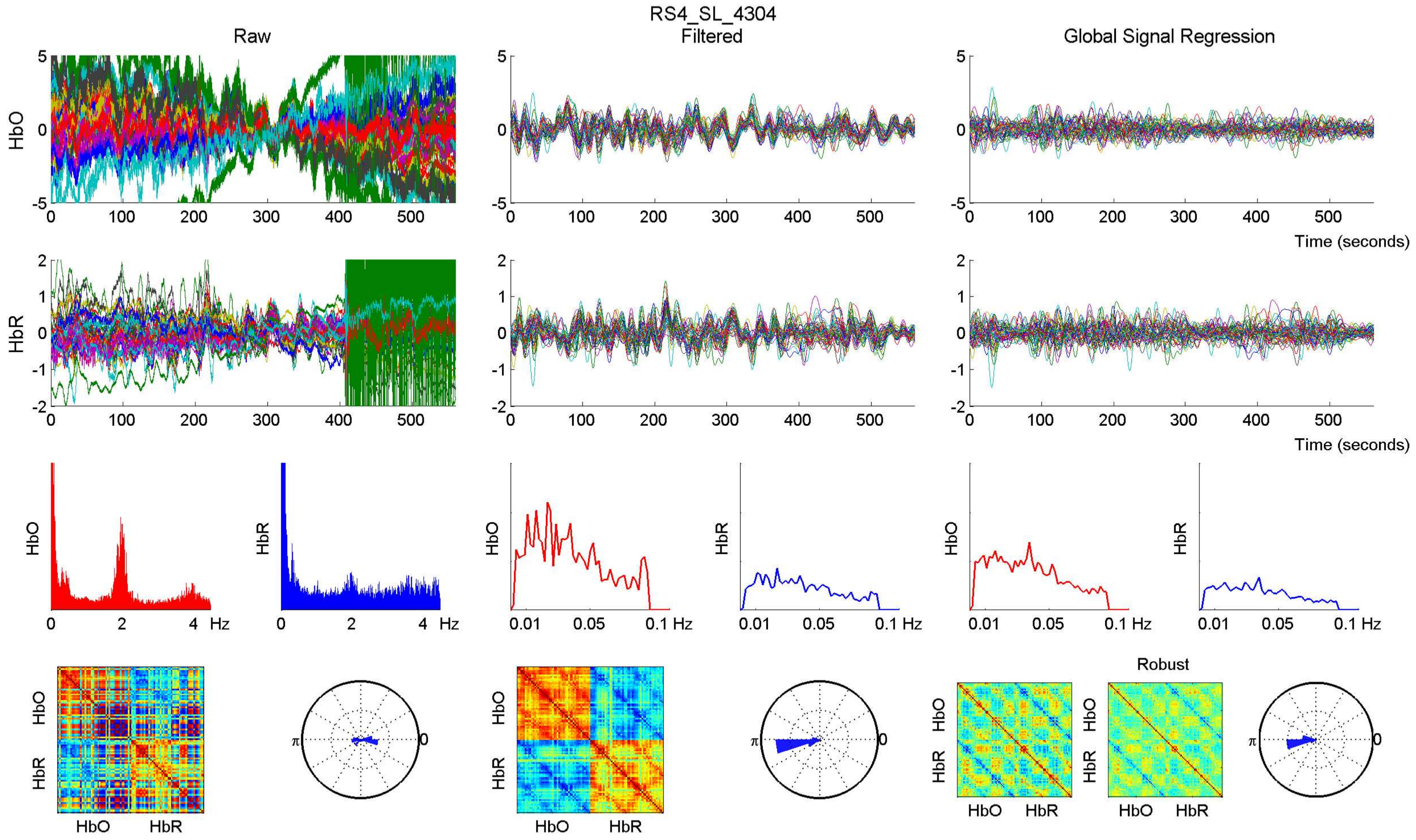
RS4_SL_4304
Raw
Filtered
Global Signal Regression
HbO
HbR
Time (seconds)
Hz
Robust
HbO
HbR

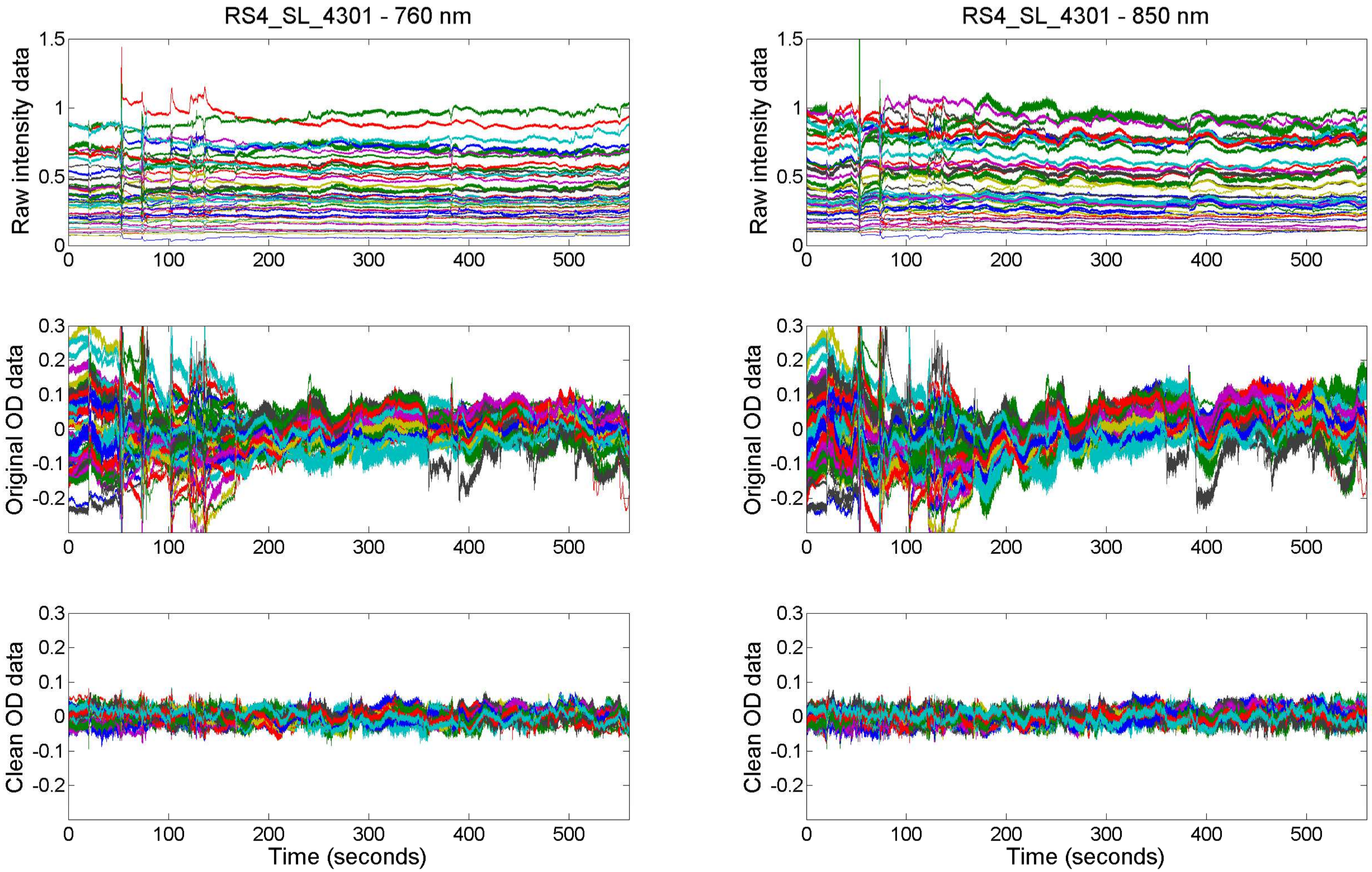

RS4_SL_4301 - 760 nm
RS4_SL_4301 - 850 nm
Raw intensity data
Original OD data
Clean OD data
Time (seconds)

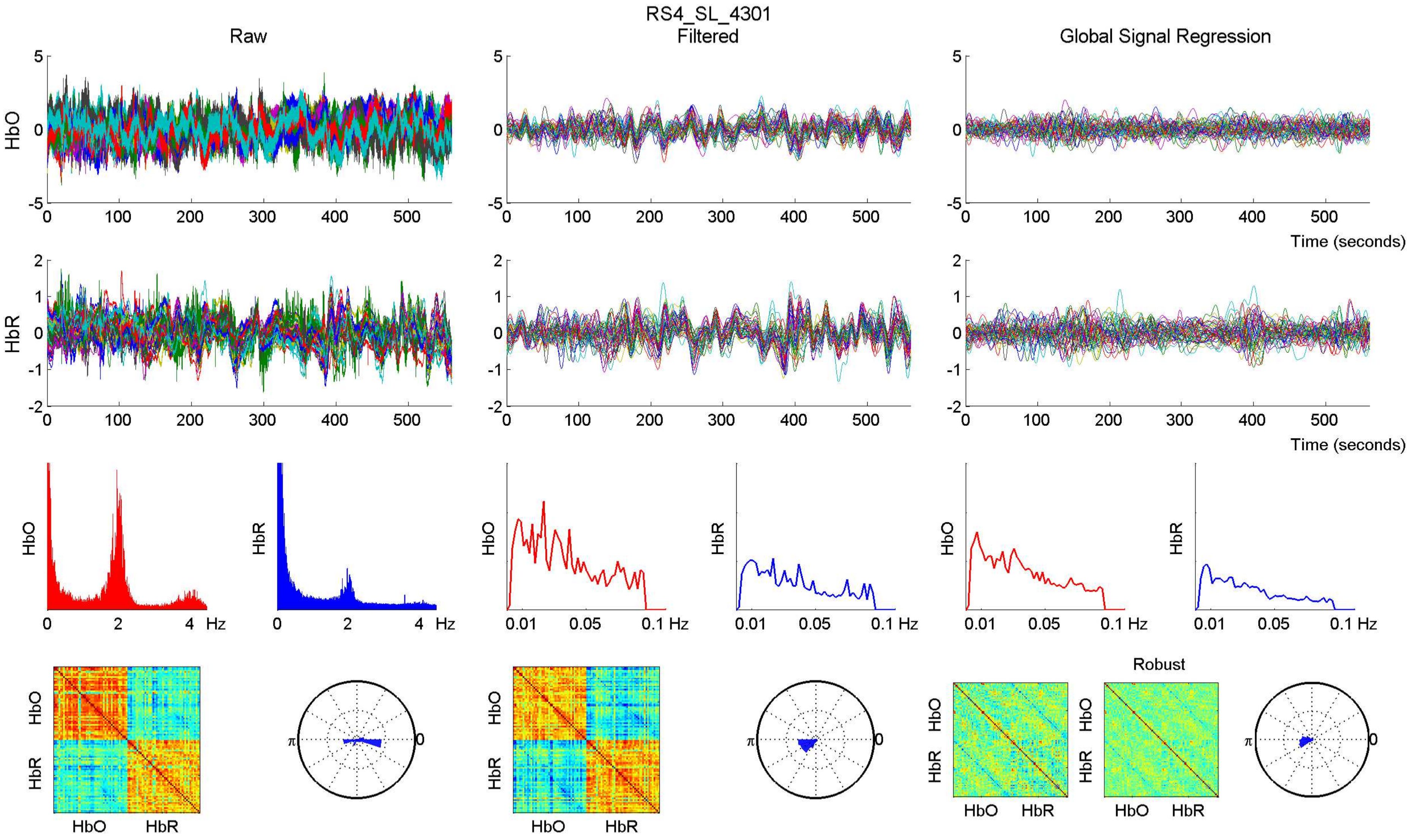
RS4_SL_4301
Raw
Filtered
Global Signal Regression
HbO
HbR
Time (seconds)
Hz
Robust

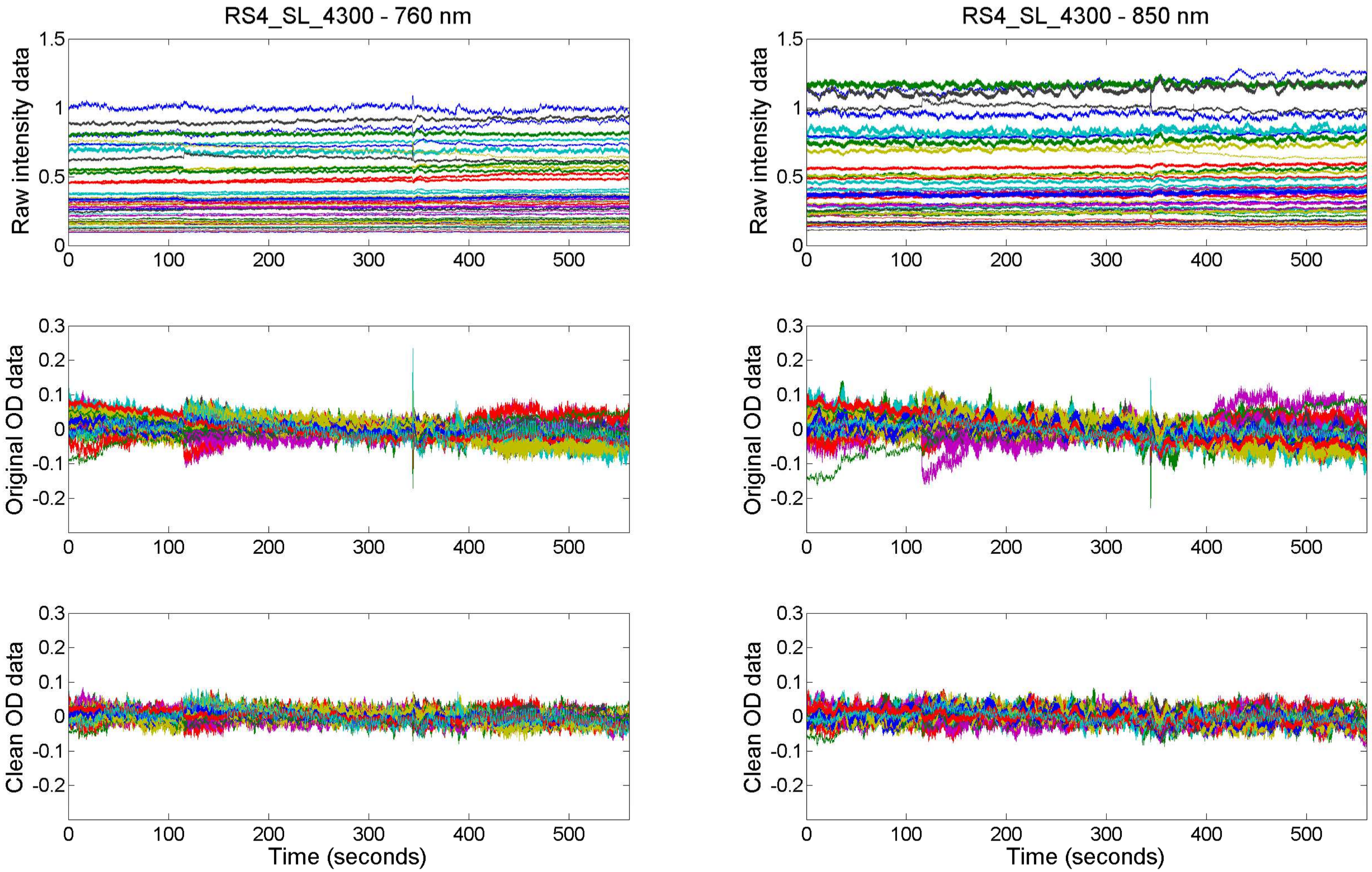

RS4_SL_4300 - 760 nm
RS4_SL_4300 - 850 nm
Raw intensity data
Original OD data
Clean OD data
Time (seconds)
1.5
1
0.5
0
0.3
0.2
0.1
-0.1
-0.2
100
200
300
400
500

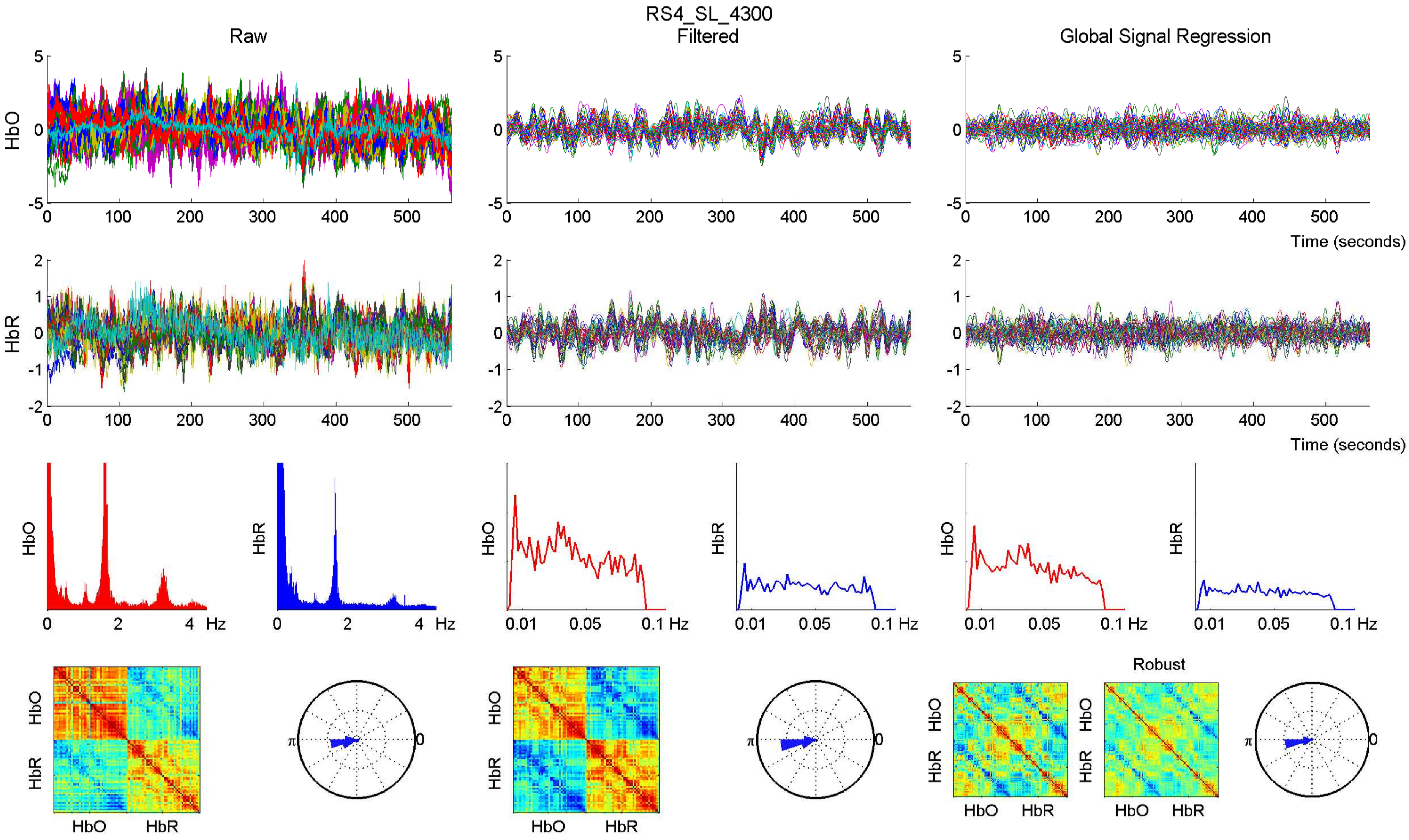

RS4_SL_4300
Raw
Filtered
Global Signal Regression
HbO
HbR
Time (seconds)
Hz
Robust

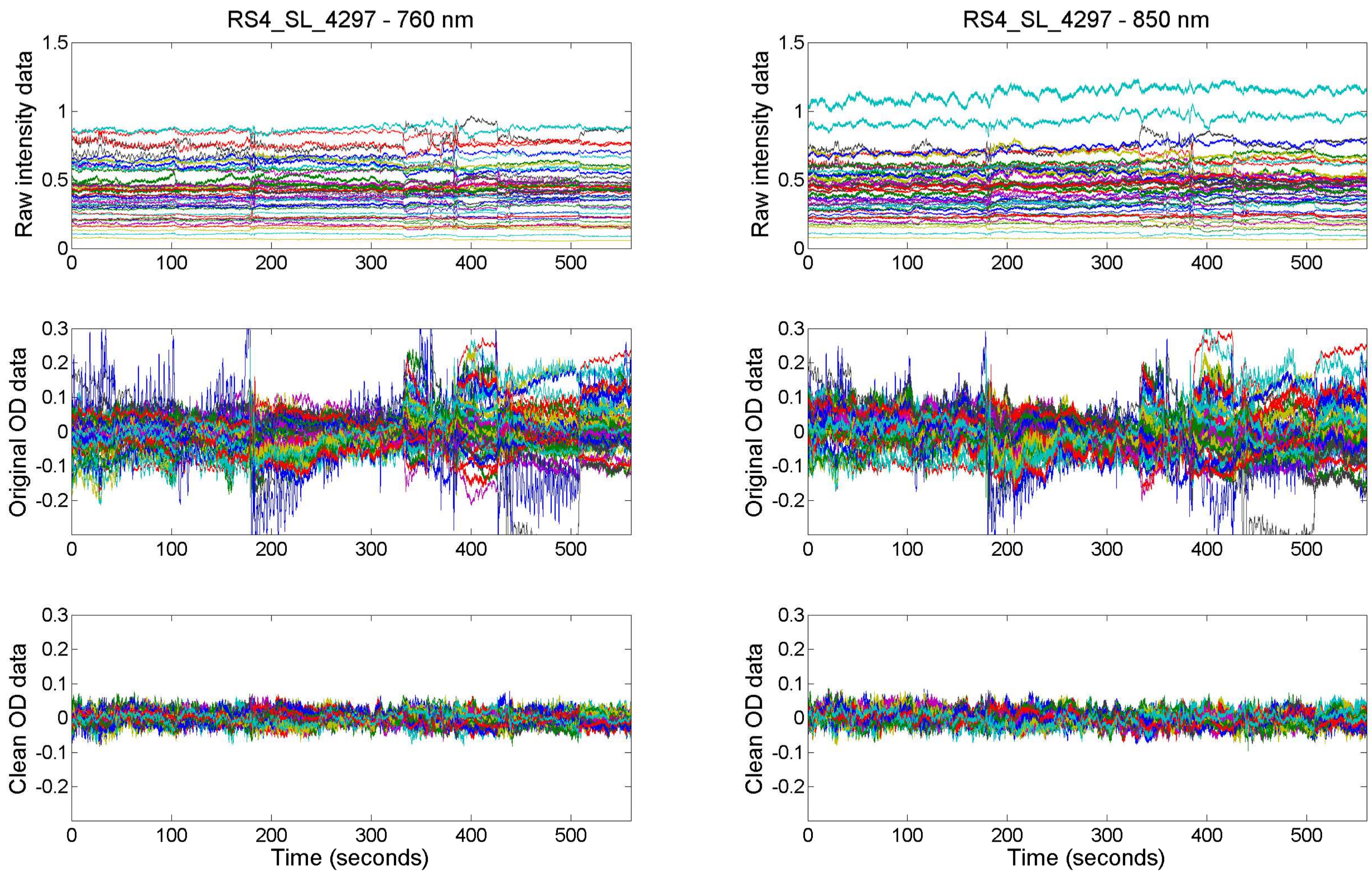
RS4_SL_4297 - 760 nm
RS4_SL_4297 - 850 nm
Raw intensity data
Original OD data
Clean OD data
Time (seconds)
0
100
200
300
400
500
1.5
1
0.5
0.3
0.2
0.1
-0.1
-0.2

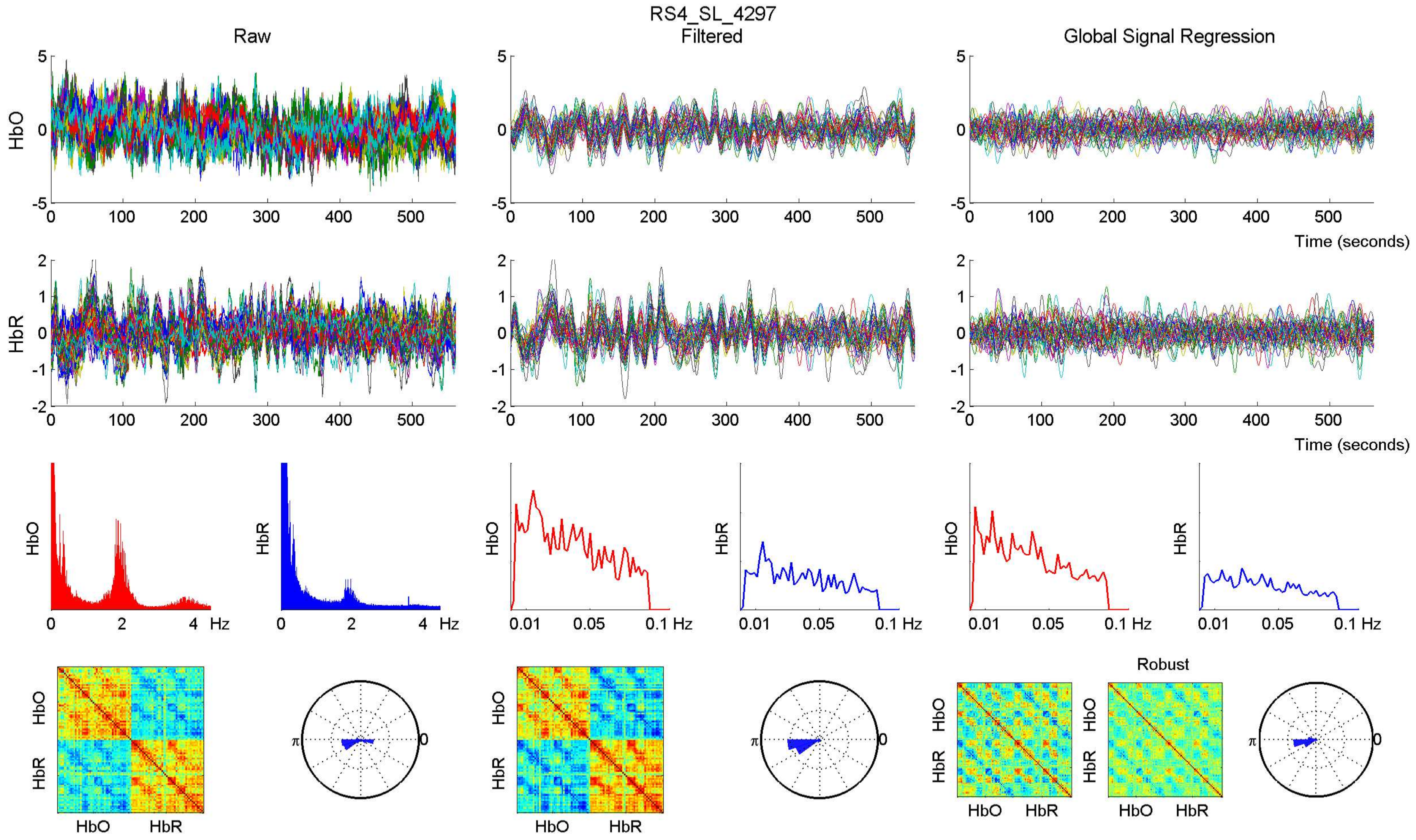

RS4_SL_4297
Raw
Filtered
Global Signal Regression
HbO
HbR
Time (seconds)
Hz
Robust
HbO
HbR

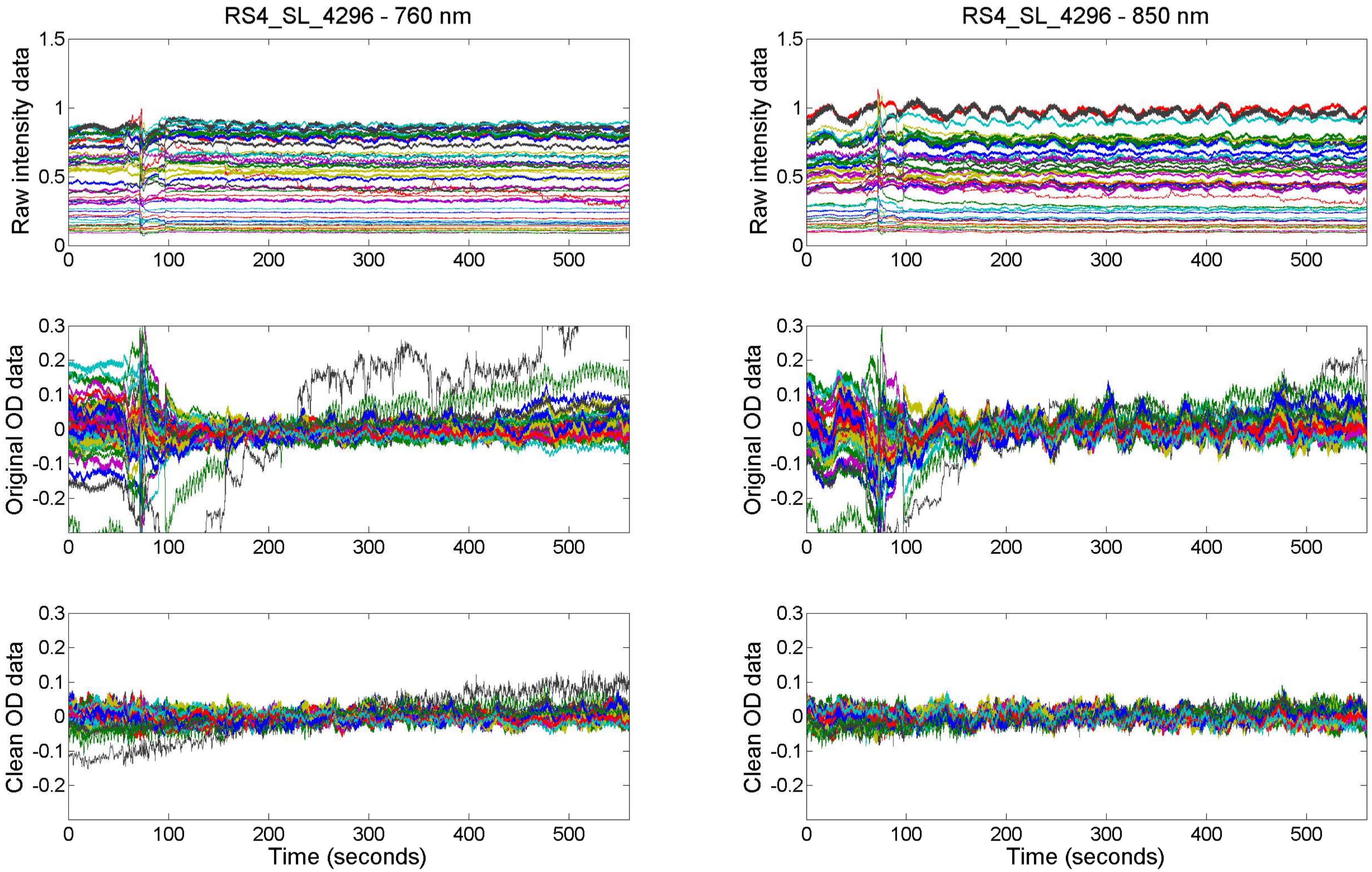

RS4_SL_4296 - 760 nm
RS4_SL_4296 - 850 nm
Raw intensity data
Original OD data
Clean OD data
Time (seconds)

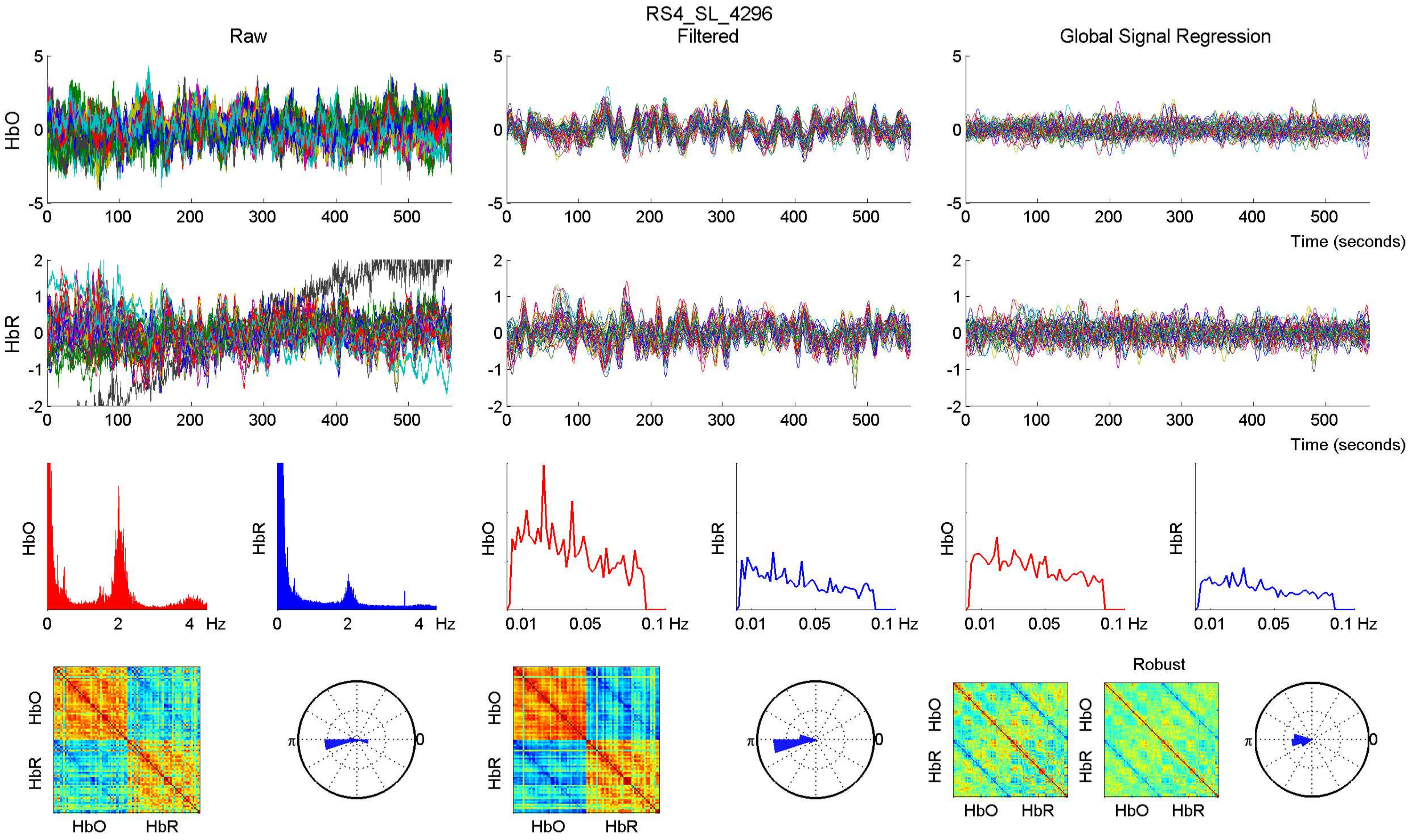
RS4_SL_4296
Raw
Filtered
Global Signal Regression
HbO
HbR
Time (seconds)
Hz
Robust
HbO
HbR

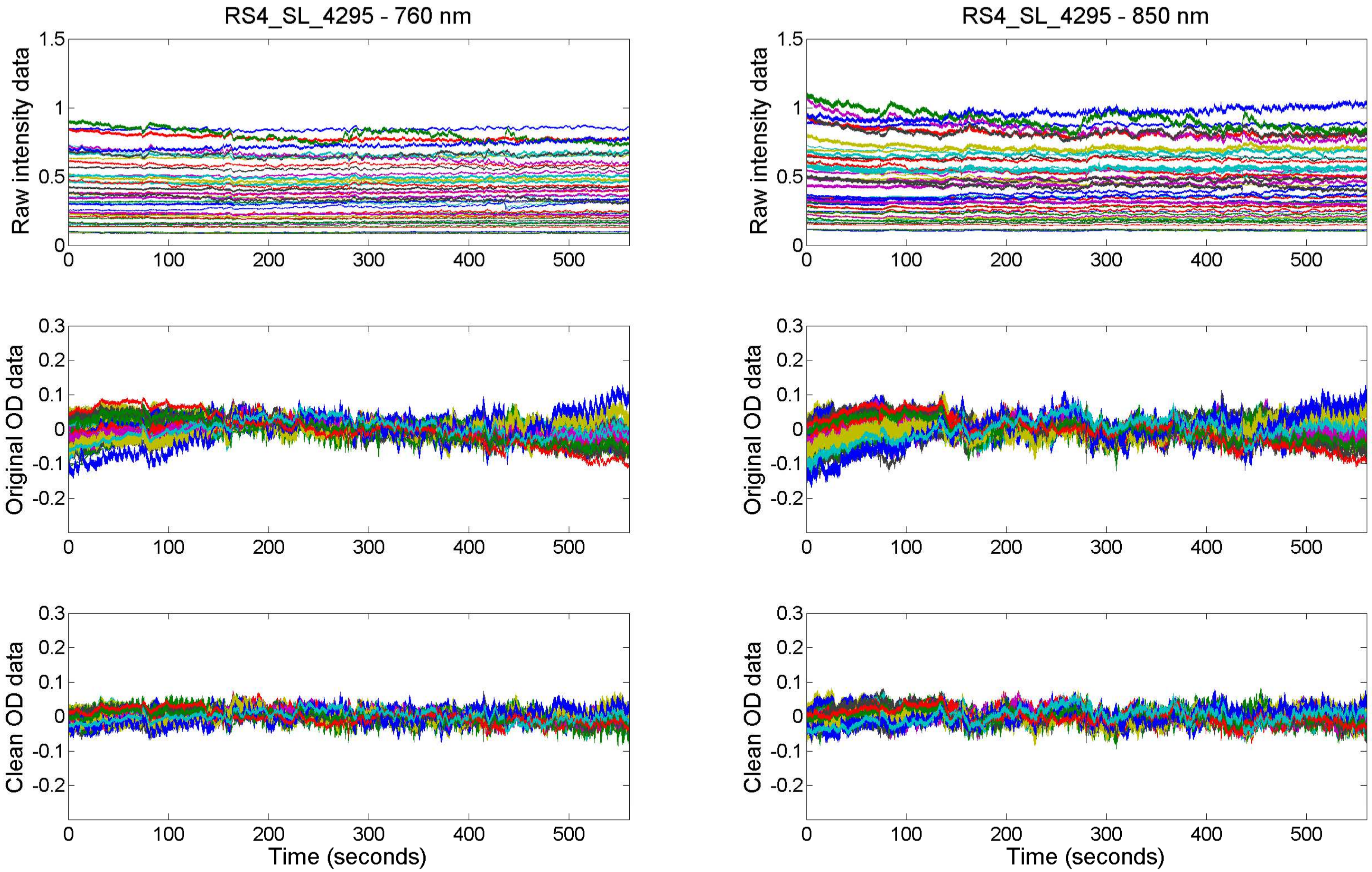
RS4_SL_4295 - 760 nm
RS4_SL_4295 - 850 nm
Raw intensity data
Original OD data
Clean OD data
Time (seconds)
0
100
200
300
400
500
1.5
1
0.5
0.3
0.2
0.1
-0.1
-0.2

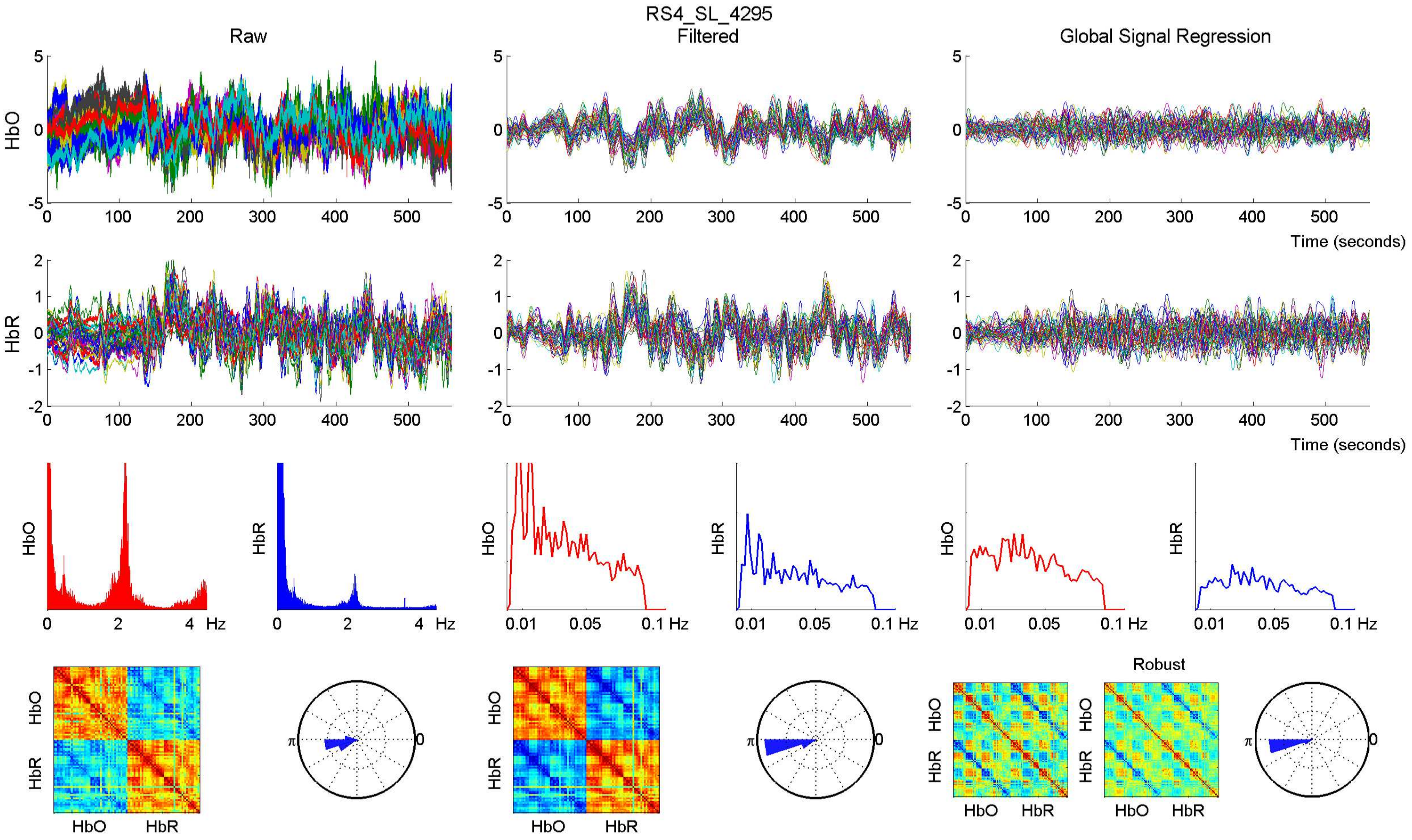

RS4_SL_4295
Raw
Filtered
Global Signal Regression
HbO
HbR
Time (seconds)
Hz
Robust

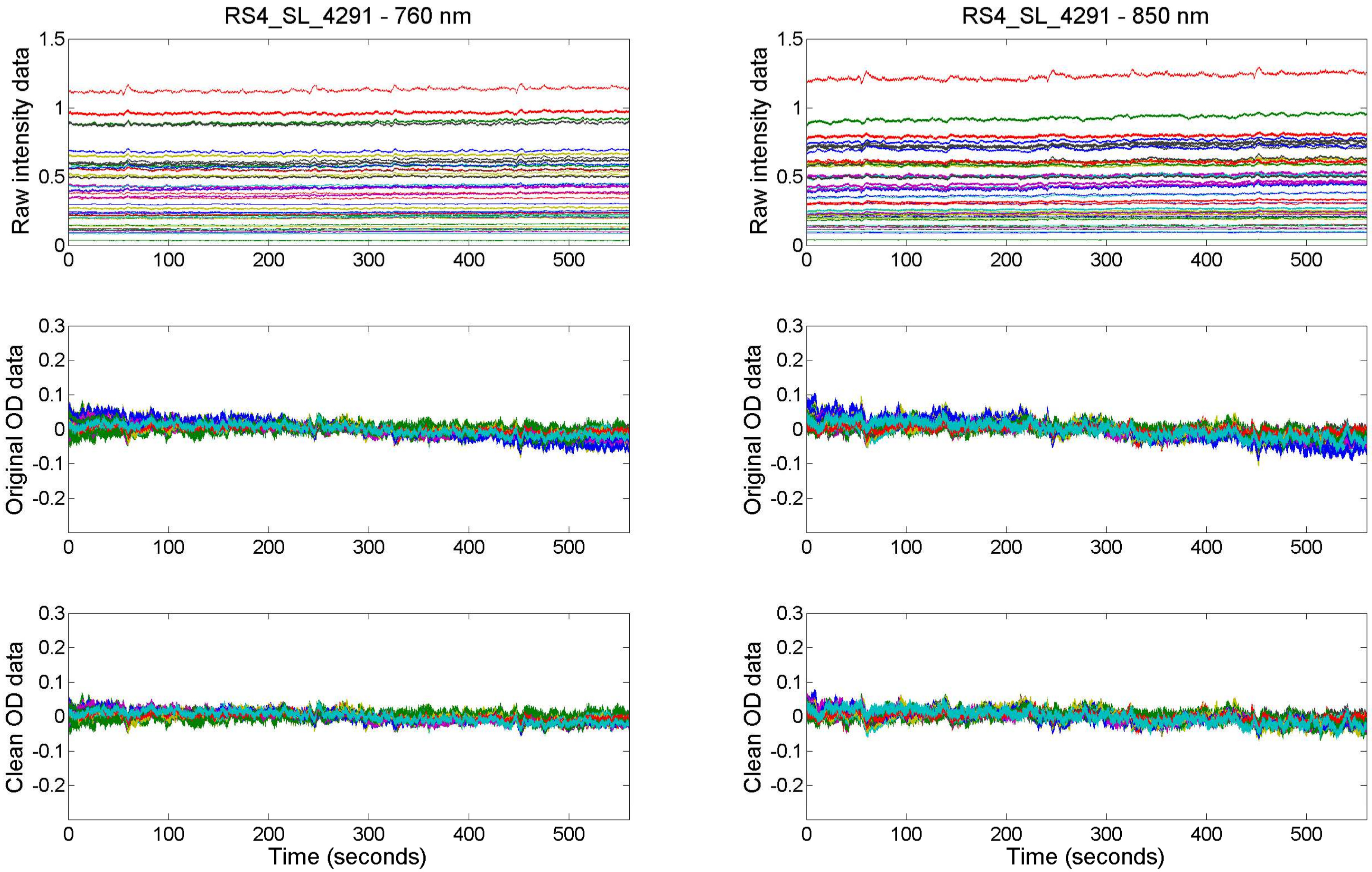

RS4_SL_4291 - 760 nm
RS4_SL_4291 - 850 nm
Raw intensity data
Original OD data
Clean OD data
Time (seconds)

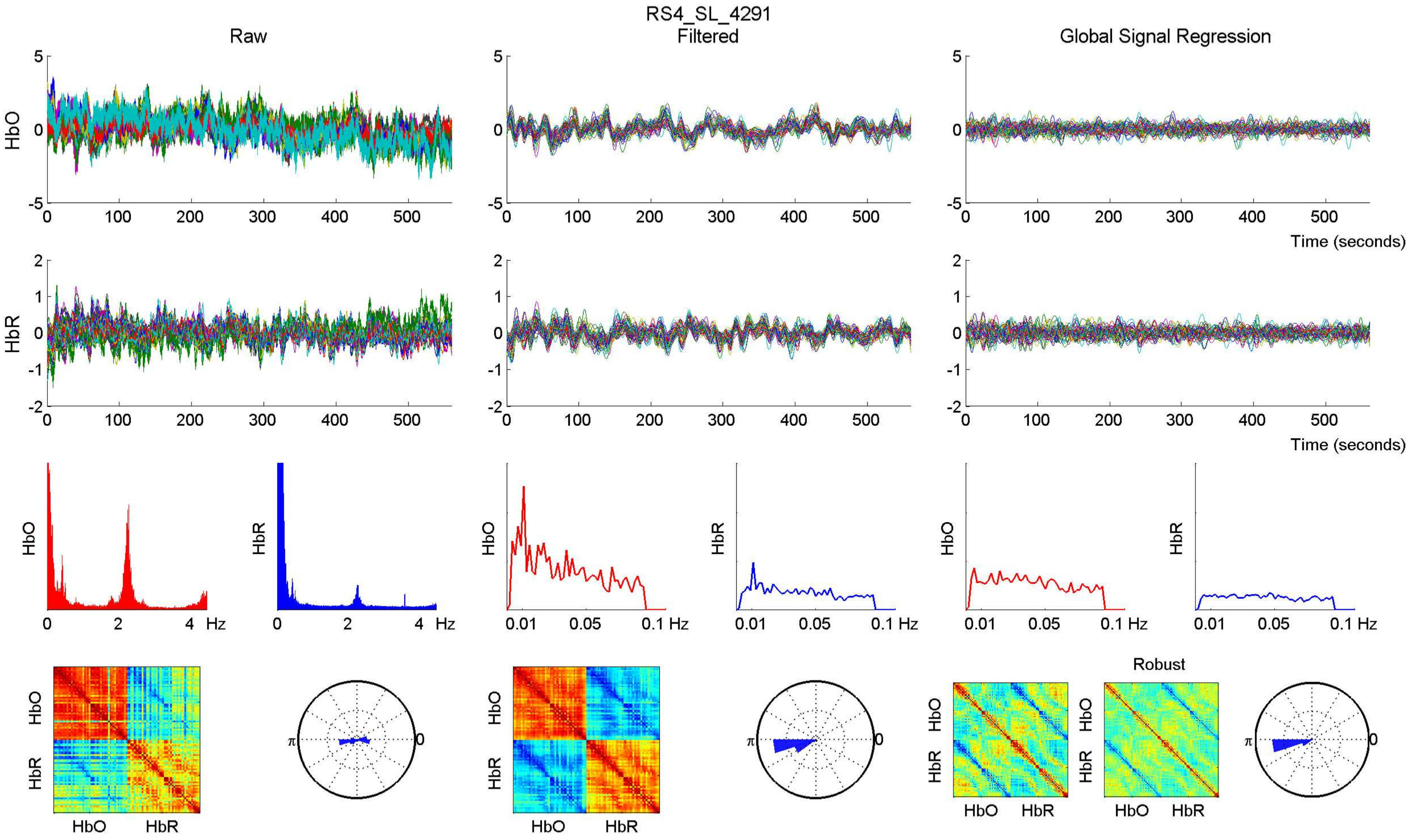

RS4_SL_4291
Raw
Filtered
Global Signal Regression
HbO
HbR
Time (seconds)
Hz
Robust
HbO
HbR

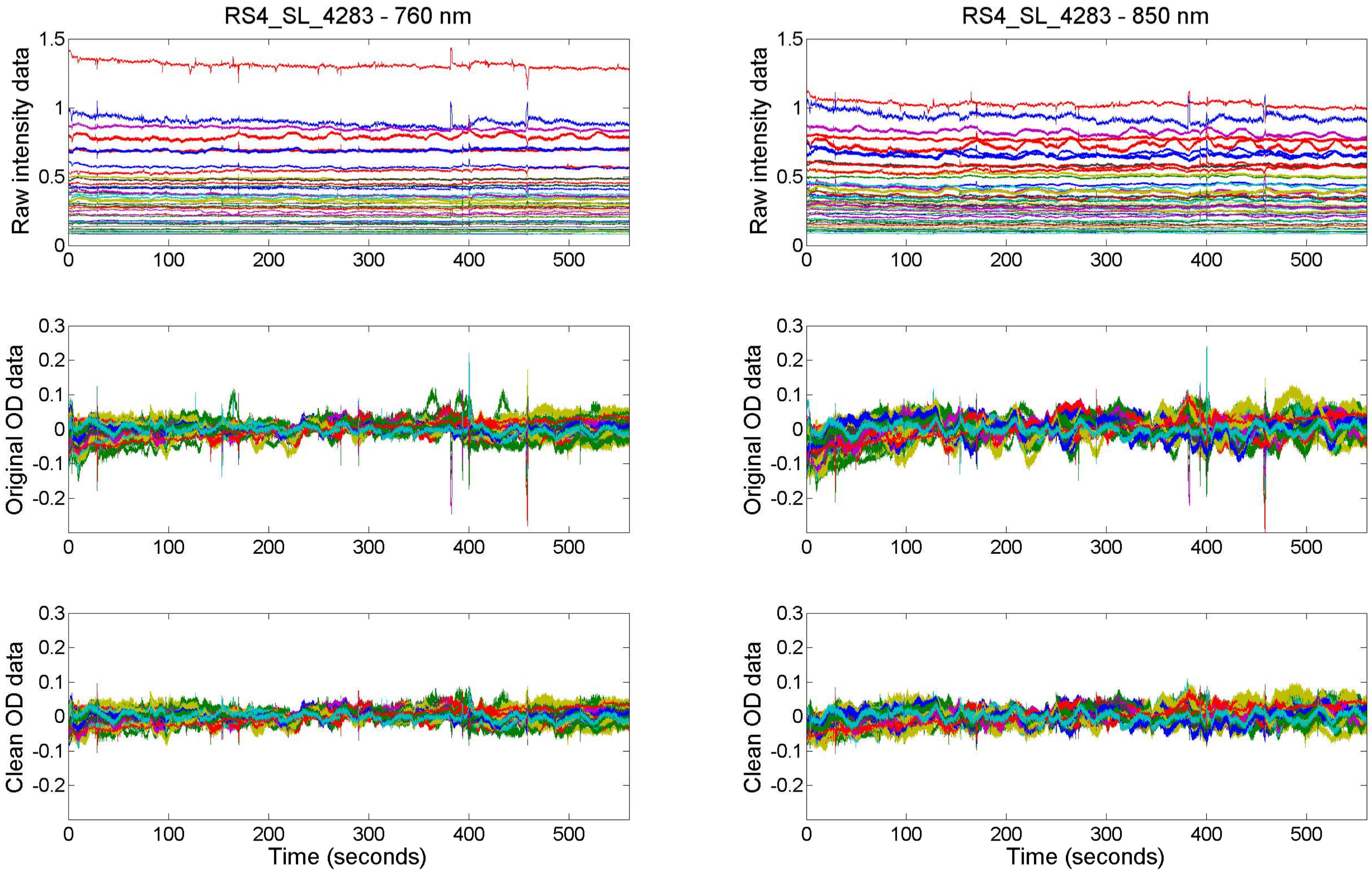
RS4_SL_4283 - 760 nm
RS4_SL_4283 - 850 nm
Raw intensity data
Original OD data
Clean OD data
Time (seconds)

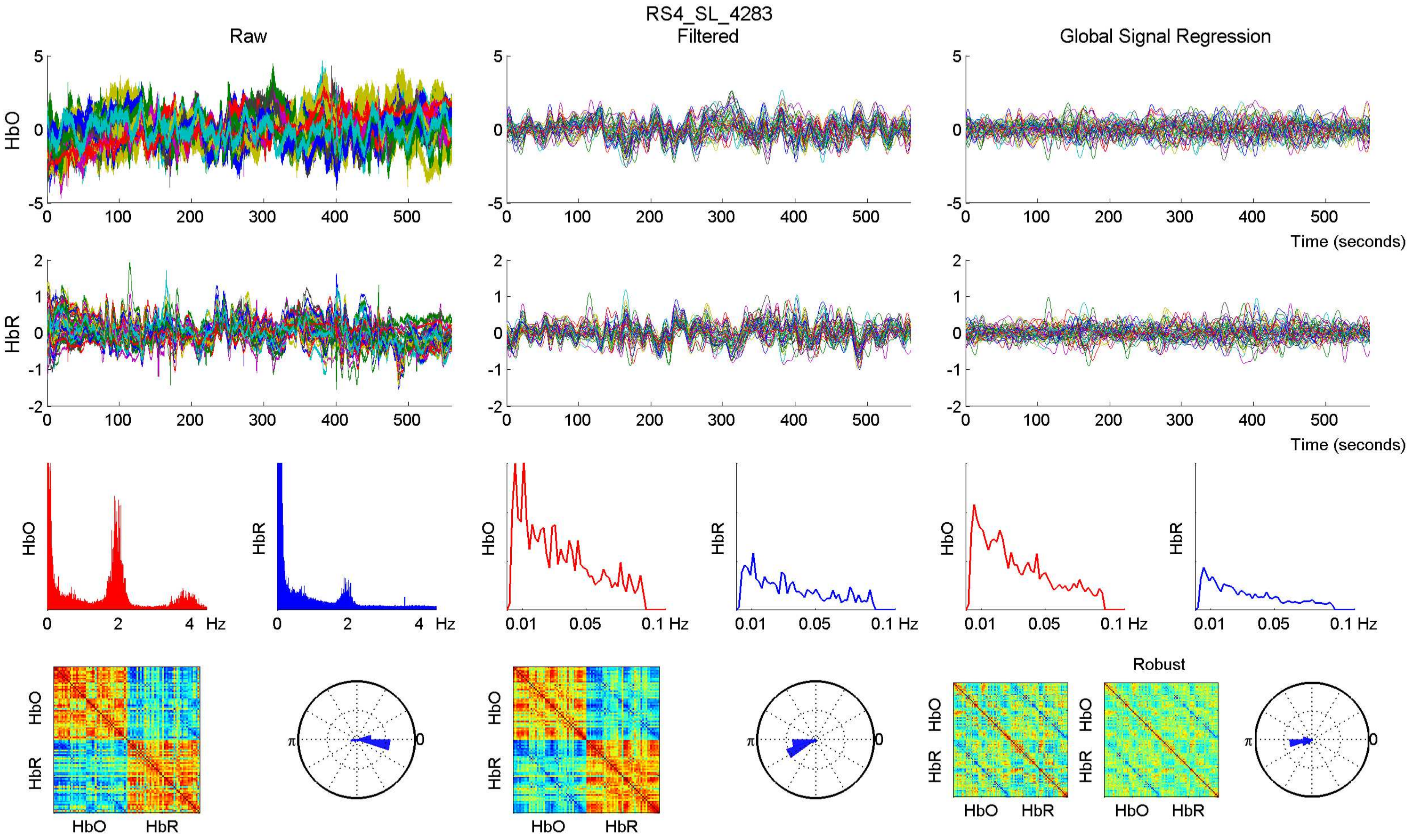

RS4_SL_4283
Raw
Filtered
Global Signal Regression
HbO
HbR
Time (seconds)
Hz
Robust

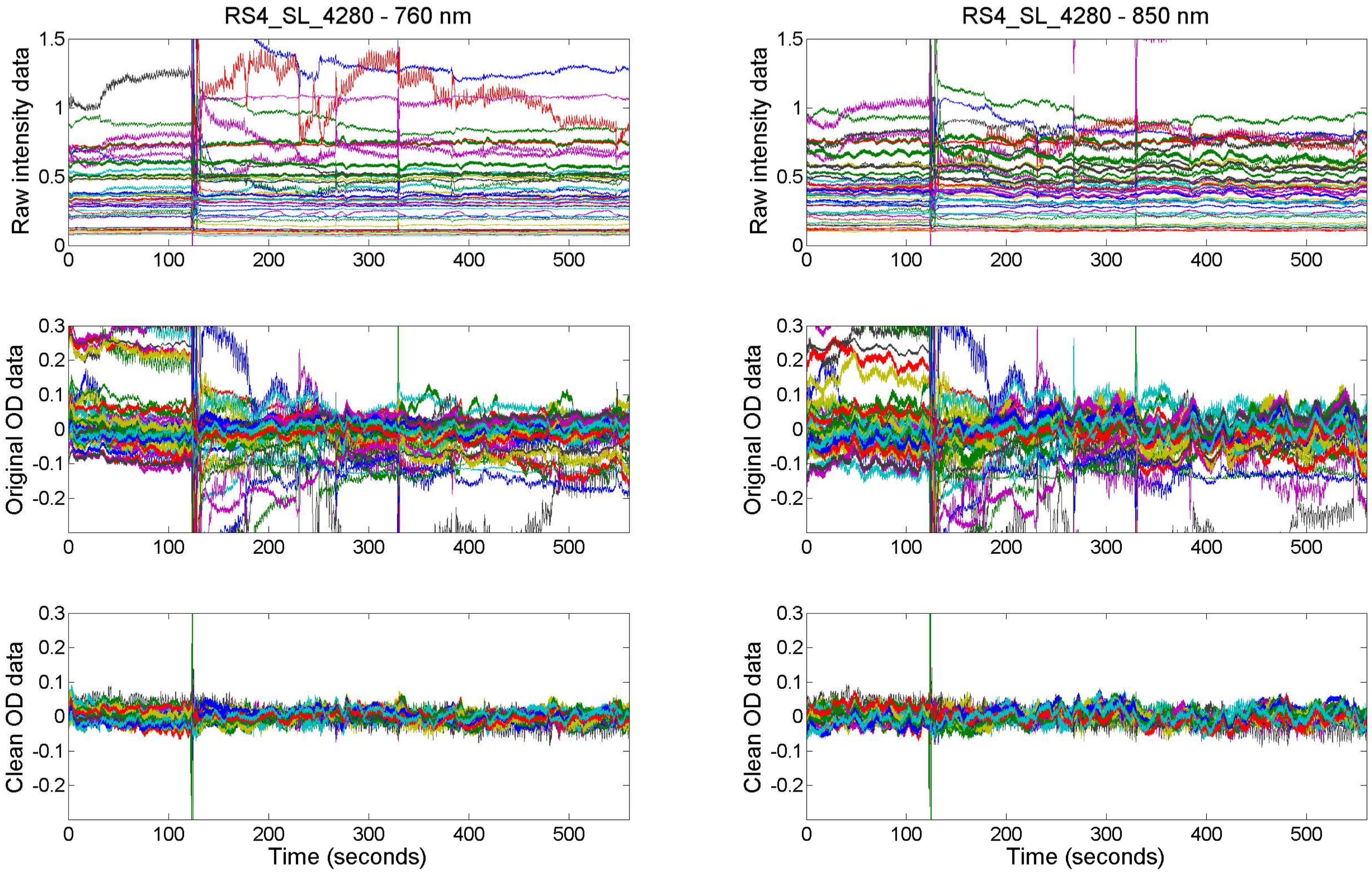
RS4_SL_4280 - 760 nm
RS4_SL_4280 - 850 nm
Raw intensity data
Original OD data
Clean OD data
Time (seconds)

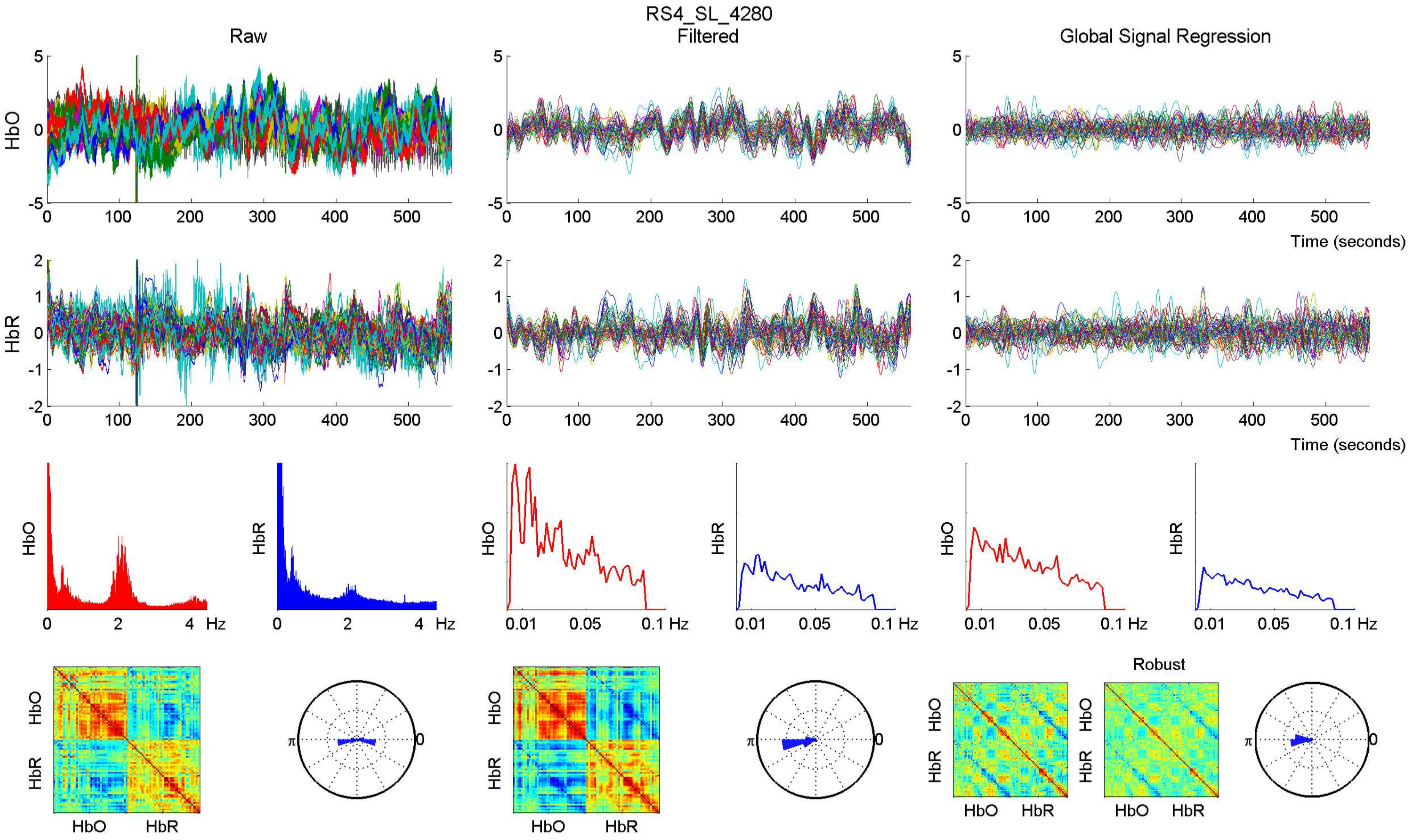

RS4_SL_4280
Raw
Filtered
Global Signal Regression
HbO
HbR
Time (seconds)
Hz
Robust

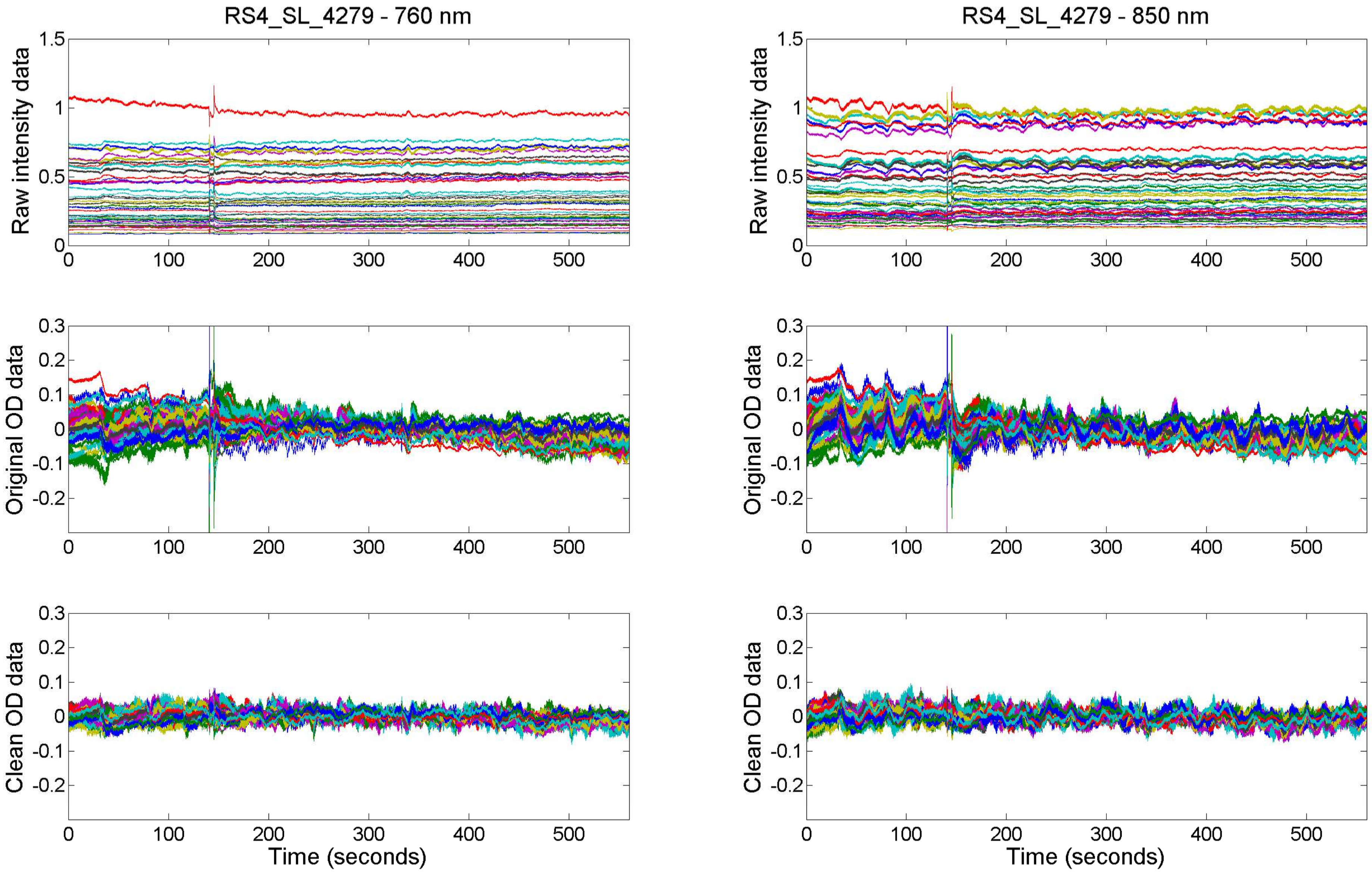

RS4_SL_4279 - 760 nm
RS4_SL_4279 - 850 nm
Raw intensity data
Original OD data
Clean OD data
Time (seconds)

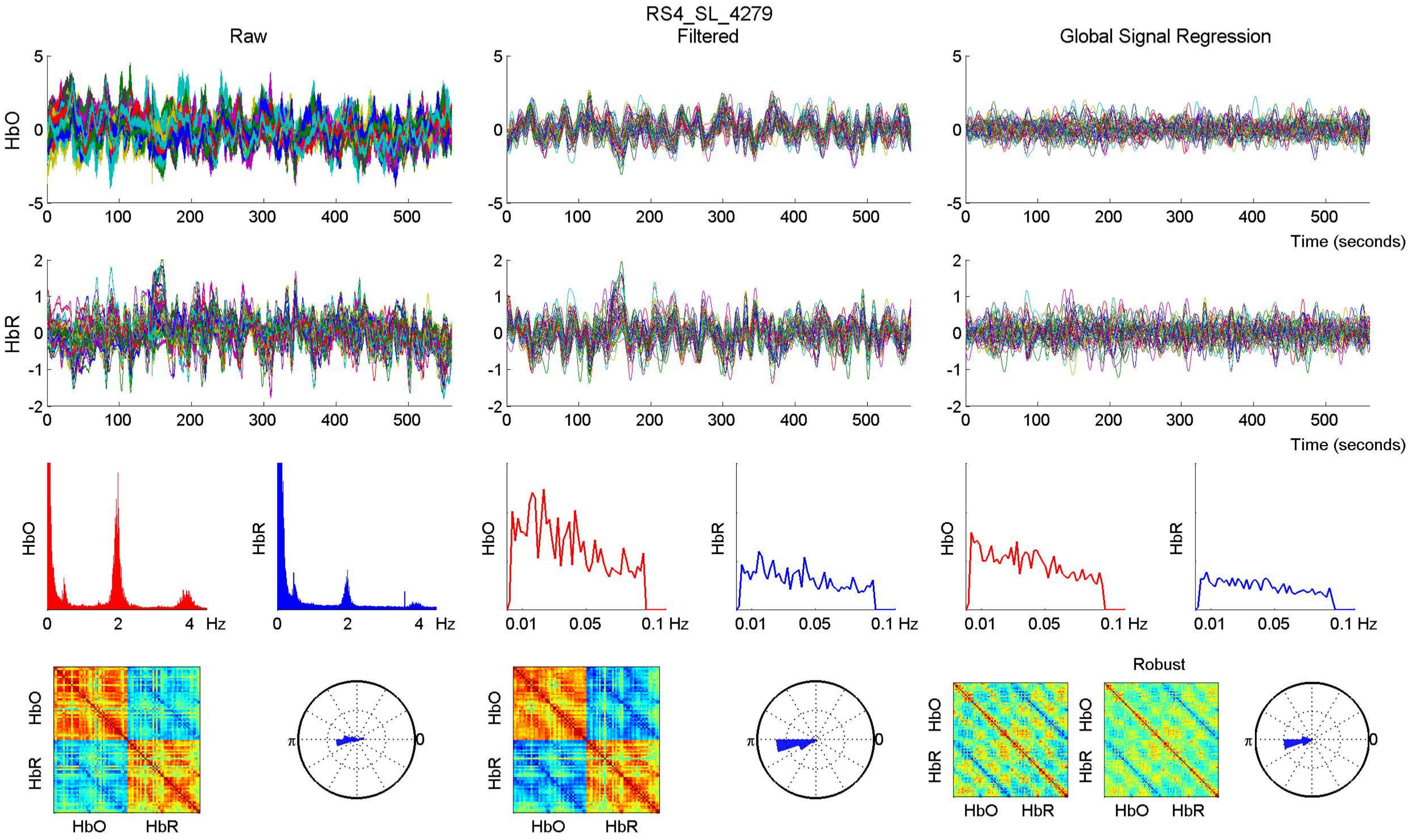

RS4_SL_4279
Raw
Filtered
Global Signal Regression
HbO
HbR
Time (seconds)
Hz
Robust

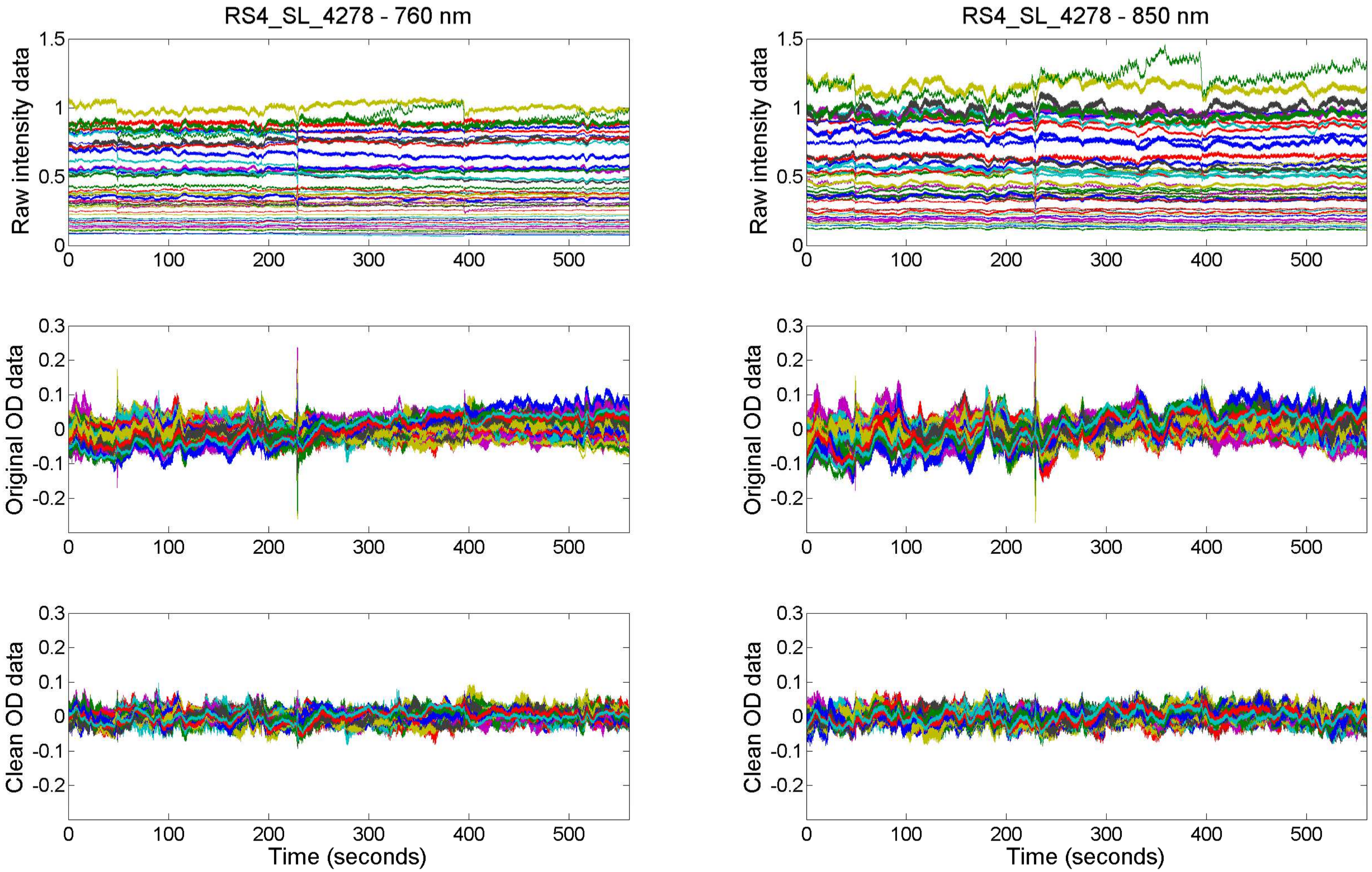

RS4_SL_4278 - 760 nm
RS4_SL_4278 - 850 nm
Raw intensity data
Original OD data
Clean OD data
Time (seconds)

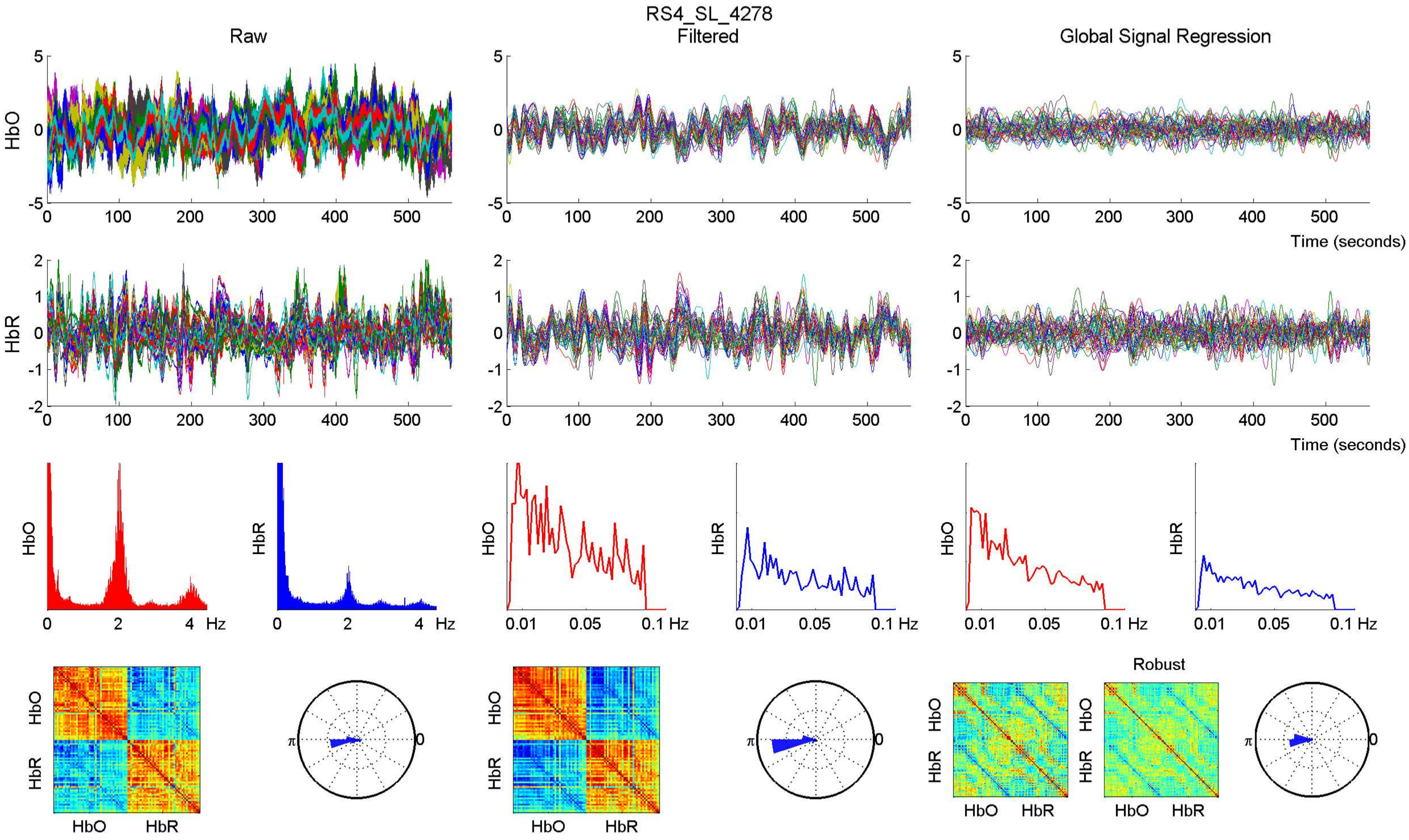
RS4_SL_4278
Raw
Filtered
Global Signal Regression
HbO
HbR
Time (seconds)
Hz
Robust

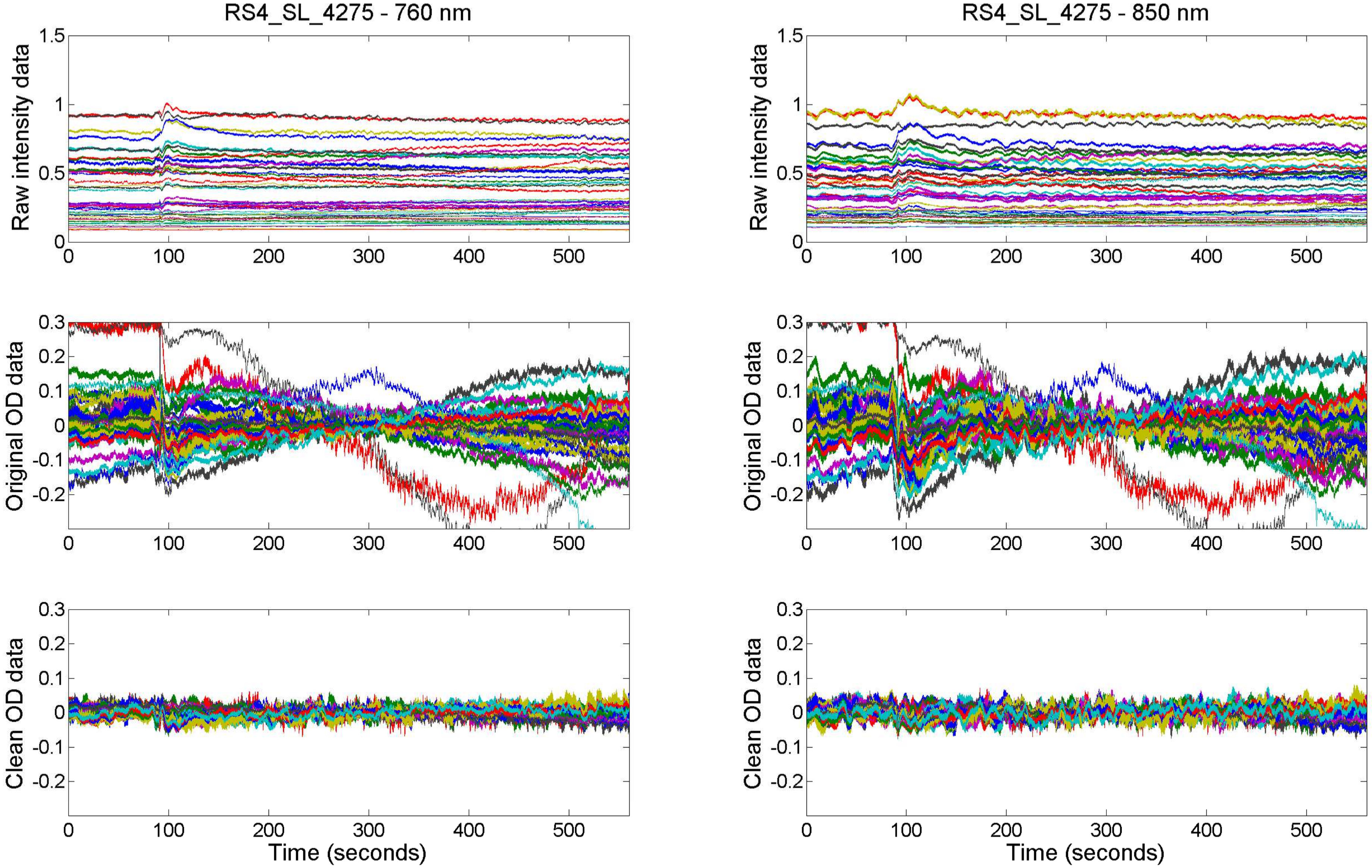

RS4_SL_4275 - 760 nm
RS4_SL_4275 - 850 nm
Raw intensity data
Original OD data
Clean OD data
Time (seconds)

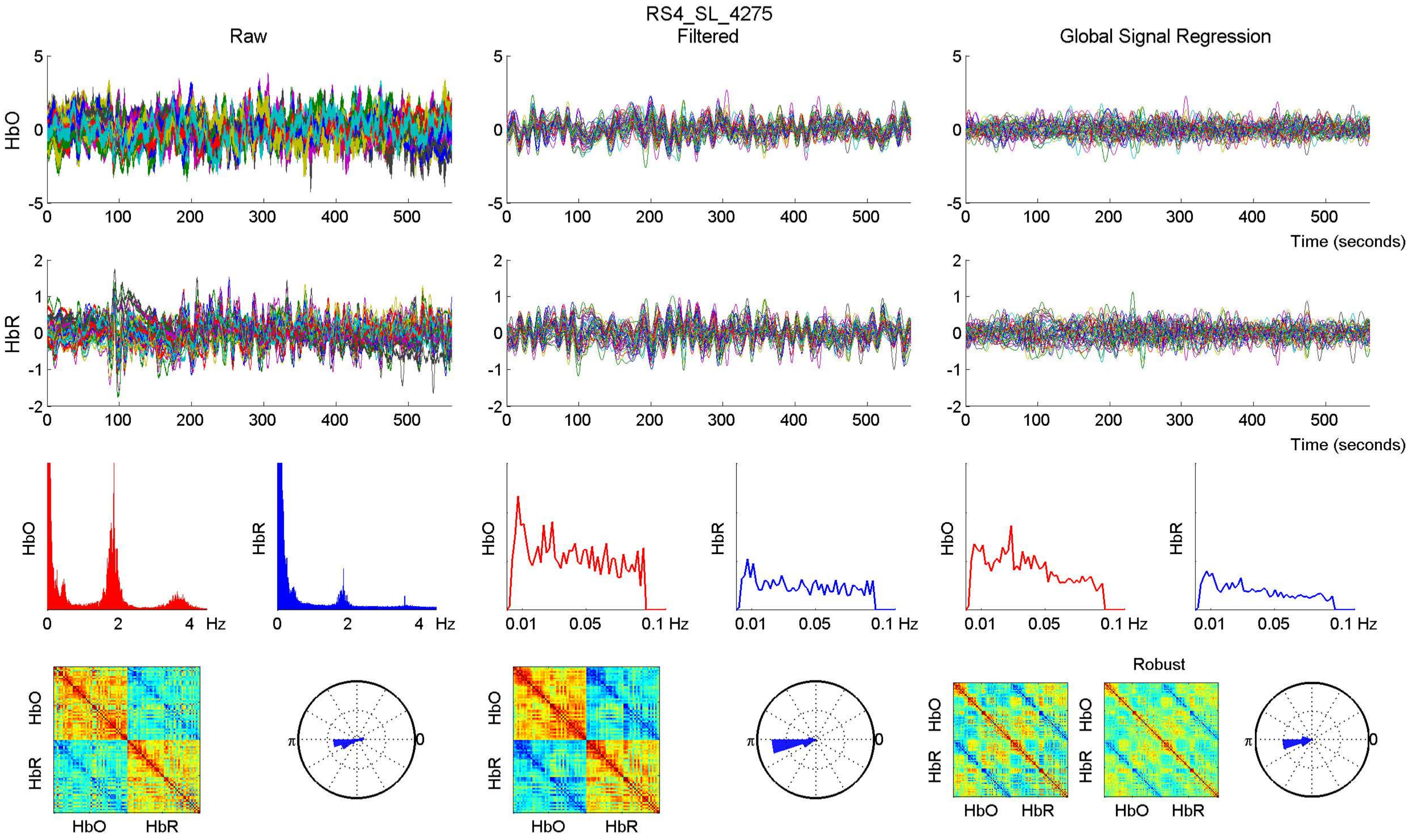
RS4_SL_4275
Raw
Filtered
Global Signal Regression
HbO
HbR
Time (seconds)
Hz
Robust
π
0

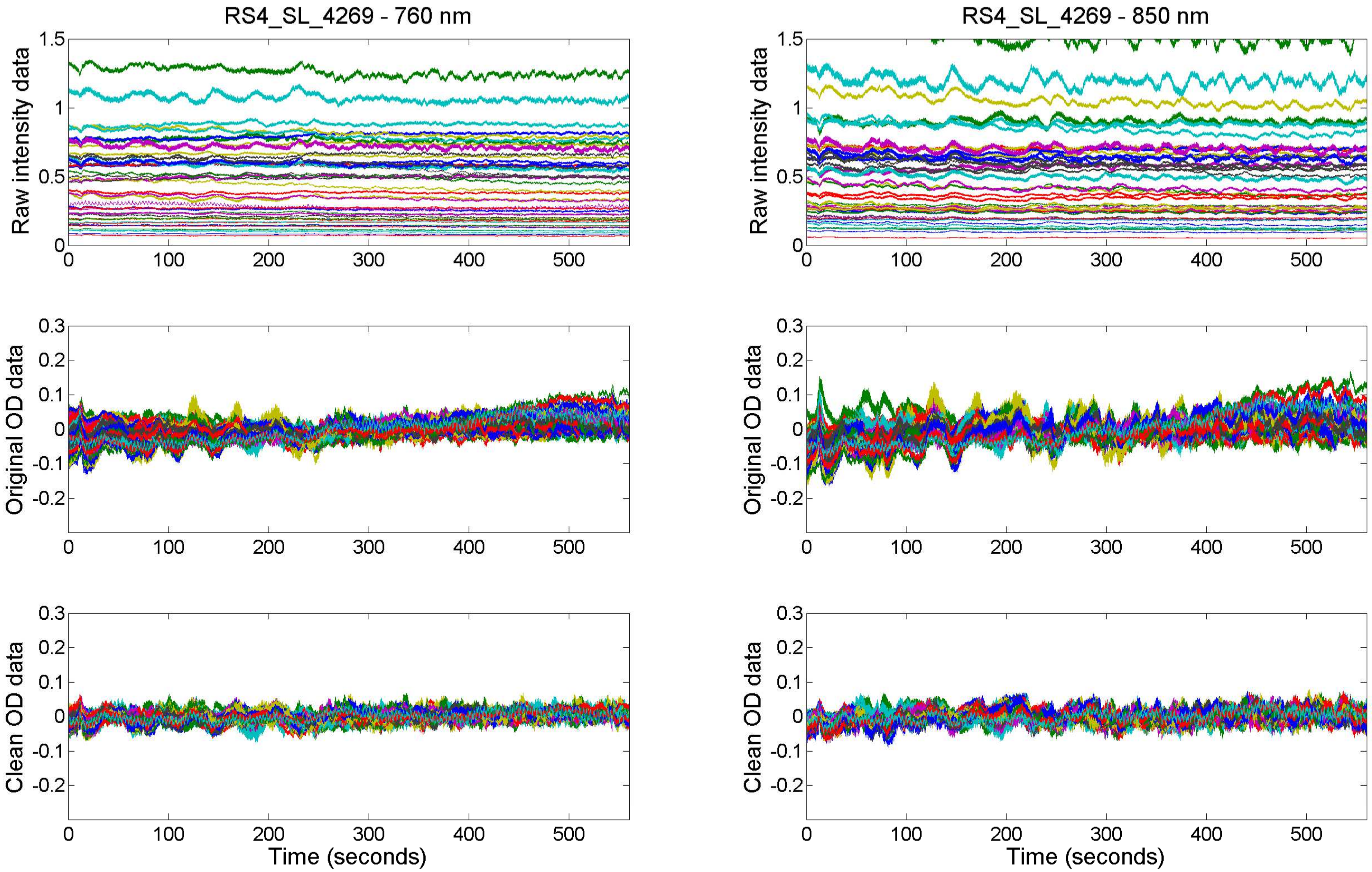

RS4_SL_4269 - 760 nm
RS4_SL_4269 - 850 nm
Raw intensity data
Original OD data
Clean OD data
Time (seconds)

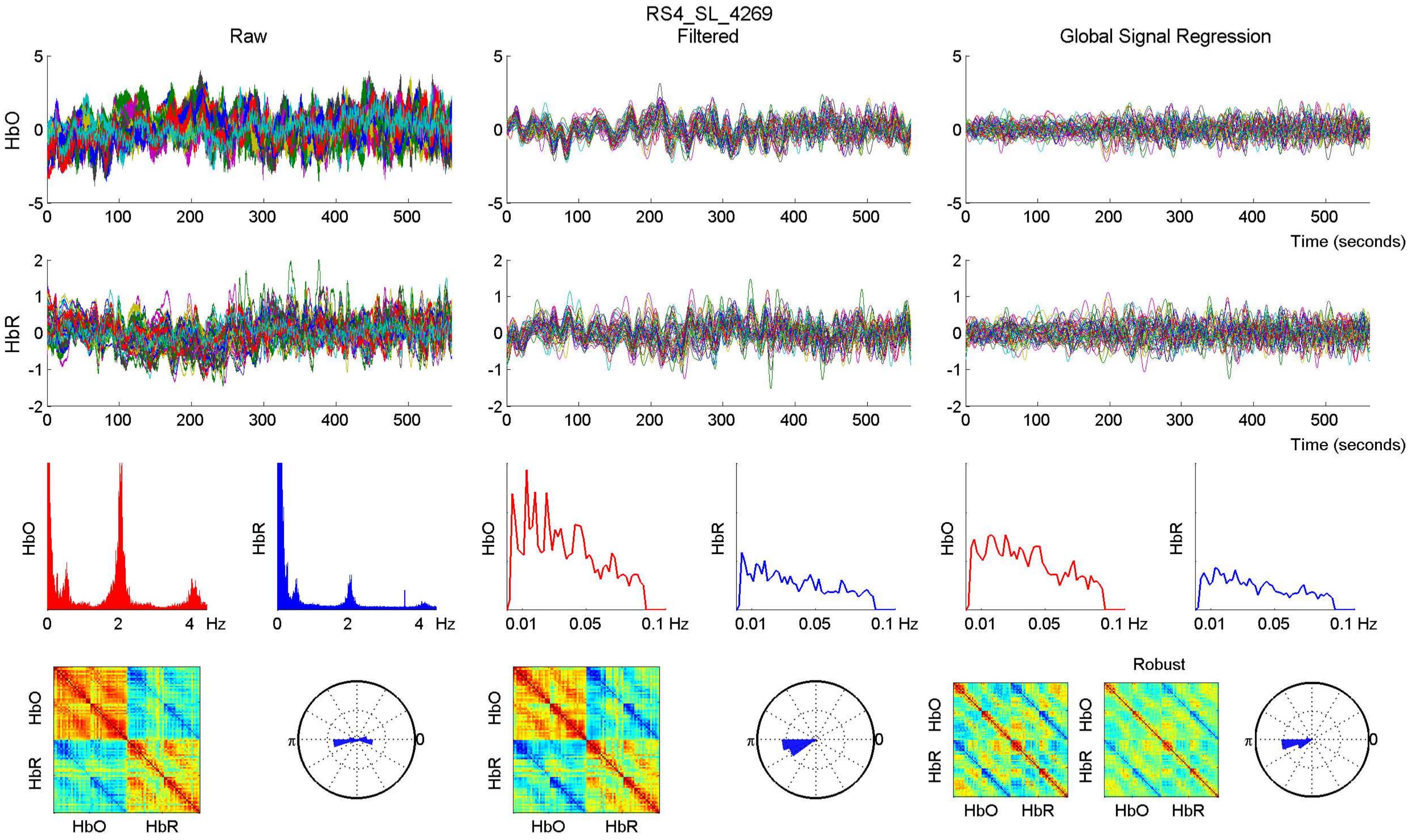
RS4_SL_4269
Raw
Filtered
Global Signal Regression
HbO
HbR
Time (seconds)
Hz
Robust

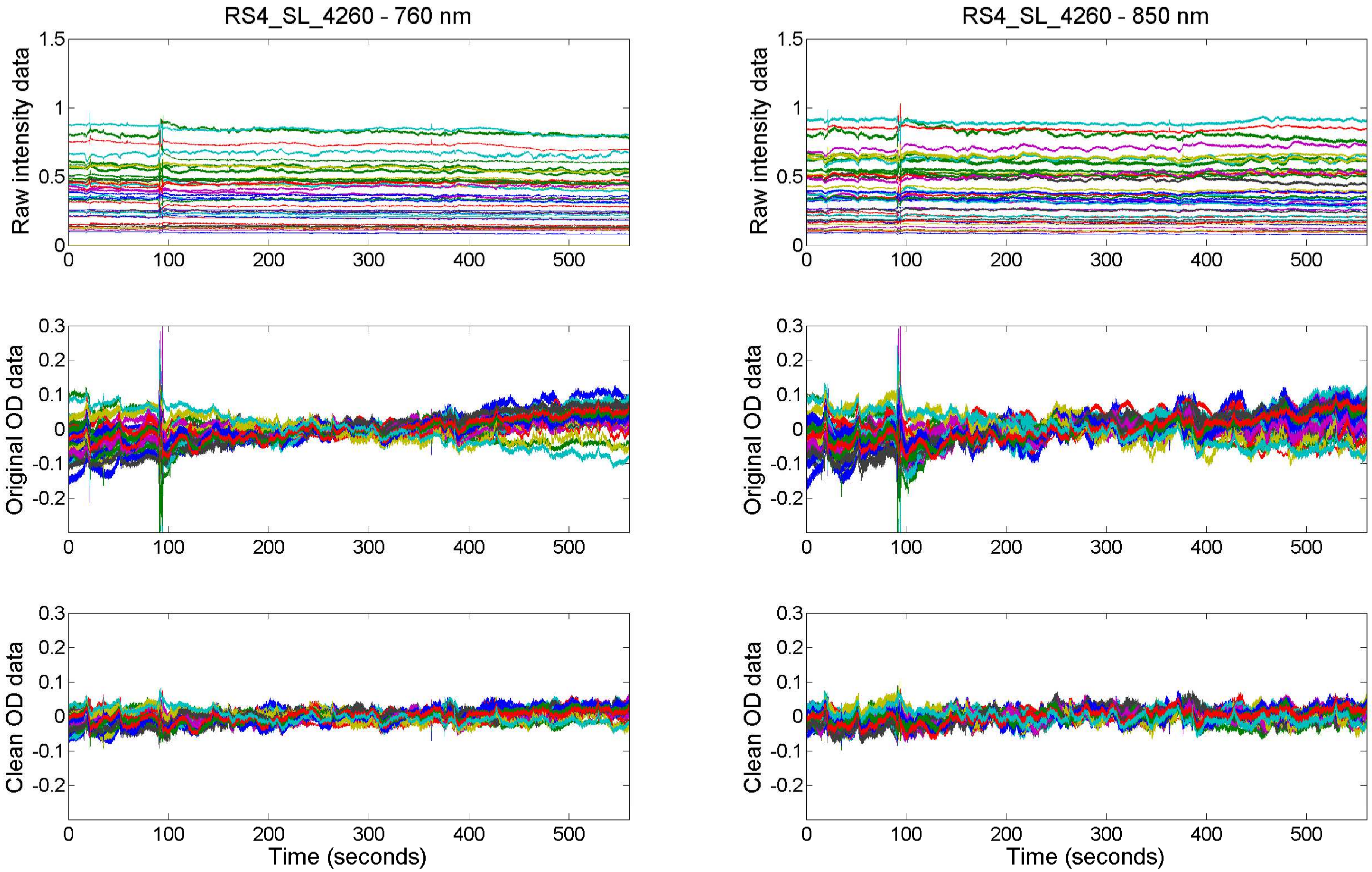

RS4_SL_4260 - 760 nm
RS4_SL_4260 - 850 nm
Raw intensity data
Original OD data
Clean OD data
Time (seconds)

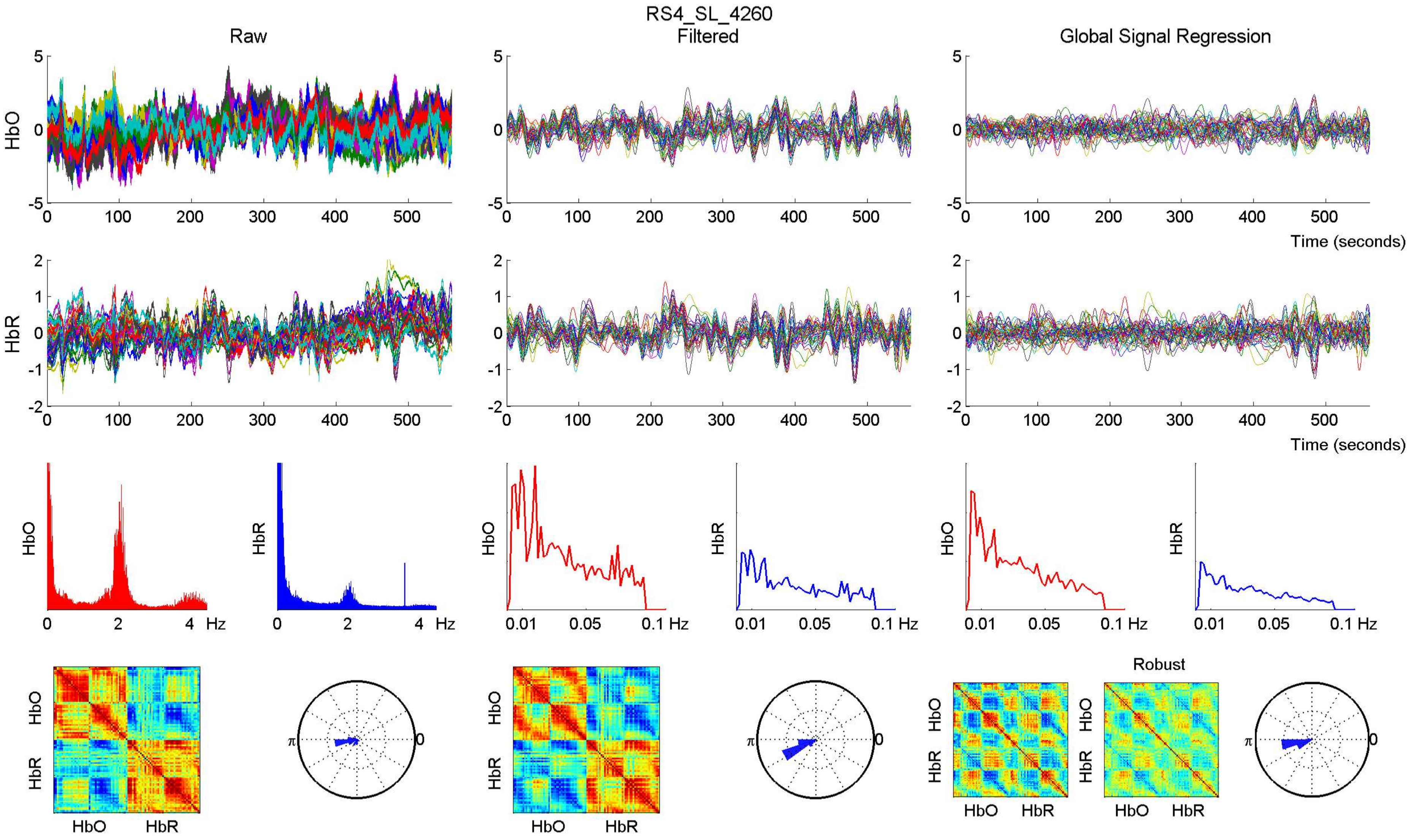

RS4_SL_4260
Raw
Filtered
Global Signal Regression
HbO
HbR
Time (seconds)
Hz
Robust
HbO
HbR

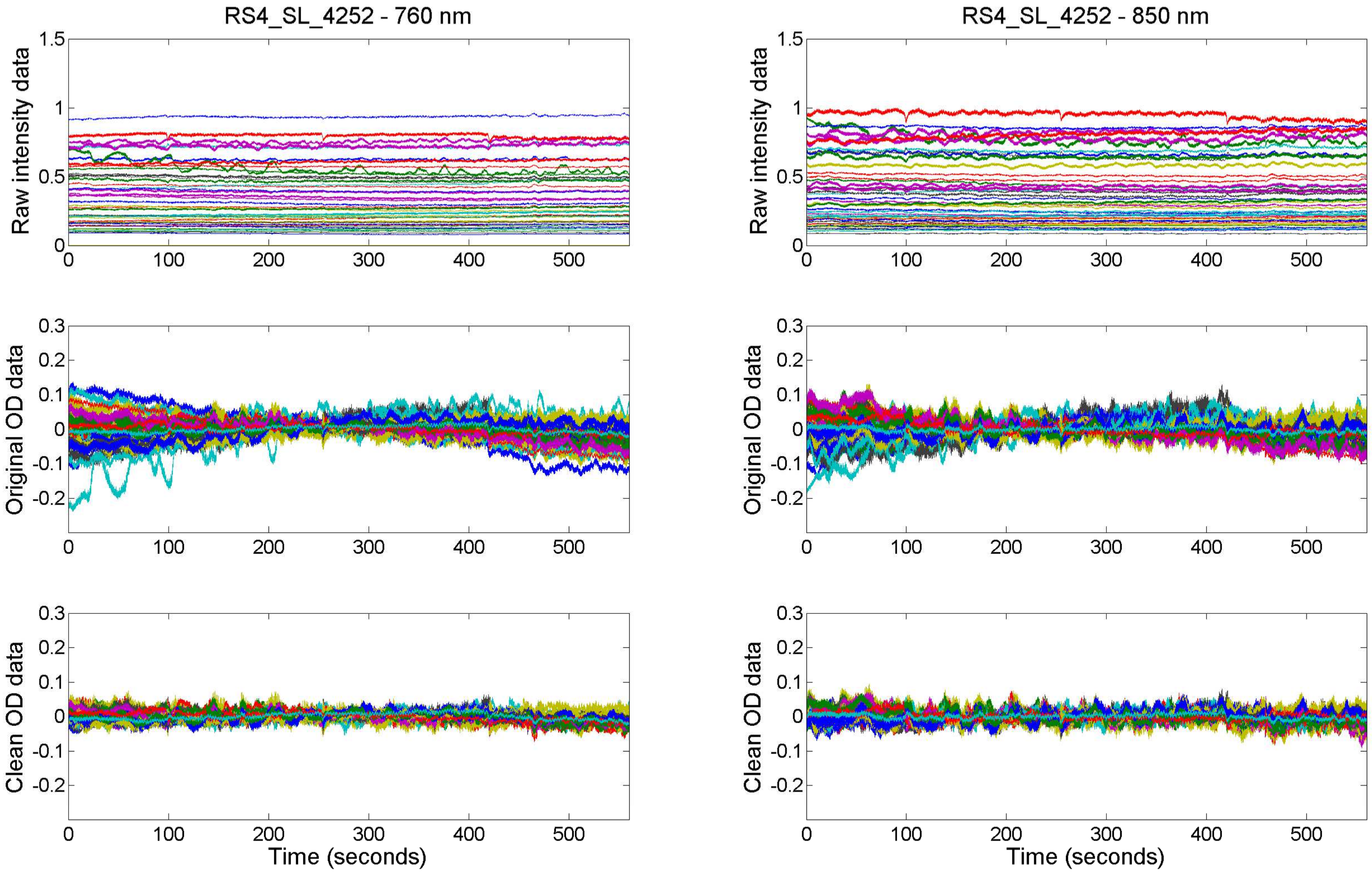

RS4_SL_4252 - 760 nm
RS4_SL_4252 - 850 nm
Raw intensity data
Original OD data
Clean OD data
Time (seconds)

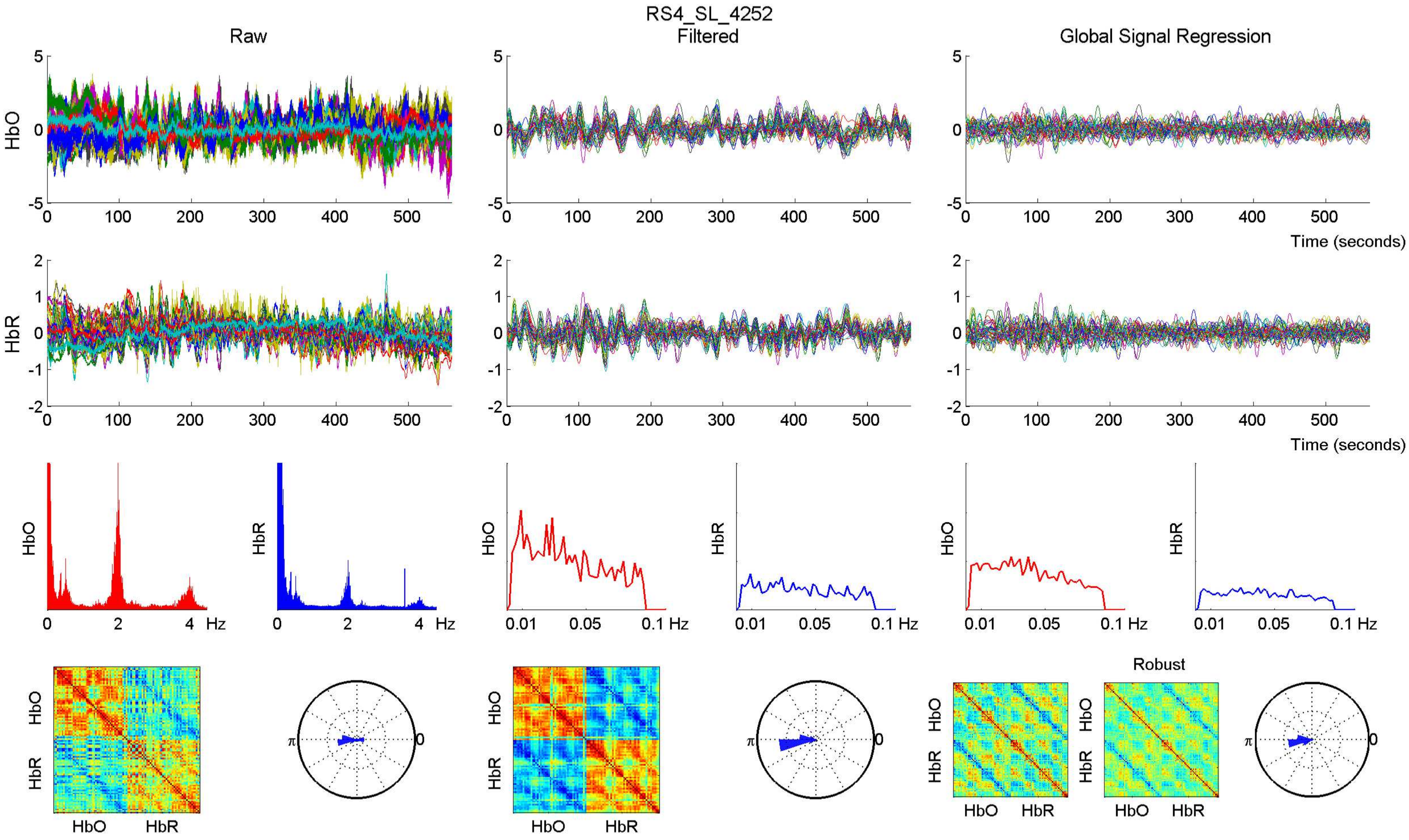

RS4_SL_4252
Raw
Filtered
Global Signal Regression
HbO
HbR
Time (seconds)
Hz
Robust

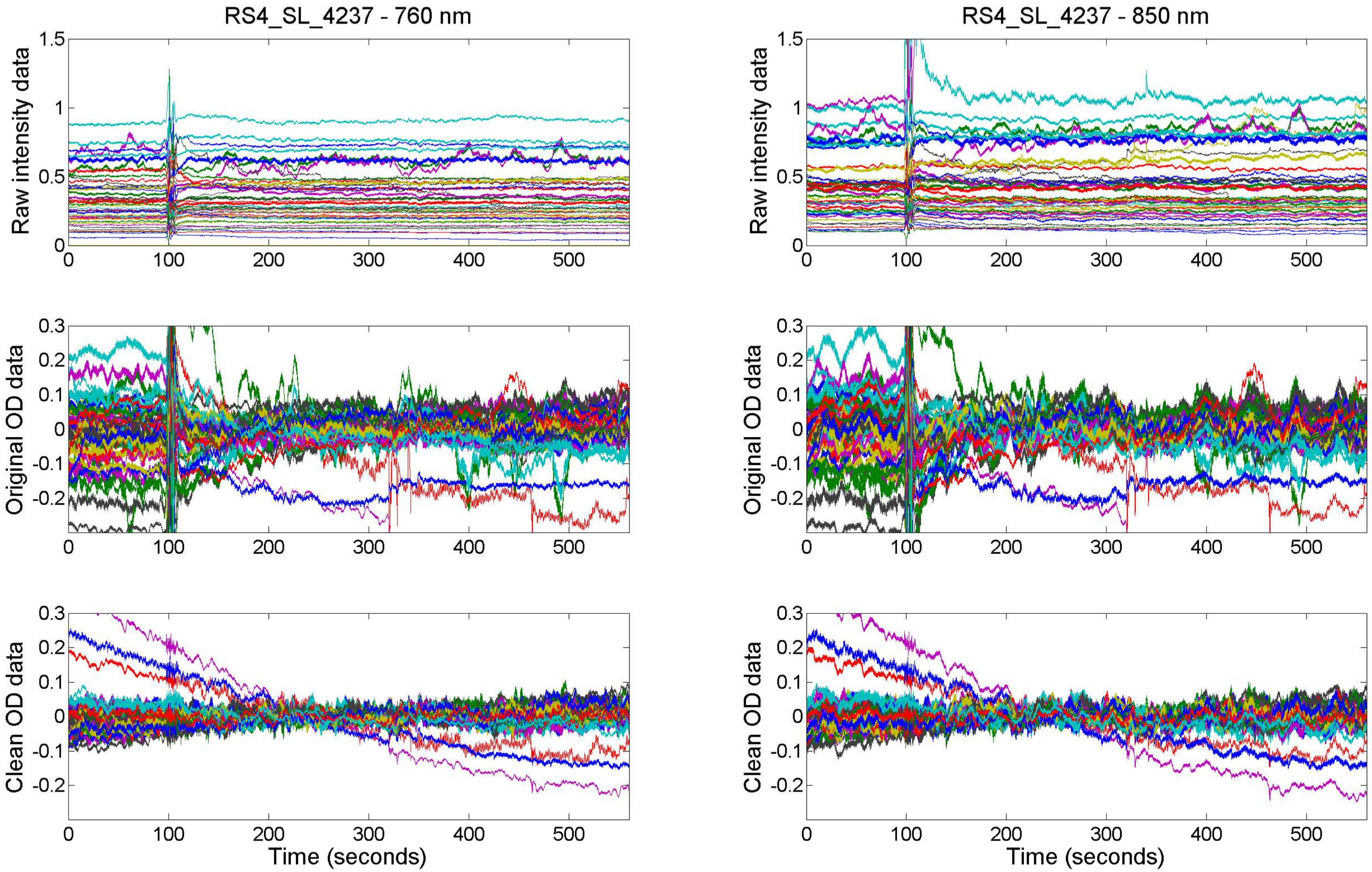

RS4_SL_4237 - 760 nm
RS4_SL_4237 - 850 nm
Raw intensity data
Original OD data
Clean OD data
Time (seconds)

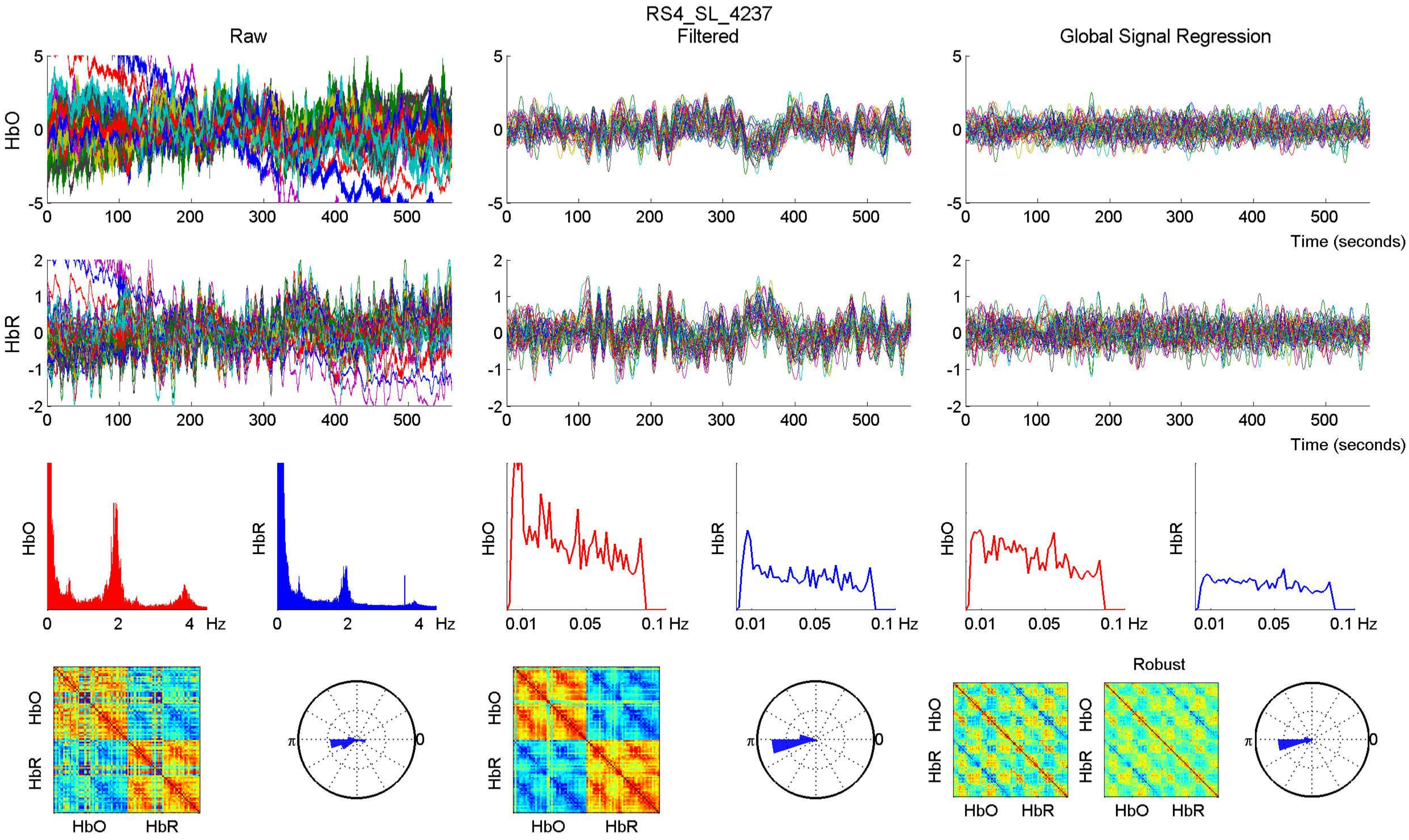

RS4_SL_4237
Raw
Filtered
Global Signal Regression
HbO
HbR
Time (seconds)
Hz
0.01
0.05
0.1 Hz
Robust
π
0

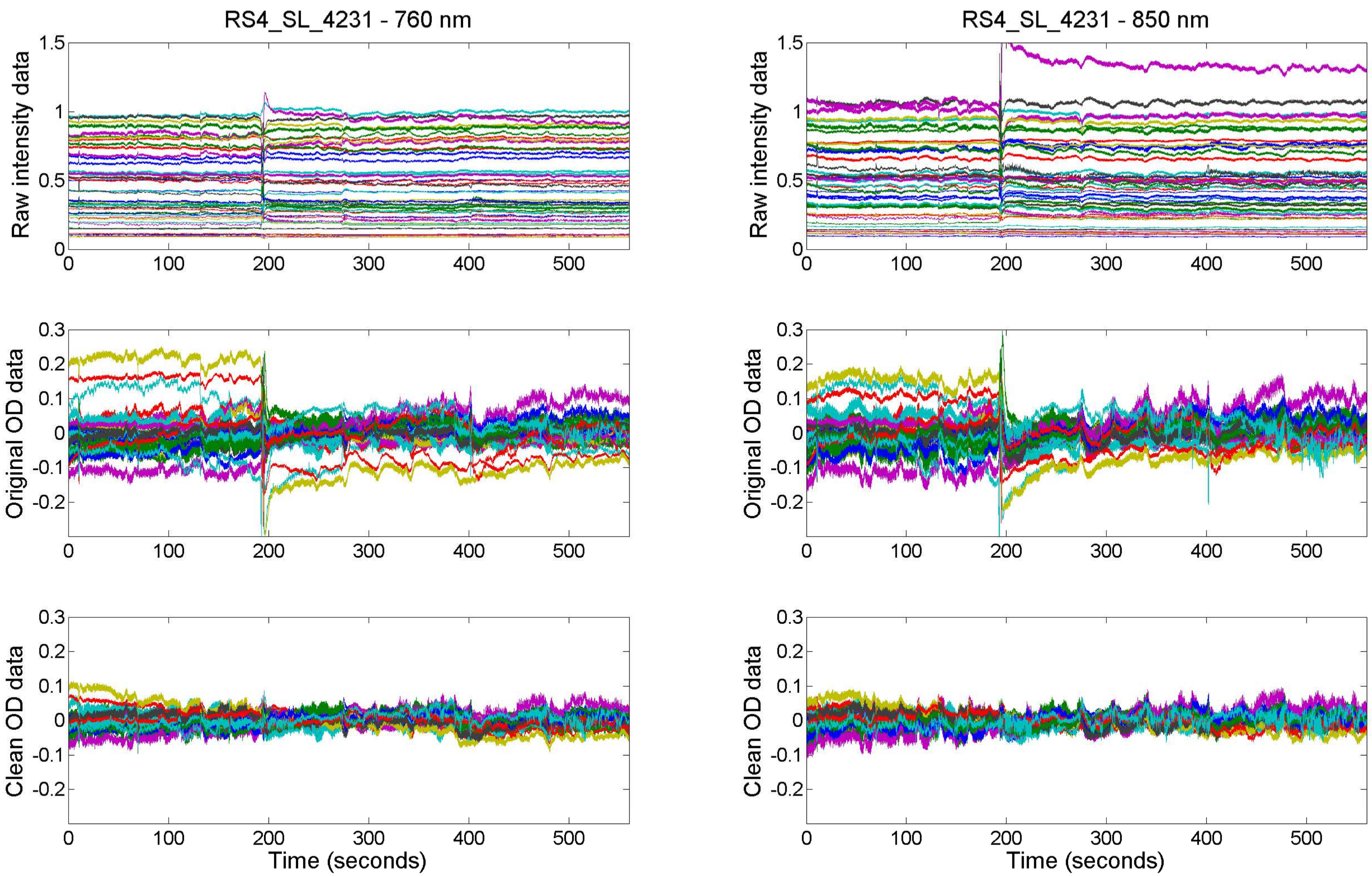
RS4_SL_4231 - 760 nm
RS4_SL_4231 - 850 nm
Raw intensity data
Original OD data
Clean OD data
Time (seconds)
0
100
200
300
400
500
1.5
1
0.5
0.3
0.2
0.1
-0.1
-0.2

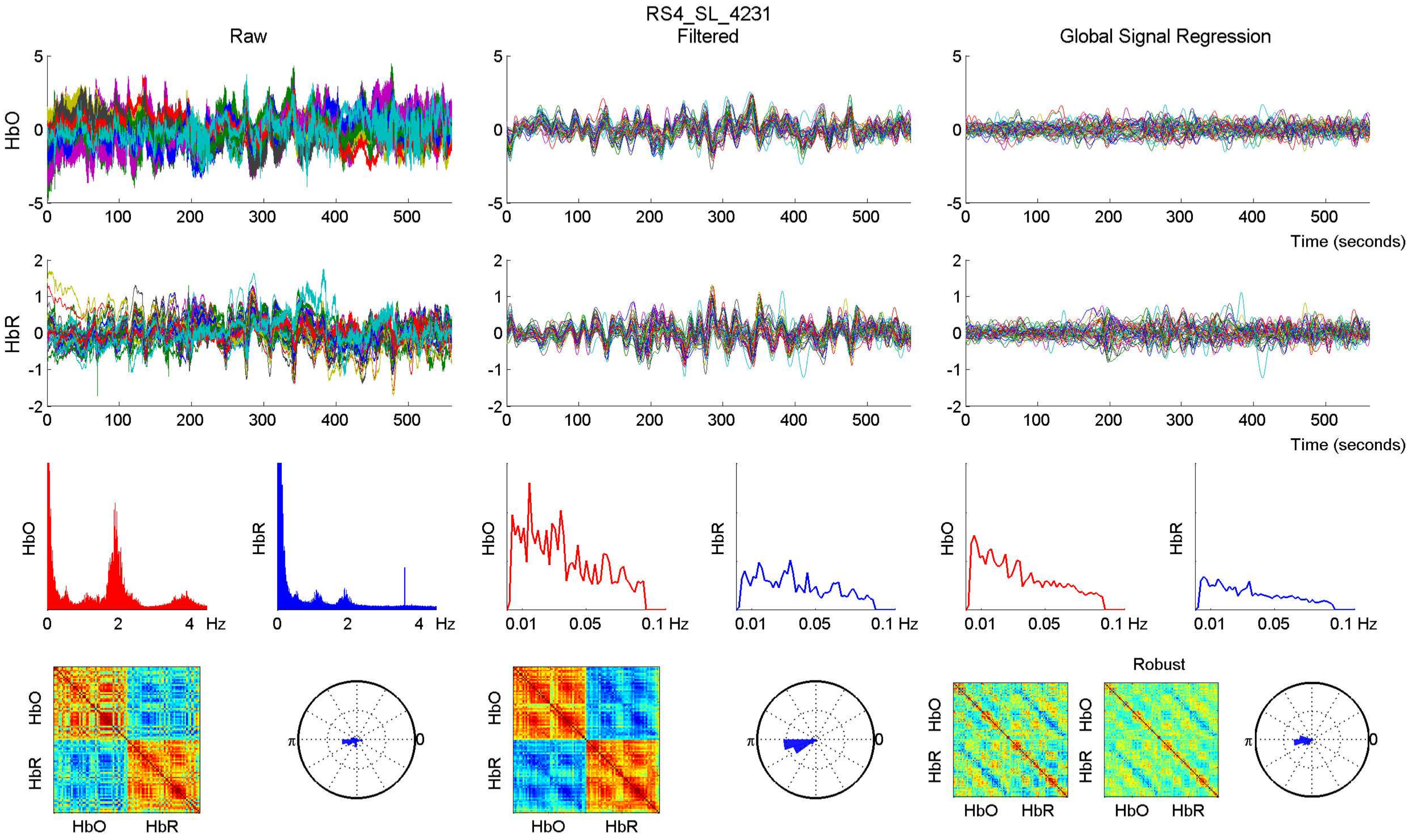

RS4_SL_4231
Raw
Filtered
Global Signal Regression
HbO
HbR
Time (seconds)
Hz
Robust
HbO HbR
π
0

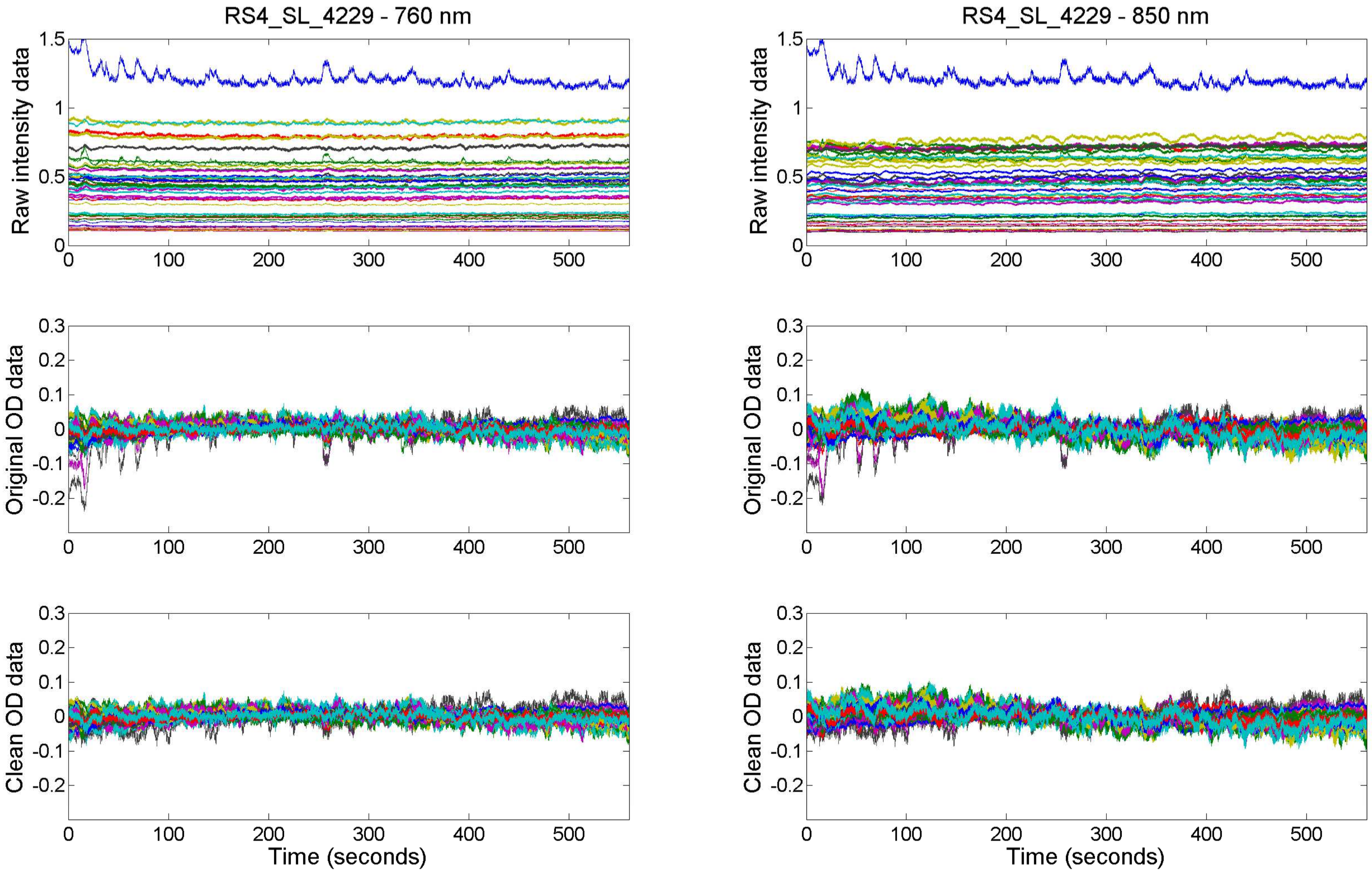
RS4_SL_4229 - 760 nm
RS4_SL_4229 - 850 nm
Raw intensity data
Original OD data
Clean OD data
Time (seconds)
0
100
200
300
400
500
1.5
1
0.5
0.3
0.2
0.1
-0.1
-0.2

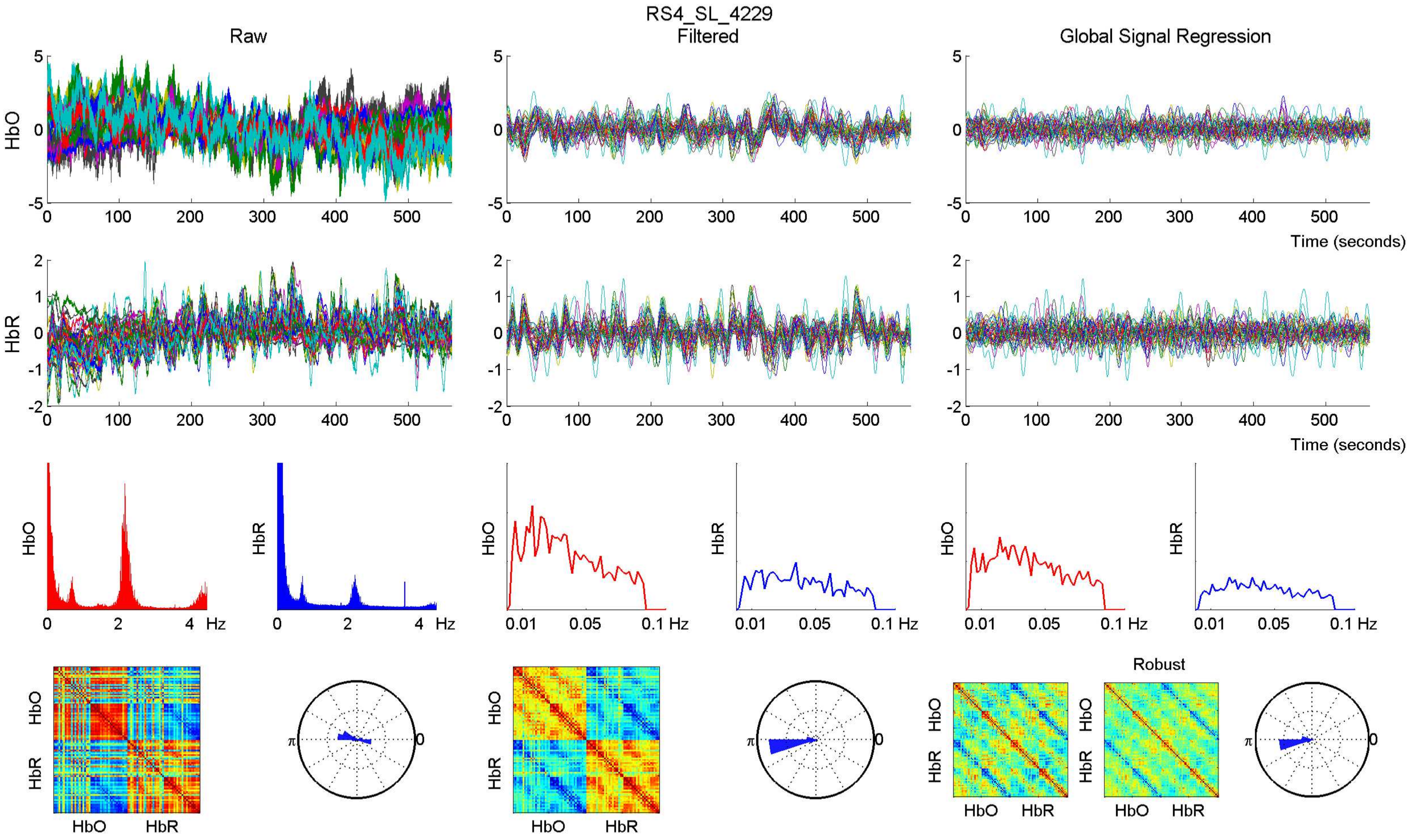
RS4_SL_4229
Raw
Filtered
Global Signal Regression
HbO
HbR
Time (seconds)
Hz
Robust

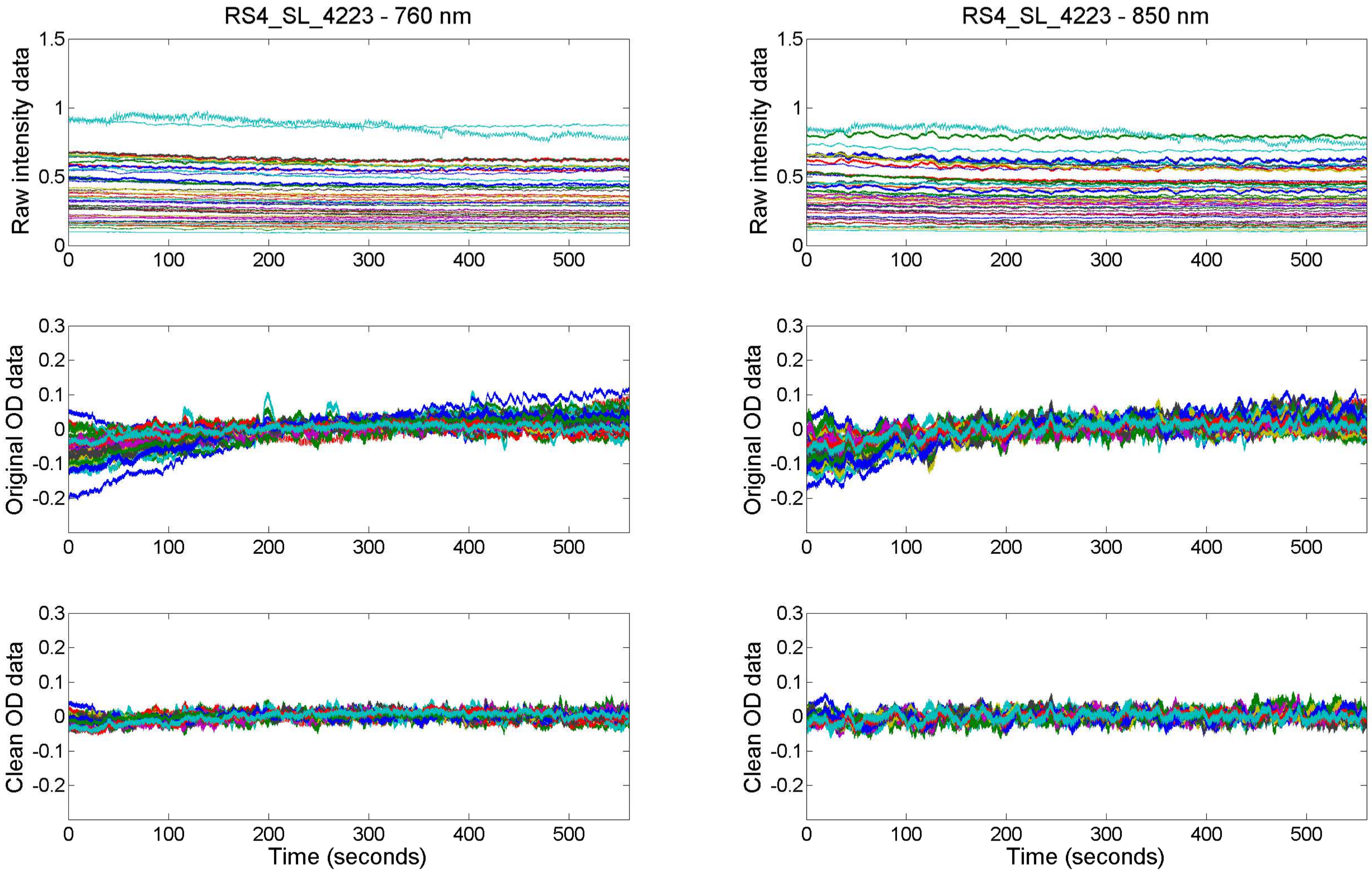
RS4_SL_4223 - 760 nm
RS4_SL_4223 - 850 nm
Raw intensity data
Original OD data
Clean OD data
Time (seconds)
1.5
1
0.5
0
0.3
0.2
0.1
-0.1
-0.2
100
200
300
400
500

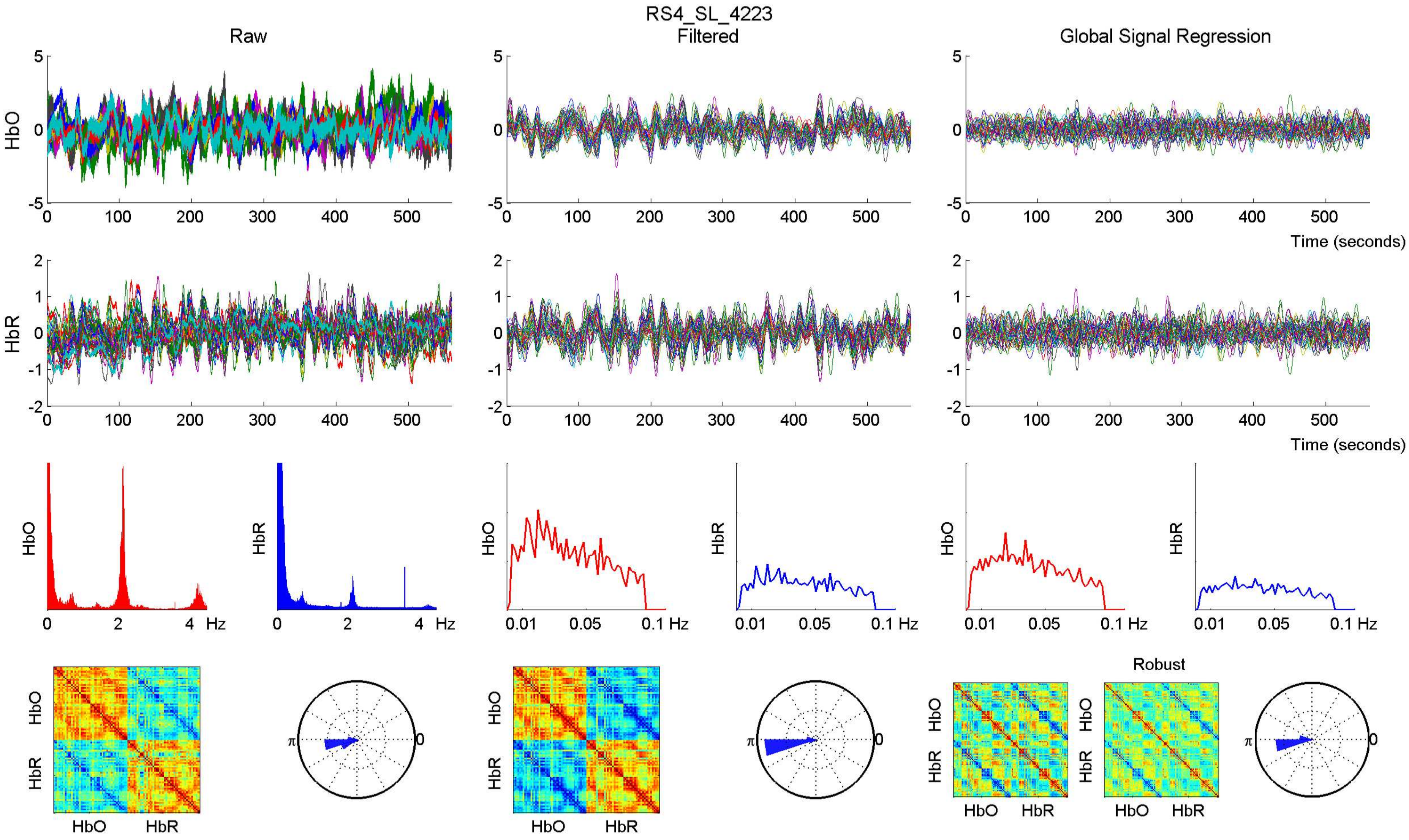

RS4_SL_4223
Raw
Filtered
Global Signal Regression
HbO
HbR
Time (seconds)
Hz
Robust

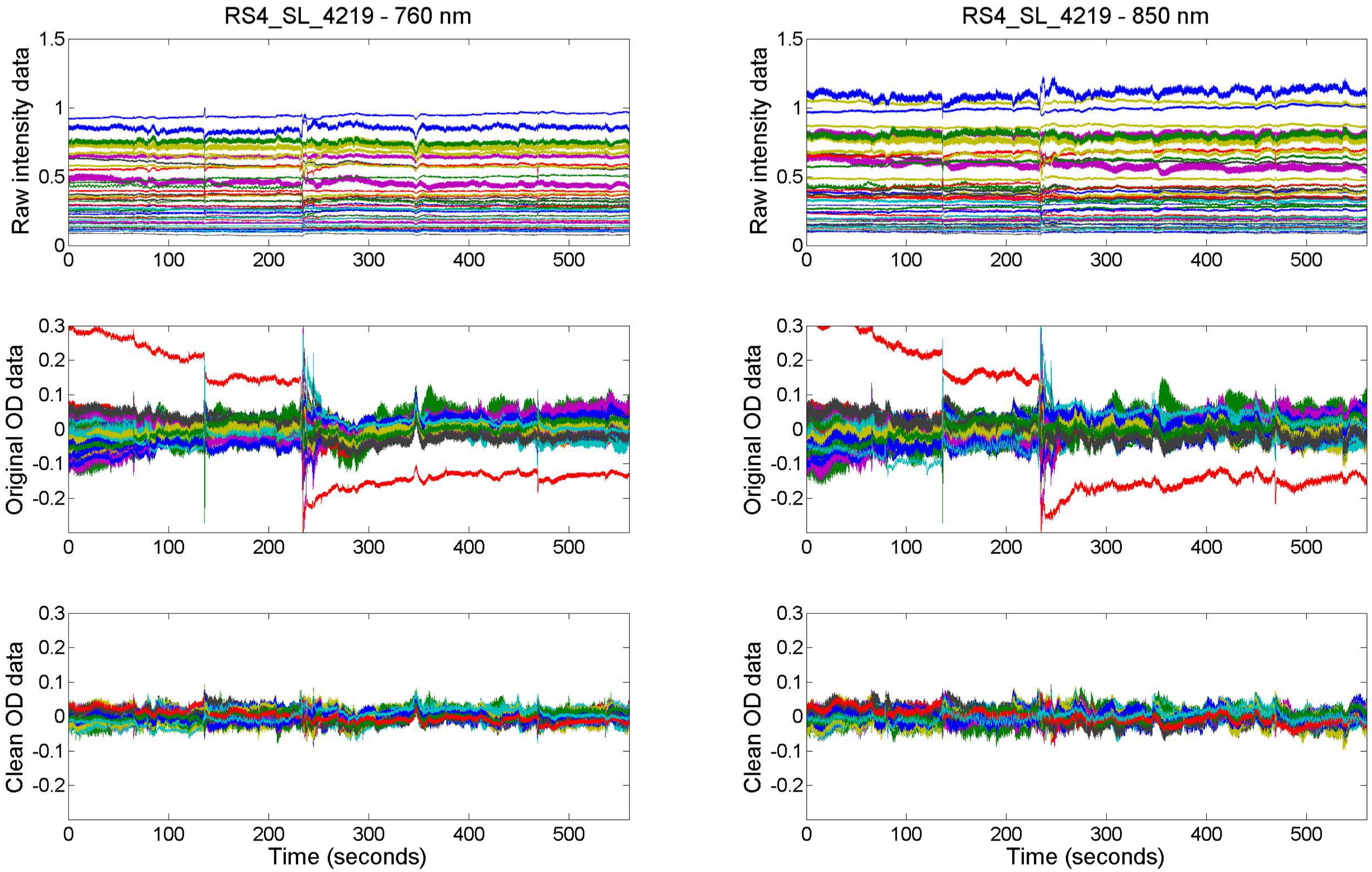
RS4_SL_4219 - 760 nm
RS4_SL_4219 - 850 nm
Raw intensity data
Original OD data
Clean OD data
Time (seconds)

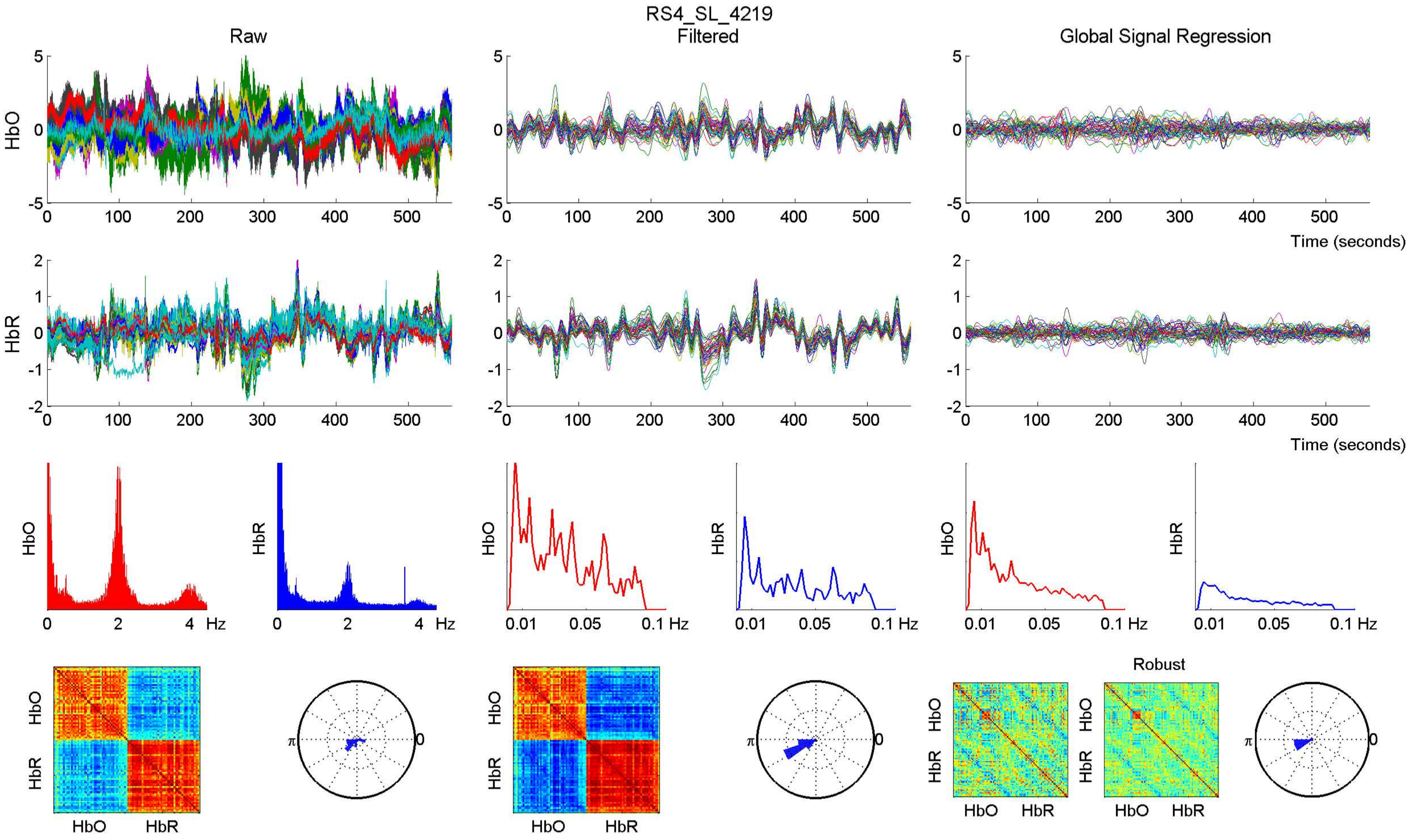

RS4_SL_4219
Raw
Filtered
Global Signal Regression
HbO
HbR
Time (seconds)
Hz
Robust

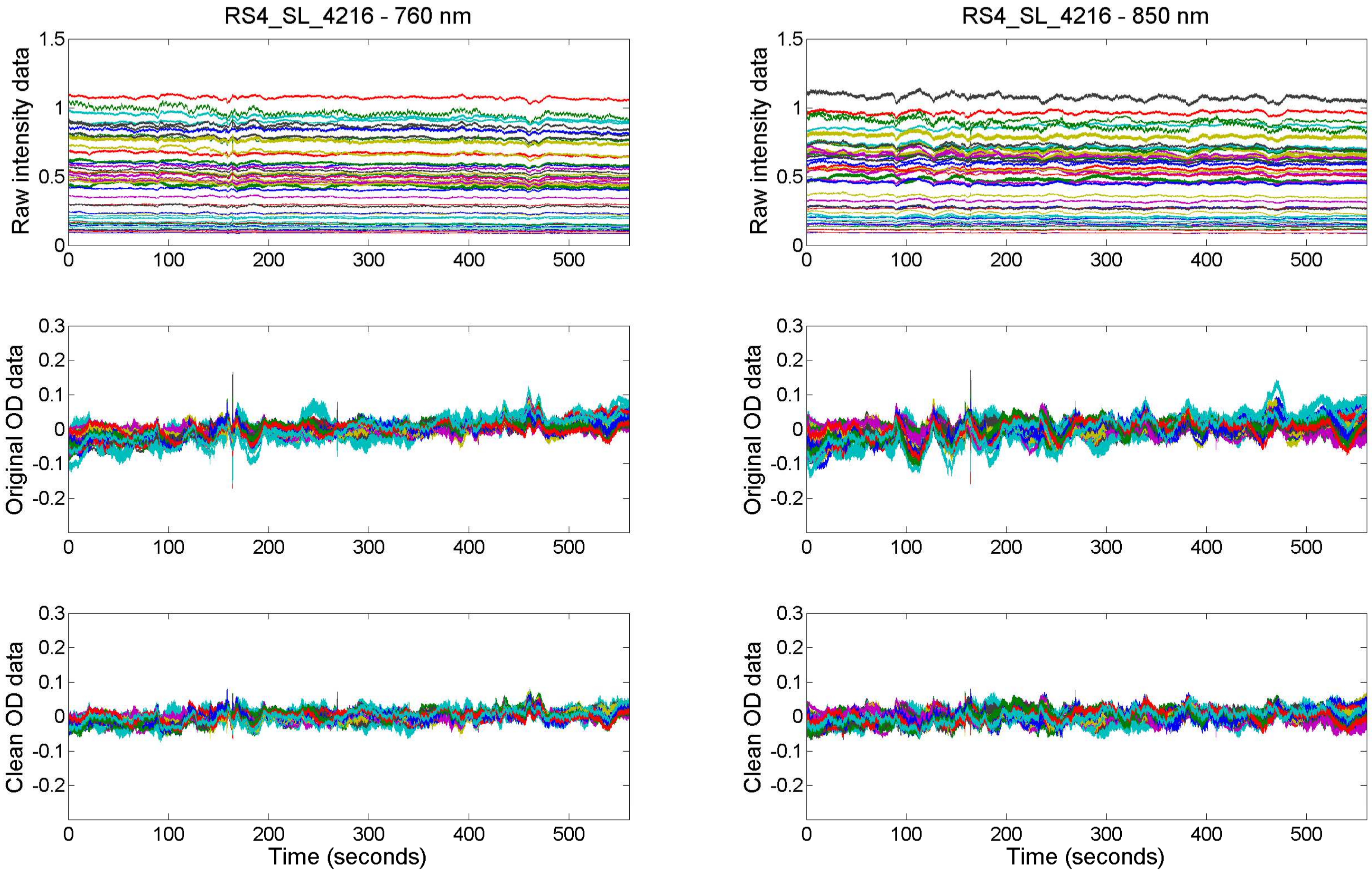

RS4_SL_4216 - 760 nm
RS4_SL_4216 - 850 nm
Raw intensity data
Original OD data
Clean OD data
Time (seconds)
1.5
1
0.5
0
0.3
0.2
0.1
-0.1
-0.2
100
200
300
400
500

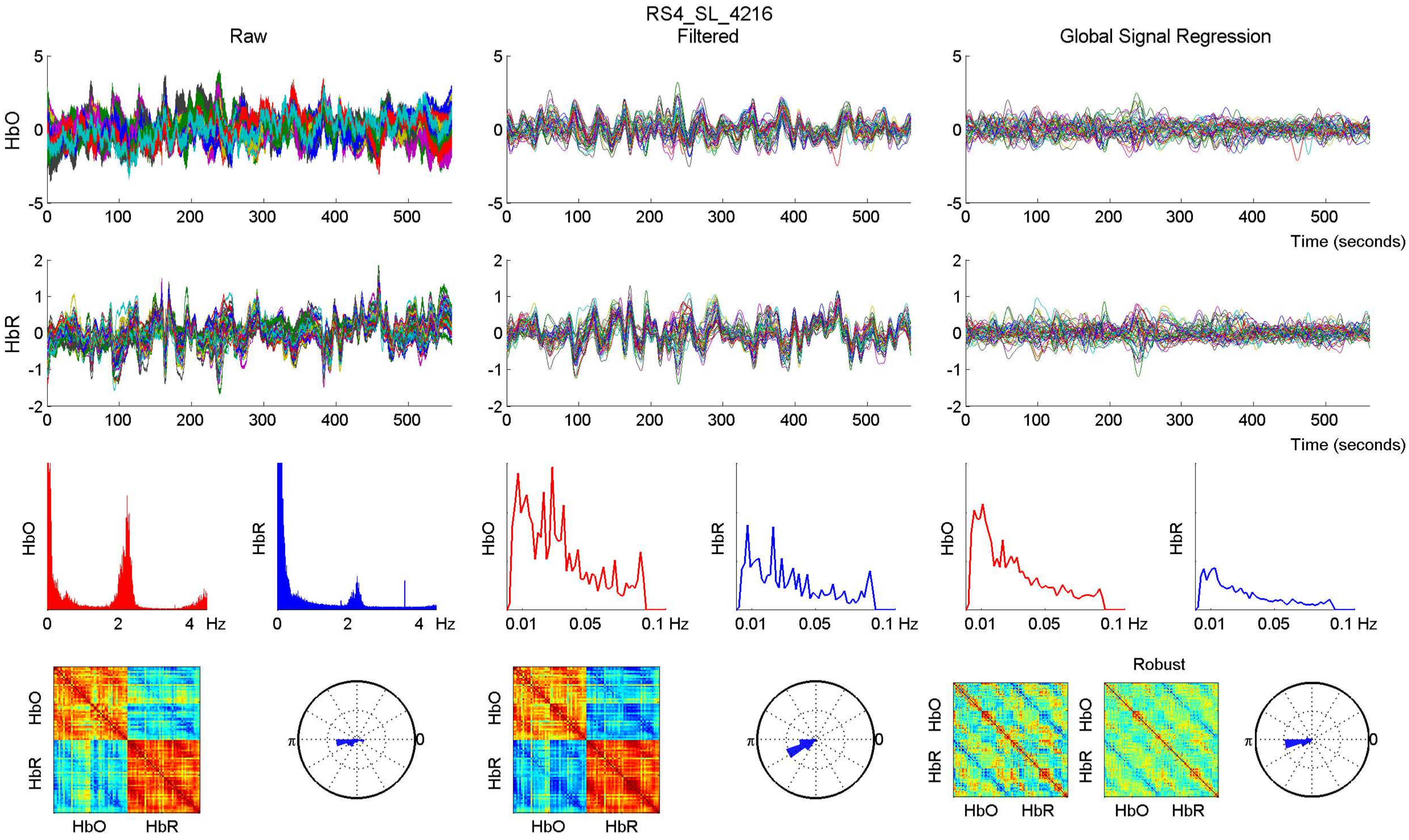

RS4_SL_4216
Raw
Filtered
Global Signal Regression
HbO
HbR
Time (seconds)
Hz
Robust

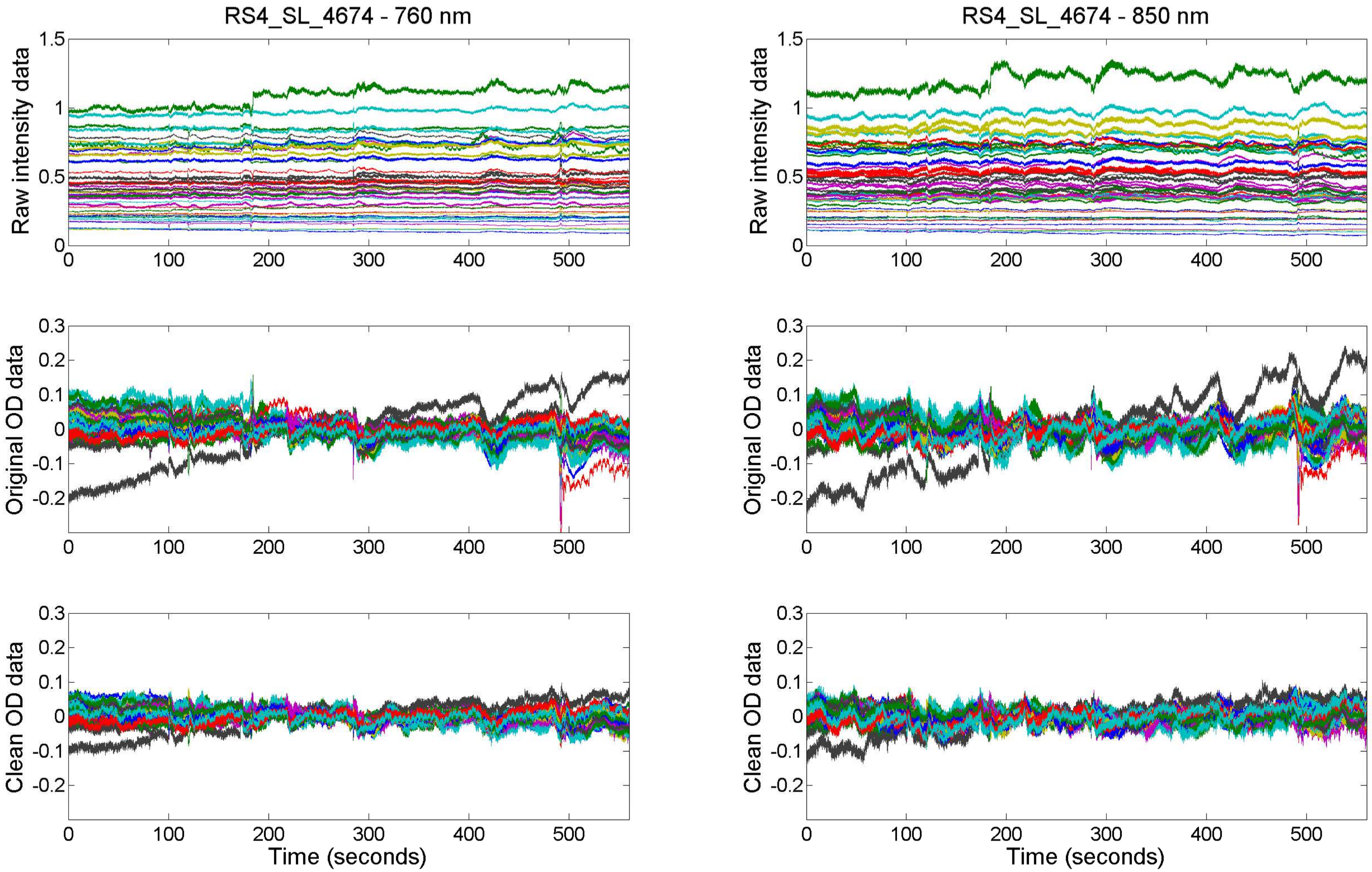

RS4_SL_4674 - 760 nm
RS4_SL_4674 - 850 nm
Raw intensity data
Original OD data
Clean OD data
Time (seconds)

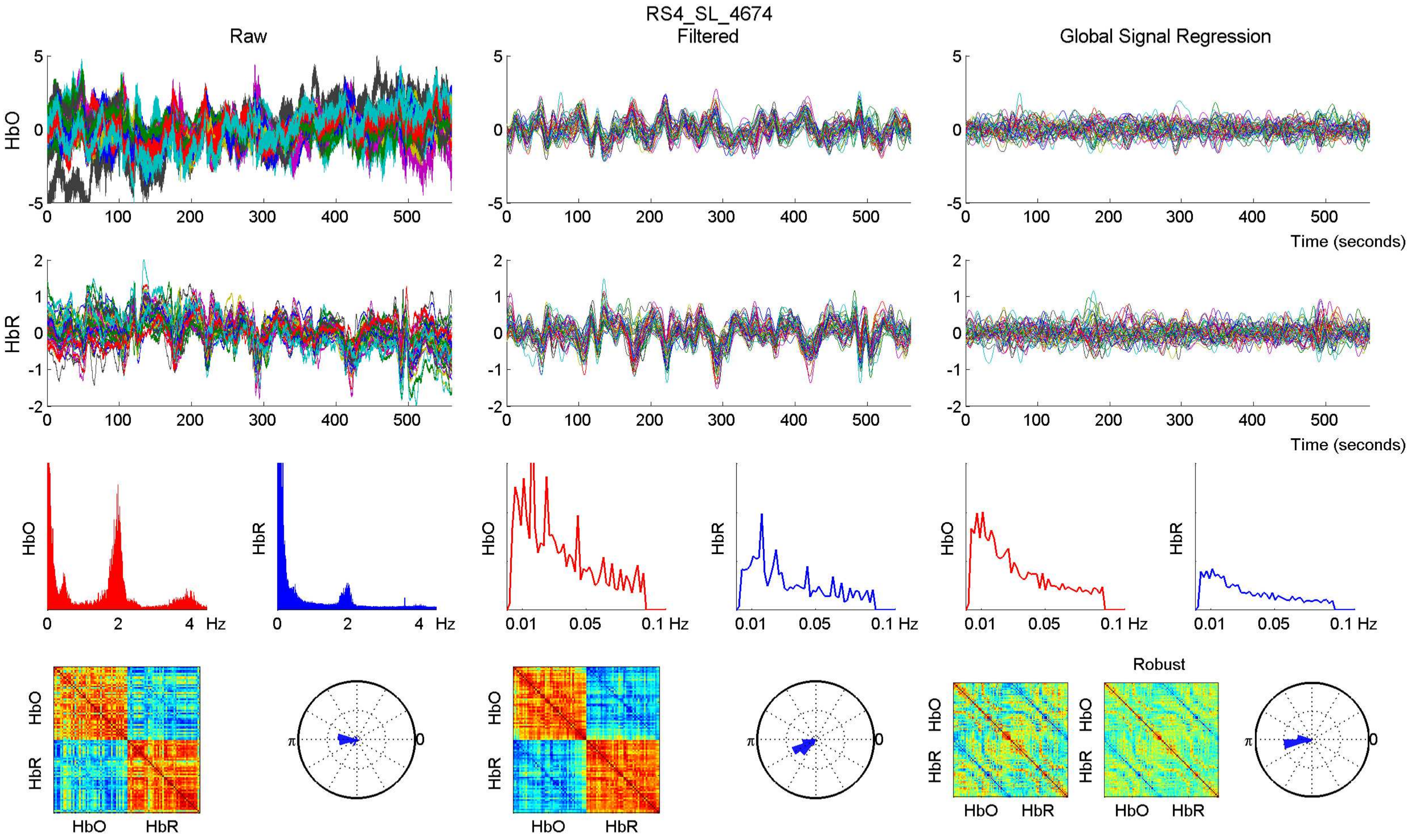

RS4_SL_4674
Raw
Filtered
Global Signal Regression
HbO
HbR
Time (seconds)
Hz
Robust
HbO
HbR

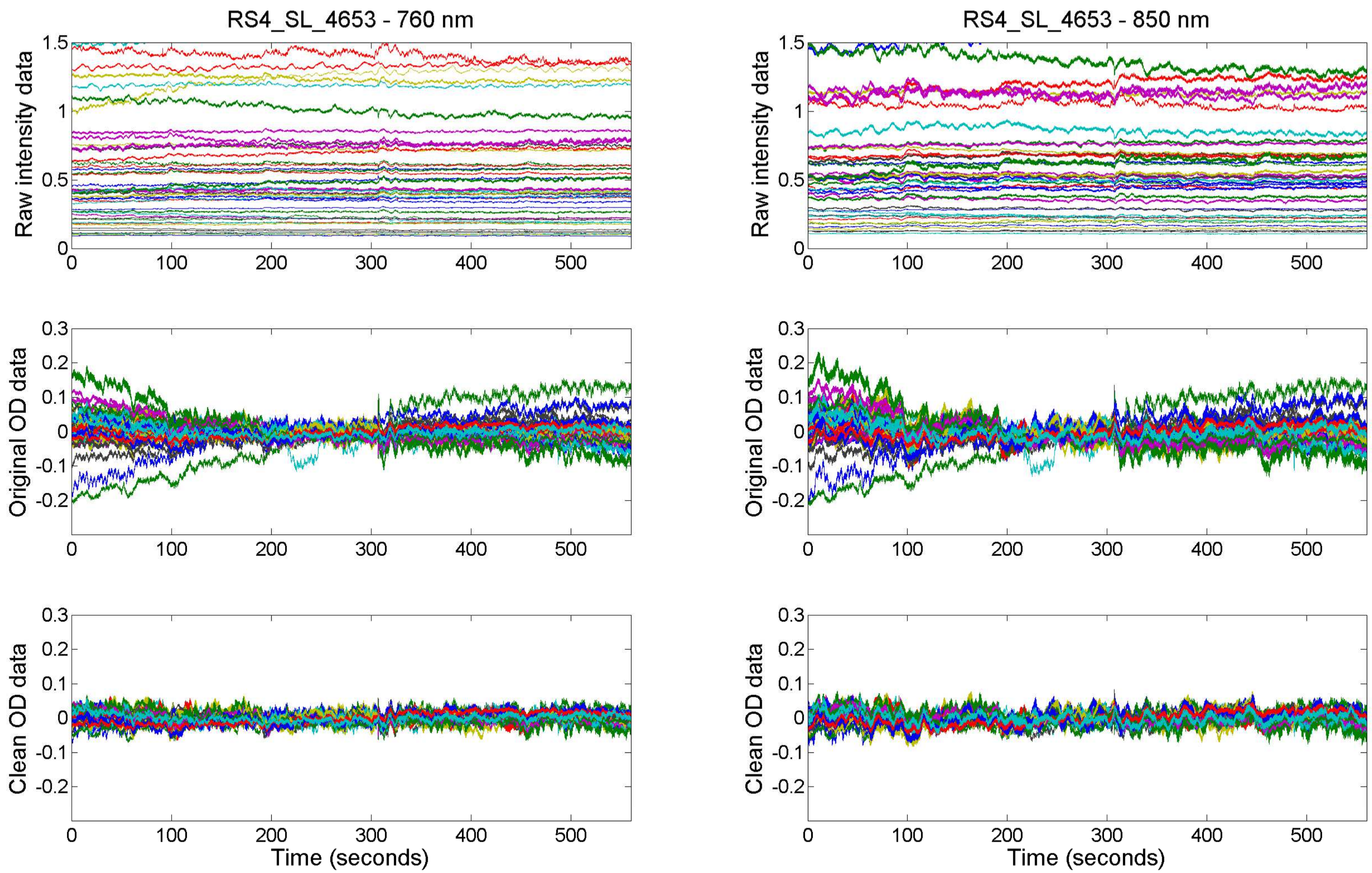
RS4_SL_4653 - 760 nm
RS4_SL_4653 - 850 nm
Raw intensity data
Original OD data
Clean OD data
Time (seconds)

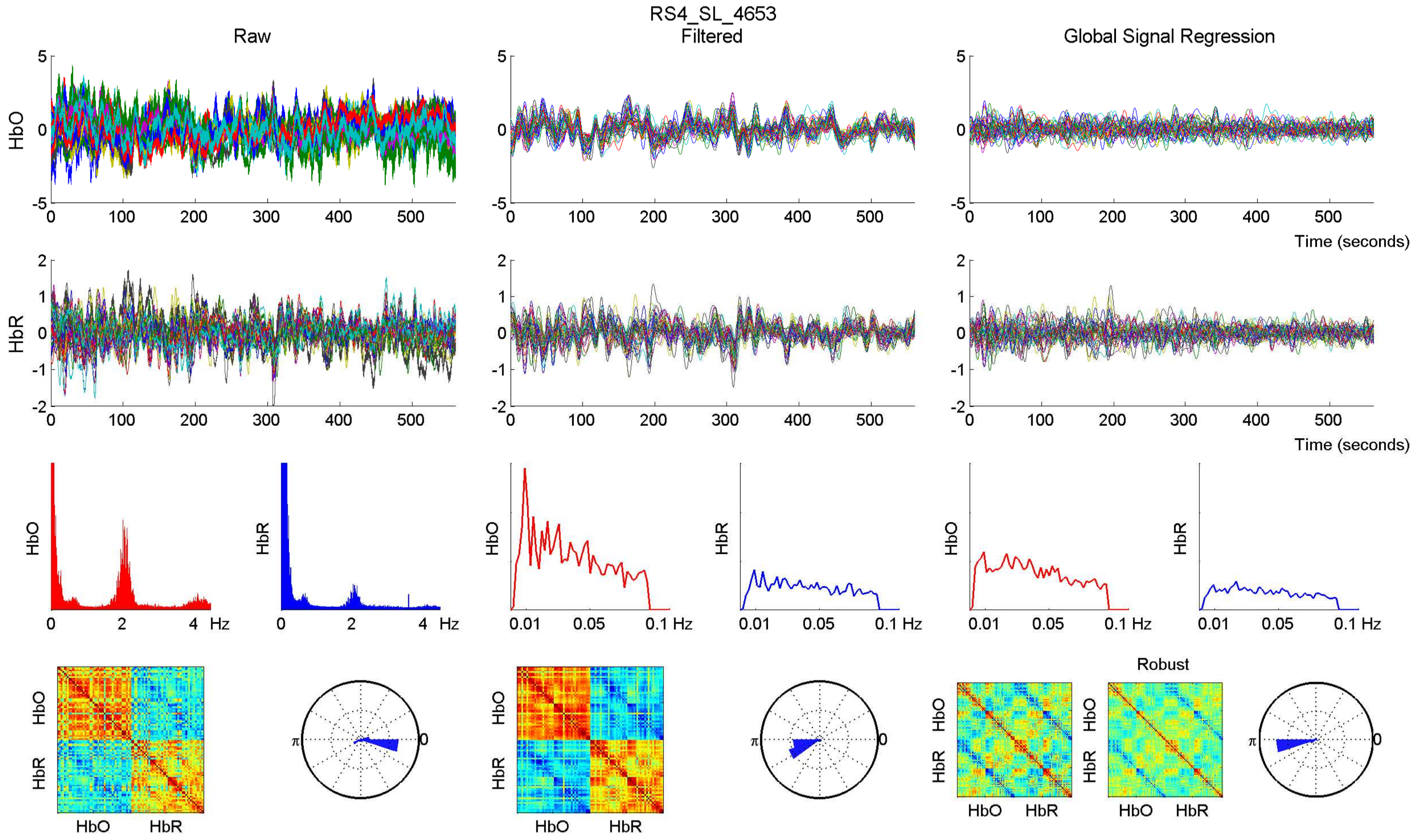

RS4_SL_4653
Raw
Filtered
Global Signal Regression
HbO
HbR
Time (seconds)
Hz
Robust
HbO
HbR

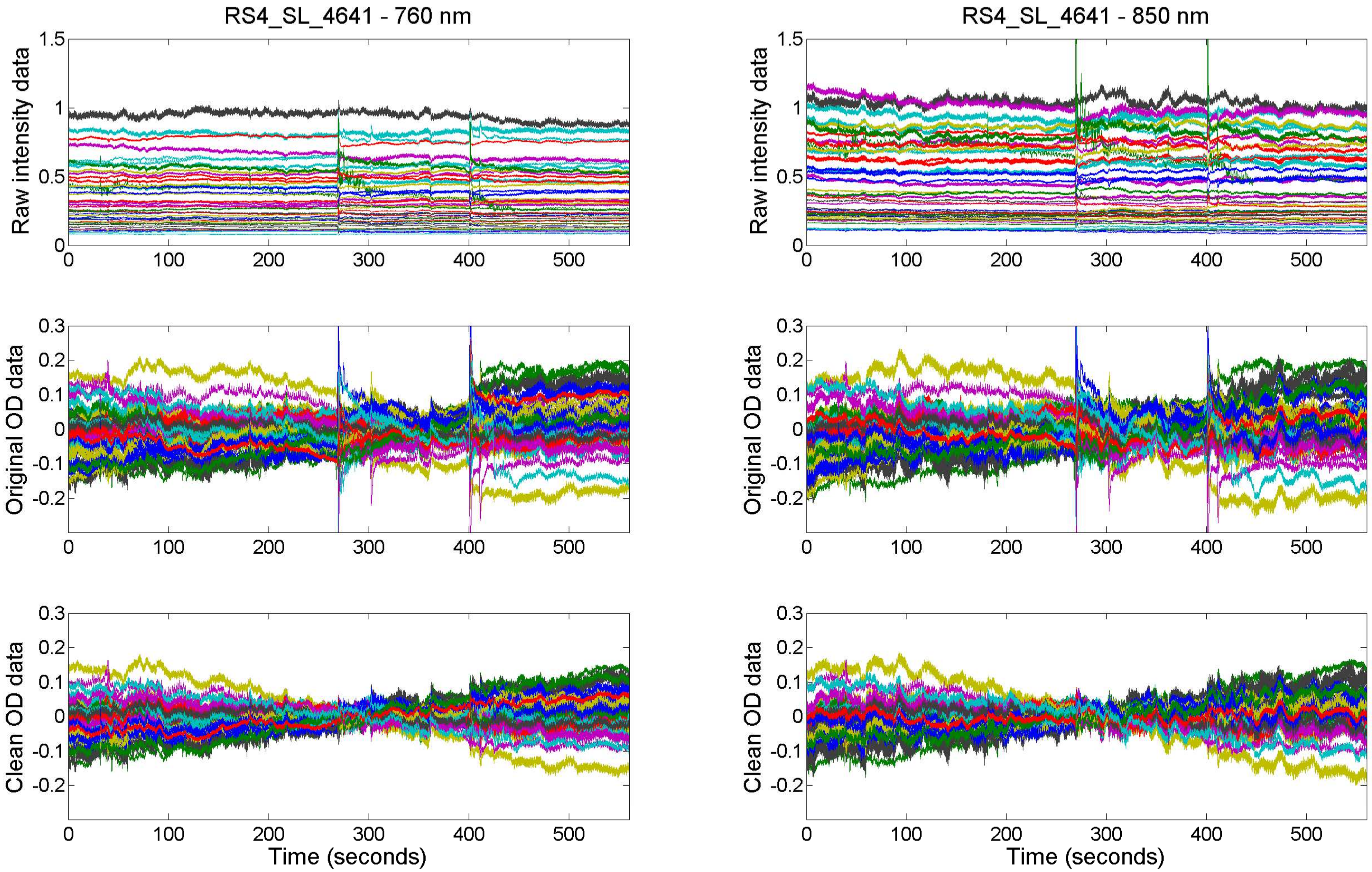

RS4_SL_4641 - 760 nm
RS4_SL_4641 - 850 nm
Raw intensity data
Original OD data
Clean OD data
Time (seconds)

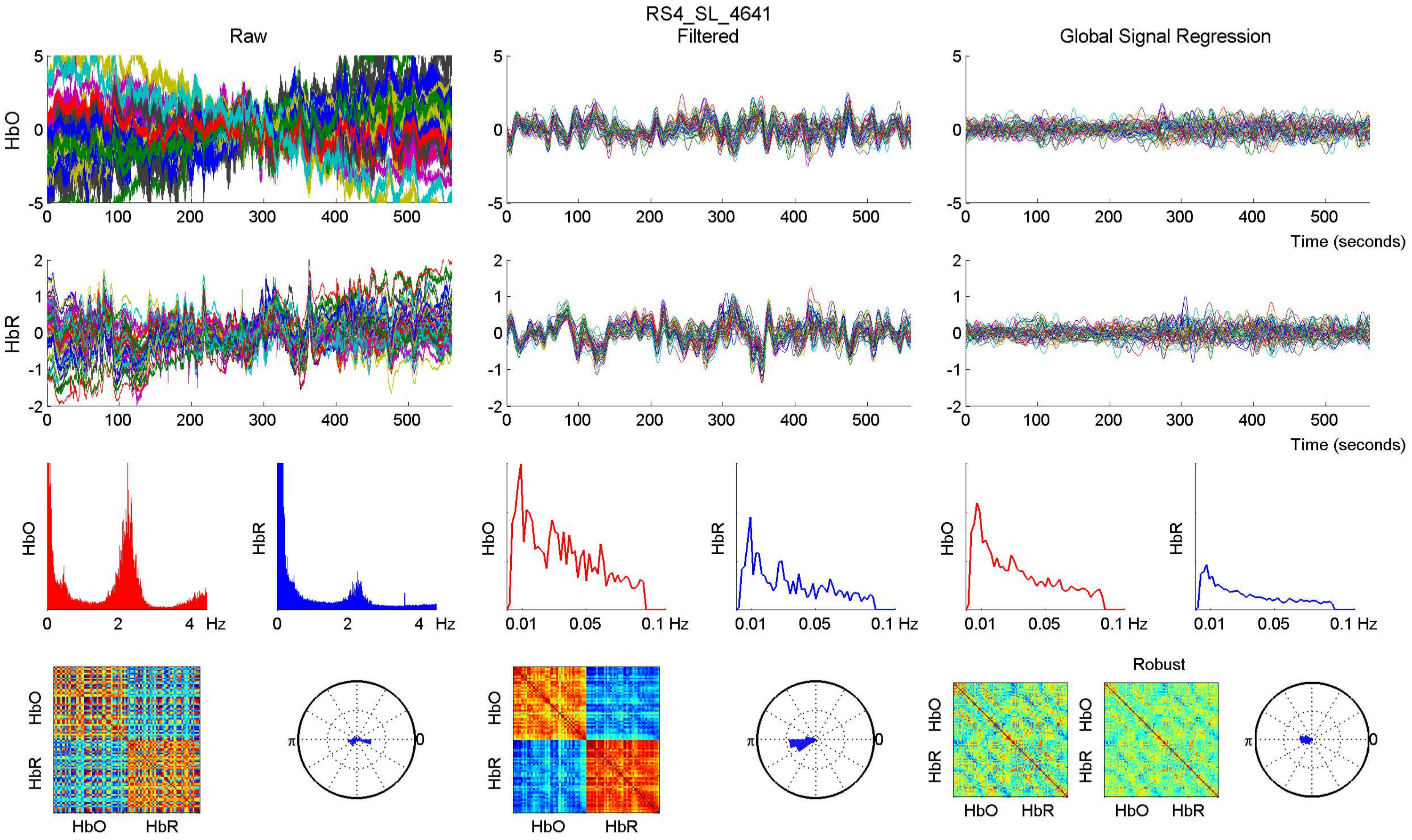

RS4_SL_4641
Raw
Filtered
Global Signal Regression
HbO
HbR
Time (seconds)
Hz
Robust

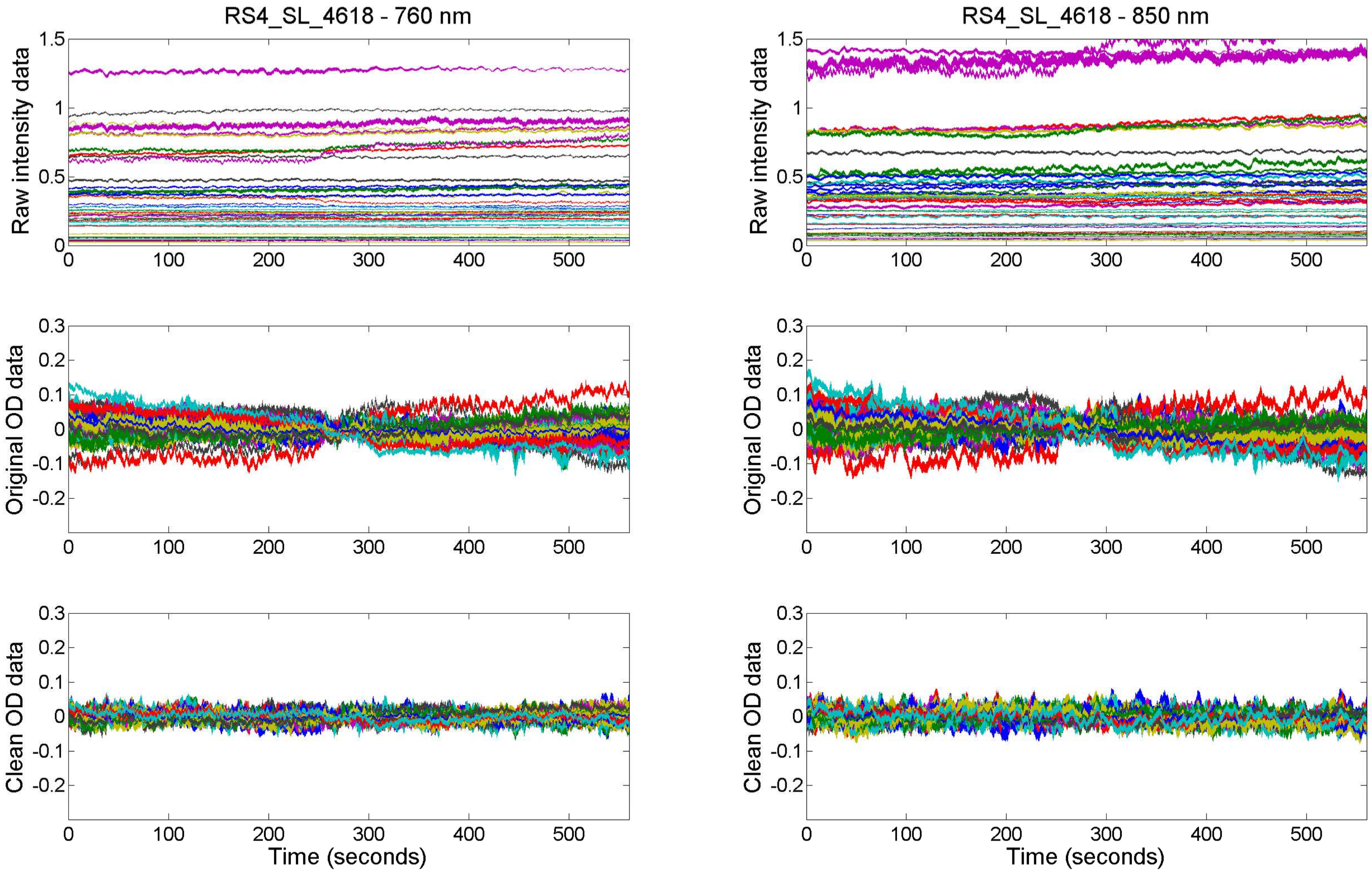

RS4_SL_4618 - 760 nm
RS4_SL_4618 - 850 nm
Raw intensity data
Original OD data
Clean OD data
Time (seconds)

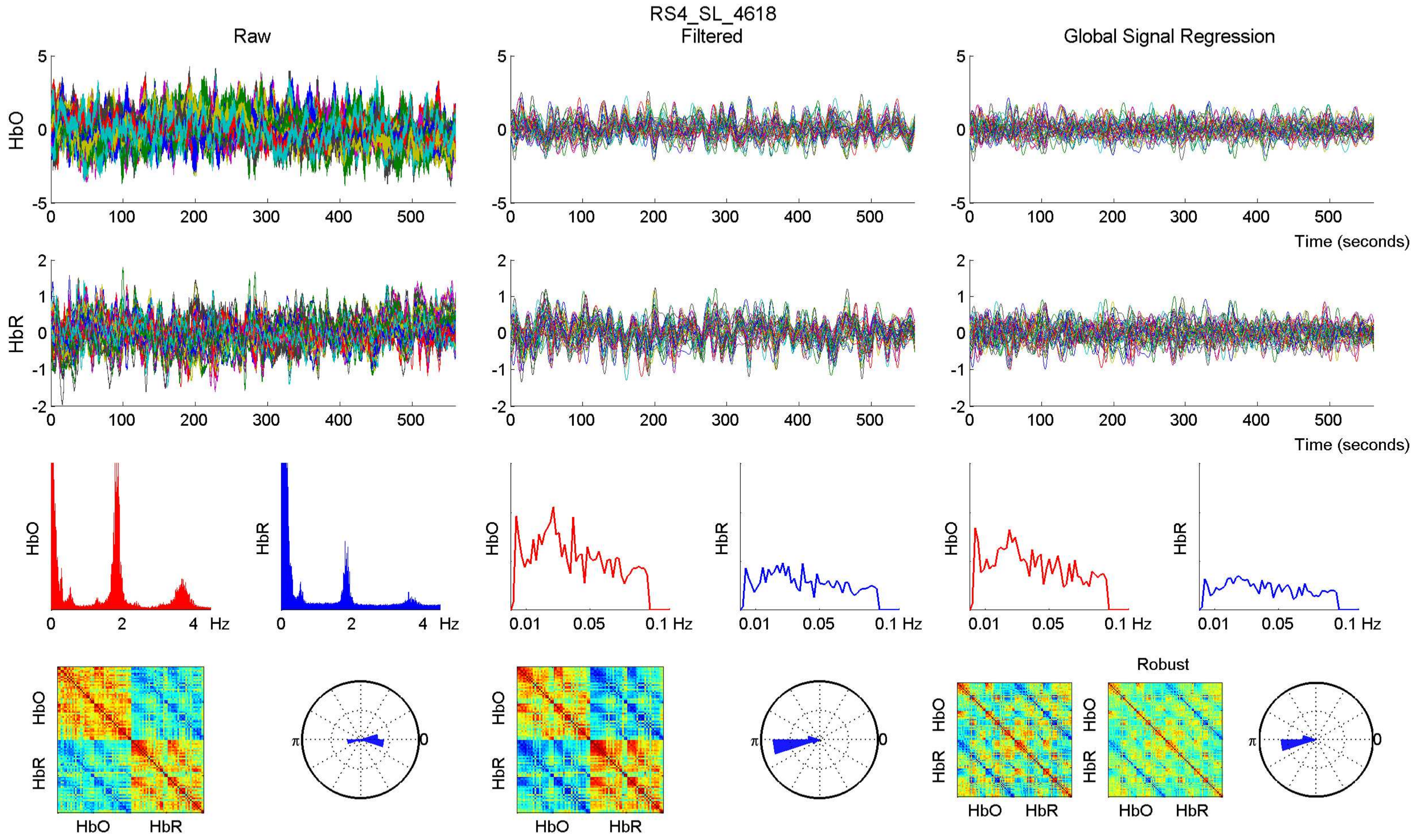

RS4_SL_4618
Raw
Filtered
Global Signal Regression
HbO
HbR
Time (seconds)
Hz
Robust

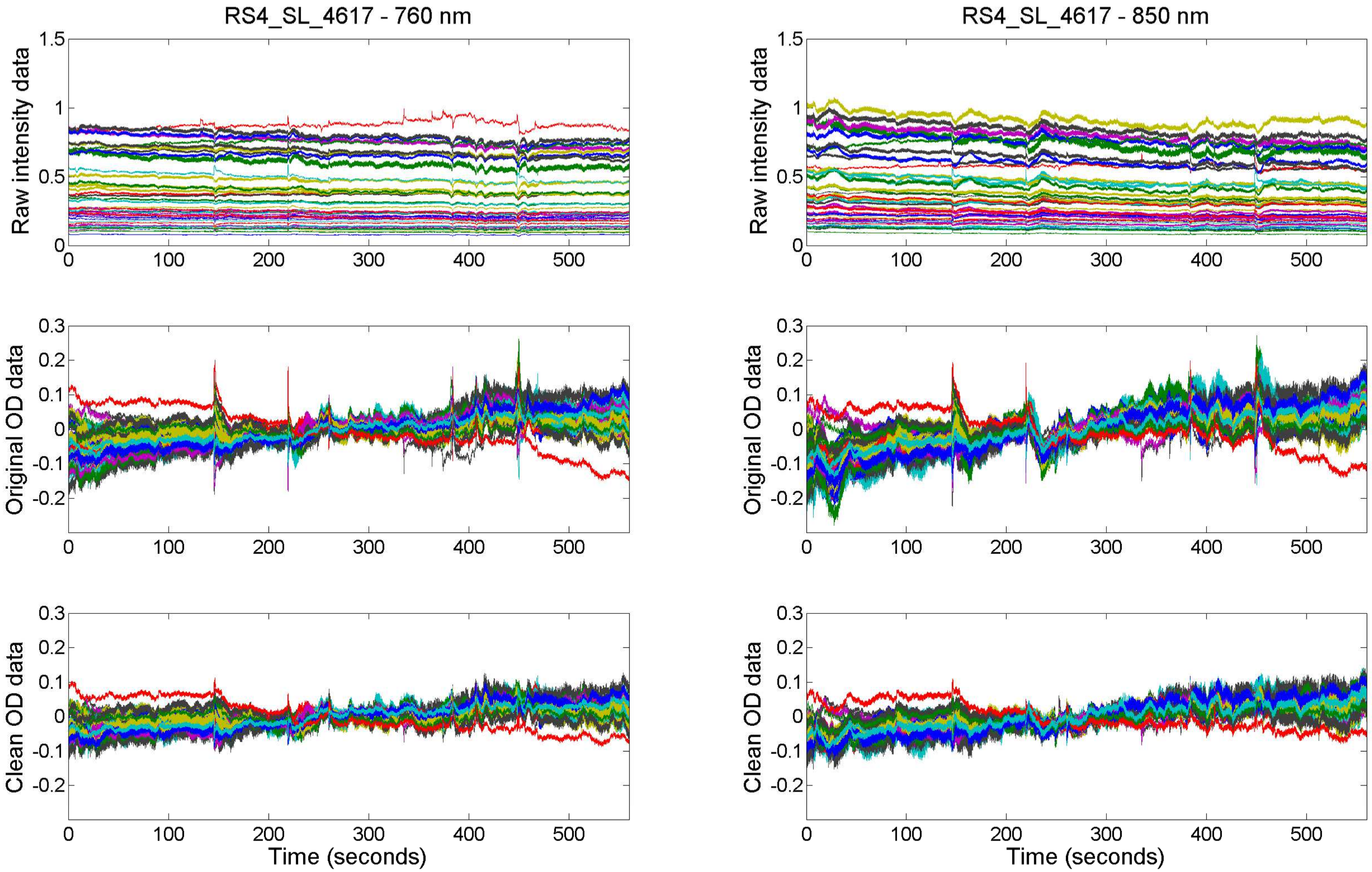

RS4_SL_4617 - 760 nm
RS4_SL_4617 - 850 nm
Raw intensity data
Original OD data
Clean OD data
Time (seconds)

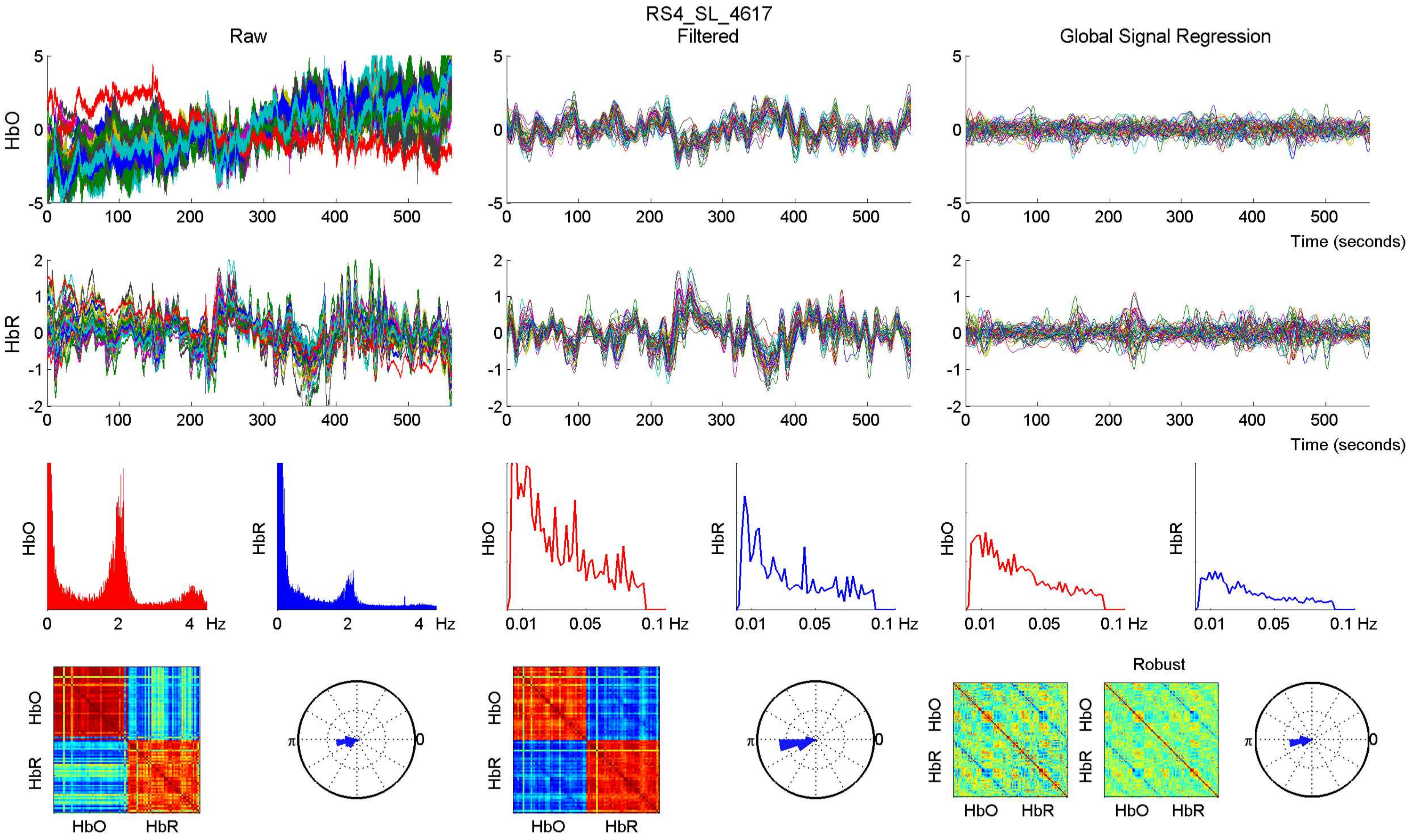

RS4_SL_4617
Raw
Filtered
Global Signal Regression
HbO
HbR
Time (seconds)
Hz
Robust
HbO
HbR

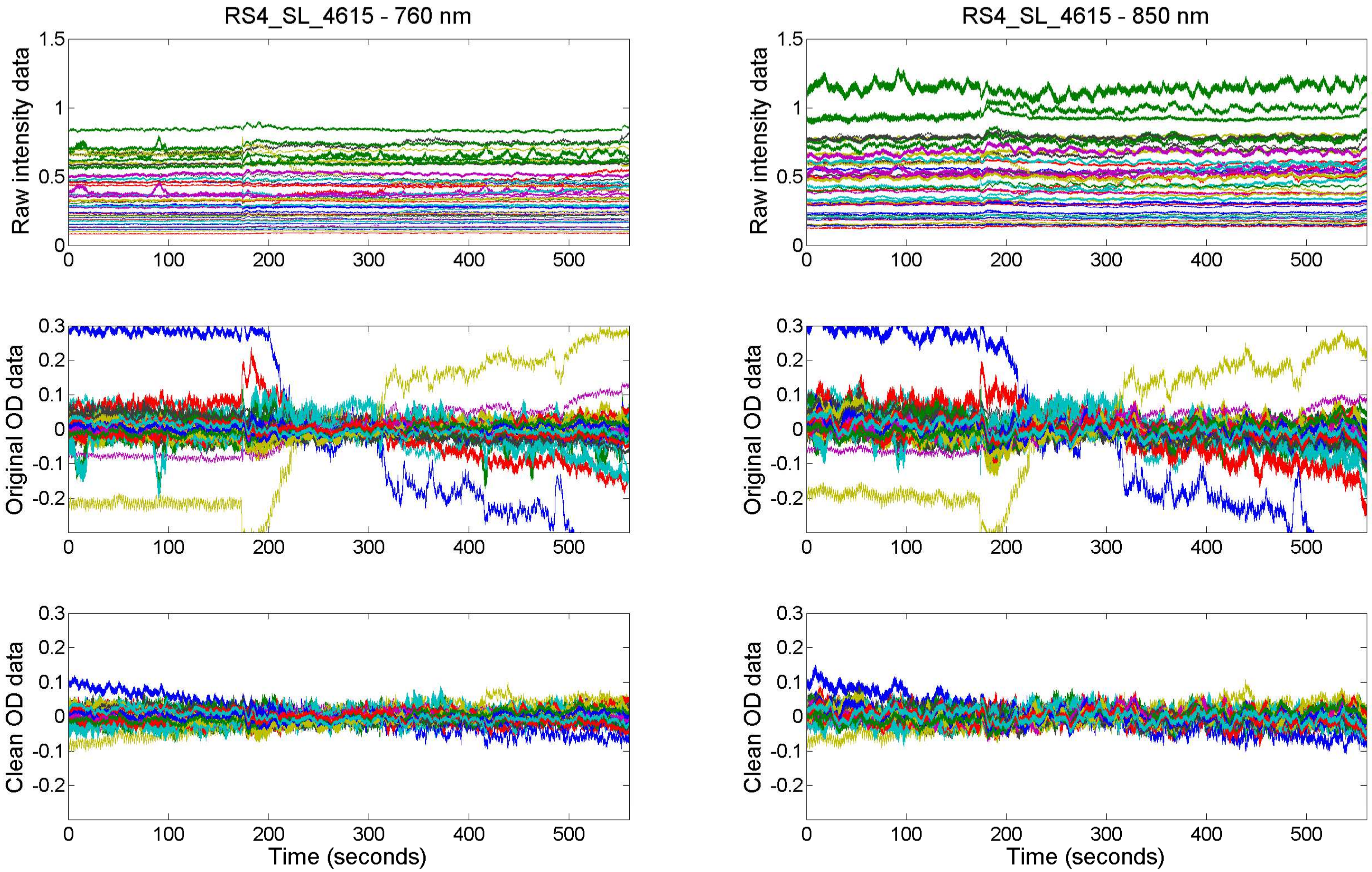

RS4_SL_4615 - 760 nm
RS4_SL_4615 - 850 nm
Raw intensity data
Original OD data
Clean OD data
Time (seconds)

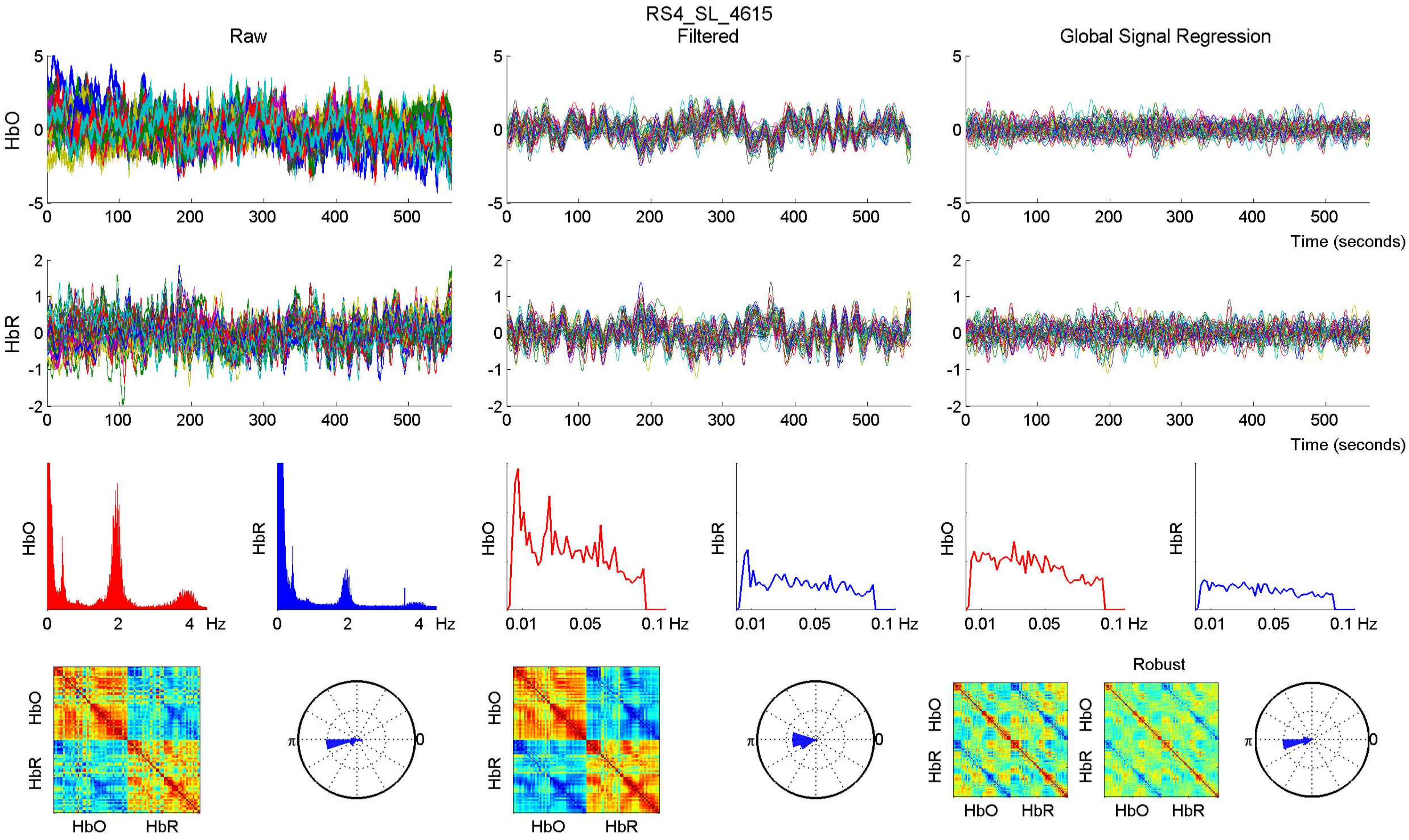

RS4_SL_4615
Raw
Filtered
Global Signal Regression
HbO
HbR
Time (seconds)
Hz
Robust
HbO
HbR

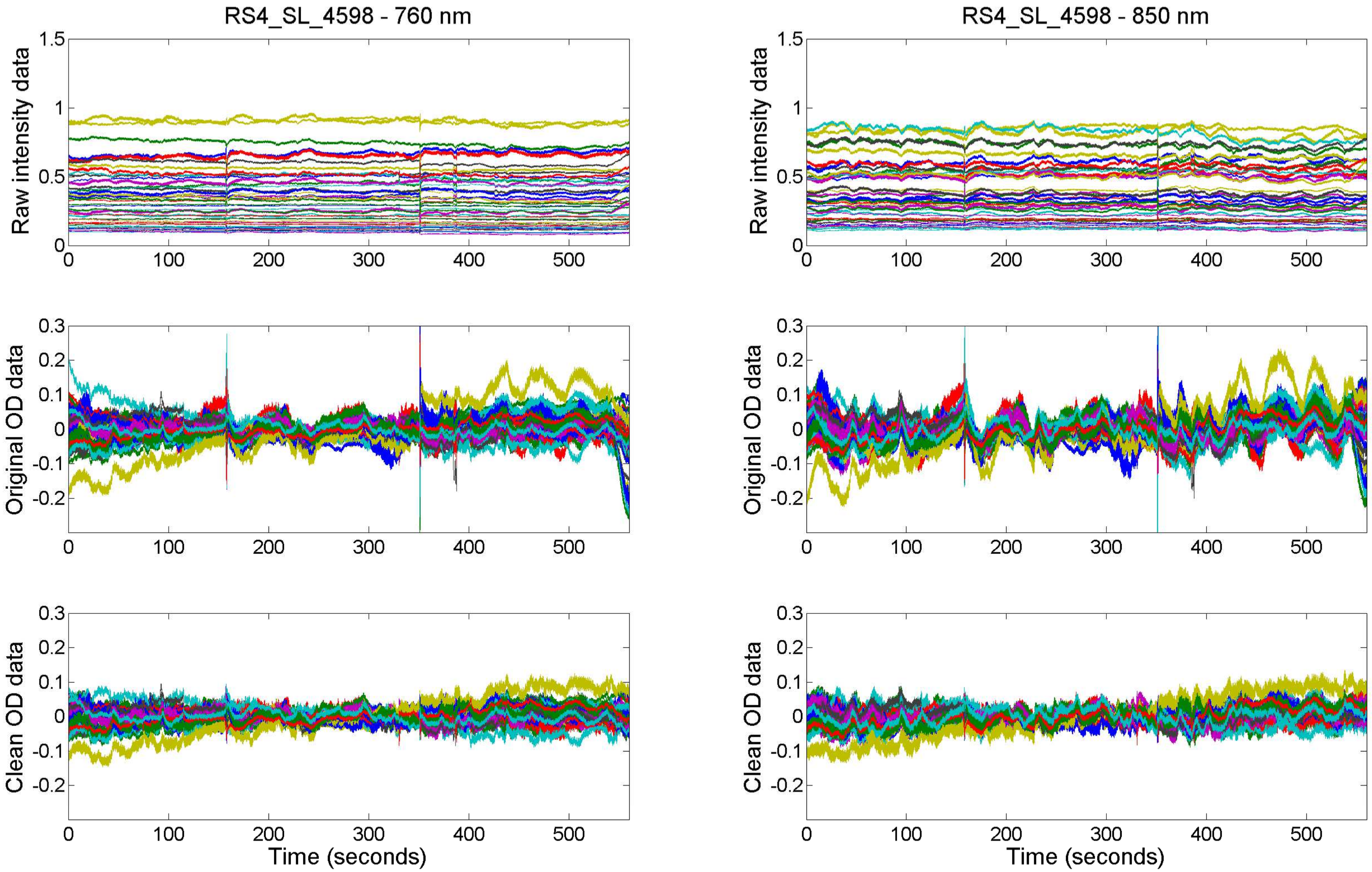

RS4_SL_4598 - 760 nm
RS4_SL_4598 - 850 nm
Raw intensity data
Original OD data
Clean OD data
Time (seconds)

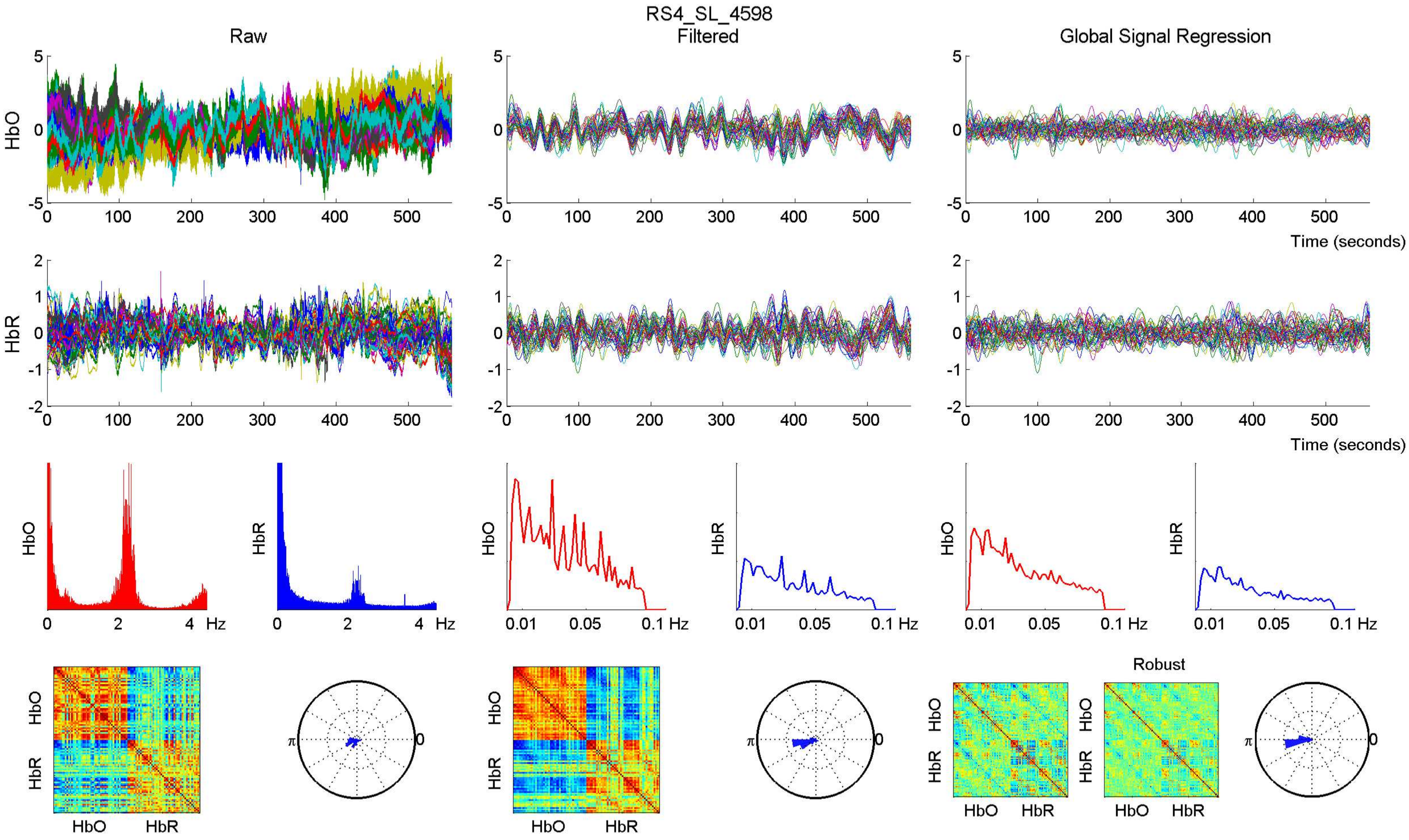

RS4_SL_4598
Raw
Filtered
Global Signal Regression
HbO
HbR
Time (seconds)
Hz
0.01
0.05
0.1 Hz
Robust
π
0

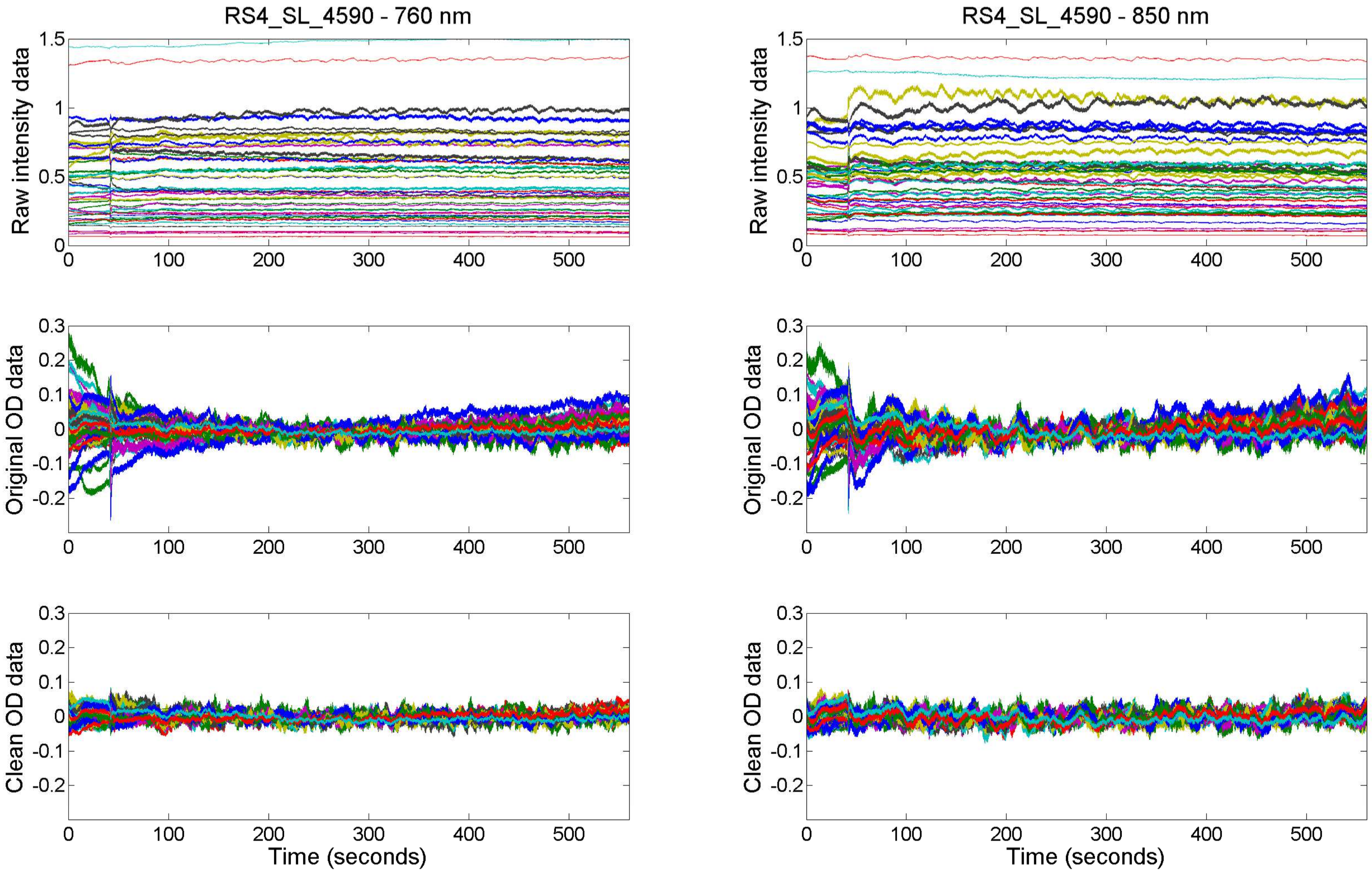

RS4_SL_4590 - 760 nm
RS4_SL_4590 - 850 nm
Raw intensity data
Original OD data
Clean OD data
Time (seconds)

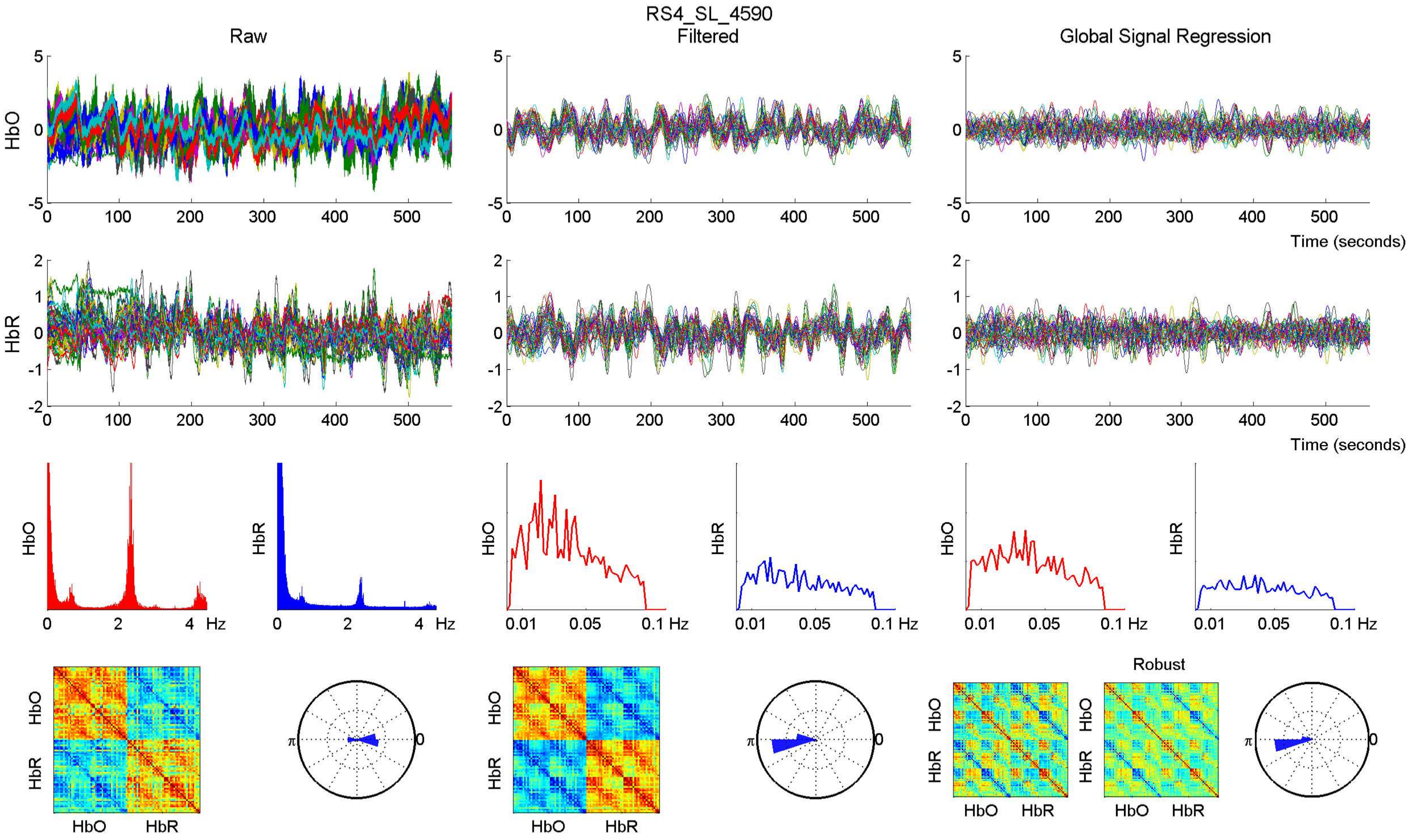

RS4_SL_4590
Raw
Filtered
Global Signal Regression
HbO
HbR
Time (seconds)
Hz
Robust

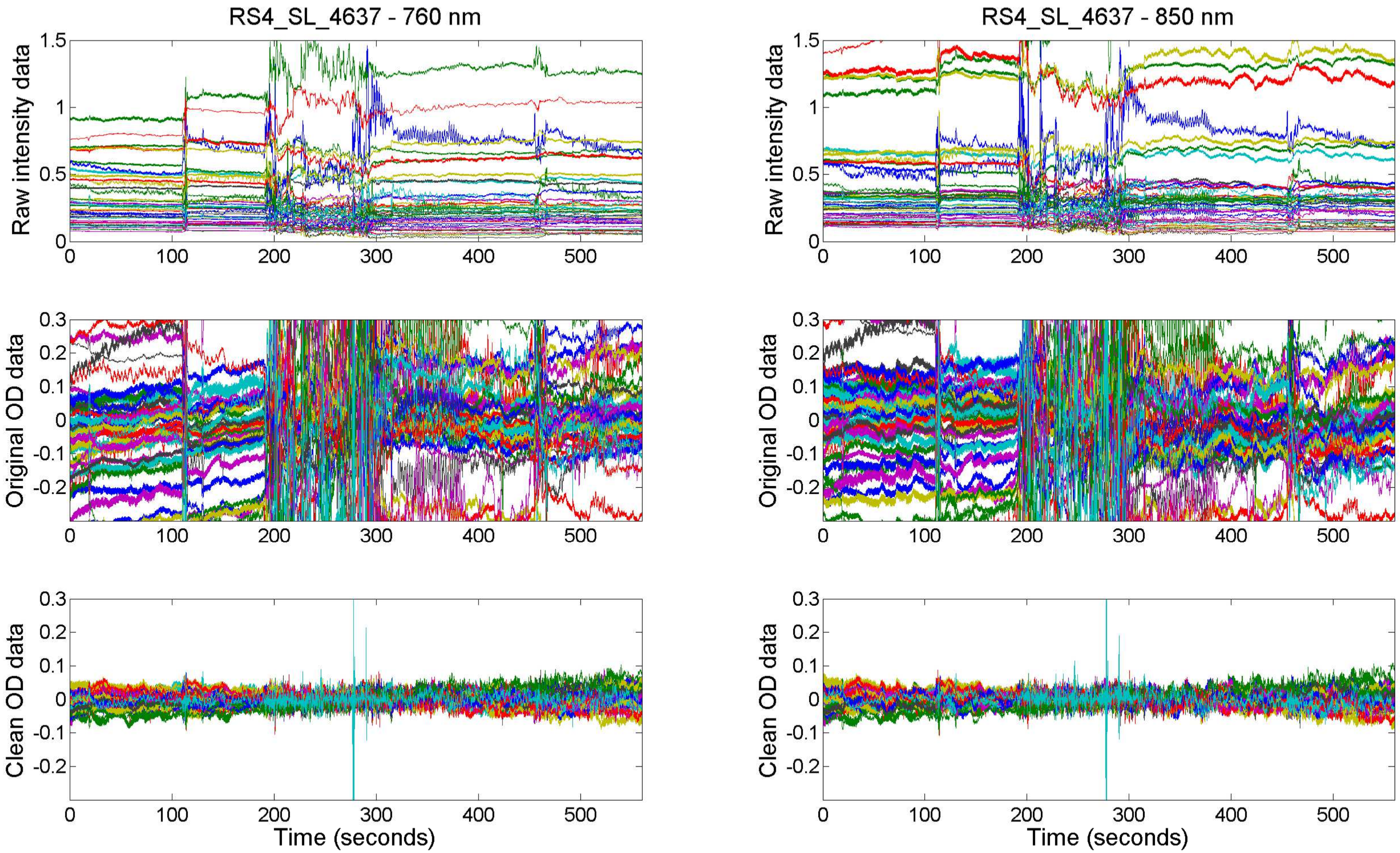

RS4_SL_4637 - 760 nm
RS4_SL_4637 - 850 nm
Raw intensity data
Original OD data
Clean OD data
Time (seconds)

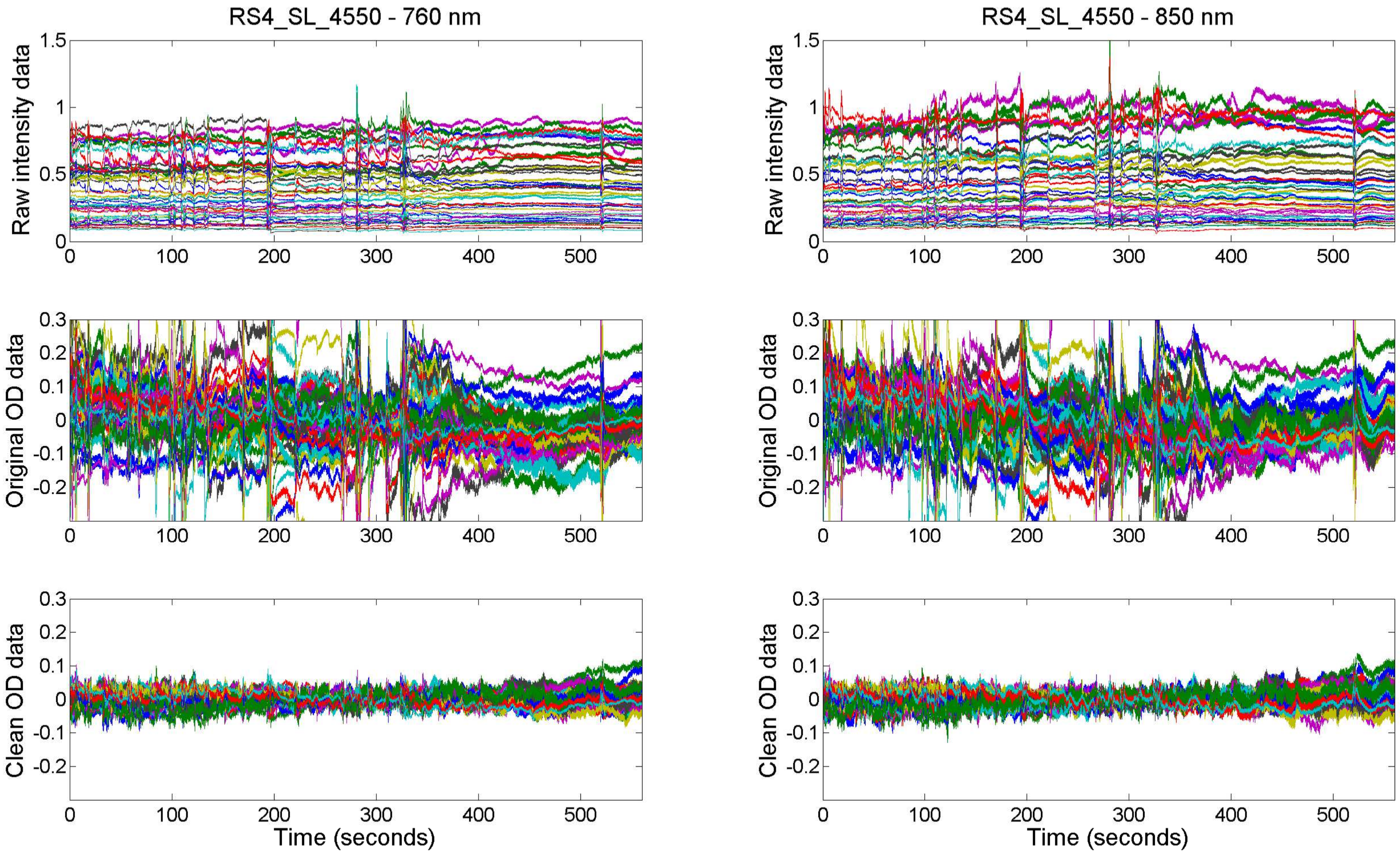

RS4_SL_4550 - 760 nm
RS4_SL_4550 - 850 nm
Raw intensity data
Original OD data
Clean OD data
Time (seconds)

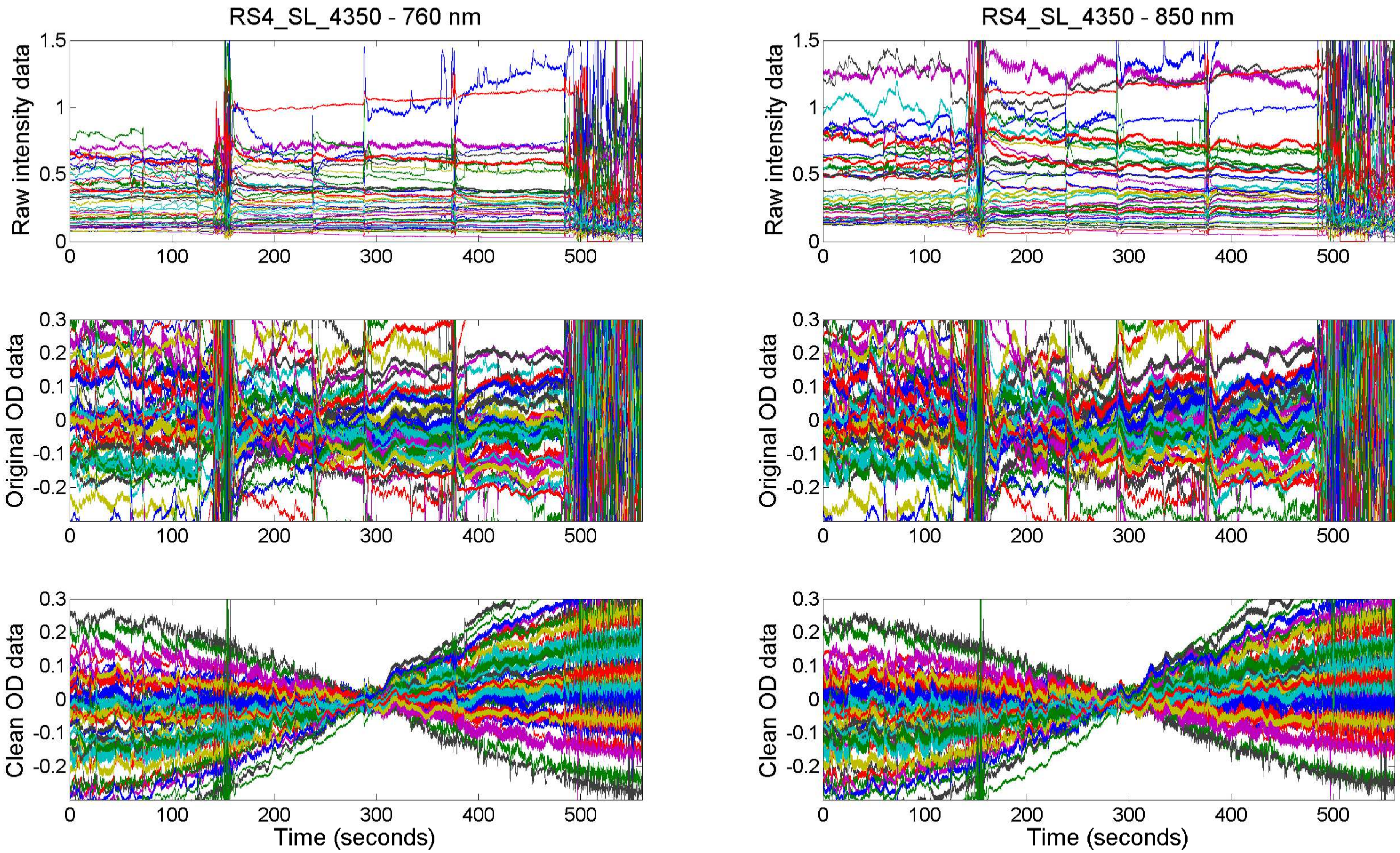
RS4_SL_4350 - 760 nm
RS4_SL_4350 - 850 nm
Raw intensity data
Original OD data
Clean OD data
Time (seconds)
Time (seconds)

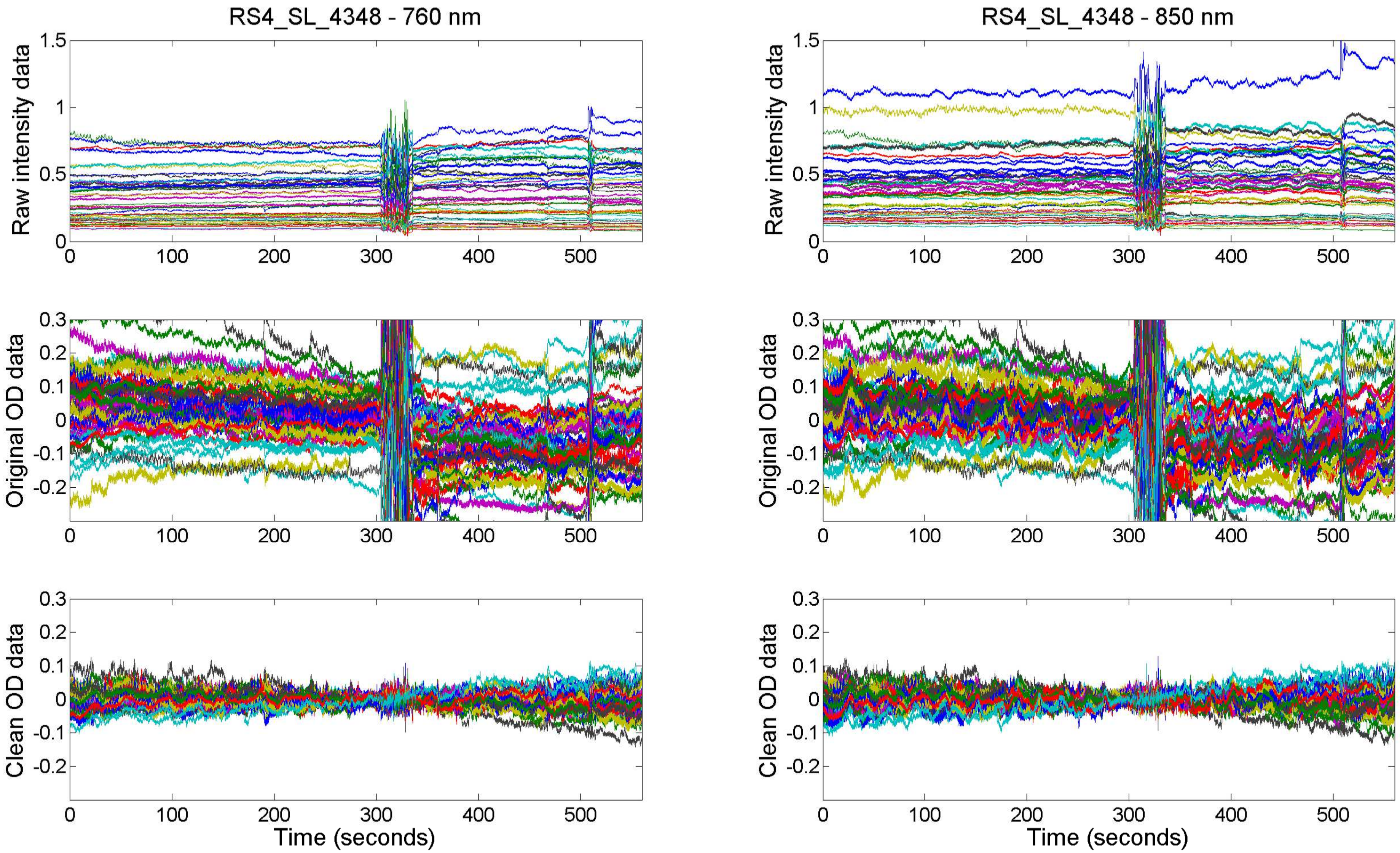
RS4_SL_4348 - 760 nm
RS4_SL_4348 - 850 nm
Raw intensity data
Original OD data
Clean OD data
Time (seconds)
Raw intensity data
Original OD data
Clean OD data
Time (seconds)

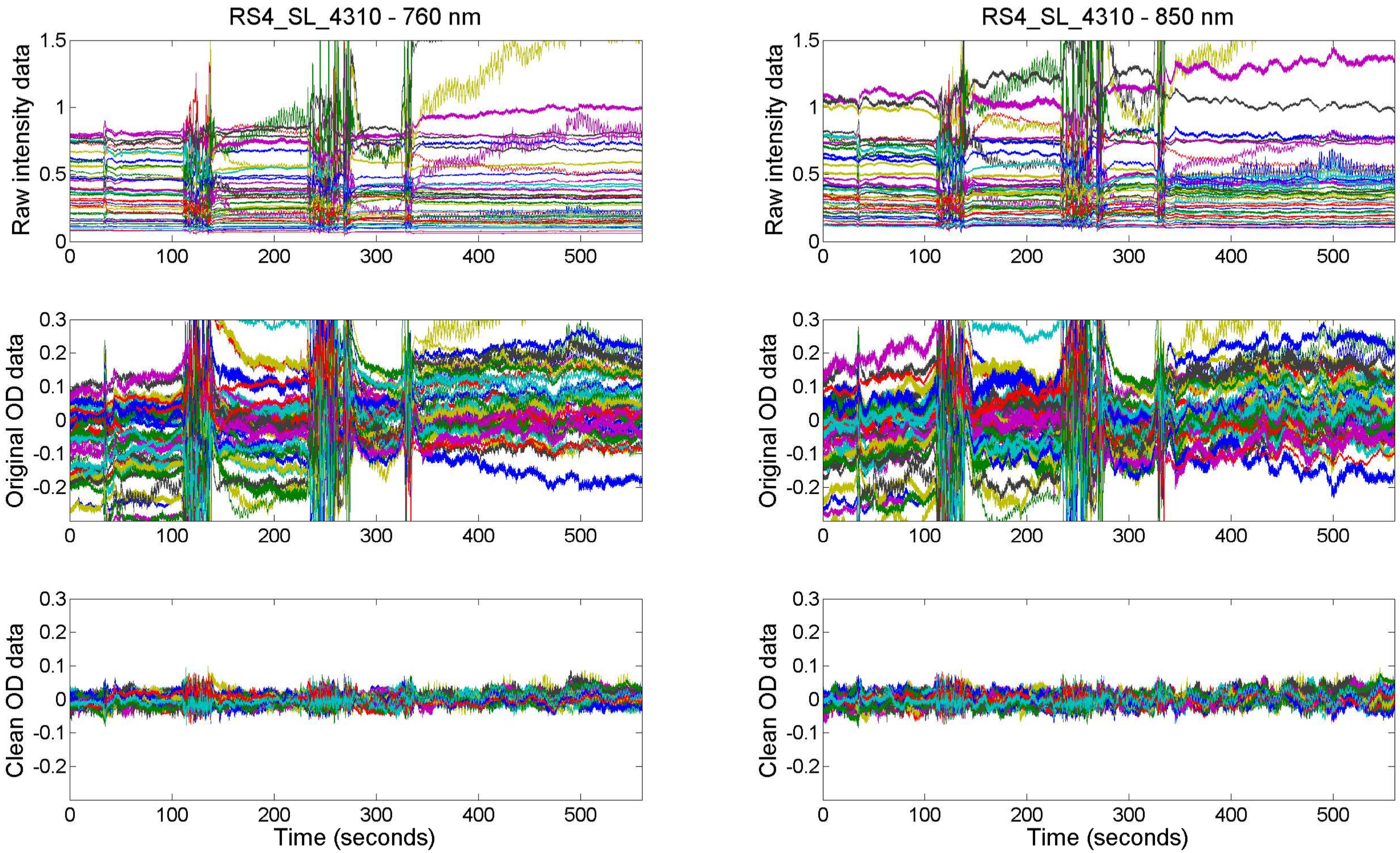

RS4_SL_4310 - 760 nm
RS4_SL_4310 - 850 nm
Raw intensity data
Original OD data
Clean OD data
Time (seconds)
